# Rapid detection of DNA and RNA shrimp viruses using CRISPR-based diagnostics

**DOI:** 10.1101/2022.12.14.520450

**Authors:** Samuel R. Major, Matthew J. Harke, Roberto Cruz-Flores, Arun K. Dhar, Andrea G. Bodnar, Shelly A. Wanamaker

## Abstract

Timely detection of persistent and emerging pathogens is critical to controlling disease spread, particularly in high-density populations with increased contact between individuals and limited-to-no ability to quarantine. Standard molecular diagnostic tests for surveying pathogenic microbes have provided the sensitivity needed for early detection, but lag in time-to-result leading to delayed action. On-site diagnostics alleviate this lag, but current technologies are less sensitive and adaptable than lab-based molecular methods. Towards the development of improved on-site diagnostics, we demonstrated the adaptability of a loop-mediated isothermal amplification-CRISPR coupled technology for detecting DNA and RNA viruses that have greatly impacted shrimp populations worldwide; White Spot Syndrome Virus and Taura Syndrome Virus. Both CRISPR-based fluorescent assays we developed showed similar sensitivity and accuracy for viral detection and load quantification to real-time PCR. Additionally, both assays specifically targeted their respective virus with no false positives detected in animals infected with other common pathogens or in certified specific pathogen-free animals.

**IMPORTANCE:** The Pacific white shrimp (*Penaeus vannamei*) is one of the most valuable aquaculture species in the world but has suffered major economic losses from outbreaks of White Spot Syndrome Virus and Taura Syndrome Virus. Rapid detection of these viruses can improve aquaculture practices by enabling more timely action to be taken to combat disease outbreaks. Highly sensitive, specific, and robust CRISPR-based diagnostic assays such as those developed here have the potential to revolutionize disease management in agriculture and aquaculture helping to promote global food security.

## INTRODUCTION

The ability to rapidly detect disease is a major challenge not just in human health but across all farmed and wild animal health monitoring. While methods for disease detection have advanced the field of molecular diagnostics to the global role it plays today, limitations posed by laboratory-based technologies hinder the ability to quickly respond to disease outbreaks. For example, culturing and molecular diagnostics using PCR-based methods provide the high sensitivity needed for early pathogen detection, but they require laboratory processing (e.g. sophisticated equipment, expensive reagents, and specialized training) leading to long turnaround times for results (several days) and delayed action for disease mitigation (1–4). Contrastingly, immunochromatographic antigen diagnostics are field-deployable, affordable and provide rapid results, but comparatively lack sensitivity and require more time for assay creation resulting in a longer time to respond to emerging pathogens (5). Alternative molecular diagnostics that require less sophisticated equipment like loop-mediated isothermal amplification (LAMP) have recently been developed, but their superior amplification efficiency can stochastically yield false positive results (6). In human health, CRISPR-based molecular diagnostics, such as the SHERLOCK (Specific High sensitivity Enzymatic Reporter UnLOCKing) technology, have emerged to overcome the limitations of PCR and antigen tests to enable rapid, sensitive, inexpensive, and accurate on-site detection of viruses and other pathogens (7). SHERLOCK combines the efficiency of isothermal amplification with the specificity of CRISPR detection for highly sensitive, specific, and robust diagnostics that can be used in low resource settings (8). Applications beyond human health in veterinary pathology and disease diagnostics in agriculture and aquaculture settings could have a tremendous impact on domestic and global food security.

The need for rapid, sensitive, and accurate diagnostics in aquaculture is imminent with production expected to double by 2050 to meet the demand of the growing population and being greatly impeded by infectious diseases (9, 10). For example, Pacific white shrimp (*Penaeus vannamei*) production, that accounts for ∼25% of all seafood products consumed annually, has suffered major economic losses due to infections caused by viruses like White Spot Syndrome Virus (WSSV) and Taura Syndrome Virus (TSV) (11–13). These diseases have resulted in up to 100% mortality in as few as seven days (14, 15). The DNA virus WSSV has been the leading cause of economic losses in shrimp aquaculture globally since its emergence in 1991, costing ∼$1 billion and reducing global shrimp production by ∼10% annually (11). TSV, a single stranded RNA virus, has caused mass mortality events on at least five continents since its emergence in 1991, with an estimated total loss of $2 billion from 1992-1996 (16–18). Although Taura syndrome caused by TSV is no longer causing economic losses to the extent it has in the past, with the current availability and farming of TSV-tolerant shrimp lines (19), WSSV-tolerant shrimp are still under development and are not commercially available (20). The domestic development and international import of specific pathogen-free (SPF) shrimp stocks has mitigated some disease outbreaks. However, even with routine rigorous screening disease continues to be a major threat. For instance, shrimp stocks in Brazil and Colombia were reported to be contaminated from imported SPF shrimp with underlying TSV infection that went undetected (21). The more recently documented spread of WSSV in Australia, India, and Ecuador has resulted in the virus being labeled as the most devasting pathogen in the aquacultural history of shrimp farming (22–24). To ensure globally distributed stocks are disease-free and to mitigate production losses for improved food security, it is crucial that rapid, accurate, and sensitive diagnostics are available to shrimp farmers (25, 26).

To address the need for rapid diagnostics in shrimp aquaculture, SHERLOCK technology has recently been applied to the detection of common and deadly shrimp viruses but remains to be optimized. For example, a CRISPR-based diagnostic was developed to detect *Vibrio spp.* that cause acute hepatopancreatic necrosis disease in shrimp but requires a two-step process of first amplifying pathogen DNA with recombinase polymerase amplification (RPA) and then detecting the amplicons with Cas12a in a separate reaction to detect down to 1000 copies per reaction (20 copies per µL in a 50 µL reaction) in 2 hours (27). Our lab developed a different variation of a CRISPR-based diagnostic to detect WSSV in shrimp tissue that uses RPA to amplify WSSV DNA, RNA polymerase to transcribe amplicons to RNA, and Cas13a to detect transcribed amplicons (SHERLOCKv1) all in one tube (one-pot format) (28). This one-pot format reduces time-to-result and was able to detect 10 viral copies per reaction in 60 minutes. Other reports have found LAMP to be superior to RPA in speed and sensitivity (29, 30), and a second-generation SHERLOCK assay that couples LAMP with a heat tolerant Cas enzyme, Cas12b, showed highly sensitive, rapid, and robust detection of viral RNA from SARS-CoV-2 (31).

Building upon these successes, and incorporating second generation SHERLOCK technology, we demonstrate the adaptability of this technology for both DNA and RNA viral targets and present two second-generation SHERLOCK (SHERLOCKv2) assays that detect WSSV and TSV. These robust, sensitive, and specific lab-based fluorescent assays can detect viral target in shrimp tissue in 30 minutes without the need for thermocycling. They also enable field-deployable conversion for improved diagnostic capabilities of viral outbreaks in aquaculture settings. This demonstration of assay adaptation for diverse shrimp diseases paves the way for next generation assay development in veterinary pathology and animal diagnostics for improved disease management in aquaculture and agriculture.

## RESULTS

### Diagnostic assay development

Combining the robustness of LAMP with the specificity of CRISPR and detection of Cas12b collateral cleavage activity, we developed sensitive and specific one-pot SHERLOCKv2 assays for the detection of DNA and RNA viruses that negatively impact shrimp aquaculture worldwide, WSSV and TSV (**Figure 1**). In an initial screen to identify an optimal LAMP primer set targeting a conserved TSV genome region, we compared the amplification activity of three primer sets as well as a previously published LAMP primer set (32). We found the previously published set targeting the capsid protein gene between VP1 and VP3 domains showed the fastest, most sensitive detection of TSV and selected this set for the SHERLOCKv2 assay (**Figure 2a and c**). In designing the WSSV assay, the highly conserved viral protein 28 (VP28) was targeted to limit the ability of WSSV strain to influence detection. Of the four LAMP primer sets designed and screened, one set (VP28.31) showed superior speed (**Figure 2b and d**). The addition of poly-T linkers between the two annealing sites within the FIP and BIP primers increased both the speed and sensitivity of the primer set down to 100 copies (**Figure S1**). Of the 12 guide RNAs designed to target within the TSV LAMP target region (**Figure S2a**), one guide RNA (cr9225) produced cleavage signals before 30 minutes with low variance among replicate reactions and minimal background signal and was selected for the TSV SHERLOCKv2 assay (**Figure 3a**). Of the 11 guide RNAs designed to target within the WSSV LAMP target region (**Figure S2b**), one guide RNA (VP28_27) showed the highest cleavage signal with the greatest signal-to-background ratio and was selected for the WSSV SHERLOCKv2 assay (**Figure 3b**). Furthermore, we found that the addition of glycine, a reaction temperature of 62°C, 2 µM reporter concentration, and a guide RNA-to-Cas12b ratio of 1:1 consistently gave optimal signal-to-background in the WSSV SHERLOCKv2 assay. Because the addition of carrier nucleic acid has shown improvements to sensitivity and specificity in RT-LAMP reactions (33), we tested this in the TSV SHERLOCKv2 assay and found that the addition of 5 or 10 ng of non-target carrier RNA produced the highest signal with the least variance among replicates (**Figure S3**). For WSSV SHERLOCKv2 assay, we decided to use 20 ng of non-target carrier DNA because that was the upper limit of input material reported in the WSSV SHERLOCKv1 limit-of-detection experiment and this amount would allow input for subsequent experiments to be standardized. A step-by-step protocol for SHERLOCK assay development is provided in the **Supplemental Material** along with a table of all oligos and DNA sequences used (**Table S1**) and a table detailing all reaction components (**Table S2**).

**Figure 1.**
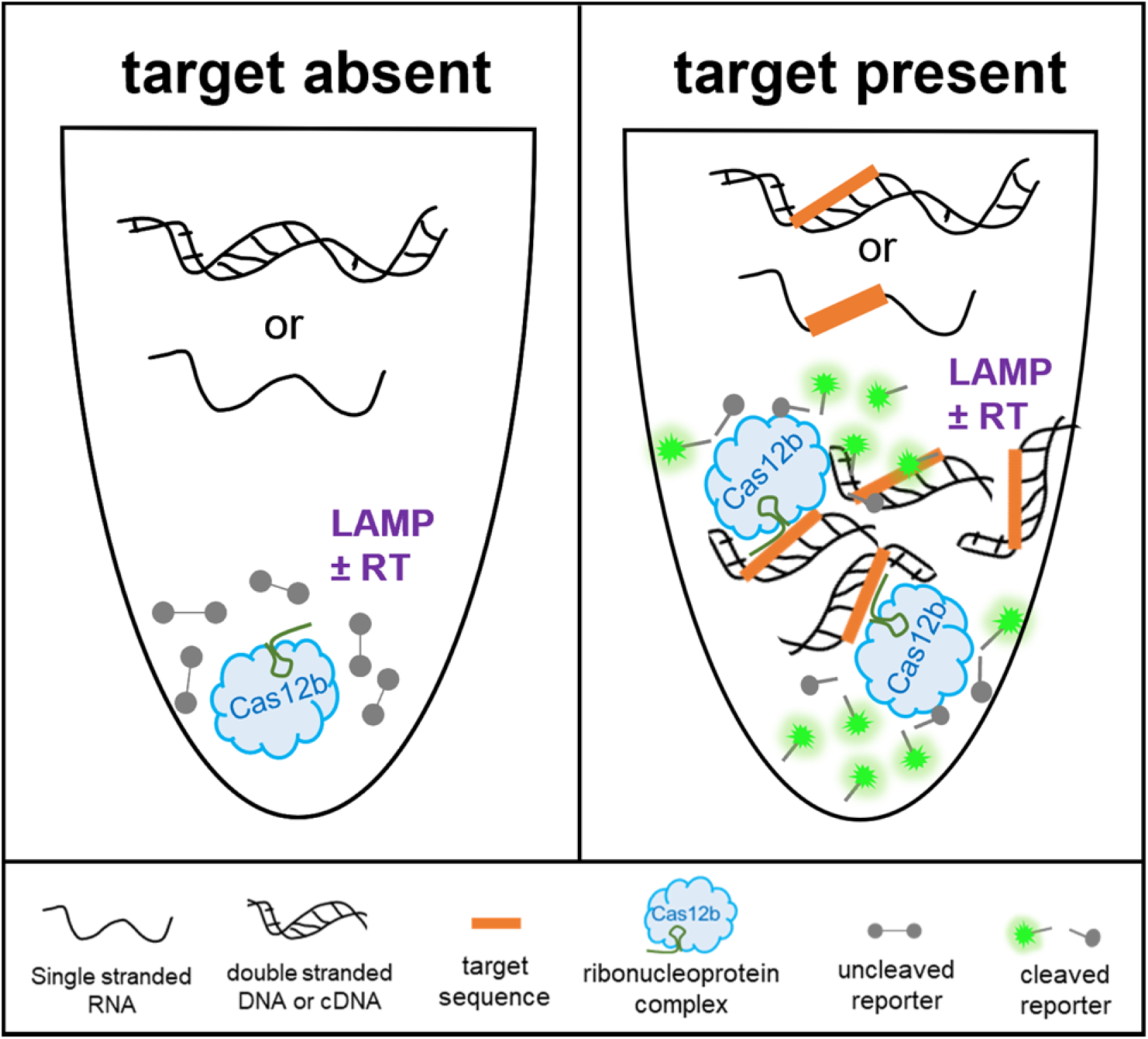
Schematic for CRISPR-based DNA and RNA detection in a one-pot reaction. When the TSV or WSSV target sequence is absent, there is no amplification and no CRISPR-Cas mediated cleavage of the reporter molecule. When the TSV or WSSV target sequence is present, there is isothermal amplification of the target sequence and simultaneous detection by CRISPR-Cas target binding and collateral cleavage of a fluorescent reporter. LAMP ± RT signifies the need for reverse transcriptase (RT) and LAMP for the amplification of the TSV RNA target or need for LAMP only for the amplification of the WSSV DNA target.

**Figure 2.**
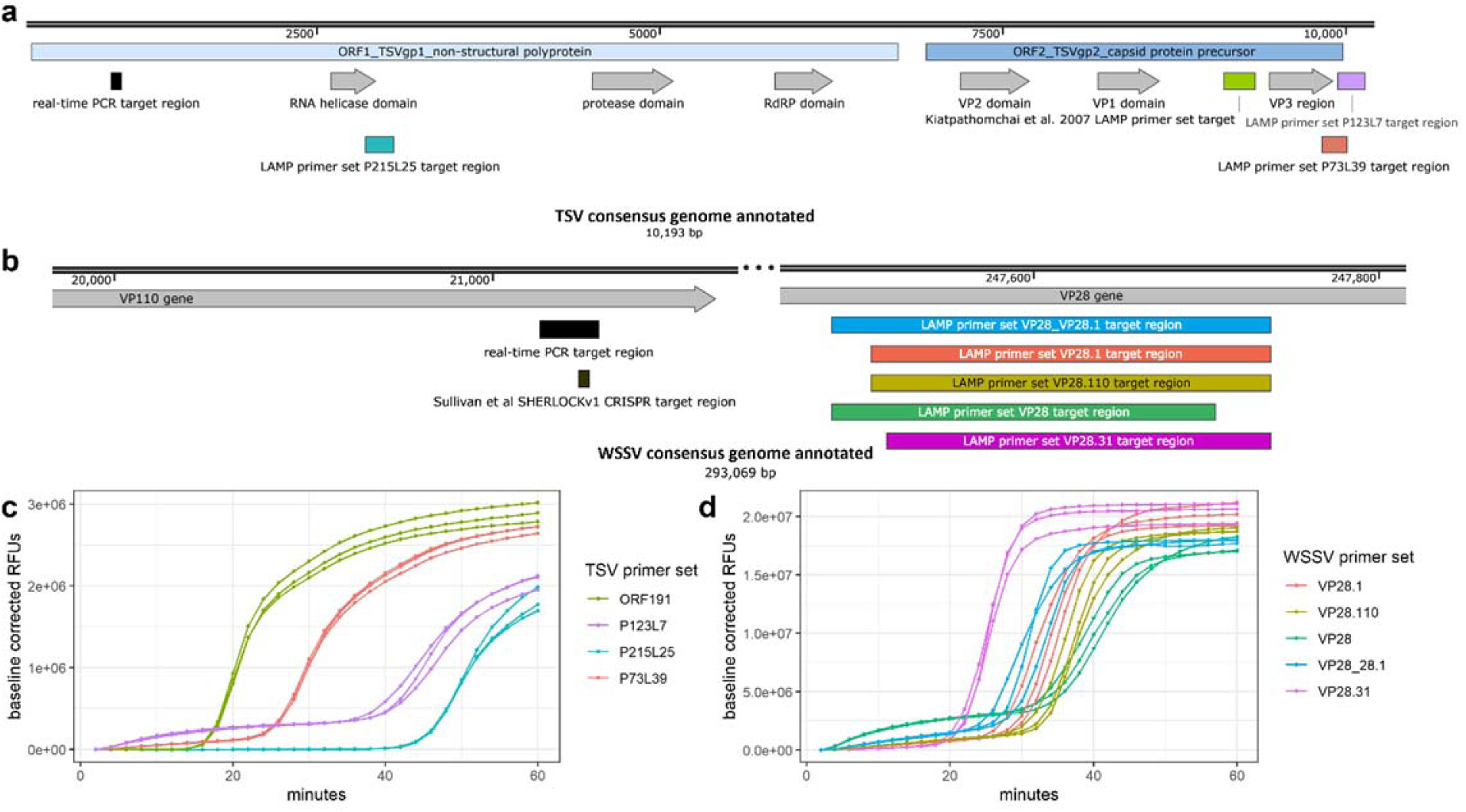
TSV and WSSV LAMP primer design and screening. (**a**) Annotated TSV consensus genome showing OIE standard diagnostic real-time PCR target region in black and LAMP target regions in teal, green, purple, and coral. (**b**) Annotated portions of WSSV consensus genome showing OIE standard diagnostic real-time PCR target region in black, the SHERLOCKv1 target region in brown, and LAMP target regions in blue, coral, gold, green, and magenta. (**c**) Amplification plot of triplicate LAMP reactions screening four different TSV primer sets (indicated by different colors) using 1,000,000 target copies of TSV. (**d**) Amplification plot of triplicate LAMP reactions screening five different WSSV primer sets (indicated by different colors) using 1,000,000 target copies of WSSV.

**Figure 3.**
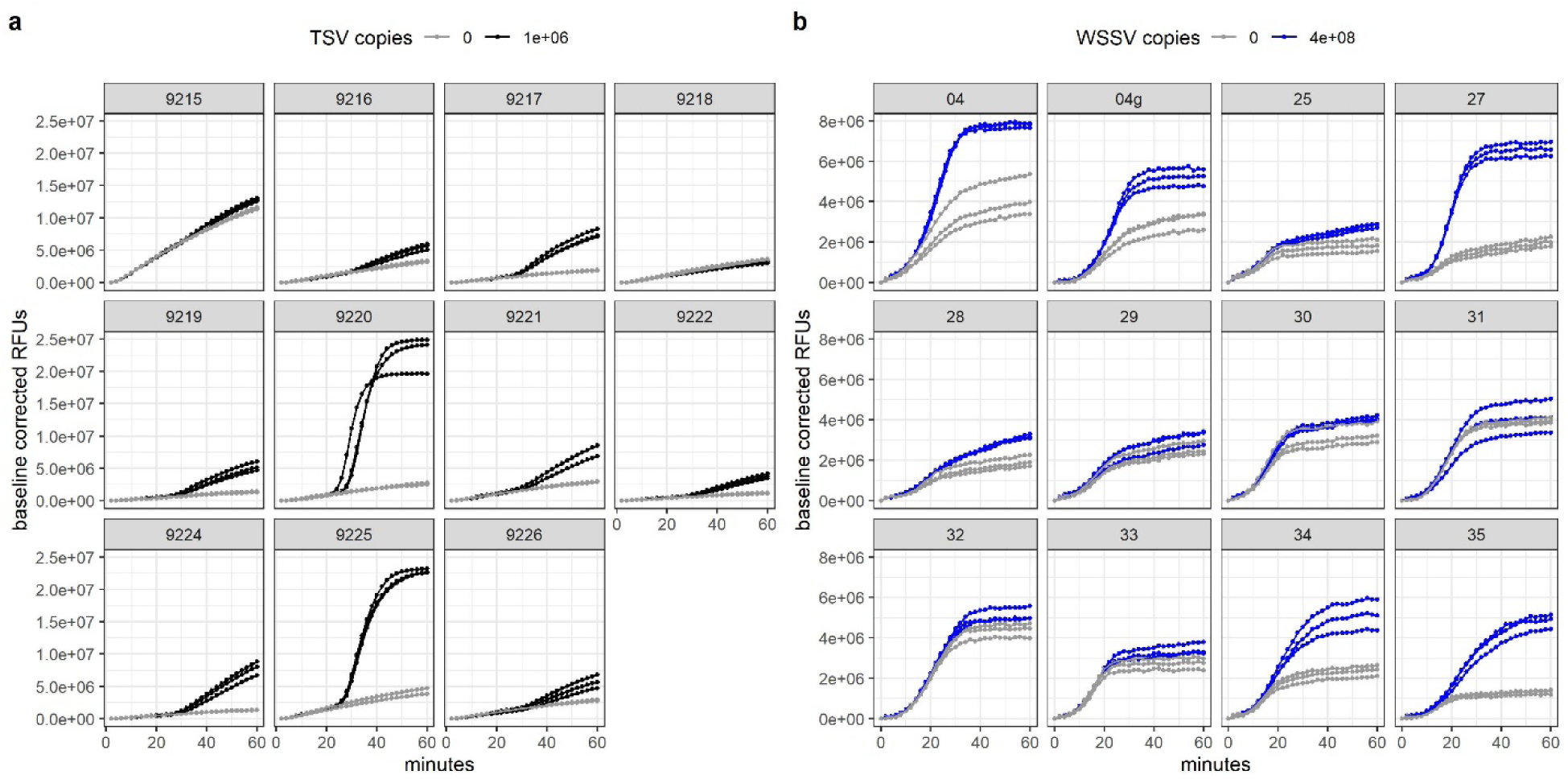
TSV and WSSV guide RNA screening. (**a**) Plots showing Cas12b collateral cleavage activity of each different TSV guide RNA when 1,000,000 (black) or 0 (gray) copies of TSV RNA target is present. (**b**) Plots showing Cas12b collateral cleavage activity of each different WSSV guide RNA when 400,000,000 (blue) or 0 (gray) copies of WSSV DNA target is present. ‘04g’ is the same sequence as ‘04’ but it was transcribed from a gBlock^TM^ instead of an amplicon (see **Materials and Methods** for more details).

### Assay sensitivity and specificity

To evaluate assay sensitivity, we analyzed standard curves generated from 10-fold dilutions of the synthetic TSV target RNA (8 independent experiments) and synthetic WSSV target DNA (6 independent experiments), and both assays consistently detected down to 100 copies for WSSV (200 copies for TSV) with a strong correlation to the amount of nucleic acid input (**Figure 4a–b and S4**). We found synthetic target to be comparable to genomic target isolated from shrimp with high TSV and WSSV viral loads previously quantified by qPCR, confirming that synthetic target can be used for estimating TSV and WSSV copy number quantity in genomic samples (**Figure S4**). In experiments evaluating assay specificity, both SHERLOCKv2 assays showed high specificity for their target pathogen. Only TSV-infected and WSSV-infected shrimp samples showed Cas12b collateral cleavage signal compared to samples from shrimp infected with other common shrimp pathogens (IMNV, IHHNV, EHP and *Vibrio parahaemolyticus* causing AHPND/EMS) (**Figure 4c–d**). Furthermore, both assays showed no false positive detection in samples from specific pathogen-free shrimp (**Figure 4c–d**). While we did observe stochastic non-specific LAMP signal in some true negative samples, this did not lead to Cas cleavage and did not result in positive viral detection (**Figure S5**).

**Figure 4.**
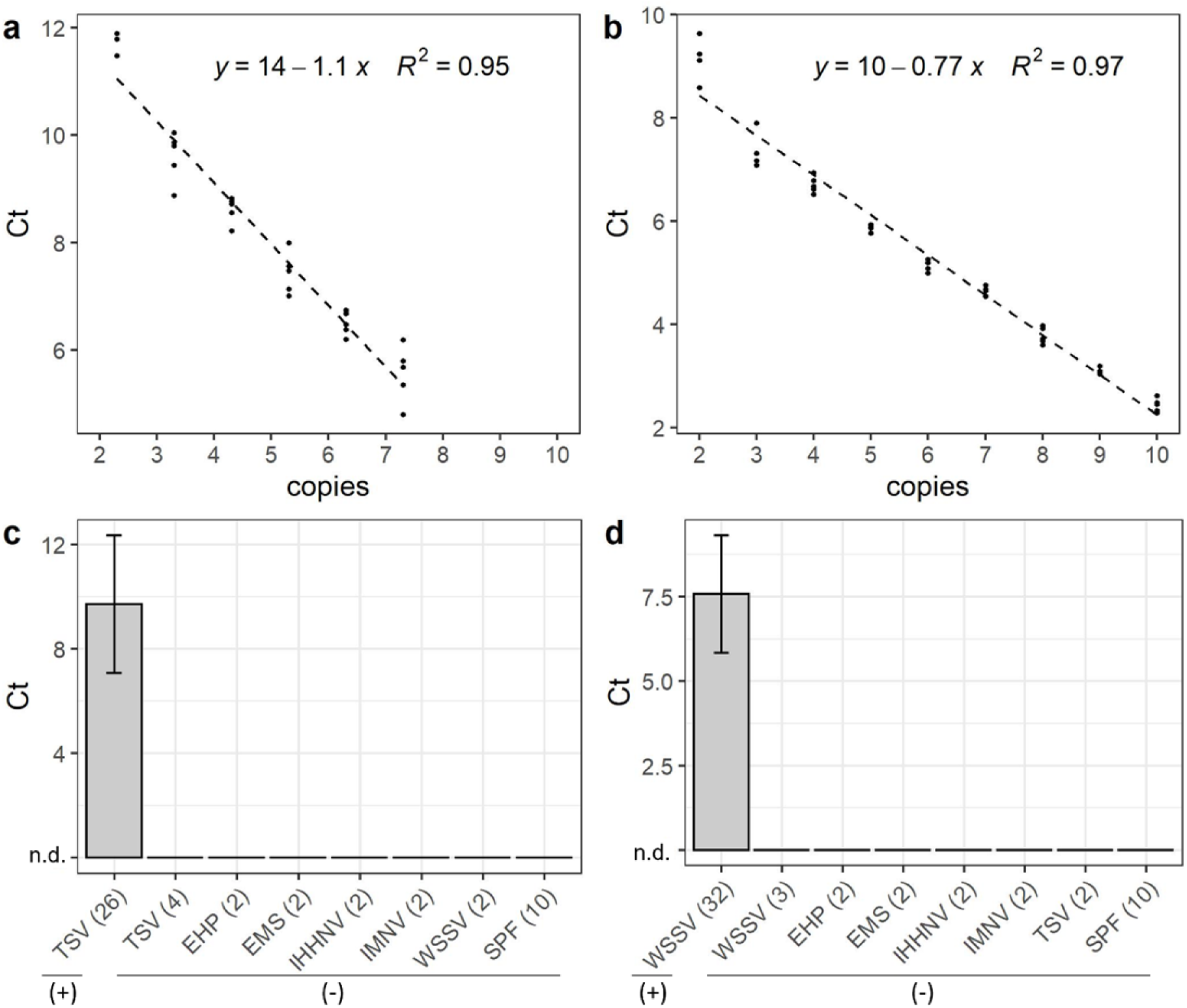
Sensitivity and specificity of SHERLOCK assays. SHERLOCK assay 10-fold standard curve showing Ct values for synthetic (**a**) TSV RNA and (**b**) WSSV DNA target with the dashed line indicating linear regression. Ct values (mean +/-s.d).; n.d., not detected) from SHERLOCK specificity tests evaluating the detection of (**c**) TSV and (**d**) WSSV in shrimp samples known to be infected with TSV, WSSV, and other common white shrimp pathogens including *Vibrio spp*. (causing EMS), EHP, IHHNV, IMNV, and pathogen-free shrimp (SPF). (+) and (-) indicate SHERLOCK results.

### Assay validation

To estimate the diagnostic sensitivity of the SHERLOCKv2 assays, we tested shrimp samples from different TSV and WSSV challenge experiments in SHERLOCKv2 assays and in the OIE (World Organization for Animal Health) recommended real-time PCR diagnostic for TSV and WSSV (34). Assays were run alongside standard curves to quantify and compare viral copy number determined by each method (real-time PCR hereinafter referred to as qPCR). With regards to the TSV challenge experiments, there are lines of shrimp that show a continuum of susceptibility and resistance depending on the line’s genetics and the TSV strain used to evaluate susceptibility (35). We therefore used two different genetic lines with two different isolates that represent two extremes of disease continuum. The Belize isolate used to infect the TSV-tolerant *P. vannamei* line is known to be highly virulent and has led to TSV infection in TSV-tolerant commercial *P. vannamei* lines, where the TSV-susceptible Kona line has shown high rates of infection regardless of the TSV isolate used in challenge experiments (36). Following viral challenge, the TSV-susceptible line displayed clinical manifestation of disease including pathognomonic lesions of TSV characterized by necrosis, pyknosis, and karyorrhexis of the cuticular epithelium within the appendages, gills, and exocuticle (**Figure 5a**). In addition, some of these showed mortalities during the course of the experiment. However, the TSV-tolerant *P. vannamei* showed no TSV-associated pathognomonic lesions or mortalities (**Figure 5b**). Because of the wide dynamic range in viral load expected in these samples, we did a titration experiment to determine an optimal amount of sample input for TSV detection with the TSV SHERLOCK assay. We determined that 10 ng of TSV-infected shrimp genomic RNA led to the least variable and highest signal (**Figure S3**), and this amount was used for subsequent TSV SHERLOCKv2 and qPCR experiments. The qPCR detected TSV in all TSV-challenged shrimp samples (**Figures 5c**, **Table S3**). The TSV SHERLOCK assay detected TSV in 26 out of the 30 TSV-challenged shrimp samples (87% positive percent agreement) (**Table S3**). The four samples for which SHERLOCK did not detect TSV had low viral load extrapolated from a qPCR standard curve (three had < 1000 copies/µL and one had ∼5000 copies/µL). Both qPCR or SHERLOCK TSV assays showed a significant effect of challenge experiment on the viral load (Mann-Whitney-U test *P* = 0.006395 and *P =* 0.003415, respectively), with the susceptible shrimp (Kona line) injected with the TSV El Salvador isolate generally showing a higher titer than the tolerant shrimp injected with the TSV Belize isolate. A Pearson’s correlation analysis followed by a t-test showed a statistically significant correlation between viral copy concentration predicted by SHERLOCK that predicted by qPCR (*R =* 0.93, *p* = 1.5×10^-11^; **Figure 5d**) with SHERLOCK tending to underestimate viral copies relative to qPCR (**Figure 6a**).

**Figure 5.**
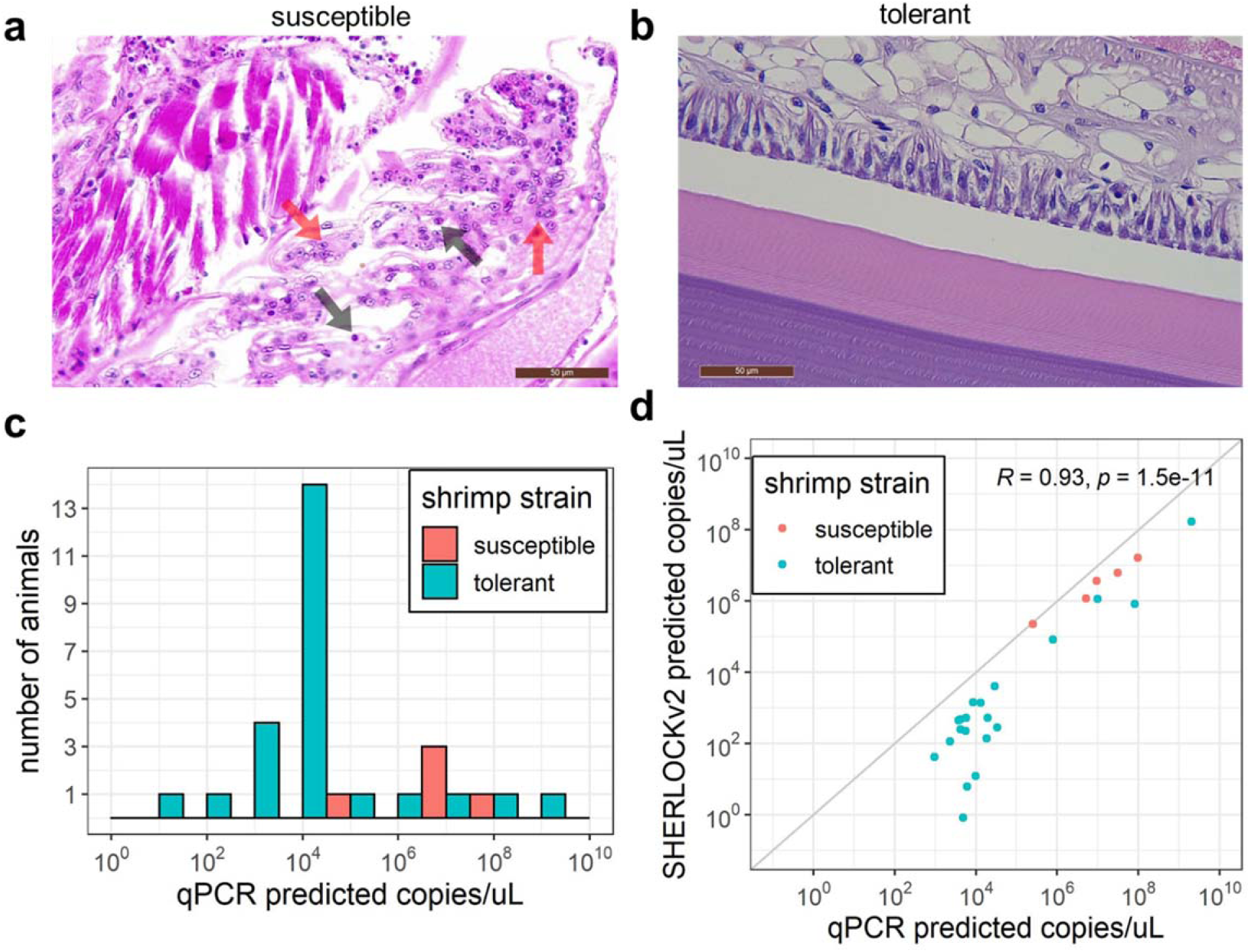
TSV SHERLOCK validation. (**a**) H & E-stained representative section from *P. vannamei* susceptible line after TSV exposure showing focal acute-phase infection in the gill characterized by spherical intracytoplasmic inclusions bodies (black arrows), pyknosis (red arrows) and karyorrhexis. (**b**) H & E-stained representative section from *P. vannamei* resistant line after TSV exposure showing no pathological signs characteristic of TSV infection. (**c**) TSV load quantified by qPCR in resistant (coral) and susceptible animals (teal). (**d**) Correlation between qPCR and TSV SHERLOCK quantification. Gray diagonal line indicates a 1:1 correlation.

**Figure 6.**
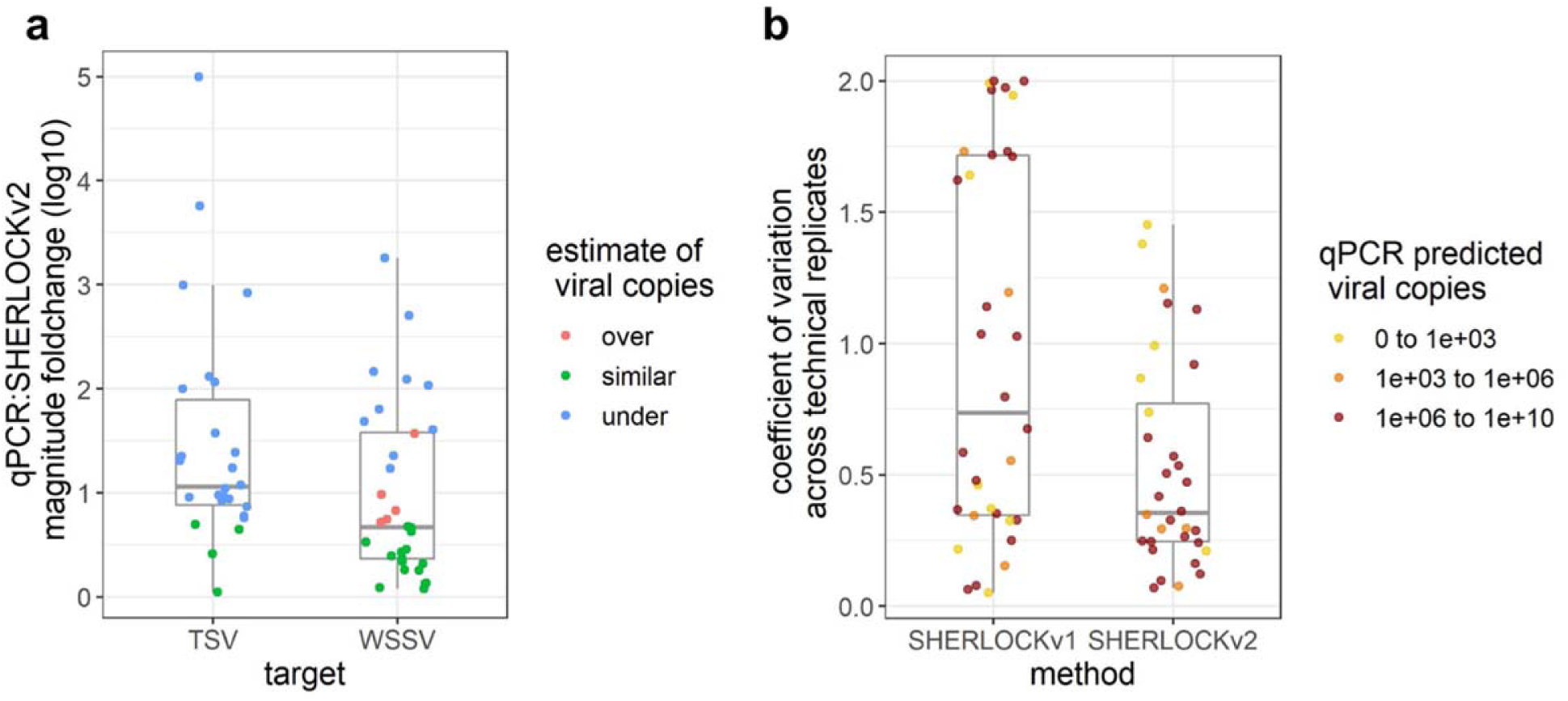
Comparison of SHERLOCKv2 assays to existing molecular assays. (**a**) Magnitude fold change in viral copies estimated by SHERLOCKv2 assays relative to qPCR assays for samples tested in both qPCR and SHERLOCK methods. (**b**) Variation across technical replicates for WSSV samples tested in both SHERLOCKv1 and SHERLOCK v2 assays.

For WSSV, we tested the same WSSV-infected *P. vannamei* samples previously quantified by the OIE-recommended WSSV qPCR and the SHERLOCKv1 assay (28). Given that these samples previously showed high viral loads and sample material was limited, 1 ng of WSSV-infected shrimp genomic DNA was used in each SHERLOCKv2 reaction. Of the 35 WSSV-infected *P. vannamei* samples tested alongside standard curves, we detected infection in 32 samples (91% positive percent agreement) (**Table S3**). The three samples where no WSSV was detected were expected to have low viral copies based on qPCR estimations (∼40, 50, and 800 copies/µL). Moreover, viral copy concentrations measured by WSSV SHERLOCKv2 were mostly similar to those quantified by qPCR (**Figure 6a**). Compared to the previous version WSSV SHERLOCKv1 assay that used RPA and Cas13a enzymes (28), SHERLOCKv2 showed less variation across technical replicates than SHERLOCKv1 (**Figure 6b**). A Pearson’s correlation analysis followed by a t-test showed a statistically significant correlation between SHERLOCKv2 and qPCR viral copy quantifications (*R =* 0.79, *P* = 9.8×10^-8^), and between SHERLOCKv2 and SHERLOCKv1 viral copy quantifications (*R =* 0.75, *P* = 9.6×10^-7^) (**Figure 7**).

**Figure 7.**
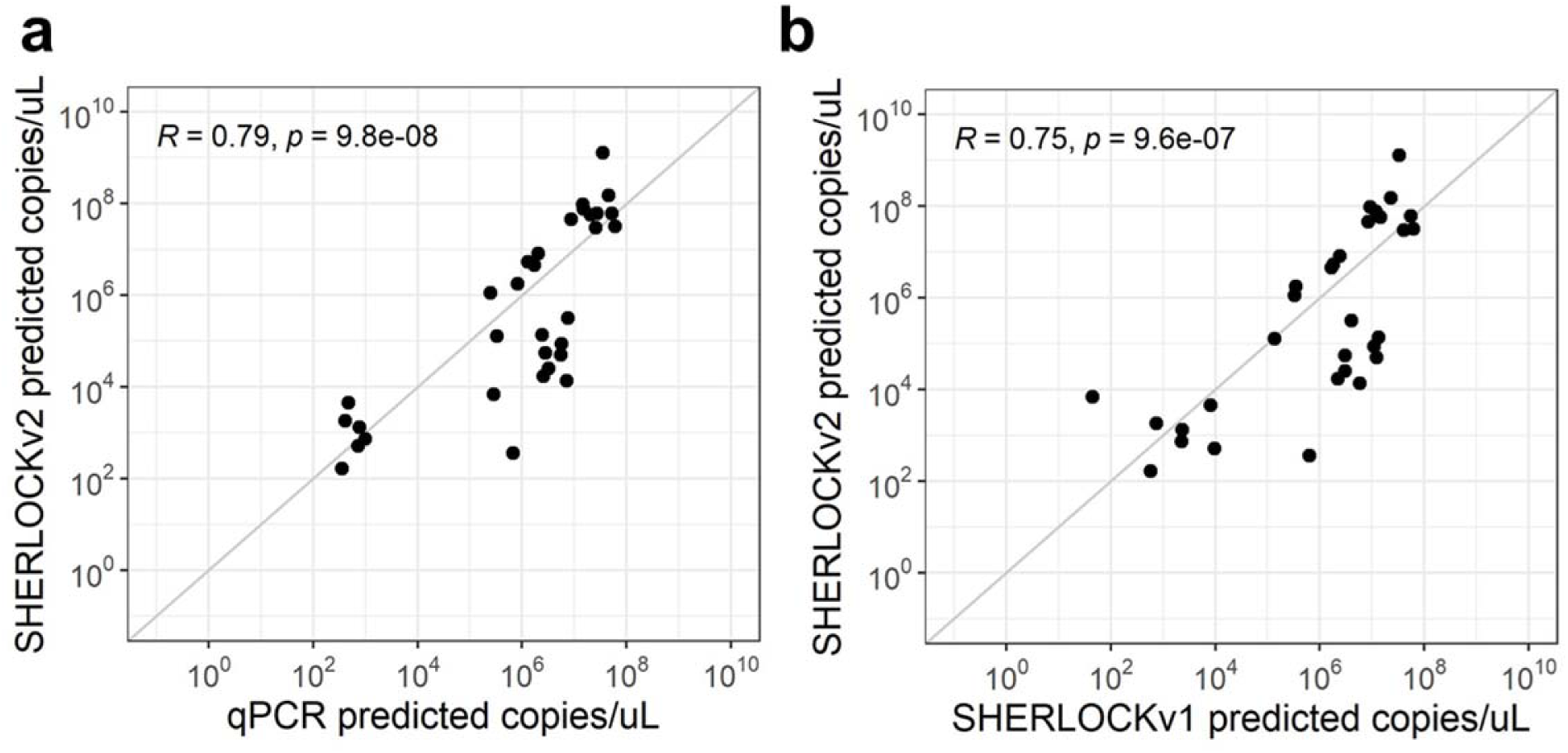
WSSV SHERLOCK assay validation. (**a**) Correlation between qPCR and WSSV SHERLOCKv2 assay quantification. (**b**) Correlation between WSSV SHERLOCKv2 and WSSV SHERLOCKv1 assays. Gray diagonal lines indicate a 1:1 correlation.

## DISCUSSION

With the goal of adapting cutting-edge human diagnostic technology to diseases impacting food security, we developed two different rapid and highly specific lab-based one-pot SHERLOCK fluorescent assays for the detection of common shrimp pathogens. We demonstrate the adaptability of the SHERLOCK platform for targeting either a common RNA virus (TSV) or a common DNA virus (WSSV) that have impacted shrimp aquaculture globally. We found the TSV and WSSV SHERLOCKv2 assays to have a quantitative capability on par with the OIE-recommended qPCR assays for these diseases. Moreover, both assays can assess infection status for a variety of samples in as little as 30 minutes, improving upon the time-to-result of qPCR. Without the need for thermocycling and with the demonstrated ease of converting a one-pot fluorescent SHERLOCK assay to a field-deployable assay (8, 28), these assays lay the foundation for rapid, specific field diagnostics in shrimp aquaculture.

Building on an earlier iteration of a WSSV SHERLOCK assay, we were able to simplify assay components leading to more robust and repeatable detection of both DNA and RNA viral targets. The previous WSSV SHERLOCK assay used RPA to amplify DNA target, RNA polymerase to convert the amplified target from DNA to RNA, and Cas13a to detect RNA target (28). The SHERLOCKv2 assays presented in this study use LAMP (with or without reverse transcriptase) and Cas12b to detect DNA target from RNA or DNA input, following the successful SARS-CoV-2 SHERLOCK assay design (31). By eliminating the need for converting input material to RNA and relying on Cas13a for RNA detection, these SHERLOCKv2 assays enable more stable and robust target detection. This is supported by the reduced variation across technical replicates observed for WSSV SHERLOCKv2 compared to WSSV SHERLOCKv1. Moreover, the use of LAMP in our assays makes them well-poised for use with a variety of sample types including crude sample preparations. Although we did not test crude sample lysates in this study, we have previously successfully demonstrated a paper matrix DNA extraction method to be compatible with WSSV SHERLOCKv1. Given our past success and that the LAMP enzyme (Bst 2.0 polymerase) is highly tolerant to inhibitors (37, 38), we foresee these SHERLOCKv2 assays to be compatible with various sample types.

Pairing LAMP with CRISPR/Cas12b enabled these assays to overcome non-specific signal that can occur in assays using LAMP alone. In the SHERLOCKv2 assays, stochastic non-specific LAMP products did not lead to Cas cleavage signal. Although isothermal amplification-based methods recently developed for aquaculture diagnostics have shown that sequence-specific probes can enhance assay specificity (39, 40), the combination of primers and probes cannot always be relied on for sequence-specific detection (41). Cas detection systems rely on sequence-specific target binding to induce conformational change that triggers cleavage activity (42), and combining isothermal amplification methods with this additional layer of specificity is becoming more common in aquaculture molecular diagnostic approaches (28, 43). Despite reports of single insertion and deletion tolerance in some Cas complexes (44), we found Cas12b to be highly specific, supportive of previous reports of Cas12b specificity down to the single nucleotide level (45).

Although disease diagnosis typically only requires a binary result, the ability to quantify viral load can be useful. The significant correlations we found between SHERLOCKv2 assays and qPCR assays suggest SHERLOCKv2 can be semi-quantitative. The TSV SHERLOCKv2 assay tended to underestimate viral copies relative to qPCR while the WSSV SHERLOCKv2 assay mostly showed similar viral copies to qPCR. These discrepancies could be explained by differences in amplification efficiencies of different genomic target sequences. The TSV qPCR assay targets near the beginning of the ORF1 (TSVgp1) region whereas the TSV SHERLOCK assay targets near the end of the ORF2 region. The primers used for the OIE-recommended WSSV qPCR protocol were designed to target a genomic region ∼20 kb upstream of the VP28 gene region, while the LAMP primers and guide RNA for the WSSV SHERLOCK assay target within the VP28 gene region itself. As amplification efficiency is affected by differences in structure and sequence length in target regions as well as primers, it can be expected that quantifications would differ between methods targeting different regions. Additionally, the structural differences in the synthetic target standards used in the SHERLOCK assays (200-300 DNA or RNA fragments) and qPCR assays (plasmids) could have contributed to discrepancies in results (46). Differences in data analysis between qPCR and SHERLOCKv2 methods could have also contributed to discrepancies in viral detection among the infected shrimp samples. Samples assessed in the qPCR assays were run in triplicate and considered to be positive if at least one replicate had a Ct value ≤ 40 for diagnosis (34), while samples assessed in the SHERLOCKv2 assays were run in quintet and considered to be positive if at least three replicates passed a baseline and an outlier threshold. The intention of this SHERLOCK scoring criteria was to rigorously control for technical variation and precisely and confidently measure viral presence and abundance in the development of these new assays. If the criteria were relaxed to require only two samples passing the baseline threshold and not include an outlier threshold, TSV infection would have been found in 29/30 samples (97% positive percent agreement) and WSSV infection would have been found in 35/35 samples (100% positive percent agreement) (47). In the future, the SHERLOCK scoring criteria used here can likely be relaxed following the testing of larger number of samples. Similar to existing reports of the proportion of replicates testing positive being relative to target concentration in both SHERLOCK and qPCR assays (48, 49), the few samples with virus detected by qPCR and not by SHERLOCK were indeed closer to the LODs of both TSV and WSSV assays. Despite the differences in analysis methods and in the nature of template used to optimize the SHERLOCK assay compared to the qPCR assay, the binary qualitative results in detecting virus in shrimp tissue showed positive percent agreements (87% for TSV SHERLOCKv2 and 91% for WSSV SHERLOCKv2) and negative percent agreements (100% for both assays) within or close to the strict range required by the FDA for clinical SARS-CoV-2 assays (90% positive percent agreement and 95% negative percent agreement) (50). Moreover, the quantitative loads of the corresponding viruses as measured by the two assays were also comparable. This gives validity to the SHERLOCKv2 assays when compared to the current gold standard of TSV and WSSV detection recommended by the OIE.

With molecular diagnostics rapidly advancing alongside outbreaks of existing and emerging disease, the transfer of technology from human diagnostics to animal fields is critical, particularly to those that support global food security. With a growing need to support aquaculture production, we demonstrated the ability to adapt a SHERLOCK assay developed for human diagnostics for detecting RNA and DNA shrimp pathogens. This development of novel (± RT)-LAMP + CRISPR/Cas diagnostic assays for TSV and WSSV monitoring in *P. vannamei* and potential for field-deployable conversion will accelerate biomonitoring for shrimp aquaculture and pave the way for the developing rapid, efficient diagnostics across the agriculture field.

## MATERIALS AND METHODS

### Viral challenge experiments

For TSV, two independent challenge experiments were conducted using *P. vannamei* lines that differ in TSV susceptibility and TSV isolates that differ in virulence. The purpose of this was to generate tissues with a wide dynamic range of viral load. In one experiment, TSV-susceptible specific pathogen free (SPF) *P. vannamei* (Kona line) shrimp (N = 120) were injected with TSV inoculum (El Salvador isolate, Lightner et al. unpublished) intramuscularly and reared until animals were moribund (4 days). In a second experiment, individuals from a commercial line of TSV-tolerant SPF *P. vannamei* (N = 30) were injected with a TSV inoculum (Belize isolate (36)) intramuscularly. Shrimp were reared for 4-8 days in a single 1600 L tank containing 1000 L artificial sea water maintained at 27 ± 1 °C and a salinity of 30 ppt. No water change was done due to the short duration of the experiments. Shrimp tissues were processed for histopathology analysis following a standard procedure (51). Briefly, tissues were preserved in Davidson’s AFA fixative for 48 hours after which the fixative was replaced with an equal volume of 70% ethanol. Samples were then washed in a series of alcohol/xylene solutions, embedded in paraffin, sectioned to 4µm using a microtome. The tissue sections were stained with hematoxylin and eosin, and examined using a bright field light microscope. Histopathological evaluations were performed using a subset of samples from the experimentally infected tolerant *P. vannamei* (N = 6) and susceptible *P. vannamei* (N = 5). The severity of the TSV infection was graded based on a semi-quantitative scale ranging from Grade 0 to Grade 4: Grade 0 showing no signs of infection; Grade 1 showing signs of infection by the pathogen but at levels that may be below those needed for clinical disease classification; Grade 2 showing moderate signs of infection based on the number and severity of pathogen-caused lesions; Grade 3 showing moderate to high signs of infection based on the number and severity of pathogen-caused lesions; and Grade 4 showing high signs of infection based on a high number of pathogen-caused lesions and tissue destruction (52). Total RNA from experimentally TSV-challenged shrimp (N = 35) was extracted with the Total RNA Purification Kit (NORGEN BIOTEK CORP) following the manufacturer’s recommendations. Quality and quantity were assessed using a NanoDrop™ (Thermo Fisher). The RNA concentrations of 30 samples were normalized to approximately 15 ng µl^-1^ and TSV was detected and quantified from 2 µl of sample using a TaqMan™ Fast Virus 1-Step Master Mix (Applied Biosystems™) following the OIE recommended protocol and triplicate standards (mean R^2^ = 0.93 ± 0.06, mean efficiency 105.3% ± 15.28%) (53). All samples were run in triplicate alongside plasmid standards with known copy numbers (100 - 1,000,000 copies). TSV load in the experimentally challenged shrimp was calculated by extrapolating Ct values in the standard curve generated using plasmid DNA template.

Unlike TSV, all commercially available *P. vannamei* lines are highly susceptible to WSSV with shrimp mortality occurring 2-3 days post-challenge in routine bioassays conducted in the Aquaculture Pathology Laboratory at University of Arizona. Details of the viral challenge experiment with WSSV of *P. vannamei* are reported in Sullivan et al. 2019 (28). Briefly, specific pathogen free shrimp (N = 137) were challenged by feeding WSSV (China isolate CN95 (54)) infected tissue and reared under the same conditions and duration as described above. Infection was confirmed in moribund animals by histology as described above. Genomic DNA was extracted with the DNeasy blood and tissue kit (Qiagen), quantified using a NanoDrop™ (Thermo Fisher), and quantified by qPCR using 1 µl of sample with TaqMan Fast Virus 1-Step Master Mix run in triplicate on an ABI StepOne Plus system and following the OIE protocol using triplicate standards (mean R^2^ = 0.99 ± 0.0007, mean efficiency 90.6 ± 12.0%) (55).

### TSV and WSSV SHERLOCK assay development

To identify genomic targets for TSV and WSSV viral detection, publicly available viral genome sequences were searched for consensus regions. For TSV, eight genomes were retrieved from GenBank (https://www.ncbi.nlm.nih.gov/genbank/; JF966384.1 (56), NC_003005.1 (57), JX094350.1 (58), AF277675.1(57), GQ502201.1 (59), DQ104696.1 (60), AY997025.1 (61), DQ212790.1 (62)) and aligned with MAFFT (63) using the default parameters to generate a consensus sequence (**Table S1**). For WSSV, the highly conserved viral envelope protein 28 (64) was selected as a target and conservation across multiple strains was confirmed by surveying a consensus genome compiled from 10 different isolates as previously reported (28).

To generate candidate LAMP primer sets, target sequences were input into the NEB LAMP Primer Design tool (https://lamp.neb.com). For TSV, three LAMP primer sets were selected as well as a previously published LAMP primer set (32) for screening. For WSSV, four different candidate LAMP primer sets were selected and the primer set was optimized by testing sets containing different combinations of primers as well as primer modifications including the addition of a poly-T linker between the F1c/B1c and F2/B2 annealing sites of the FIP and BIP primers (**Table S1**).

Guide RNAs for Cas12b collateral cleavage were designed using a sliding window of 20 bases in length and advancing one base at a time across the region between the F2 and B2 annealing sites of the LAMP FIP and BIP primers. To synthesize guide RNAs, a gBlock™ (IDT) containing the *Aap* Cas12b sgRNA scaffold sequence was first amplified using a forward primer containing the T7 promoter sequence and reverse primers containing each different spacer sequence. Amplification reaction conditions were as follows: 1X HF Buffer (NEB), 200 µM dNTPs (NEB), 0.5 µM forward primer, 0.5 µM reverse primer, 0.01 ng template DNA, and 1 unit Phusion DNA polymerase (NEB) in 50 µL. Reactions were run on a thermocycler (Applied Biosystems) with the following parameters: 98°C for 2 minutes, 30 cycles of 98°C for 10 seconds, 58°C for 30 seconds, and 72°C for 15 seconds, 72°C for 5 minutes, and a hold at 10°C. Products were gel purified using the QIAquick Gel Extraction kit (Qiagen) with a 2% agarose 1x TAE gel and products were eluted in 30 µL of Buffer EB prewarmed to 50°C. Concentrations were measured using the Qubit Broad Range dsDNA kit (Life Technologies). Next, *in vitro* transcription reactions were performed using the Ampliscribe T7 flash transcription kit (Lucigen) in a 40 µL reaction with 50 ng of DNA template, 9 mM of each ribonucleotide, 9 mM of DTT, 1 µL of RiboGuard RNase inhibitor, 4 µL of T7 RNA polymerase, and 1X transcription reaction buffer. Reactions were run at 42°C for 4 hours, incubated with 2 µL of DNase I (Lucigen) at 37°C for 15 minutes, and purified using the RNA Clean and Concentrate Kit (Zymo Research). Guide RNAs were quantified using the Qubit Broad Range RNA assay kit (Life Technologies) and molarity was estimated using the NEB copy number converter (https://nebiocalculator.neb.com/). All sequences from the consensus genome, gBlocks™ (IDT), primers, and guide RNAs are listed in **Table S1**.

### Evaluating LAMP primers

For TSV, primer sets were screened against a synthetic TSV RNA target *in vitro* transcribed from gBlock™ DNA (IDT, **Table S1**) using the Ampliscribe T7-Flash Transcription kit (Lucigen Corporation) and purified using the RNA Clean and Concentrator kit (Zymo Research) following manufacturer’s instructions. For WSSV, primer sets were screened against either a synthetic WSSV DNA target (gBlock™, IDT; **Table S1**) or genomic DNA isolated from shrimp with a high WSSV load. Target nucleic acid was diluted in background nucleic acid from SPF shrimp as detailed in **Table S2e** to improve the specificity of the LAMP primers. Total RNA from 10 SPF shrimp was isolated using the RNeasy blood and tissue kit (Qiagen) and genomic DNA from 10 SPF shrimp was isolated using the DNeasy blood and tissue kit (Qiagen). LAMP reactions were prepared as detailed in **Table S2a** and run at 60°C and 62°C (TSV and WSSV; respectively) for 1 hour on a thermocycler for real-time PCR (QuantStudio 12K flex, Thermo Fisher) with fluorescent measurements taken at 2-minute intervals. Successful primers were determined by their ability to start amplification before 30 minutes.

### Evaluating guide RNAs and Cas12b detection

Guide RNAs were systematically evaluated with Recombinant *Aap* Cas12b (GenScript) for efficacy using TSV LAMP product or synthetic WSSV target DNA (gBlock™, IDT) in 20 µL (TSV) or 10 µL (WSSV) reactions described **Table S2b**. Prior to adding template and reporter, reaction mixtures were incubated for 15 minutes at room temperature to facilitate the formation of the CRISPR/Cas ribonucleoprotein complex. For each guide RNA, a negative control was run that did not contain target nucleic acid to measure background collateral cleavage, and all reactions were run in triplicate. Reactions were run in a 384-well black clear bottom plate (Corning) at 60°C for 1 hour with measurements recorded every 2 minutes using a Spectramax iD3 plate reader (Molecular Devices). Effective guide RNAs were identified as those that generate a signal well-distinguished from the negative control in less than 30 minutes.

### Evaluating sensitivity and standard curve generation

To produce standard curves for each assay, one-pot LAMP-Cas12b SHERLOCK reactions were run using either *in vitro* transcribed synthetic TSV RNA target, synthetic WSSV DNA target (gBlock™, IDT; **Table S1**), or genomic DNA isolated from shrimp with a high viral load. Ten-fold dilution series of synthetic DNA or RNA were made ranging from 10^1^ to 10^6^ copies µL^-1^ (TSV) or 10^1^ to 10^10^ copies µL^-1^ (WSSV) with all samples diluted in specific pathogen free shrimp genomic RNA (5 ng per 20 µL reaction for TSV) or DNA (20 ng per 20 µL reaction for WSSV). Negative control reactions that did not contain any target nucleic acid were included in each experiment. Reactions were run with 5 replicates per sample prepared as detailed in **Table S2c** and run at 60°C (TSV) or 62°C (WSSV) using a QuantStudio 12k Flex thermocycler for real-time PCR (Applied Biosystems) for 60 minutes, with fluorescence readings every 2 minutes. Prior to adding dNTPs, *Bst* 2.0 DNA polymerase (New England BioLabs), primer mix, fluorescent reporter, SYTO-82 dye, or DNA, other reaction components were combined and incubated for 15 minutes at room temperature to facilitate the formation of the CRISPR/Cas ribonucleoprotein complex.

### Evaluating SHERLOCK quantification

To compare quantification of viral loads estimated by the OIE-recommended qPCR and SHERLOCK, one-pot TSV SHERLOCK reactions were run on the same RNA samples from 30 TSV-infected shrimp quantified by qPCR described above. For WSSV, one-pot SHERLOCK reactions were run on the same DNA samples from 35 WSSV-infected shrimp previously quantified by qPCR (28). To standardize input across samples, 10 ng of RNA (TSV) or 1 ng of DNA (WSSV) was added per 20 µL SHERLOCK reaction. SHERLOCK reactions were prepared as detailed in **Table S2c**, and all samples were run alongside standard curves as described above. A linear regression analysis was performed on Ct values from the standard curve samples calculated from fluorescent measurements that were corrected to a baseline defined by cycles 1-2 for WSSV and 1-4 for TSV, and viral copies were estimated for each sample from the extrapolated standard Ct values. For each sample, including standards, any replicate outlier based on a Ct value beyond the sample median Ct ± one cycle was excluded and only samples with at least three out of five replicates were considered in downstream analyses. Viral copies estimated by SHERLOCK one-pot reactions were then compared to those estimated by qPCR using a Pearson correlation analysis followed by t-test with the stat_cor function in the R package ggpubr (65).

### Evaluating specificity

The specificity of each one-pot SHERLOCK assay was tested against either RNA or DNA extracted from shrimp tissue infected with infectious myonecrosis virus (IMNV; muscle tissue), infectious hypodermal and haematopoietic necrosis virus (IHHNV; muscle tissue), *V. parahaemolyticus* causing acute hepatopancreatic necrosis disease/early mortality syndrome (AHPND/EMS; hepatopancreas tissue), and *Enterocytozoon hepatopenaei* (EHP; hepatopancreas tissue). RNA was extracted using RNeasy mini plus kit (Qiagen) and DNA was extracted using the Qiagen DNeasy blood and tissue kit following manufacturer’s protocols. One-pot reactions were prepared as detailed in **Table S2c** with a total of 10 ng of sample RNA (TSV) or 1 ng of sample DNA (WSSV) in each reaction. To estimate false positive rates, one-pot reactions were prepared as detailed in **Table S2c** and run as described above with 5 ng of total RNA or 20 ng of genomic DNA isolated from muscle tissue from 10 individual specific pathogen-free shrimp.

### Data availability

Raw fluorescence data from all qPCR and SHERLOCK experiments, intermediate files, and R code used to generate plots and statistics are publicly available at https://doi.org/10.5281/zenodo.7438263 (47).

## ACKNOWLEDGEMENTS

We thank Veronica L. Pereira and Zachary G. Dench (Gloucester Marine Genomics Institute) for their contributions towards previous iterations of assay development that inspired the assay design described here. We also thank New England BioLabs and Sherlock Biosciences for helpful discussions and advice.

## FUNDING

This work was supported by the United States Department of Agriculture NIFA AFRI Exploratory Research Program (grant # 2018-08980)

## AUTHOR CONTRIBUTIONS

S.R.M. and S.A.T. designed and performed molecular experiments, analyzed the data, and wrote the manuscript; M.J.H provided input throughout the project and edited the manuscript; R.C.-F. conducted histopathology and preliminary qPCR experiments; A.K.D. provided all laboratory challenge samples, developed the experimental design, managed the histopathology verification, provided input throughout the project, and contributed to manuscript revisions; A.G.B. conceived the project, provided input throughout its progress, and edited the manuscript.

## Notes

https://doi.org/10.5281/zenodo.7438263

### Competing Interest Statement

The authors have declared no competing interest.

### Summary of Updates

figures, text, tables, and supplemental material for clarity

