## supplemental material for "Rapid detection of DNA and RNA shrimp viruses using CRISPR-based diagnostics"

**Table of contents**

**Supplemental Protocol.** One-pot SHERLOCK protocol for WSSV and TSV detection

**Supplemental Figures**
Figure S1. Effects of WSSV LAMP primer modifications
Figure S2. Target regions of guide RNAs screened for TSV and WSSV assays
Figure S3. Optimizing amount of non-target and target DNA for the TSV SHERLOCK assay
Figure S4. Comparison of synthetic and genomic target dilution series
Figure S5. Non-specific LAMP does not give rise to Cas cleavage

**Supplemental Tables**
Table S1. Consensus genome, gBlock oligos, primers, sgRNA, and reporter oligo sequences used for developing TSV and WSSV assays
Table S2. Composition of LAMP primer evaluation reactions, Cas sgRNA evaluation reactions, and one-pot SHERLOCK reactions, template dilutions, and 10x primer mix
Table S3. Viral concentrations estimated by qPCR, SHERLOCKv1, and SHERLOCKv2 assays

**Supplemental Protocol. One-pot SHERLOCK protocol for WSSV and TSV detection**

*Materials*

- Nuclease free water
- 10X Isothermal Amplification Buffer (New England BioLabs, Cat# B0537S)
- 100 mM $MgSO_4$ (New England BioLabs, Cat# B1003S, supplied with M0538L)
- 10 mM dNTPs (New England BioLabs, Cat# N0447L)
- Bst 2.0 WarmStart DNA Polymerase (New England BioLabs, Cat# M0538L)
- WarmStart RTx Reverse Transcriptase (New England BioLabs, Cat# M0380L)
- 500 mM Taurine (Millipore Sigma, Cat# 86329)
- 2 M Glycine (ThermoFisher, Cat# BP381-500)
- *Aap* Cas12b enzyme (synthesized by GenScript)
- 5'FAM/TTTTT/3'IBFQ reporter (100 µM, Integrated DNA Technologies)
- SYTO-82 Fluorescent Stain (ThermoFisher, Cat# S11363)
- Sample DNA or RNA
- 10 µM Guide RNA
    - Guide RNA can be prepared ahead of time and stored at -80°C before use.
    - Prepare guide RNA as follows:
        1. Amplify guide RNA template using the following oligos and conditions:
            - *Aap Cas12b sgRNA scaffold gBlock* 5'-TGTAAAACGACGGCCAGTcatataTAATACGACTCACTATAGGGGTCTAGAGGACAGAATTTTTCAACGGGTGTGCCAATGGCCACTTTCCAGGTGGCAAAGCCCGTTGAGCTTCTCAAATCTGAGAAGTGGCAC GTCATAGCTGTTTCCTG-3'
            - Cas12b_sgRNA_fwd primer 5'-CATATATAATACGACTCACTATAGGGGTCTAGAGG-3'
            - Cas12b_sgRNA_rev primer(s): Replace x's with 20 base target region 5'-xxxxxxxxxxxxxxxxxxxxGTGCCACTTCTCAGATTTGAGAAGCTC-3'
            - 1X HF Buffer (New England BioLabs), 200 µM dNTPs, 0.5 µM forward primer, 0.5 µM reverse primer, 0.01 ng gBlock DNA, and 1 unit Phusion DNA polymerase (New England BioLabs) in 50 µL.
            - Run reaction on a thermocycler (Applied Biosystems) with the following parameters: 98°C for 2 minutes, 30 cycles of 98°C for 10 seconds, 58°C for 30 seconds, and 72°C for 15 seconds, 72°C for 5 minutes, and a hold at 10°C.
        2. Gel purify PCR products on a 2% agarose 1x TAE gel and using the QIAquick Gel Extraction kit (Qiagen) following manufacturer's instructions.
        3. Transcribe guide RNA using the Ampliscribe T7 flash transcription kit (Lucigen)
            - In a 40 µL reaction, combine 50 ng of gel purified PCR product, 9 mM of each ribonucleotide, 9 mM of DTT, 1 µL of RiboGuard

RNase inhibitor, 4 µL of T7 RNA polymerase, and 1X transcription reaction buffer.
    - Run reactions at 42°C for 4 hours
    - Incubate with 2 µL of DNase I (Lucigen) at 37°C for 15 minutes
    - Purify using the RNA Clean and Concentrate Kit (Zymo Research) following manufacturer's instructions
    - Store purified guide RNA at -80°C.
- 10X LAMP Primer Mix
  - 10X Primer mix can be prepared ahead of time and stored at -20°C before use.
  - Prepare a 10x primer mix of the LAMP primers as follows:

| **10X Primer Mix** | | | |
|---|---|---|---|
| **LAMP Primer** (100 µM) | **10x concentration** (µM) | **Reaction Concentration** (µM) | **Volume** (µL) |
| F3 | 2 | 0.2 | 2 |
| B3 | 2 | 0.2 | 2 |
| BIP | 16 | 1.6 | 16 |
| FIP | 16 | 1.6 | 16 |
| LF* | 4 | 0.4 | 4 |
| LB* | 4 | 0.4 | 4 |
| dd$H_2O$ | | | 56 |
| Total | | | 100 |

* If not included in LAMP primer set, replace with water

**Procedure**

1. Prepare sample DNA or RNA
   - For synthetic target, dilutions are made as follows:
     - For TSV, combine RNA stock containing $10^{10}$ copies TSV target, nuclease free water, and specific pathogen free shrimp genomic RNA (5 ng per 20 µL one-pot reaction)
     - For WSSV, combine DNA stock containing $10^{9}$ copies WSSV target, nuclease free water, and specific pathogen free shrimp genomic DNA (20 ng per 20 µL one-pot reaction)
   - For genomic DNA or RNA extracted from shrimp, dilutions are made as follows:

| **Template dilution component** | **WSSV** | **TSV** |
|---|---|---|
| Sample DNA/RNA | 0.25 ng/µL (1ng) | 2 ng/µL (10 ng) |
| Pathogen free shrimp nucleic acid (DNA for WSSV, RNA for TSV) | 5 ng/µL (20 ng) | 1 ng/µL (5 ng) |
| Nuclease free water | variable | variable |
| Total volume/one-pot reaction | 4 µL | 5 µL |

2. Prepare the master mix by combining the reagents in the following order:

| One-Pot SHERLOCK Reaction component | Initial concentration | Final concentration | WSSV reaction volume (µL) | TSV reaction volume (µL) |
|---|---|---|---|---|
| water | | | 3.44 | 1.46 |
| Isothermal Amp Buffer | 10x | 1x | 2 | 2 |
| $MgSO_4$ | 100 mM | 8 mM | 1.6 | 1.6 |
| glycine | 2 M | 200 mM | 2 | |
| taurine | 500 mM | 50 mM | | 2 |
| Cas12b | 8.73 µM | 200 nM (150 nM for TSV) | 0.458 | 0.344 |
| sgRNA | 10 µM | 200 nM (600 nM for TSV) | 0.4 | 1.2 |
| dNTPs | 10 mM | 1.4 mM | 2.8 | 2.8 |
| *incubate at room temperature for 15 minutes to facilitate formation of CRISPR-Cas complex* | | | | |
| WarmStart RTx Reverse Transcriptase | 15000 units/mL | 150 units/mL | | 0.2 |
| WarmStart Bst 2.0 | 8000 units/mL | 320 units/mL | 0.8 | 0.8 |
| Primer mix | 10x | 1x | 2 | 2 |
| SYTO-82 LAMP dye | 100 µM | 0.5 µM (1 µM for TSV) | 0.1 | 0.2 |
| 5'FAM/TTTTT/3'IBFQ reporter | 100 µM | 2 µM | 0.4 | 0.4 |
| Sample DNA/RNA | | | 4 | 5 |
| Total | | | 20 | 20 |

3. Run on real-time PCR thermocycler at 62°C for WSSV or 60°C for TSV for 1 hour taking a reading every 2 minutes.

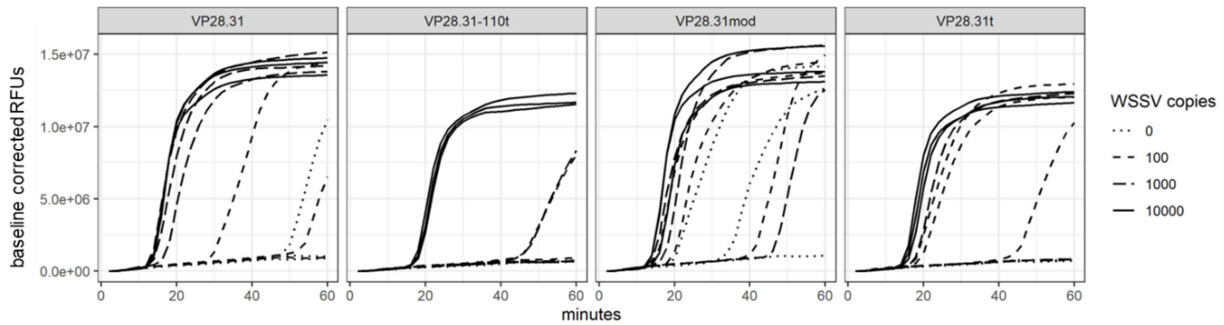


**Figure S1.** Amplification plots of triplicate LAMP reactions showing the effects of different modifications (indicated by plot titles and shown in **Table S1**) to the VP28.31 primer set for WSSV.

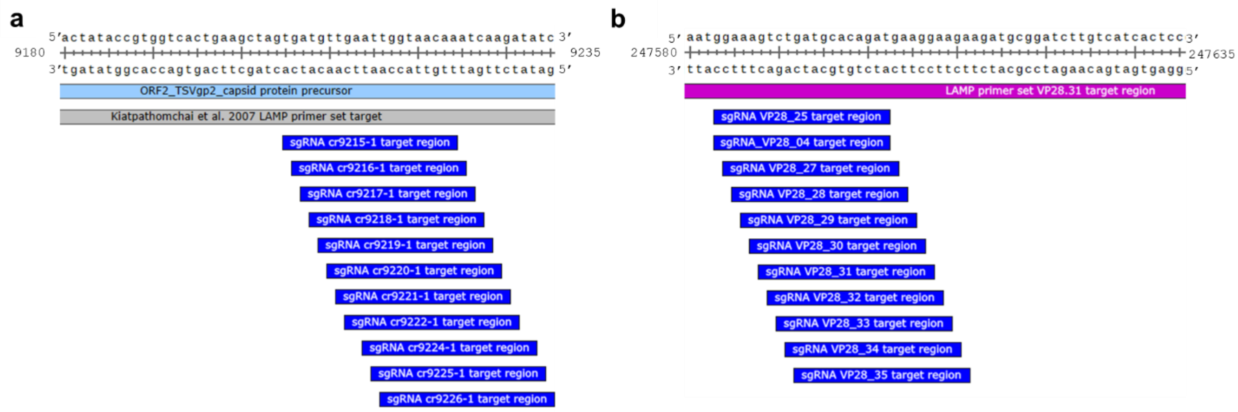


**Figure S2.** Diagrams of annotated consensus genome segments showing target regions of guide RNAs (dark blue) screened for **(a)** TSV or **(b)** WSSV.

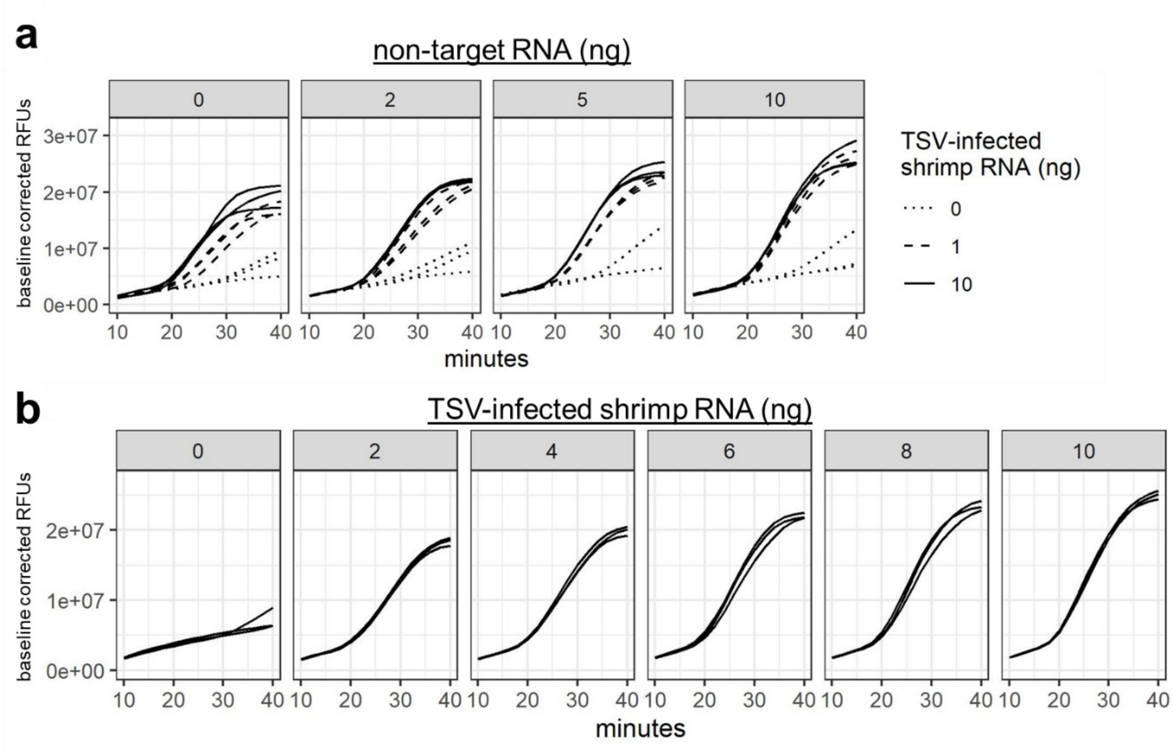


**Figure S3.** Optimizing amount of non-target and target RNA for the TSV SHERLOCK assay. (**a**) Cas cleavage activity plots of TSV SHERLOCK reactions containing varying amounts (ng; nanograms) of non-target RNA and varying amounts of target RNA (TSV-infected shrimp genomic RNA). (**b**) Cas cleavage activity plots of TSV SHERLOCK reactions containing varying amounts (ng; nanograms) of TSV-infected shrimp genomic RNA (gRNA).

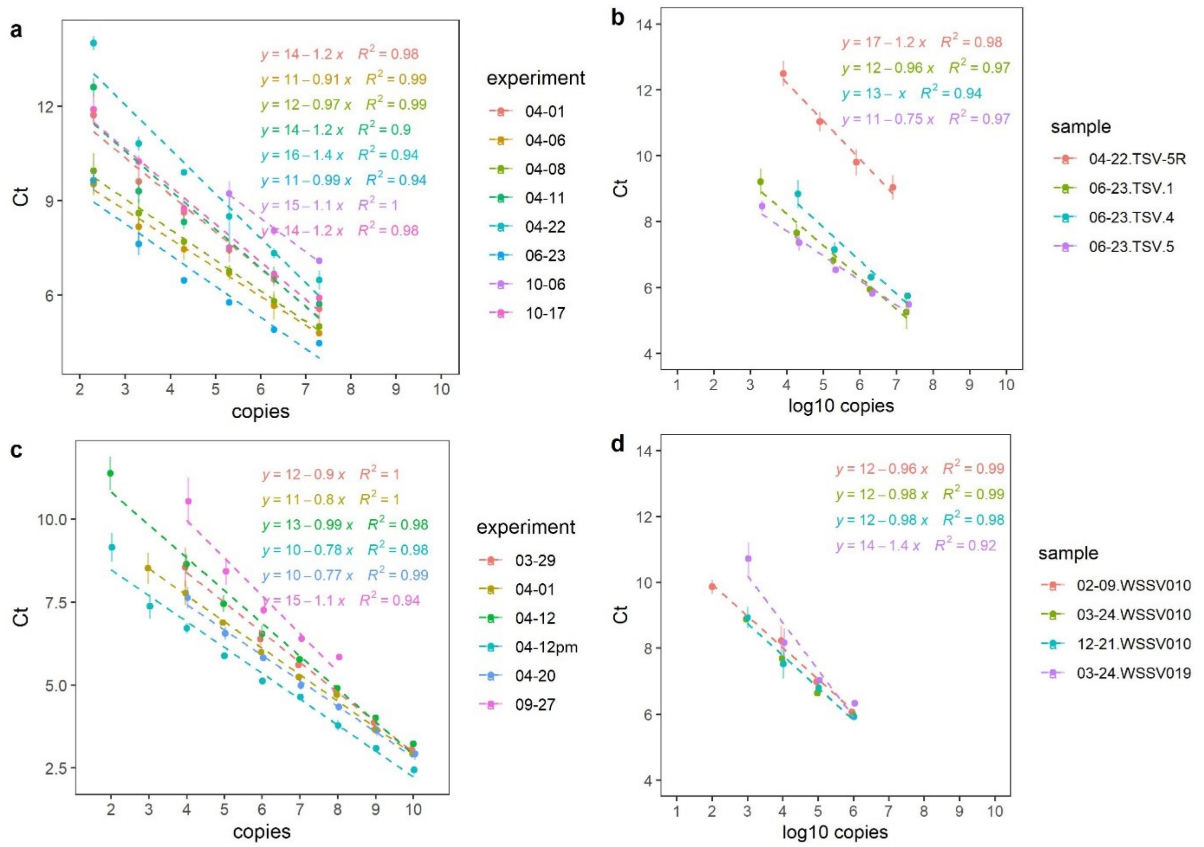


**FigureS4.** Comparison of synthetic and genomic target dilution series**.** (**a**) TSV SHERLOCK standard curves generated with synthetic target for eight different experiments. (**b**) TSV SHERLOCK curves generated with a 10-fold dilution series of genomic target isolated from samples with high viral load. (**c**) WSSV SHERLOCK standard curves generated with synthetic target for six different experiments. (**d**) WSSV SHERLOCK curves generated with a 10-fold dilution series of genomic target isolated from samples with high viral load. Linear equations are shown for each regression model (dashed line) with color corresponding to experiment number.

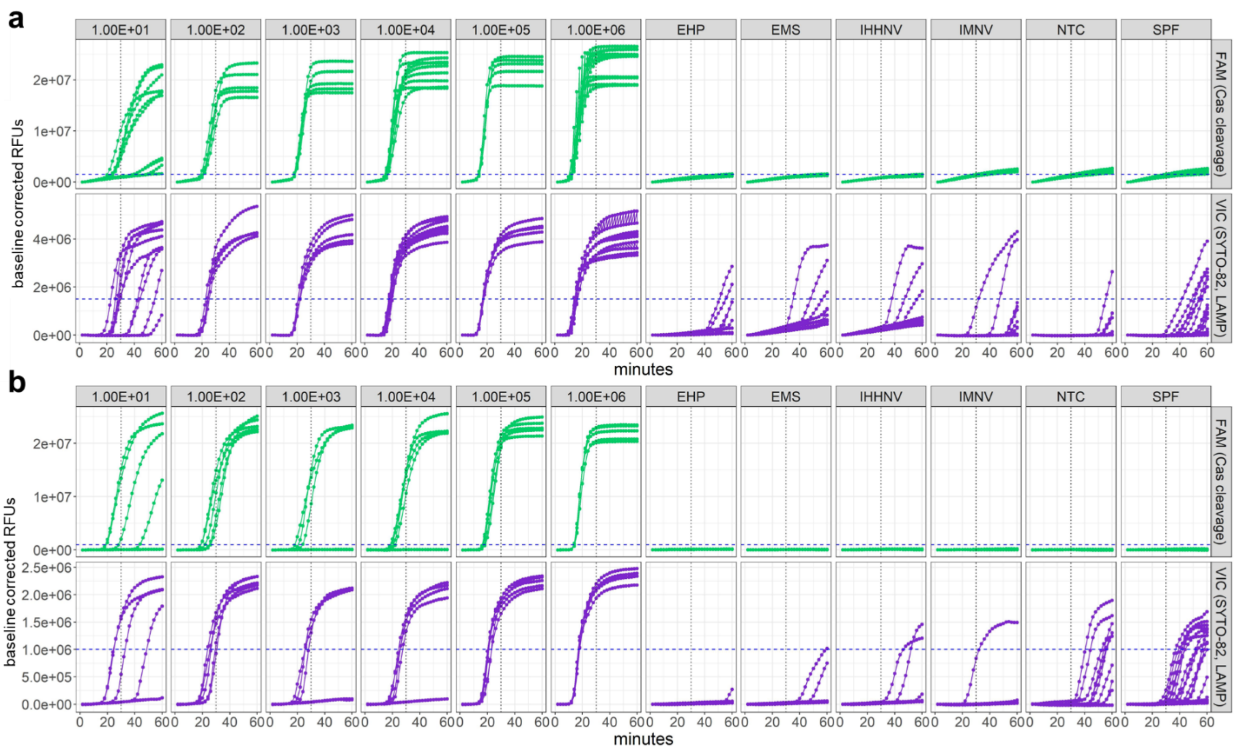


**Figure S5.** Non-specific LAMP signal does not give rise to Cas cleavage signal. (**a**) TSV and (**b**) WSSV SHERLOCKv2 assay fluorescence shows only samples containing target sequence produce a Cas cleavage signal (green) above the CT (blue dashed line), and non-specific LAMP signal (purple) produced in samples without target does not lead to Cas cleavage. Plots titled 1e+01 through 1e+06 indicate viral copies present in each reaction; 'NTC' signifies no template control; SPF signifies specific-pathogen free shrimp samples; EHP, EMS, IHHNV, and IMNV signify sample infected with nontarget viruses.

**Table S1.** Consensus genome, gBlock oligos, primers, sgRNA, and reporter oligo sequences used for developing TSV and WSSV assays. Bold indicates oligos used in final versions of assays.

| Virus | Oligo type | Name | LAMP Primer Mix Name | Notes | Sequence (5'-3') |
|---|---|---|---|---|---|
| **TSV** | **LAMP Primer** | **TSV_ORF-191_F3** | **TSV_ORF191** | | **CAATTGAAATTCTGAGATTAGAGTC** |
| **TSV** | **LAMP Primer** | **TSV_ORF-191_FIP** | **TSV_ORF191** | | **CTAGCTTCAGTGACCACGGTATAGTTTTATTTTGAGTCCAAAGCTCCA** |
| **TSV** | **LAMP Primer** | **TSV_ORF-191_LF** | **TSV_ORF191** | | **TACCTGGAGTAAATTTCAC** |
| **TSV** | **LAMP Primer** | **TSV_ORF-191_B3** | **TSV_ORF191** | | **GGTACATATCGAGCCACTC** |
| **TSV** | **LAMP Primer** | **TSV_ORF-191_BIP** | **TSV_ORF191** | | **GCGAACCCATGCGGGTATAGTTTTCAATGCGACCAATGACTG** |
| **TSV** | **LAMP Primer** | **TSV_ORF-191_LB** | **TSV_ORF191** | | **ATGAAGAACCGCCAGTTAAGCAA** |
| TSV | LAMP Primer | TSV_P123L7_F3 | TSV_P123L7 | | GTCCAGCTTTCACTTTTGG |
| TSV | LAMP Primer | TSV_P123L7_FIP | TSV_P123L7 | | ACAAATTCATGGCACCTCAACCCCTGCCGAATTGGCTTAA |
| TSV | LAMP Primer | TSV_P123L7_LF | TSV_P123L7 | | TGACGGGTCTACACCTGAAGG |
| TSV | LAMP Primer | TSV_P123L7_BIP | TSV_P123L7 | | TGGTGCATTTATCTCAACAGTTTTCTCTCTCAGAGTAAAAACCCTG |
| TSV | LAMP Primer | TSV_P123L7_B3 | TSV_P123L7 | | CTCTAATAGACCTTAGTGAGCA |
| TSV | LAMP Primer | TSV_P73L39_F3 | TSV_P73L39 | | TTCCAAGTCTTTCCCTCAA |
| TSV | LAMP Primer | TSV_P73L39_FIP | TSV_P73L39 | | ATCACCAGCACTTGCAAGCAATCAATTAGTAATTTGGAAAGGACG |
| TSV | LAMP Primer | TSV_P73L39_LB | TSV_P73L39 | | TCACTTTTGGACCTGCCGA |
| TSV | LAMP Primer | TSV_P73L39_BIP | TSV_P73L39 | | GACCACACTTTCTCTCAATTGGCCTGAAGGGTGTTAAATTAAGCC |
| TSV | LAMP Primer | TSV_P73L39_B3 | TSV_P73L39 | | ACAATGACGGGTCTACAC |
| TSV | LAMP Primer | TSV_P215L25_F3 | TSV_P215L25 | | GTTGAACAACAGTGATCCAT |
| TSV | LAMP Primer | TSV_P215L25_FIP | TSV_P215L25 | | TTGAGGTTGCAAAAACAAAAGATGGGTACATATGTCCAGTGTCAGG |
| TSV | LAMP Primer | TSV_P215L25_LB | TSV_P215L25 | | GATGCATTCAGACGGCGTTTAG |
| TSV | LAMP Primer | TSV_P215L25_BIP | TSV_P215L25 | | GCACGTATGTCCCCAAGTCTACTTTAACATCTACATAAACGCATAG |
| TSV | LAMP Primer | TSV_P215L25_B3 | TSV_P215L25 | | CGATGCGTGCAAACTCAT |
| WSSV | LAMP Primer | VP28_F3 | VP28 | initial design | CTCCGCATTCCTGTGACTG |
| WSSV | LAMP Primer | VP28_B3 | VP28 | initial design | CCACACCTTGAATGTTCCCT |
| WSSV | LAMP Primer | VP28_FIP | VP28 | initial design | GCCCAAGGTGTCGCTGTCAAACTGAGGTTGGATCAGGCTAC |
| WSSV | LAMP Primer | VP28_BIP | VP28 | initial design | CTTGTCATCACTCCCGTGGAGGAAGGTGAGATTCTGCCCCA |
| WSSV | LAMP Primer | VP28_LF | VP28 | initial design | GGACACATCAGTCATCTTG |
| WSSV | LAMP Primer | VP28_LB | VP28 | initial design | GCCGAGCACTCGAAGTG |
| WSSV | LAMP Primer | VP28.1_F3 | VP28.1 | initial design | GGTTGGATCAGGCTACT |

| | | | | | |
|---|---|---|---|---|---|
| WSSV | LAMP Primer | VP28.1_B3 | VP28.1 | initial design | CCAGTGATGTTGATCTTTCTTGATG |
| WSSV | LAMP Primer | VP28.1_FIP | VP28.1 | initial design | CCATTGCGGATCTTGATTTTGTCAAGATGACTGATGTGTC |
| WSSV | LAMP Primer | VP28.1_BIP | VP28.1 | initial design | GCCGAGCACTCGAAGTCACACCTTGAATGTTCCCTCA |
| WSSV | LAMP Primer | VP28.1_LF | VP28.1 | initial design | GGTGTCGCTGTCAAAG |
| WSSV | LAMP Primer | VP28.1_LB | VP28.1 | initial design | GTGGGGCAGAATCTCA |
| **WSSV** | **LAMP Primer** | **VP28.31_F3** | **VP28.31** | **initial design** | **AGGCTACTTCAAGATGAC** |
| **WSSV** | **LAMP Primer** | **VP28.1_B3** | **VP28.31** | **initial design** | **CCAGTGATGTTGATCTTTCTTGATG** |
| WSSV | LAMP Primer | VP28.31_FIP | VP28.31 | initial design | CAGACTTTCCATTGCGGATTGATGTGTCCTTTGACAG |
| WSSV | LAMP Primer | VP28.1_BIP | VP28.31 | initial design | GCCGAGCACTCGAAGTCACACCTTGAATGTTCCCTCA |
| **WSSV** | **LAMP Primer** | **VP28.31_LF** | **VP28.31** | **initial design** | **ATTTTGCCCAAGGTGTCG** |
| **WSSV** | **LAMP Primer** | **VP28.1_LB** | **VP28.31** | **initial design** | **GTGGGGCAGAATCTCA** |
| WSSV | LAMP Primer | VP28.1_F3 | VP28.110 | initial design | GGTTGGATCAGGCTACT |
| WSSV | LAMP Primer | VP28.1_B3 | VP28.110 | initial design | CCAGTGATGTTGATCTTTCTTGATG |
| WSSV | LAMP Primer | VP28.110_FIP | VP28.110 | initial design | GACTTTCCATTGCGGATCTTGACTGATGTGTCCTTTGAC |
| WSSV | LAMP Primer | VP28.110_BIP | VP28.110 | initial design | AGGGCCGAGCACTCGAACACACCTTGAATGTTCCCTC |
| WSSV | LAMP Primer | VP28.110_LF | VP28.110 | initial design | TGCCCAAGGTGTCGCT |
| WSSV | LAMP Primer | VP28.110_LB | VP28.110 | initial design | GTGACTGTGGGGCAGA |
| **WSSV** | **LAMP Primer** | **VP28.31t_FIP** | **VP28.31** | **optimization** | **CAGACTTTCCATTGCGGATTTTTTGATGTGTCCTTTGACAG** |
| **WSSV** | **LAMP Primer** | **VP28.1t_BIP** | **VP28.31** | **optimization** | **GCCGAGCACTCGAAGTTTTTCACACCTTGAATGTTCCCTCA** |
| WSSV | LAMP Primer | VP28_110_BIPt | VP28.110 | optimization | AGGGCCGAGCACTCGAATTTTCACACCTTGAATGTTCCCTC |
| WSSV | LAMP Primer | VP28_31mod_FIP | VP28.31 | optimization | GCATCAGACTTTCCATTGCGGGTGTCCTTTGACAGCG |
| WSSV | LAMP Primer | VP28_F3 | VP28_VP28.1 | initial design | CTCCGCATTCCTGTGACTG |
| WSSV | LAMP Primer | VP28_FIP | VP28_VP28.1 | initial design | GCCCAAGGTGTCGCTGTCAAACTGAGGTTGGATCAGGCTAC |
| WSSV | LAMP Primer | VP28_LF | VP28_VP28.1 | initial design | GGACACATCAGTCATCTTG |
| WSSV | LAMP Primer | VP28.1_B3 | VP28_VP28.1 | initial design | CCAGTGATGTTGATCTTTCTTGATG |
| WSSV | LAMP Primer | VP28.1_BIP | VP28_VP28.1 | initial design | GCCGAGCACTCGAAGTCACACCTTGAATGTTCCCTCA |
| WSSV | LAMP Primer | VP28.1_LB | VP28_VP28.1 | initial design | GTGGGGCAGAATCTCA |
| **NA** | **Cas12b sgRNA scaffold gBlock** | **Cas12b_m13-T7-Handle** | **NA** | **Cas12b conserved handle with T7 promoter and m13 fwd (5' TGT AAA ACG ACG GCC AGT) and m13 rev (5' CAG GAA ACA GCT ATG AC)** | **TGTAAAACGACGGCCAGTcatataTAATACGACTCACTATAGGGGTCTAGAGGACAGAATTTTTCAACGGGTGTGCCAATGGCCACTTTCCAGGTGGCAAAGCCCGTTGAGCTTCTCAAATCTGAGAAGTGGCAC GTCATAGCTGTTTCCTG** |

| | | | | | |
|---|---|---|---|---|---|
| **NA** | **gRNA template primer** | **Cas12b_sgRNA_fwd** | **NA** | | **CATATATAATACGACTCACTATAGGGGTCTAGAGG** |
| TSV | gRNA template primer | ORF191_cr9215-1 | NA | | ttgttaccaattcaacatcaGTGCCACTTCTCAGATTTGAGAAGCTC |
| TSV | gRNA template primer | ORF191_cr9216-1 | NA | | tttgttaccaattcaacatcGTGCCACTTCTCAGATTTGAGAAGCTC |
| TSV | gRNA template primer | ORF191_cr9217-1 | NA | | atttgttaccaattcaacatGTGCCACTTCTCAGATTTGAGAAGCTC |
| TSV | gRNA template primer | ORF191_cr9218-1 | NA | | gatttgttaccaattcaacaGTGCCACTTCTCAGATTTGAGAAGCTC |
| TSV | gRNA template primer | ORF191_cr9219-1 | NA | | tgatttgttaccaattcaacGTGCCACTTCTCAGATTTGAGAAGCTC |
| TSV | gRNA template primer | ORF191_cr9220-1 | NA | | ttgatttgttaccaattcaaGTGCCACTTCTCAGATTTGAGAAGCTC |
| TSV | gRNA template primer | ORF191_cr9221-1 | NA | | cttgatttgttaccaattcaGTGCCACTTCTCAGATTTGAGAAGCTC |
| TSV | gRNA template primer | ORF191_cr9222-1 | NA | | tcttgatttgttaccaattcGTGCCACTTCTCAGATTTGAGAAGCTC |
| TSV | gRNA template primer | ORF191_cr9224-1 | NA | | tatcttgatttgttaccaatGTGCCACTTCTCAGATTTGAGAAGCTC |
| **TSV** | **gRNA template primer** | **ORF191_cr9225-1** | **NA** | | **atatcttgatttgttaccaaGTGCCACTTCTCAGATTTGAGAAGCTC** |
| TSV | gRNA template primer | ORF191_cr9226-1 | NA | | gatatcttgatttgttaccaGTGCCACTTCTCAGATTTGAGAAGCTC |
| WSSV | gRNA template primer | Cas12b_sgRNA_rev_VP28_25 | NA | | ATCTGTGCATCAGACTTTCCGTGCCACTTCTCAGATTTGAGAAGCTC |
| **WSSV** | **gRNA template primer** | **Cas12b_sgRNA_rev_VP28_27** | **NA** | | **CATCTGTGCATCAGACTTTCGTGCCACTTCTCAGATTTGAGAAGCTC** |
| WSSV | gRNA template primer | Cas12b_sgRNA_rev_VP28_28 | NA | | TCATCTGTGCATCAGACTTTGTGCCACTTCTCAGATTTGAGAAGCTC |
| WSSV | gRNA template primer | Cas12b_sgRNA_rev_VP28_29 | NA | | TTCATCTGTGCATCAGACTTGTGCCACTTCTCAGATTTGAGAAGCTC |

| | | | | | |
|---|---|---|---|---|---|
| WSSV | gRNA template primer | Cas12b_sgRNA_rev_VP28_30 | NA | | CTTCATCTGTGCATCAGACTGTGCCACTTCTCAGATTTGAGAAGCTC |
| WSSV | gRNA template primer | Cas12b_sgRNA_rev_VP28_31 | NA | | CCTTCATCTGTGCATCAGACGTGCCACTTCTCAGATTTGAGAAGCTC |
| WSSV | gRNA template primer | Cas12b_sgRNA_rev_VP28_32 | NA | | TCCTTCATCTGTGCATCAGAGTGCCACTTCTCAGATTTGAGAAGCTC |
| WSSV | gRNA template primer | Cas12b_sgRNA_rev_VP28_33 | NA | | TTCCTTCATCTGTGCATCAGGTGCCACTTCTCAGATTTGAGAAGCTC |
| WSSV | gRNA template primer | Cas12b_sgRNA_rev_VP28_34 | NA | | CTTCCTTCATCTGTGCATCAGTGCCACTTCTCAGATTTGAGAAGCTC |
| WSSV | gRNA template primer | Cas12b_sgRNA_rev_VP28_35 | NA | | TCTTCCTTCATCTGTGCATCGTGCCACTTCTCAGATTTGAGAAGCTC |
| WSSV | gRNA template primer | Cas12b_sgRNA_rev_VP28_04 | NA | | GGAAAGTCTGATGCACAGATGTGCCACTTCTCAGATTTGAGAAGCTC |
| TSV | gBlock | P215L25 | NA | | CCTAGGTCTAGGGCGGCacaagggaccttacgatgacccttagatccagctacgatgcgtgcaaactcatctttaacatctacataaacgcatagatctaaacgccgtctgaatgcatctgcactatgaatagacttggggacatacgtgcctggatttacgtttgaggttgcaaaaacaaaagatggtgctatgaatgtatttgccttatccctgacactggacatatgtacttgatatggatcactgttgttcaacctaataaattcgaataactcaggCCCTATAGTGAGTCGTATTACTCCTATTTTTATAGGTTAATGTCATGATGGTTTCTTAGACGTCGGAATTGCCTGT |
| TSV | gBlock | ORF191 | NA | | CCTAGGTCTAGGGCGGCagatcgcgaagaatacgtctgataagcttattgggtacatatcgagccactctacggacaatgcgaccaatgactgattgcttaactggcggttcttcatctatacccgcatgggttcgcgggttacgctcatttactgtgatatcttgatttgttaccaattcaacatcactagcttcagtgaccacggtatagttacctggagtaaatttcactggagctttggactcaaaatctgactctaatttcagaatttcaattgtctcgtccacgaattgCCCTATAGTGAGTCGTATTACTCCTATTTTTATAGGTTAATGTCATGATGGTTTCTTAGACGTCGGAATTGCCTGT |
| TSV | gBlock | TSV-9807-10157 | NA | | CCTAGGTCTAGGGCGGCactaatgtctctaatagaccttagtgagcaggcatttatctctcagagtaaaaaccctgccgcgcaacgcgcggttagggaaaactgttgagataaatgcaccatgaatgacaaattcatggcacctcaacccacgcgtcacaatgacgggtctacacctgaagggtgttaaattaagccaattcggcaggtccaaaagtgaaagctggacatcctgccaattgagagaaagtgtggtcatcaccagcacttgcaagcacaaagtactcattttccgtcctttccaaattactaattgataaatcacttaattgagggaaagacttggaatcaataactaaattttctgctCCCTATAGTGAGTCGTATTACTCCTATTTTTATAGGTTAATGTCATGATGGTTTCTTAGACGTCGGAATTGCCTGT |
| WSSV | gBlock | VP28_gBlock | NA | synthetic target for WSSV VP28 | CTCCGCATTCCTGTGACTGCTGAGGTTGGATCAGGCTACTTCAAGATGACTGATGTGTCCTTTGACAGCGACACCTTGGGCAAAATCAAGATCCGCAATGGAAAGTCTGATGCACAGATGAAGGAAGAAGATGCGGATCTTGTCATCACTCCCGTGGAGGGCCGAGCACTCGAAGTGACTGTGGGGCAGAATCTCACCTTTGAGGGAACATTCAAGGTGTGG |

| | | | | | |
|---|---|---|---|---|---|
| WSSV | gBlock primer | VP28-1_gBlock_B3 | NA | primer for amplifying WSSV VP28 synthetic target for VP28.31 primer set | CCAGTGATGTTGATCTTTCTTGATGTGTTGTTCCACACCTTGAATGTTCCC |
| **NA** | **reporter** | **FAM_5bpDNA_Reporter** | **NA** | | **/56-FAM/TTTTT /3IABkFQ/** |
| NA | reporter | FAM_FluorDNA_Reporter | NA | | /56-FAM/TTTTTTTTTTTTTTTTTTTT/3IABkFQ/ |
| NA | reporter | Fluor_DNA_Reporter | NA | | /5HEX/TTTTTTTTTTTTTTTTTTTT/3IABkFQ |

> TSV | consensus genome | tsv_consensus_20201004

TTTAAAAGTCGTGCGTGGCTTCACCACGCACGATCAGTACTATCAGTTAACCACTCT
TGAATATGCTCAATGACCCTATTCAACACTGGTGTCTCTTAGTACATTATTTAGCAC
TTAACGTGCATGAGTTTTGCCCATTTCTTTCAAAAATGAGTATTCGAGGAGACGTCC
CGCTCCCCGTCTTATTTCAACCGTAGACTCGACATCTATTGGTGGACATTTAATTCCA
GTCGCCGTAAGTTGCTTCTGCCCCGCGCTATATTTTCTTATACTTATGGTTCTATAGG
TCTGGTTTAAAACGTAAATAGACGGCCCACAAACTATAGAACGCGTACCCGGAACG
CCAATCCCGGATAAGTCCCTGGATATATAGATGCACCGCAATATAAGCCTGCAGACT
GGCTCATATACTATGGCCTCTTATTATCTAAACATTAAAACCCACAACCTCCGCAGA
ACCCCTGTTGCCGACCGAGCCTTCTATGTGATGAACGATGATGGAGAAAACCGGAT
ATATTCGTTAATTGGAACTTTGCGACGGGCCCCTGCCTTTAAGGTTGGTTCACGACG
CTACAAATCACACATTCCATACAGACGTGAAGCCACGTTTGCAGAATTGTGCAACCA
GTTTCATGATAGAGTGTTACCGTTTGCTAACCCTCGGGTCTGGAAAGAAGTTATTTC
GGAGAATAAAGTGCAGCCTGATTCAATGCTTAAGGCTGCATTTGACAACTGGGAAG
AATGGCCAAAGGCTAAAGTATGTGAAGAACTATACTCTGAGTGTGAATGTGGATAT
GTGGGCACCTGTTACGTGTCAGTCGATTGGTTGAGGCCTCAGGCAACGAAGTGTAAT
GATTGCATTTTGAAGATGAATAGGAAAGTTGAATACCCATATCACACTATAGGAGTT
TCAGGAAATGTTGTTACTAATACAGATATTGTTTATACAGGATATGCAGACGTCTTC
AAGTGTGAACAGTGTGATTTGCTGATGGGTGCTTGGGCACCAAACGACATTCCAGCA
CTGACGCACAATATTCGAGCATCACAGTGTGTCCAGTTTAAGCTCCCCACAGAAAAT
TTAGTGGCACGTAATTATGTGTTGTTGTGTGAGGAAATTGAGCGAGAAAATATTCCT
GTAATTTTCCAGGATTATAGTGAAGGGAATGTCTTTACGTGTCGGATTGTTAGTGGA
GATCTAACTGCTGTTGGTACAGCATCCAATATGTATACAGCTCGAGATGTAGCGTCT
AAAAATTTGTTAGATCAGCTACATAACACCCCCAATGTGCACATGCACTCTCTGCAT
TCCCTCCCGTATGAGAATTTCCCTTGTGAAGCTCTGGAATTTGCAGTTGAGCAAGGT
ATTATCCCCCCTGTGACCTTTGATGAGGTATTTGCTAATGATGAATACGTTATTACTA
TTTCATGTAGCCTATTAGTTGTTGCTGACGTTGGCCCCACCCAAGCAGTTGCACGAG
AAAGAGCTGCAAAGAGATTTTTGAAGATGTATGATTATTCTGCAAGTTATCCTAGTA
CCCACATGTTTACTTTATCCACACTCCCCCAGAGATCAGGTGAAACGCTAGAGTTGG

CTAATGCCACATTGAACCATGTGAATAATGTGATTGACCGACACGATGAAGCAATA
AGTAATGTCAGGCAAAATGTTGAAGTGAAGTTGACGGATGTGTCCCGACAAGTTGG
TGCTATGTTGCCGAAAGTAGAAACTGCTATTGACGACGTATCTTCTACTCTATCTTCC
TTTAGGGGAGTATTAGATAAGATTTCAGCATGGATGCCTTCATCAAACCCTAAGATA
ATTGACCTCATTAAGGAGACTTTTGTATCACTGTTTTTTGCTATTCTAACTAAGTCTT
TGTATCCTATAATTCAGGGTATATCTAGTTATGCTCTTCGTAACAATTTGATGGCTAA
CCATCTGACTGCCTTGTCAGAATGGTTAATGACACTCAAGTATGATTCACCGGATGA
AGAGGAGATGCCAAGTACACACGGTTTCATGGATGACTTAACTAGTCGCCTTCCAGG
ACTAAATGCTGCCAAGGCGCAGGCTGCGACTATATATGAGTCTATAGGAACAGGCT
TATGCGTAGCATTGTCTGGTATATTGTCATTTATAGCAGTCATGTGTCTTGGAATCAC
TGATTTATCTGCTGTTACATTCAATAAGCTGCTCACGCAGTCTTCATTGGTTGGACGA
GCTTTAGTCGGTGTGCGTAGCTTTAAAGATGTTTTCTTTGGCATCTGGGATTATGTAG
ACAATCAAGTGTGTGAAATTCTTTACGGGAAAAGTCGCAAGAACTTGGATTTGTTGA
AAGAATATCCGAGTTTGGACTCGTTGTTGTCCATCTTCAACTACTTTCACGACACGGT
AGATGCTAATGTGCTCATTAGTTGTAACCGTGCAGCATGTGAGCTGTTGGTTAAAGC
AGATAACCTGTACCAAGGTTACCTAGATAAATCGATAACTCTAATGCACCGAGAGA
TTTCGTCGCGACTCAAGGAGGCACGCAATTCAGTTAAGGACTTAATTGCAAAAGCTC
AGGTCTATCTGACATGTGGTGATGGTAGTCGAGTTCCCCCGGTGGTAGTGTACATGT
ATGGTGATGCTGGGTGTGGCAAAACAGAATTGTCGATGGCGTTACAGGATCACTTTG
CAACTAAGTGTTTTGGAGAAGTACCCAAGAAAGACGTGATATATTCCAGGAAAGCT
GAAAATGAATTTTGGGATGGTGTGAAGCAATCACATAAAATTATAGCTTATGATGAT
GTATTGCAGATAGTGGATTCGGCCCAAAAGCCAAATCCTGAGTTATTCGAATTTATT
AGGTTGAACAACAGTGATCCATATCAAGTACATATGTCCAGTGTCAGGGATAAGGC
AAATACATTCATAGCACCATCTTTTGTTTTTGCAACCTCAAACGTAAATCCAGGCAC
GTATGTCCCCAAGTCTATTCATAGTGCAGATGCATTCAGACGGCGTTTAGATCTATG
CGTTTATGTAGATGTTAAAGATGAGTTTGCACGCATCGTAGCTGGATCTAAGGGTCA
TCGTAAGGTCCCTTGTGAACAGAAGATATGGCTCCATCAGAATCCAGGAAAGACGC
AGCAAGACATGAAGCAGGAAATTATTGCAGGAACATACAAGATCACTCCGGAGACA
GCTGTGTATGAGTTGCATGTTGACACTACATTAGCAGGCAATGCTCAGTCTAAAGTC
TGTGCTTATGATGGTTTGGTGTCACTGATTGAACAGGTGAGGAAATTGCGTGTTGCG
GCCCATAGCGATAAGGTGGAAACTGACGTCCCAGTCCTTCCCACTAGACTGCACGA
GTTATCGCAAGAAACTTTTCCCAATACACATGCCGGTGTAGGATTTCAATTTGCAAC
TGATTGGTTGGGCGATTTCGATCGGCCAGTGGAAGCATTATCCTATTTAAATGAAAC
ATTGGAAGCTCATTTTGTCTCGCGGAGTGCGAACGATGGAAGCATGTTCATCCCAGC
CAGTGAGGTTGCTGATCTGTTGTGTCAGAGACATAACAATACGAATTTGAATGAGGA
ACTGGTGTATTTGACGTGGATGACGCAGATCACAGATAAGGAGTTAGCCTCGAGTTT
AGTATATTTCACAAATAACGGAATGGATAAGTCAATTTGGAAACAGAGCGCCGAGC
GTTCAGCACAGGCCATTAGTCAGTGTAAGAATGCTTGGACGCGCATAAACGATTTCT
TGAAGAATCATTGGATTTCCATATCTGCCGTTATAGGATCAGCTCTCCTAATAGGGG
GAGTGTCGAGTGCAGTGAAGTGTGCAACGAAGTGTAGGGTTAGGAAGATATTGCAG
GATGGAGGTTCGATCATGCAACTTGTTGGTGTACGTTCATGTATGTACGCATGCCAG
TTATGCAAACGCATCAAGAACGGTGATTTGCGCCTACGTGTCCGAAATCGCTCGGAA
GGTGTTACTACGTTTGTACCAGGTGATGTTAGGCGAGTTGCGCGCCACGTGATATCC

GCTGCGGATGTGTGTGAAGTGCCTGTTCATCACTCATTTATACAGTCACTGTGTGAT
GAGGCTTTTACCGTACATTCGGATAAGGAAGAAACGTTCTCCATCCTTGATTTCACC
CCAGAGGCGAAAGGTAGAAACCCTCTCGAAAGTGCTGTAGTTGAATCCCATCAGGA
CTATAGAGCTAAGACTGCTGTGGTGGAATCCCATCAGGACTTTAAACCCAAGGGCG
CAATCGTAGAATCTACCAGAGACACTGTATTTACTGAGTCTCATCAAGACGTCAGGG
TAAAGTTGCACCCACAAGTTGAATCGCATCAGGACTTTAGAGCCAAAAATCCGACA
GTTGAAAGTAGAAAACCAGACTATCAGGTGGAGTGGACCGATTTGAGAACTGAATC
TTCCGACGACAGAAACGCTCAAGACATAAGTAACAGGATCCTATCTAGGAATTTTGT
GAGGTTATATGTCCCAGGATCAAGTCTATATACACATGGTTTATTCGCGTATGGACG
AATGTTGTTGATGCCTAAACACATGTTTGACATGTTAAATGGCAGTGTAGAAATAGT
TAGTATAGCAGATAAGGGTAACACTCGGGTTCACGTCAAGATACAATCCCACAAAA
CTGTGACAAGGGGTGGCTATGAAGTGGATATTGTAATATGTGAAATGGGAAATTCT
ATTTCAGCACGCAAGGACATAACTTCATATTTCCCTACGGTGAAGGAACTTCCAGGA
TTAACAGGTATGATATCTTCTGGGCGGATGAGAGTCTTTTCGACCGCTAAGTTCAAG
GCATCAGATTCATGTTCGTACTTGATGCCACAAGACTTTGTAGCCAAGTACATTGCT
GCAGTTGATCATATAACATCCAAGTCCCCAGAAAAGAGAAGTTATTTTATACGACAA
GGCTTTGAAGCAGAGAGTGATTCCATGCAAGGCGATTGTTGTTCACCTTATGTACTG
TTTAATTCAGCATCGAGAGCTAAGATTGTTGGATTACACTGTGCAGGATTCGATGGA
ACAGCAAGAGTGTTTGCCCAGATAATTACTCAGGAAGACATAATGGCCGCCACGCC
GACAACTCATGCAGGTCGCGTGACTACTGAATTTCCCCATACATCACTGCGGGATTC
TCCTCTCCCTAATTCAATGGCCATTGGTTCCGTTAAGACAGCACCCAATCCAACAAA
ATCTGAAATTACTCGGAGTCCTATCCATGGATGTTTCCCCGTTCGTACAGCCCCCGCT
ACCTTGTATAGCCCAACAGAGAACTTATTAATCAAGAACGCAATGAAAGTAACAAA
GAATGTGGAGTTGCTGGAAGAAGACCTAATTGATGCCTGTGTTCATGACGTAAAGC
GAATTTTGAATGCTCCAGGAGTGTCTGATGCGGAGAAGAGAGTTTTGACGCATGAG
GAATCCATTACGGGTATTGAGAATCGTCAGTACATGAATGCATTGAATCGAAGCAC
GTCAGCAGGTTTTCCCTACAGTTCTCGCAAGGCGAAAGGGAAGAGCGGAAAGCAGA
CGTGGTTGGGCTCTGAGGAATTCATTGTTGACAACCCGGATTTAAAAGAACATGTTG
AGAAAATCGTGGACAAGGCCAAGGATGGCATAGTAGATGTTAGCTTGGGTATTTTT
GCGGCTACATTGAAAGATGAAAGGCGCCCTTTGGAGAAAGTACAGGCCAATAAGAC
ACGCGTGTTTGCTGCTTCAAATCAAGGTTTAGCCTTGGCACTAAGAAGATATTACCT
AAGTTTCTTGGACCATGTGATGACGAACAGGATTGACAACGAGATTGGTTTAGGTGT
AAACGTGTATTCGTATGATTGGACGCGCATAGTTAATAAGCTTAAGCGCGTTGGTGA
CAAGGTGATTGCTGGTGATTTCTCAAATTTTGATGGTTCATTGAATTCTCAGATTTTA
TCACGAGTATCTGAAATTGTCACTGATTGGTATGGAGATGATGCAGAAAATGGTCTG
ATCAGACATACACTACTTGAGTACTTGTTTAATGCCACCTGGCTTATGAATGGTAAG
GTTTTCCAACTCAACCATTCTCAGCCTTCCGGCAATCCATTAACTACTCTCATCAACT
GTGTATATAACATGATCATTTTTAGATATGTCTACCTTCTAGCTCAGCGAGAAAACG
GGTTTCCCATGACGCTCTCTGGATTTACTACAAACGTAGCTTGCATTTTCTATGGTGA
CGATTCATTGTGTAGTGTGTCAGATAAAGTGAGTGAATGGTTCAACCAGCACGTAAT
AACCCGATTGATGGCTGCTACTGGACATGAATACACGGACGAGACTAAGAGTGGTT
CCCCCCCCCCATACCGCTCCTTAAGTGAGGTTACCTTTCTCAAGCGTGAGTTTGTGCT
AAGAGATCATTTTTGGATTGCACCCCTATCCCGGAATACGATTGAAGATATGTGCAT

GTGGAGTAGAAAGAATATCGATGCGCAGGATGCATTACTGCAAACAACGCGCATTG
CTTCTTTTGAGGCTTCGCTGCATGAGAAGAGTTATTTCTTAATGTTCTGCGATGTCAT
TAAGAAAGCGTGTAGGAACGCAGGGTACAAGGAAGCATGTTTACATGAGTTGGATT
GTAAGAGCTTCCTTTTGGCCCAGCAAGGTAGAGCTGGAGCTCATGATAGTGAGTTCC
TAAGTCAGCTATTGGACTTAAACTAATAGCACCACCCGATCGTAAACTCCATGTATT
GGTTACCCATCTGCATCGAAAACTCTCCGAACACTAGGTGCAGTAAGGCTTTCATGG
AGTGGTTTGCTATTTAGCGTACGTGTACCATAGGCAGCCCCAAAAACACGTGTGAGG
AGAAAGTCCCAGTCACTTTGGGCAAAGTAGACAGCCGCGCTTGCGTGGTGGGACTT
AATTAATGCCTGCTAACCCAGTTGAAATTGATAATTTTGATACAACAACCAGTGGAG
GACTAATTCCAGGAGGTAGTGTTACAAACAGTGAAGGTTCTACAATCTTGATGAATG
ATATCCCAATCACTAATCAGAATGTAGTGCTGTCTAAGAATGTAACAGATAACCTGT
TTGAAGTCCAGGACCAAGCTCTCATTGAATCTCTCTCTCGCGACGTTTTACTTCATAA
CGACAGTTGGACATCTAGTGATGATGAAATTGGCACAACTATGACGCAGGAACAGC
TTGCAACAGAATTCAATCAGCCACACTTATATGAAATTTCCCTACCTGATGACATTG
TACGTAAATCGCTGTTTATGTCTAATAAATTAGCGAATATTGCATATATGCGATGTG
ATTACGAGGTTACTGTACGAGTACAAGCCACGCCCTTTTTACAAGGAGCATTGTGGC
TGTGGAATAAGATGAATGCTAAGCAGACATCAATTATTCGACGCACTCTTACAGAAC
ACCTACGCTCTATTACATCATTTCCTGGCATTGAGATGAACTTGCAGTCTGAAGCTC
GAGCTATTACTCTTTCGATTCCTTATACTAGTGAGTTGCAGGTTTTTAACCCCAGAAA
TGTGAATAACCTAAATTCTATTCGGCTTTCCGTTCTGAGTCAATTGCAAGGCCCTGA
AGATGTAGAATCCGCATCTTATTCTATTTATGGCAGGTTGAAAAACATCAAGCTATA
TGGGCATGCCCCATCTGTGACATCTTCAGTATATCCGTCTACTCAGTCCGGATATGAT
GATGATTGTCCCATTGTGCATGCGGGAACTGATGAGGATTCTTCTAAACAGGGGATT
GTCTCAAGGGTTGCAGACACCGTTGGTGCGGTGGCAAATGTAGTAGATGGGGTAGG
AGTACCTATTCTATCCACAATTGCCAAGCCTGTTTCCTGGGTGTCGGGCGTAGTGAG
TAATGTAGCTTCAATGTTCGGATTTTCAAAAGATAGGGATATGACGAAAGTCAACGC
ATATGAGAACTTACCTGGTAAGGGCTTCACTCATGGTGTTGGCTTCGATTATGGCGT
ACCCCTGTCTCTTTTCCCTAACAATGCCATTGATCCCACAATTGCAGTGCCTGAAGG
ATTGGATGAGATGTCTATTGAATACTTAGCACAGCGACCATATATGCTCAACAGATA
CACTATCAAAGGTGGTGACACTCCTGATGCGCATGGAACAATTATTGCAGATATTCC
AGTGAGTCCTGTCAATTTTAGTTTGTATGGTAAAGTTATTGCTAAGTATCGCACCCTA
TTCGCTGCCCCAGTTAGTCTAGCTGTAGCAATGGCCAATTGGTGGCGTGGAAATATT
AACCTTAATCTTCGCTTTGCTAAGACGCAGTACCATCAATGCAGATTGCTGGTGCAA
TATCTCCCCTATGGTAGTGGTGTTCAACCAATAGAAAGTATCCTTTCACAGATCATC
GACATCTCACAAGTCGATGATAAGGGTATTGACATTGCTTTTCCTTCCGTCTATCCCA
ATAAGTGGATGCGAGTGTACGATCCAGCGAAAGTTGGGTACACGGCAGATTGTGCC
CCAGGCCGAATCGTCATTTCCGTTCTCAATCCACTTATCTCAGCTTCGACAGTCTCTC
CTAATATTGTCATGTATCCTTGGGTGAATTGGAGCAATTTAGAGGTTGCTGAACCAG
GTACGCTTGCTAAAGCAGCCATCGGCTTCAATTATCCAGCAGATGTTCCTGAGGAGC
CCACTTTTTCAGTAACGCGTGCTCCAGTATCTGGAACACTGTTTACGTTACTCCAGGA
TACGAAGGTGTCTTTGGGGGAAGCTGACGGTGTATTCTCATTATACTTTACGAACAC
TACCACTGGTAGAAGGCACAGACTAGCTTATGCCGGACTGCCTGGTGAACTCGGTA
GTTGTGAGATAGTGAAGCTACCTCAAGGGCAATATTCAATTGAATACGCAGCTACCA

GCGCTCCAACTCTTGTTCTCGATAGACCTATCTTTTCTGAACCGATTGGCCCCAAGTA
TGTAGTCACTAAAGTTAAGAACGGTGATGTTGTTAGTATTTCCGAGGAGACGTTAGT
AACATGTGGTAGTATGGCAGCGATTGGTGGAGCTACGGTCGCATTGCAATTCGTGGA
CGAGACAATTGAAATTCTGAAATTAGAGTCAGATTTTGAGTCCAAAGCTCCAGTGAA
ATTTACTCCAGGTAACTATACCGTGGTCACTGAAGCTAGTGATGTTGAATTGGTAAC
AAATCAAGATATCACAGTAAATGAGCGTAACCCGCGAACCCATGCGGGTATAGATG
AAGAACCGCCAGTTAAGCAATCAGTCATTGGTCGCATTGTCCGTAGAGTGGCTCGAT
ATGTACCCAATAAGCTTATCAGACGTATTCTTCGCGATCTATCGCAATCTCCATGTAT
ATATCCATCCACACATGCTGGTCTGGACTACTCCAGCTCAGACACATCTACAATGTT
GACTACAATGGGTGAACAGTTTGTCTCTCTCAGAATGTTAACTAGACGTTCCAGTCC
CGTTGACATTCTTAGAGGCGATTTGGTTACTTTGCCCGGAATTTCCTTTGGCACAGAT
AATTCATTACGCCAGAGCTTAGTTAACATTATTTCATATATGTATAGATTTACTCATG
GTAGCATATCCTACAAAATTATTCCTAAGAATAAGGGCGATCTATATATTACTACAA
CGAGCCCAGATTCGATCGAAACTAGTACTAGTGCTTATCAGTTTGATACTAACCGTG
CTATGCATTATATTAACACATCCCTGAACCCTATGGCTCAAATTAGCTTGCCTTATTA
TAGTCCAGCAGAAAATTTAGTTATTGATTCCAAGTCTTTCCCTCAATTAAGTGATTTA
TCAATTAGTAATTTGGAAAGGACGGAAAATGAGTACTTTGTGCTTGCAAGTGCTGGT
GATGACCACACTTTCTCTCAATTGGCAGGATGTCCAGCTTTCACTTTTGGACCTGCCG
AATTGGCTTAATTTAACACCCTTCAGGTGTAGACCCGTCATTGTGACGCGTGGGTTG
AGGTGCCATGAATTTGTCATTCATGGTGCATTTATCTCAACAGTTTTCCCTAACCGCG
CGTTGCGCGGCAGGGTTTTTACTCTGAGAGATAAATGCCTGCTCACTAAGGTCTATT
AGAGACATTAGTATGATCCGGCTAATAGTCGCTTTGGATGACCTCCAAAGGGC

> WSSV | consensus genome | WSSV_Consensus_Genome

atggcgtacattgaccaaggggcgtttggagccaaatctgtgatgcagcaacaataccttcttccaccctctaataggctctccaaacaacaa
cctccactggcatcatcttctcttcaaccttcctcttctaataaacctaggagtacatcaagagtaacagacatatttgttgtgataacttgtgttgtt
ttaattgctgcttttataagcaactcatttagtgttacaaaaaatgtggtaaaactttctaaagaacaaaccgagaaaatactagaaaaagatttg
cctgataaggtgtacaaattgtttgaaaatttaaaggatggaacatttggtataggggaagatgaggaggaagaaagggaggagagggag
gaaggggaggaagagcttgaaacaaaaaacataagacttaaaagaaagtggaaagaaatgactgaacaaggggatcaaggaataaaag
taaggaaattacatggcccgagaggagaaagaggagaaactggtccagcaggagcagttggccctgcaggccctcaaggagaaagag
gagcaattggaccggcaggaaaggatggagcagttggccctgcaggccctcaaggagaaagaggagcaattggaccggcaggaaag
gatggagcagttggccctcaaggccctccaggagaaagaggagaaaatggacgcccaggaagagatggagcagttggccctcaagga
gaaagaggagcaattggaccggcaggaaaggatggagcagttggccctcaaggagaaagaggagcaattggaccggcaggaaagga
tggagcagttggccctgcaggccctcaaggagaaagaggagaaaatggacgcccaggaagagatggagcagttggccctgcaggccc
tccaggagaaagaggagcaattggaccggcaggaagagatggagcagttggccctgcaggccctccaggagaaagaggagcaacag
gtataccaggaagggatggcgtggacggttctgtgggccctcaaggagaaagaggagaaattggacgcccaggaagagatggagcagt
tggccctgcaggccctcaaggaagaagaggagcaacaggacgcgcaggaaaggatggtgcagttggtcctgcaggccctcaaggaga
aaaaggagaagctggtaaggacggttctatagggcctcaaggaatacaaggcccaagaggagagactggaccaccgggaagggacgg
cactgcagcagaaagaggagaaagaggcttcccaggaccaccaggcgaaactggaccaccaggaaaggatggtgtggatggttctgag
ggccctcaagggaaaagaggagaaacaggacccgttggacctaggggtgaaccaggtctagctggcctcccaggaagagatggagca
attggccctgcaggccctccaggagaaagaggagcaactggtctaccaggaaggaatggtgtggatggttctatcggcccccaaggaag
aagaggagcaacaggccgcgcaggaaaggatggggcagttggccctgcaggccctccaggagaaagaggagcaacaggtataccag
gaagggatggtgtggacggttctgtgggccctccaggagaaagaggagaaactggaccagcaggaagggacggttcagttggccctgc

tggccctcaaggagaaagaggagaaaatggacgcccaggaagagatggggcaactggccctataggtcctgctggtcctcaaggagaa
aaaggagaaaatggacgcccaggaagagatggagcaactggccctataggccctagaggagaaactggtgcaatgggaaagaatggc
gtggacggttctatgggtcctcaaggaagaagaggagcaacaggccgcgcaggaaaggatggggcagttggccctgctggccctccag
gagaaagaggagaaactggaccagcaggaagggacggttcagttggccctgctggccctcaaggagaaacaggattaactggcagccc
aggaagagatggagcaactggccctataggtcctgctggccctcaaggagaaaaaggagaaaatggacgcccaggaagagatggagc
aactggccctataggtcctgctggccctcaaggagaaaaaggagaaaatggacgcccaggaagagatggagcaactggccctataggtc
ctgctggccctcaaggagaaacaggattaactggacgcccaggaagagatggagcaactggccctataggtcctagaggagaaactggt
gcaatgggaaagaatggtgtggacggttctacgggtcctcaaggaagaagaggagcaacaggccgcgcaggaaaggatggagcagtt
ggccctgctggccctccaggagaaagaggagaaaatggacgcccaggaagagatggagcaactggccctataggtcctgctggccctc
aaggagaaacaggattagctgggctgccaggaagagatggagcaattggtcctcaaggagaaaagggagaaaatggacgcccaggaa
aggatggggcaactggccctatgggtcctccaggagaaaggggagagactggtcctataggtcctgctggccctcaaggagcaactggt
cttccaggaagggatggtgtggatggttctgttggccctcaaggaaaaagaggattaatagggcgcacaggaagggatggggcaattggc
cctgtaggtcctgcaggccctaaaggagaaacaggattagctggcctgccagggatagatggaaaggacggttccgtgggtcctcaagg
agcaattggacctataggcccacgaggagaaagaggagaaactggacgaccaggaagggacggtgaggatggttccacaggccctatg
ggcccccaaggactaagaggagctacgggagctccaggaccgcaaggagaaagaggattaaagggacggccaggaaaagatggtga
aacaggtcctccagggcgacaaggaagggatggaataatgggtcctaggggtcttcgaggagaaaaaggagcacctggtaatgatggtc
tagagggacctgaaggaagagatggtgcacctggtcccgctggccctattggacctcaaggaataagaggattaaaaggtatccagggac
gaccaggaagagacggagaaatgggaccagccggcaaggacggaatagaaggccctagaggtcaagatggaacaactggcgctaaa
ggacctagaggattaagaggttttcaaggaagaacaggagaaactggtgcacaaggatctagaggagaaaaaggcgatagagggctaa
caggccctcaaggaagagacggtccacccggtgaagaaggtcctcaaggtcttagaggagaaaggggagcacctggccctagaggtcc
tagaggtattcgtggccgttcaggacctcaaggaagtaacggcgtgcaaggacctcgaggtccccgaggaacaaaaggaagaacagga
atacaaggcctcactggcatagaaggtcctcgaggtcctagaggtatacaaggaaaggaaggaagaatggggaaaattggacatcgagg
agaaaagggtgataaaggagaccgtggagaacaaggcatcgctggagcagacggggaaaaaggtccaagaggtttacgaggaattcg
aggccctattggtgctcctggtaagcctggcacggaaggggttagaggtcctagaggggtgagaggtgttcctggctatcctggcgcacaa
ggggaattaggtccccaaggaccaacaggtcctcaagggccagcaggtcctcaagggccgatggggcgtacaggagatactggtccca
tgggccctcctggagcagtgggaccaagaggagagaaaggaggtagaggaagaaagggaaaaaatggccctaaaggagcggacgga
aaagatgccgtaaatatcatacaaaaatattcaatcacccatgctcgtgcagagataatgtgggaaggaaatgaaatcggagaagcatacat
tggaagatcttatggaactgatacaatccctgtgatgatagaaaatagaatagggatgacaaatgaggacaaaaaaaacgaatattgtataca
agtaatgacaatgcactcaataacaactagaggaagaacatcgggtgtttttgtggtaagcaataagacagattatatccttttagttactttact
gatgccagaaagtgtttcctgtagaacagatgtcagtacaaatgcgaggtcagagagggtgaatgctgttagagaaagagaaagcaaatcg
tacagatttattaggccgtctgaccaatctataggtactcattcacgttcaaaaattgccgtggtaatgtatccagacgcaagcatgagttactca
gttgatacattagacgctgatgtggcgcgaagagaaacaacgtctgtgcttttattagcagaaaccatacacggggaaaaagatagaggttt
ctatgctgatagaggaactgtagggaggttgatggtacctcccactgaagaagagttattggtattgcaagcggatccagacatcaccccaa
caatcgcagacccgattacagtagcagaagttggaaaccctgtcgtgaggataaaaactccttcatgtgatgagttgaataaattcatgagatt
ctctactaatttctcatttttaaaatacgtcaaacggtttatggcgcagagagcttcatctaatagtaggaatgatatgtccgctttcttattagattt
gatttttgaaaaattgatagagtttttattatcaacaggatggaatagagaagctctgggaggaactactggggctattcttatgatcaggacgg
caacttgtcgtctagcgcaatcttattaccccctccctgaaaccgttggtatggactttttgcaaacacctgggtttgactacaataaatctttccc
taacaatgaaaggtatatatatgaatttcaagctacaatgtaaataattggttaataaaataaaggtatatttttaaaaaatgtgtttatttttccccaa
ccttaaacagatcattgccaggagaaaatcgcatacttaacagataatgcctatttgtatcatcgatgtcttcgtcaaacgttgctccaaacaca
agtgtgttgatcctatttttcttcaaaactgttggtttgaaatataatacaaatttgttagctgaaaattcccttcctgtagcatttagacctggagac
agtgaaacagagtccactgttatatccctctttgtattattgtatactcttgcatacaattctaattctttagattcggctttgtttacatttagtgtgtgg
tctccgtctccaagtaaagatgtgaggattatgtctcctttctttttttaccagttcagaatcggacgttccttggccagctacatatacctttccata
gttcctggtttgtaatgtgccgtttacaaagttaaagtggtttcctctagcaacaccgttaatggtcaagtttattatttcagattcaacagggtagg

cgtctttatccttcttgtccaatttcttgttaagttctatgttggttacaactatagatataaccacgagtatcaatgtcaaacccgccagtatagcg
gcgtaaaccccccacatgtgcattttgaagaaagttgtacaaaggtcatgagagaaaaaaacagatattaaagtttgttatattttatttagtcgt
agaaatcaccactgagatatattgtgagcagcaactgccgtttcagcatactcgaaagaagaatagttgcgatcctttgaagccggcatcttg
aaatgatgtgcagtattatttctgctagttttgttgaagtactggatagcctttactgagttgaaagtttcacgggaacagcttgaacagttgatgg
aggaagaagttcgcctgcactggcaattcacaaaggagtgctgattattgttgttatgatggtacattttgaaggtgagagcacgtctttcttca
accaaaggtgtttgtctaatgtggctcaggtacatccacgctcaccttttataccaacgtcttacagttccttcagatcttccgtcatggctttaatc
ttgttggttatccctcgatctacgaagaatctaatatctccctcagatctgttcaaaactttccacgactccattgggtccagagtataatctgacc
tttctatgtttttgacaactacaggatctggttgaatatatgcagatgatgcatcaaaatctacttcttcgccgctcatgcctgtttctagtcttgaaa
ggagtgtgtccaaatcttcctcttcttcgtcttcttccatttcttctccactatctaaaaattcgtctatatcattatcatcaaagtctatgtctatatcat
catcatcatcttccccaagagatgaagaaggagatggggttcgtgtcgggggaggtggaggagtaggagtaggagtaggtggaggagta
ggagtaggagtaggagggggtggaggtaacaaaccttctccaggcacagttcctgttcctccacctcctgtattggacccagtatttacactc
ggttgttcaaattcaaaaacttcctcctcttcttttccgtcttctacagcaccttcctcgtctcttccgatcggcctattttcaggttcccttcgcccta
atacagctccttcggtgtcatcttctttttccttctctttatctttctttgttttttttattggatgttgtagtagttttatttggtcgtgacatgaaaatcacac
caaccaccacacccacaaaacacacaaaaagtatgagagataataaaaggaacatcattattatccttattttttttacattcaattatggcacca
gaaaaataaaatgatttgtacaaacacgagttacccctccacacgcctgacctctgccgcgaattcttcatacccttccatattctttgctgcaaa
ggtcacgtcatgttgatgagctaccatttcatatcgcatatctcgccaaaaagatccatccgcaacagcggccgtgcattcttcttcaatggca
gaagtcatgtcagctgtcatttttttaataagatcacgtagtatttcctccctatacaaatctataaccttctgcctaaagacaggcctctctgctaa
aagagaaggattttctgtaagctggtccatggtgtaataaagtgttcatttatctaacttttaaaacccaaactcttctaaaatatctacagattcca
ggtactctgccgccattgttgtaatcattgtatttgccttgtctatcactgcttgtcttttatcttccacatctcccacctcgtcttcgtgtttttttcctttct
tcttctagaagccgtctaaagacagcttctctaattgaaggtattagcccctcaactacctgagaagcttcctgttttacatcattcacaaattcgg
gcgattgtagtacgagcgtgatcgcgctcttgtagatgtcttccaaaatgtccatttttgttgtttagacctagatagagagtgggagacgctatt
aattttttttgttcatgtggctcatgacctcatgtccgatatatgtattggttgggtgtaatacaagtctctacaaagcaatagaaggggtaaaaaca
aaacacaccatggtgctaactctttcctgtacgactaggcgcgtagcgtctagcaaggggaatttctctaaggaagatgcggtgttggggaa
ccagttccccattttaaagaaatcaaacaacttgtcaattgcaagacctccctcaatagaatctttttctgcatcagtggaaaaaatattcaggga
atggaacgaaagtgggggagaaaaaattttcgacatatctcagaatgaagaagaatggatggatatcatatccttagtggaaagtgtatatga
acctgtattttctaaatcacttaaacctgataaattggcagataaaacatgtctaaccgcggctgcctttgcagcactagcttctgccgtggatga
aaaattgacaatcttatcaggtagtgatgggagtgtgcttcaacgtacaacaaaggttatgaaaaaggaccccaaaaaaatagcagaatctct
tttaaataatgaaaaatggacatctattttattggacaggttaaaaacagccaagaaacttctaagcagacgaggtgcactgaaaagcgccga
aagagtagaagtacttcatcggttgaataaactcaaggaggctcctcttccccaccatccgagcctatttgataattttagtggaggaaaaaca
tcagcagtatctgctggaacagtcatcgcatcagatatgcatttcaaattggttgaacatatttttaaggtctcctttagaaaatggggtccctgtg
gagataaaactgaaagcggggaagaagaagatgaggaagaagaagaagaagaaaagaaacattccatatcaagattcgtgcttcaattta
tgaacggacacaacgggcaacattatcataggcccgaaagtgcttctgtttacttttgtgattattatgactatttggcctacaggaatctcccta
atgagtacaaattatcgtcaatgcatcctggcacattcaatatggaggatttacctttccgccctttcgcagtaccttcaacttataagacagaatt
agagtacaaaaggtttgtgcaatcaacaaatcttccccagctaagtttcgactatggggagtttttatgttactgtatcttcggagcagattggta
caaacacctgggggatgtggtagattctctagaaaatagttccatgatatcatttgattctcagacattgagtggtgtgtataagaacactgcta
attacaaaaggttggggaagaaaagaaacggaatagccgatttggccgttaggagtatggcagaatttatccgcactgaagcgcataaggc
attgacagcagaagagatggaagaagaagaagaggaagaagaagcggaagaagaagcgatggaccaggagcctgcagaagtagact
tcctttcagtgcctcatttacgccgtaaaattcgtcaagctgtttctgtgttaaataactttgtggagaacgatctttctatattggtttctaacttcaa
gaatgtgttaaccgatgatactgtatcaggaacagatacggacaactttggttctagtggagaatttgaagcattatcttcccatttattcctttca
agaatattggatgaagtgcacattcttaggaatactgatatacaaagaaccctattttcaacgcacgtgtctctgtcggataaatctccccctag
ccgtgtccgtggaagcaatgtcaactttaataataacgctgggaacatttcttccctgcaaacgtatggcggtatagaagagttgcctgaaaat
gtactagtcggtttgtccggaggatttgaagataccgacatgtattccggagaggatgttgttgtcgtatgggatggttgtgatggaggaaaag
tgctaagtgtcaccttcaattgtggtgataattttatccagctccatgaaaaaacagcagaaacttttaaggatgatacggatttagttgaacgaa

taagagatgtgcttcagactgcaagtaagaccggaaaccttaacaaaaaagcatattcaaggaagaacatctacgctgttttgcgtgaaaatg
gcattgagcgccccggggacgattttacagaaaaggggattgctctcaaggataaaacaaatcaacccccctccccctgcaagaagtgcca
agataacggttgaaggagtcaagggatttttcagcggttttcgtgacattttggagacgagggcgctcaccacatatagtgcagaaaccttca
gagatttaggccaaggcatagtaaaagagaccgaaggactgacagctgcaacagtggcagaaacatccttctctgaaggtttagctgaaag
tttaaggtctgatgcgaatctaggtctagaattttcagaggacgccaaaacggttgtattcaaaaatgacacctctcgttctttattggaagaaac
tagggcattaagagcaaacaatacttctttttcgtcgtttgcaagggacatgggcgtccaagttagtgccgatttagatgctgaatttgctgcag
agatgagagaaacataccccgatgcagcccttgaacaaaacttgaaagatctcgacaaattcgaagagactataccagaaagtcaagtgaa
gaaactaaagaaaatagacagttatttgacagagaatccagaaagggctggcaaagaaattaacgacactgaactgtcaaaggctacagat
tcagtattggggaagaaactaggcaatgcagttacagtgttgatgaacaactttggaaaggttacaattgtagtaggggcttctgtggtggcc
gggtttttaggtccagccgctgtcgccctggtgcatgcgtccagaggggcacatctcaacgtcgtggaccacaccagccctaaaggtgtca
tcagttataaaattgtggacttttcttgtgcagatagaaacaccggatgggctaagccaaccaagcacccgttcagggaagaaatagaccat
gttatcgcactagatgcatcattcttaactgaaaatggagcatatgtattccctgaagacggaggacccaaatcgaaatataaggcctacgca
ccaatctgtggaacaaaagatgctgctcaaggagaatgtggatcttgggcaacattcgacgacccgcattctgtattgccttgggtggcaag
catgaaagatttgcctaaaggacaatccctctcctgcgataaagggatgtccactttaaaggcagtttcttccgttcttttgtccataggaaagg
atgttgcagaggctatttttgaggttgcagaggacgccgtggtggggttggcgagcaaggcaatttcagctgtaataaataaccccttgttcat
atttggagtgcctcttggatttggtatagctgctacacgcctcaatccatccaactggaaaactggcctcattgtattttcaatactactagtggtc
atactgatagttcgcttctttgcagggtcgggcccgctaaccctgaattggtttggtgcaaagaattcagctaaaaggaaacagactgaacaa
ttcgaagacgggggaggaaatcgttcaaaaatagtattggcagaaaaggacaacgccaatagtaaacttcaatcgaggaggaatgaaact
gggcccatgagattagaggagcttcctgggcatgaagatttgcgcccagttttcttccctgccacaacaaattattccaaatctgccaagattct
gggctacaaatctaaacccttcaacgacttttatacaaaaataataaacacagacatcataaaaatggataggtaaaataacacattaaatgtat
attttatatatttttattttagtaacaataaagattatacatgtataaaaattgcattgttttattcctatacaccaacaatagatatagacttatcactaa
gattataagtttctgatgacgtttttccatcacccattctgattttattcctattcttctccaaaacagattgcattccgtgtccatcaacatccgtgttc
ttcttgaagatgtgggttatatatggctccgacatatcttgtactgtcgattggatagtttcagctagggaatgtttttctggtccaaattctcctctc
ccataacggatgcgacgataagcttcagcagaagccagcaggcaaaagtgggcccagccagcctcgtcagtacacatggggtgcttcag
agtttttgttctattgtgcacatcaacatatgccataaaatcggaatgggtcaagtgcatgtcctttaatttacccgtttttgacaagtaagtgactg
gcaactgtttggcagtaaattgttcattgatatgagaagtggagtatttttcttgtttagaagcatgtcgtatatagagagcgacagctaaggcag
ttattgctagaataaagaaaacaccaacaaccataggccaattcattcgtttagtagttattacaattttgttccacatggtcataatttctcagaaa
agttcctcaacaggggaattatagtacgtgatagaattgcagtgataggcacaggcggatattccgacggtcctcttccgagaatattttgtct
agattgaagcattgcagcaatctcctgtttagcatggacaacctgtatatcatcacgaggcaaaagattgtcctttgatccttttgataaagcttc
gatgtttataaaggaacctaaaagtttccacattatagcaatgtgataaaagctgtccaacgaagcatatacagcagctgggaaagttgtccat
attaattgcccactttgtaacgcagaggctaaattttcaaccccgatcttgttgctttggcagatgttcgccatccaatctagttttgttatcgtatct
agaactggggtattcttagccagtttgactgcctcccctttcggttctttggggaagtaaactggctccttttcagtagagttggtttttatgtttttct
ctgatgcagattctccaaaatataacagttggcgaataatggagtcaacttcaatagtatcactcttcactaattctatgtccctttcgattagttgg
cacacgtctttatctagcctcatcctgttagagtttcctgtttcggctgctggtgctgtcgcagacgcagttgctattagtccatcagatgcgtcca
gtatggaattgggaggaggagctgcaacagataacatggcagtaatatcatggatgattttggagtagtccagttctaatacatcaaaatcctg
ttcttccccttcctccattttgttggtgtccatgtttgaggagggatggttgttgatactacccgagctcatgccgtacaccgaatgacgagagg
aggtaggagcaggccctttttatagccctaaagggccattatcgacagtactattatagaagttatttttatccctgatattatacaagcagagaa
aaatttcagtctacttgcaggagacatctttttaataaatttagggagtcttctccaagttttcttcttattggttttctgtttcttgtctgttggtttttga
gtcttttgttgtttagcttatgtatttggggttgccgattacggtgcctcttcttccttttattcctgtcagcaagtaaaatatccttttcgtacttgataa
atctatcaacgtactttttttattagcctgtttctggtcgtggaattccgtgaagaagatcgtagaaagtttgcatcagcagatgtggattctgtatct
gaatcttcttcctcctcctctccaagaagagaatcgacgttagtgtctgattcagagtcatattcttcttcttcttcttcctcatcgcttgaaggg
atggccgaagacgtgggagaagaagaggaggatgaggaagaagatgtagaagtagaagccatttctttgtttgttgttgttacctgcccatg
ttcctcctcgacatcatcttcttcttcaatcttattcagtgcacggttcaattgctgctgataatgccttgtgtttgttacagtgctagaccataaagc

cctcctggaagcagccgatgctccagggtccatttctgatcgcgtacatctaaccttagcaataaccagaaatacgtgcttttttttactttcag
ccagccgtgttagtcatctacggaatggtgtctttgtctttagatgaggagtcgtacaggcaaaatttgggtaaaccttcaaactcgcctgtgca
gttatcaaagcatatttgtatcaaagaaccttccgcacctattaaatttccaaccataaaagagtctatgtgttcattaacaaaaacactattttcttt
agtgagtgttagtttctttctattgcacagtattatgtccacatcgtgcccgattgtgttatttttatcattgatgacagaaatgtcgtaagcactttct
ccttcttcttccccccactccctttctgcttgttcctcctcctcgtcttcaaattcttcgtcgtcgtcttcttcttcttcctccccgccaaacctctccct
cgtttcttcctcttcgtcatttattatatgtctcctaaaattcttgtccaataattttgtaatgaaaacacactctggcttcttcctgacctttaagaaact
tgcatccatcatgctgttcattctcatatcacaaaggccatcatacctatcatctctggccatgatcgctttgacgctcaagaatatatcgtacata
atcttagactctttttttaggtgagaaaagttggttttggaactgtctaaaattaatgaaaccacgggacaatttcttagccaaacccagaagatga
gaacctccgtacccttctctgtgggaatgtgctagtgcagcattatgtatagtaaaaagcaaaccaggattttctttagccaaactctgtataaca
ttagttgccgtcactacaggtatgagatttgtcatatctaattcaggtacatctaaacattcactgataggaatcactccctctttctctccttgacc
aacaattctcaccgtcactagttgatgtgcatcttttatggctctatcataatcatgggggaagtttgatagagctaacgttgtagcattttcttcttg
caaaggtaaacacccttcgtcttcatcattgtcgggaacatcttcatcaaagtccatgccgttattgttattatatgccattatattagagaaaacat
ccatctttctattcctaggaccagaagggagcattagagcagttaatgaaggtgcatcactttctattttatttgtcttggaagtaacttcgttattttt
cgtattcaaagtgtctatcagggcagatgctgcatcacgcactcgcttctcgcgtgggagaagattgtcacaatcagatttaaagtggactaa
atgcagacacaaattagaaaaaacgaattgagatgcggatttaacaatggacaatatttgattatgaggaagtacagacgtgaaagaaagag
catgcccaagatcgtctgatgatgtgattttattttgggcacctataactactagcgctatgtgcatcgcgaggacaattacttcagtgaagatac
cattttctaacgtcaaaccttctacgaatgcaacccctcctctatttgtcctcatcatcatctttgaaaaatgttccatcaagtgtacctttgtcatact
ggctgtgacaatctgtacgatattccccatagtctttaggactaacccttgatcgataaattgttccctcataactgcggcagcaatatctatgga
attattactgtatctatcggcatatgttcctgcctttgaatctttgatcctcttatccaaactatcaaaaaatcttctctctgcgggcgagtctgtaga
gaatatgaggaactgagggactgaaagttgaccgtttctgtgagtcctaccgatagtctgtaggaaagcaatggcgttataaggtacatctaa
aacgcaatggtaacgtttagcgtacatggagttattagaagagtcatgtaaagagagaccggtattgcctttaggtcccagcattataacgtcg
acttctttcgtgttgttgaatgcactaatacatttgtttgtgttggcagttttagtgttgtttttaaccagaaacaatcccctattcgttatacgcgaac
acaatttcctgtttgttatttcagcgttactatcctcccctattgattgtgtgattgaatctatagggttgcctgctagcgccatgagcggttccgtgt
cagggactgaaggcacaagcataacgtgagctggggaagttcttctattggttttattctttaaggagattgtgaaagctgaagccatcatgac
tgcacggcacaaaagtctatagcctgagaatattgtattagcaacaggcgatgaatcgaatgtccccacgtctacgatgccgtattttgcgtca
ctttcatccttggcgatcgtttgtaagatgcgtgtagtcaaattcttgagaaaagaatccccagtttgctctagtgacatgaccatctttttttgattc
gtttgttgtccttaaacgcttcactgcatcggtgacagaagttgccttgatattcaatagtgcacctttacatgctgccacagaagtacttgctgta
ttaatgaagtattgtctcctcaacttttcaaaacactcgtcgtcaaatacttcaccccatccgtcttcttcattagcaccgtcgtcataatcttcgtta
aaatcagttttacctttggaaggtttttcctttttcttgatagcatcgtataaatcttgagcaaccatatcaatgctgcttatgtcaacttccggagac
gttattattcctcctcctccattagaaagcgcatcctttttctgtgagtaggtttttatagcatcttgtactatacttcttagtgttgttttcccatcatgtg
cgacatccctatcgcgcacaaccaccaccttaaaacgcctgtcgatattttccattacttcatataaatcatcatcatttgcagcacgtttcctcttt
ctggacggggaattaatctctgctactaatttttcaacatcttcctcttccaatattcctccttcttggtgttctcgtataatctttgtgaaggcccttct
gacatgtcctcctactttttcagaatcgatgagtacttgtctagcgtttaaacagtacgaagccaattcgtctatagcgtatttttgaagaggggga
ggctttacaattagtagtagagcagtcgactccagccatgctaatactcctactaaccaattgtcctctatttcttaaaagtttaataacctgttcca
gaaacactgtcacgtactcgctgtgctcagccatagcatctggagtagcactggacacttctttgaaggcttgtatgctcgtgtaggctggcg
cacttttccttagtatgtggtcaatcatgtgcaaatcagcattgctctggaatggcgtagcgctggccataaccgtgaacgtatcatactttaaga
tctctaagaaaatacggaacgcatcagcttgagataatagtttcaagtagttggacattgtaactcggcccttattattggatgatttcatgcgctt
cactttaaagtttctgtcgtcactcattgcactcttgaacctattaataaggtctgacactgaagggtttgccctttttttgtatgttgcgcaaaactct
agtatcttcttcctcccaagtctttgccagttcctttctaaaagagtcactaatattttgagtcttgtggaactcgtcacataataatgctgtgacaaa
gttgtcaatgggcataaccttgttggaatcgaggtctgttcctcctgttataaatttcagtacaaattccagattcgtgcgtaaatcagagtacgtc
ataaataaaatggtgggcgtttctgttaagaaatctcctattgatcttgaagaacgcttactatgaaattccttaacatccctgagagtgaaaaat
ctaatatccaccgagtctttagcagacgatacagaattttccacatctactgtcatggacccacttttcccgtcgctcttgaaagacgttggtttat
tagcacaaggcacttgtagttttctccatgaaaatttaggatcacgaagacctctagaatttgtaaccacttctcgcatgccctgttgacagctgt

tgaaaagaggcttagagcacgtcaaccatataaaaaaggggtgccttttccacacgccactgttgatggccgtaattaccttgtccgaatcttg
accaaaaattgatggccctacgtgtttttgtacatctaagagtgccttttcttgtagtatgacactcatgacgaaggctgccaattctcttgtttttcc
tacaccagtaccatctccgatgataaatcctccactcgttttcttgtccactttaaaattcctttcgaccttatctttcacaaggtctataatggacat
gttcttcttccaccctctccctttgtagaattgaatcagggctagacgcactgctagtgcctgcacagaattaatgtccatcggccgaggaaga
aatttggggtaggcatcggttacatttttgcagaagcccaccgaccggaatgtttcactctcataaagagcgaatgaaggttctgggtgggag
agggtgccaatctctttagcatcttcttcctccagaaataaaagtttgtgagccatggtaattcaaccagaaaatcaaccaatatatgttgtaaca
agaagttgccttaagaagactcttgtgtactttttttacgatacttgaaacgagtacgcctgcacggatcaaaaatttggtatactggttagctgct
atttttggcaaaataatagtttttattttcccatgctaaagagaaccaggagttacgtacagccatactagtctctacaagtaatcgtctataagcc
tcagaagtatccacaccgtccctgaaccacgaagtttcttgcacgtccgccttcttatgcccccactcgacgagtgacttcttgaatagagaga
ggggtgttattgtagctccttcgggggttgaggaaatgtttgacgcatcaaacgctagacgagggtctgactgtactaccgggccactcttgg
ggttatgatccaaacgaaacgattgcaaaccccacgcatcgtcgtcaatatccactctgtactcggatgaggaagatgatgttccgaacaattt
ttcacttccgggagttacattggagcaatcaaaatagctggggattgtttcgctgtcaaatatcctattgtggtctactgaaaaaatatcgtcggt
agctccttcttcgggaagaacagagtaggatgtggggaaagcgtactgcaaaggattggacactttcaatgtactggaagtttttgatgcagg
ttttccttttaggtaatctaaatcaaacatatctcgcgctagcacagtagcagcataaggtccgtatgaacgcattaataacccctctttcaataca
aaagtatcctcagatttttttatgatatgtcttagaaattctttcctcttcattaaactatcccaagaacttgatttgagatatatttcacttgtcctcatc
cctctcgcgtgtttctttattacgtattctgtctgagacaatctcctctcatcgttaaaattacgttcaaaaaagtcccttatttcttccagcctttggtc
acctgtagaattgtactctggagctggagtacctccaggggggagttctgatataagaggagggtttattttgactcaaaagtgtagagaataaa
agttcagatccgccgtgaacactacccgaaaccatgcgactatttcgcaaacattctgtggccaacatactatgttcatagggtttttgaggtat
gtttccggccaaaacattagctgctggctcaatgtaaaagggtgcagtacagaatggcctatgctgccccgaggaaggtttgttcaggacgg
gagctaaacaacttctatgtaacaatgaagacgggcggatgattacatggggaatattttctagaggaggtaaaacagtgctaggtattggta
caaagagcacattcttagaagtatttgttgcatccaggtttaaaggcgtgatgatgggcgcttgggccagtgcaattttcttggcagtagtttcgt
cgatacttgatttaggtctttgctggtgaatctggtcccctcttttagattggtcggccaagaaaattcccatggcaggaacagtccccactaac
ggtgtcctgtcatcgcctattattgcatgcctgtttgagtcttggggattattaacgcccggtacacacttcattgtgctcacatcaaacgtaaca
aacgaagggtctatgttgggcatcataacacaggggttttccatacattcatacatacctctgaatttcatatcctttgttacatcttgagaaagaa
ggtcgttatttagtacatccttcttattattacagaaacaggtgatggggtttttgtataaaaattcaggatcgttgcatttaattttgaattccgccaa
ggaagaagattgacgcaatttggagatgtaagaagagttgcggaattctgtatgattccgtatttgttgtttctttaatacatcataatagattggt
gtcgaagatgaaggaggtggtggtcctttgtgggttgaaggattgaaagcgcaagcagtcatcgatgttcctcccgaacctccgaaatatgtt
gggtatttaggtctgcattgccagcccccatcatctgtcagaacgactatagatgtgtgcacagaacactctttaactacacgttctccagtggt
tggggatgtttctatgtggtcaccgggcacgtttcctgaccaacaataaggtttaggatcttctccagatttatgtgtgataaatgttcctccagc
aacaaatggtgtcatcacacgaactgctctcgaggtataacacacagatgtgcaagtttgtccctctttcaaaggaactgggttggaacagta
cccgccagtattgtaggccaaaatatccacatcttcatcatgtccttctgatgtagatggaggaggaggaggagcgtcaccacttacgcctga
tgcactatggtgatggatatcatgtacaccaaataaagaatcgtacaattcagtatcgtctttttctatattatccttcaaattgaccatcctttgtcta
aactttttccccgatacagcatctagatatttgttagatttttcgatggtggcctcactcaatctatctatatggtgttcaggcacgagaagagatg
aggatctatttgcagctaaatcatgactttcgaaccgagaattcctttccttcgtttctgacggaacgttccatttattccctttaacagtcaaattca
tcatgtccaggagagcggggtgaggtgggtaattgcttgcatcttccaccccgtcatgcctacctctggcttctaggtattcttctagtgtcttttc
agggttagctccttgaactgctttcaatttacttttcatccctaattcgtgtatagcttgaaaattaacagacattggtttggctacattataggccat
tgcagcgcagactagcgcaaaaatacagcctaacacgaacacggtcacgaataacgttatttgcgcggtgaagcgcatacttcggttggta
gggtaaagtaacggatctcttcgtcgttcatccatcgttcctgatattacaatggatatcttcctagataaatctcatacaggtgccacccttttag
ggtcttataagacgcagactgtgaaaaataattctacgttaaaaaagcttaaaatgggcgcgcctaccaacgctgattttacacgcacggtatc
aggagtagcttcttctctctatcttgttaaccctggagctccttccgatagagaaaagttagttttggcctcttcttattctgactcttttgtgtacaac
tacaaggatgcagtggtgaccgctgaggctcccaagtggtgtccctttaacgagccagctcttcatgagcacatcatgaacagacttgaaaa
agctggtctaattaacagatctcgttttgtgtgtaaccctgttaaatcggctggagaagtatgcggatttcgctattctggaggaagtactcccca
gaacttaattttcccgattggagcatcagagataaattggttggtcatacttagaaatgctgctcgttttggtacagtggcagcatcggccaaag

acgccatcgaacgcattcctgatctaagagaaggtggtacaagtaaacatgtggcaaagaatgcaatgaggagacttcgtgtatggcgagc
ctttaactggatagcggaagcctccagatcagcgggcatgattcgctacgaacccttttcggttggctgtgctctatacgatcacgaggtaag
aagaaggcaactaaaaggaagttatgagcgaaatgtactatttcttggtgaccatttctgtaaagaatcaactcttctggctgatatttcaagag
ggggaagaagttctgatttctggacgatcgttgaagctgtcatccggtacaagaatagacatgctcgaacaatcagtaatgaaactaatgcca
tccctgaagactcttctataaacttggagtgggaggatgttctagtaaaaaaccaacgcgacacaaatcaaggagatgattctacgttagaaa
agaccttagaagctgccatcaaagaacatgaaagcataggagaaaaaaggaaaaaacatatcctagagtttattaaaacatgtctaacagag
gagcaacgagaaatgatttttaagggagttggaggaaaagggaatttgtccccggcccatctaacaaatctggccgatgcgatactggctaa
taatgccaaagcaggcatatgggttattatgcagagtcttttaaagcaaatcaatttctctatactccatttaatagggtacgaagctcaacgact
tttaatgttcaagttgtatatgcctgcattacttgctctttttatatctcagaggggaattggtgacgtgtttctgaatggtgtgtttaacctagaagtg
atgaaagaaagagctgcaaatgcaaaaataagagacatggtttctcgagacgcctacaagaacaatactaacgagtctaataatttgggcaa
ttattcccaattcgatatcgctacaatttatggtcaaatgtctgattgtaatgcaattcctctatctattaatattggagtacctctccataaatcgag
gatgaacatgcaagacattgaaaagaccatacaacacatagtggatatgggtttggcaaagatacgtgtagcggaagaaaatagagtagct
agacgtcttttacatagaagaaaaatgcaagaaaaggcagccagggagcgagcagctgcgagacagagaagactaagaggtgaagaag
aagaagaagaagaggaggaggaggaagagggagaagaaatggaagaagaggaagaagaagctggaacatcaggtgttaatggttctg
gttatgatcaagaagaagaagatgatggtgaggaggaggaggaggaagaggaagaagaggatgaagaagatagtgagggtgaagatat
gaatggggaaaataactcaagaaaacgtaaaaacactggtaatacttcctccactcaacaacctccccaaaagcgtcagcgaggtaaaaat
gcgcccatttcaaccaagggaaaaaggttaaaggaaagagataatattggaggatttttattagccacaattcaaaacgatgaccgccaagt
gaatgtggagagcatacaaaaacttttgacggccaggcaaagaaaatatgttaaaggtggaaaatgtggagataatggactaccagaattgt
taattgaaaaggttacaaatctactagattcagtgttccatttcagaaaggggtctattcttaacagcatacacgcaaataggcgatcagaaact
ggcgtatacaccacaaaggcaaactgtatttgtgattactatgaaagaaacgtatcaaaagacagtagtaataatacaccccattcatctgaat
gtattgagcgggcaagagaaagggatgcttcctgtgccgaatctaacaaacgccccttgtcctgtagactctaacaaccctgaagatgtggaa
caacgtatgcgggaattgatcatggaccctccttcattatcaggggtagaagattctctcgctatcgaaagagtacttcaaaatgagatattatt
cacaagtcttgtcaccaatccaattttcaatgccgtgttaggtgctgaaaaaggagatcttggaagatttatcgtattaaataacattgtaaaattt
atgaatatgactattgcttgtctagtggacggagatatgcctatgcttctagactcaagaggcaagacaaaaaaccttctagaaaagggtaca
gtgaaaaacacgagaaaatttttcaaaccaaatatgactgcagcagaattgaacgtggccactgctcaatctgcaggacaccaatacatgaa
cgctggtcattgtccagaacctggaatcaagcaaatgttactccctgattgtataatgaaactgaaatccatcgccatggaaaagggtcgagg
agggagatcggcccttcacagacagaaatgtgaccatgccttttgcaagatgctgaagtgtttattcttcaatattgacccttcaaatgctgcag
atacatttattgaccctgcgtcacgtgccaccttatttaggcttgatgatctttgtagggataggaaaaaatacaagaacattgactgggtaaag
gatttactagaccctgtaatgaagggaacaaataaatgggtgggaactggagaatacactaacattggacgtgactcaaacgtggccgccc
ctgttgatttttacacaattttgaaatacacaatgattgatgatggtgtaataagtgttccttcacggaaaccaaatgacgtgtattacagtactatt
gaaagagcggatgacttactcaccgagagcagggacgcttcatgtgaaagctaccgcccgaccttgtttgacgccagagcggtgttggaa
gtcaacggagacggacgtgtcccttacccttccagtgaaaccgtagaagacttgggaggagaagaagaagaagaggaaataacaggcgt
tattgacgacagtactgaaatagaagacgttcaagtacaggacagtaatctattcgatgtggaattattcgatatccctgaaatagaacaacat
caacagggtggagaagaagaggaacttccttcagcaatatctgaagtgtttgcttcattaccagctgataacgattcctcctcacccgcgcat
attccctcatttgggaattctgaggaggggaaaaaagtccagaaccgtacaatatttttgattctgccctcgaccaattattggatttaatagat
agcgatggaagaaacaataaccctaaaagagtcgactggaacagtgtcaccattcaagaatgaaaatattaccagaatagcgtctaattacg
tccgagcctttactgacacgtggtctcatttggtcaacatttctggagctcctctaactgctgagaagaatccaagcgctatcccagctaacga
actgaatagatactggaccaaaactaacgtgttatgcaacccactctttaaattggaggaccacataacgagggatgaagatactggtacaat
aacactaaaattcaaaatgtatatagatgataaaaatggactatatcagtctgccgttttaatgctggctctcgattcgttcgtttcgcttgcatcttt
ttcccatggagctgatttagtttccaataaaagtgaaaacaaattctgcgtaaaaattccccacgacactcgcgctgaatctttactgaataatgt
tggattccccgctggattaagtgggccttttaagagatggagcattaactacaaggctgcaaacttgagcggtaaaagcggtatagatggcct
ttcaggttccatgttgacagtactaaaaaataacaccaataaaagagcaactgatattttacatttggtgaataatgtttctgcctctgcacaaca
acttgacgattctgaaatgtctcgtacttttaaccaccaaaagaaagttggagtttgttatgatataaatgtgtccagttcaaggcaagtgaacca

gcgtaatttactccaccatcagaatataataggacaacatctgatcgaatttagaaccaagcaacttgaacgcgcacaaaataaaaaagtcaa
ggaagaggaaaatggtgagcatgaagaaatgacaagtgaggaggaagaagaggaagatgaatatgaagaaggaggttgcctatcagat
attgacgaggaagatttctatgaggatggttacgatgaagaagagggcgatgacaatagaactagaaagaagaagaaaatggaagaagat
gaagaggatgaagaagaagaatatgacgatgaagaagatgaagaagaggcagaaacttgtggtgctaatggtgttattgattgtgaagacg
atgcaatcattttccccaatggacaaaattcaaaaaggaagaaaaatggtaaaaaaacaaacattaaaaagcggtcacggaggaaggggg
agtgctctgctaacactttatcctttgtggaaaaaatacgttggaaattgtaagagcctaggtataaagccagtagggtgtccgccccccatccac
tgaatttacttccctatttatgaaggggagtgaagctgacagctgttataatacttgtcagtccaccagaggggctagccgtataaggtcactac
tcaataaatactctgttaaagatttgatgcaggtaaacagcccttcgagttggaaatgggctaaccctcccgaccgacggttcgtgctgtttgat
aagaaaactaaggaagaagttgaggttaagtttgaaattgaatgtgaaaaatccgagtattttgatgtcgtatctgaactccctagtaatattaaa
gtatggttaaaagagacggcaaaaataataaaacatttggctctgattgaagactttcttccagctatgggtgctgctaccccaaaaattcccct
caatttgattaaaactatgacgagcattttctctgttagagatattgttggatttaaaataccagaagaagtgctcagttttattcctatagaatgga
agacatctatttctgcaatggggctcctctctgtacaatttgatcgtataatagaagtgatagatttaatgataactaatggcgcctttgcgacgtc
atgcttgaacaacgcattcttcttggaaagaggagtggtgcccagagatgggagtaacacgtggctccacacggaccttgtgcaactctcca
cctccatatttagaagtattcgcaacagaggagtgaatattggcggtaacaacaacactggtagcaattcttctagttcttcttgtggagggaat
aagggcgattatggagtacgttgtggattgagtataagcaagcgtggtataaccctgaaaccaccacctgcagcgatgactaattcttcttcc
ccgtcatcctcggccatgatctcattgcctcagcccacgcgccagagcatagatctttcgataacgacaatcatccaagatttctcagaagttt
ctgggaaattgaggcttaatggattacagaaaaacatgtctgacaagagcaaagacgtgtttaatgatgcaatatacgactctggcgcattca
aggcgctcctaacatgcacagtcaacgataaaagtagacgtaaaaggaaaagaaggactttattggcatctggagagggtgtggtacgaa
gaaacctgatggtgagtcagggcaatgacgtcaatgatgcccaccagttccaggaagaatgcggaataaaaattgggggcggggcttcta
gggtgtataaaagagcccagcgccgaggttcggcagtcagttccagaagaagagtaaggaacaaaccccagtttactatagcagtctctga
cgaagacgacgactgcgaagaagaaggcgacttttcttccgagttaaatccaacgcactctcaactacttctattccaacaacggcaacaag
atagctgcacagaagacgacgacgttttagtgtctgtagaagaatataacaacagagtaagcggttcttccaccacagccggagacagagtt
cttgcaaaggatcttctctctactgtatctccgaacgaaaagaggaactctgccgccctcgccgcactcaccatatcccggcactctcttttca
acgctctatctgcaaaaacaaagttgggagaaaatggacgtttcttcctataagagcactattgactaccacaacattgaagatatggacgatc
tccagcgcgccacctacaaggatcgtatggagacggaattggtcctcgagatggctaagaaggagggaaggtacgtccgatcgttggcca
ccatggacgaattggaggtacctgaagaaccagccacttgctacacttgcggctacacctttattagacgcagggcacccccaccaaaacg
caagtcaatattcagagagccttgcgcttacccagaacttctccccgatgcaccatcccccgtccgtttagaagagcttgtcgacgtgccaga
aggagcgagtttttcacctaccctccctacgacgacggatcttctacatcgtcttcacaagccgaatgtgaagatgattatcctccaccatacg
acccatcagaaaatccacagaggtcccaagtgtgtgattattgtaccacacgtcaagtcctcagttctatgacggatcacgccagggccaac
ctcataaaaaaatctgaagagggagaagaaggccctgggtcttggccgtcgcaacaactttagctactagggttgtacaagaagaaaatagg
attgattgtatcgatgacgaactttgcatgtttttgaccattcttttgcttgctgagtgtacatagctttatatttgccaataaaacagaccccgaatg
tatttttgtgtattctttattactgcctatttttatggtttaacggaaaaaaatttttgaaaagtttttgagatggagatgaagggaaaaagagggcgct
agttcatatactgcccaataggtggtcgggtccagaaacgtctgtctgttccagaaatgtgtttagagatttctggacaagtcatttctggaaag
ggttgtaatttattatagttggtatattatttctggtatatatttctagcccctctctgtaatctgacgttgggtcgacccagcggtccaccccttcgaa
cttgacataaggcctagtcagcggtccacccccctaaactggagtgagctgaaaaatttttgaaaagtttttgagatggagatgaagggtaaaa
ggctagtaatagaaggttgtgtaactcggcataacccagaaacatatatcaaaagggatggaaacaactattctttttatacagtagttttattgt
accaaaaactcagaaatctcatcacatgacaaatctgaatgtcccattccgtggccctcaataacgtacgacactggacctgggaactccca
cctacatgttttacatacagaaaattttggcttagatgcattatcaacaaacacttttaccacattggcattcattctcttcagtttgtcacacatcca
caaagaggccatattctccatagaatcagatccatcaacaaatatatagaatgttacacccttgtcgatccagtccatggcgacaatagtaaag
tcttgtccaaaaacactgtttccgaagatttctttgtccccatccatgtattgggaaaaagtacagtatggtccacagtccttgatctcctccaacc
agtcgttcaaaaaaatccctaataaagaaagacaggcaattcgacttgatctggaaaccaatgttgagtttcttcgtattttcttcattgttggagg
ggaagaaatacgccaattgtgtatctgcttcaaatgctccacacttccctttatggatcaggcggttctccataatctccttgccagaattgggg
agtaggccttcgaagaacaggatattggggcgggtaggaactttgccagtcaagtccatgttggcaaagagatcagcagggtcttcaaaga

ctgaggccatttttttggttgttgctgctgagagaggagcaagtagtagtgtggttactagcaaggagctcacagtcttatatatagactaacac
ccccccccccaaaaattctcccacatctttcataccggggtcggctagaacccagaaacggtttgactcacatctggtacattatttctgttatac
atttctagccccttcgtgtaatctgacattgggtcgacccagcggtccacccttcgaacttgacattcgatcgactcagcggtccatcccctaa
actggagtgacccagaaaaatttttgaaaagtttttgagatgaagatgaagggggaaaagggttagtgaatacaggcagttctgccttgctcc
agaaatgattctaaagatttctgtgcattatttctggtatacatttctagccccttcgtgtaatctgacattgggccgacccagcggtccaccctcg
gaacttgacattcgatcgactcagcggtccacccccctaaactggagtgacccagaaaaatttttgaaaagtttttgagatgaagatgaagggg
gaaaaagagggtgctagggtgcgctccagacatctctagagtggctagagcccagaaacatcccaactagtttctgggacattttttctccca
ctgacgcagactatataagctaaccactaagcatattttgcacacatttcatccaccatcactgccgaatacactcttgtgttgctccgcatagc
ctaacaatggccccaaactccttccagaaatttgctccagttatcaagactgagaagaaggaagaagaaagggatgaacatgacgacccttt
acggcagattgattttagagatagaaagacattaatctgcctcactgcaaactgtgtttcgaggaagagaaaagctggatctgcacatgatcg
agtatacaaagtactacgctacgggaacccatacaagtaccgtcgccccaatagaacacatcgaggattggccctctcaatggatcaaggt
gaagtaggaacatgcctccctctgcgacccatggaagagactgaagaaaaccccatcgacaagtgcggagtggcgttcctgtactccaac
tacaatgaaggcgatggcatgacccacctttacaacgacgaagagtatataaagaagtgcaaaacaattgaaggaggaacaagaacgtgg
gtaaagaagaaccgccaagaatacttcagacaagctctagagacattgatgatgtcccattctataaaacaatattccaattttatttttttcaag
gaggatatggaggaaggatttgtgcacaaactccacacatttattaatatggtacaccctaaaaaggtgtctgttttgtgatataataaagattga
tgtttttataaatgaaacgtattatttataccaaaaaaatttattatgatgtatttcattgggatggcttttcactcaccaatctggcctatgtgtgtgttc
atgctagcccctctttcacccttcatctccatctcaaaaatttttagaaaatttttctgggtcgctcgagtttagagggtggaccgctgggtcggc
ctaatgtcaagttccgagggtggaccgctgggtcgacccaatgtcagattacagagagggtctagaaatatgtaccagaaatagtgtaccag
ttataagaaaatgtgaccctttccagaaatggcttgtccagaaatctttgaacccatttctggaacagacgtttctggagcatggtcgacccaga
tatctgttggcggctgtcaatagtagcctatgtattcaatggtcttttaccctcttctccatctcaaaaatttttagaaaatttttctgggtcgctcgag
tttagagggtggaccgctggctcggcctaatgtcaagttccgagggtggaccgctgggtcgacccaatgtcagattacagagagggtctag
aaatatgtaccagaaatagtgtaccagttataagaaaatgtgaccctttccagaaatggcttgtccagaaatctttgaacccatttctggaacag
acgtttctggagcatggtcggcccaggcagcctcacagttacctgttcacacccttcctactagcacccctctttcacccttcatctccatctcaa
aaatttttaaaaaatttttctggctcactcgagtttagagggtggaccgctggctcggcctaatgtcaagttcgaagggtggaccgctgggtag
ggcccatgtcagattacagagagggtctagaaatgtataccagaaatagtgtccgattattaaaaatttaacttttcaacatgtggcatacaccg
tatgaacgtatagccaaatatgtaccacagtcaaaacatttaggtgagaaagataatacacatgtataaatttcaaatactttatttgcatacataa
taaaaaatcatacagtttcggactcatctgtccacacatcatttattaggaactggtatctattataccctattcttctacgtttacaagatacttgttc
atcaggcactgccattcttttacaagacactattggttcaaccaacatctctttaacttgatcagcgtcaggtcttagggaagtacgttctctgcat
acaggacaccgggtaatttctccacgttgcgtactcaacaccaatctgtcccaacatttttcatcaatacaatgtccacagcgtgtcgtcctgat
gatattggtactactgaaatcttctaaacacactggacaattatcgctgtcgaaacgcacattagcattcttctcatgttcttctagagcccttttaa
tttcttccttatccccagaaattatgtgttcagcccaaggataatttgaggatgcaatataccccaagcgacaattcagtattggccaagaacga
ggtgctttattgacatcgagaccggtggcagcaacaaggggttcttcttcttggagagagaaatctaaagcagctttcaatatggtgggtatac
taggctcgtatcctctctgggcaaactgtgttaatgcggtatcattgaacatcaataatgctgcttcagttacacttaaagtactagatttaccata
caacatattaatctttgccccatgggctccaactagtgcatcagagacatcgtgtttgtgattggcgatggcaatcattaggggcgttgcaccat
cgctcgcccttctagcattcactatacttctatcgaaatcgtatattatattcttgtatttttttagcaagagcatcaagaaaattgcacgaggttgca
acagctgcagcatggaatacgttttcattattttcattaataaaggaaaaggccaagctgtttgttgttagaatttcaatacctactcttgtggtaaa
gggggaagatattgcagcgcaaaccattagaggtgacatgcctttcgcagaaggcttcagaatgttaagtttttcttgtttgactagtttgctagt
atgggaatatttgttaatgatacaataacacaatagagtattccccttttcatcaaaataagtgtgtttgagccacggattcttggtaagaaaagct
tcaagatcttccatgaaacacgagacttttacacacaatctaacatagcctttaacttcatgagggtctgttatcacaacgtttttatgtgatacgat
ccttttcacaataagattatcgcacagtttgtcggtcttttttcttaccataatctccctcaaacagatcttgaagtaattgtaatcaatctcttcaaag
ggcctactgttaatcatccacaacagtattgcaccacacaaaccactacccactgcttcgtttattgcaacatttttttagaatattgtccatcatttc
ggcagagaattttggcatcgtgtcaaatatactgtaaagggtttgaaaatgctggattgtttcagccttctctataaacacatcaacaaactctac
tgccatttcagcaacagtttttgtagtagtctcacagtactcttctaccatttcttcttctatcgaggaaagaaaagacttgataatcacactcctgt

aagcactagccctttcttcgttggtccaattcaatctagctatagatctagcgtccatgttcatttcttctctgaagtcagttgacacggatgatggt
gatgtttctaggcaagaaaaaggtctcccgataataaaattgccattggatatcagtcgttttgcctttgtaacacaaggagattcgtccacaaa
atacttgtatccgaaagatatgtcaaaaggttcaagtggtgcagatttttcatttcagccacgtaatcagaggtgatattgacgattcttgaaaa
gagcctgaatctaataacactcgaacatttttcaacgtagaaaacaataccacttcttgcagaactagtagactttttcaggctagccaaaacac
cgtccaacttcttgatccttctcataaccttctctctttcttcctcagcctgttcctttgaagtaaacttgaatccagttctgctgtcatcaccagtgcc
aaacttgatgccgtgcgtctcgcgtctcaaaaatccattatccatagagaccagaagagaatattttacgaacaaaaagtcgtcgtggatgtttt
cgtaaaggcctctgaaggttttgcagacggttgtcaatgcgttgataaaagtcattccctcgcagatgggggaagaatcagacttggtattgtt
gttgataaagaagtagataatatctctaaactcttctttattgtctaatttcttgaaactacttgaaggaacaggaggagaatttctggaggtaatta
tgtcattcagaagggccaattccctttctgtgaaaacgtccagaaatgacatatatggttcaatgttttcaagtacttcttcaagcacctgacggt
atcgtggagctgcttcagccatgttgatgatgtctcacatacgactgttgagtttatccatgcgtacgcccgcttttatacaaagatcccgtgtaa
gaaactccctccggttcagttcaggataggggtgtgtcccagttttacatccaaagttaatatattttttaatataacaaaaaaatcgtaccgctt
attggctgctataaaagagggagcacctgctcacttggacatcattaaccatcatcaatatggagggagaacatcaatatttgaacctagtca
gggagatcctagaaagaggagtgaagaaggacgatagaactggaacaggaactctatccatttttggaccccaaatgaggttctctcttcga
gacgacactattccagttctcactaccaagaaaattttctggagaggagttgtggaagaactcttgtggttcatcaggggcaatacagacgcc
aaagaattggccaagaagaagatacacatctggaacgctaatgggtcgcgggaatttttggacagtagagggttatacgatagagcagag
ggagatttgggacccgtatacggattccaatggcgtcattttggtgctgaatatgatacctgttcttccgattatactggaaagggtattgatcaa
ttggccaatatactaaagaccctgagagaaaatccagatgatagaaggatgattatgacggcatggaatcctatggatcttcaccttatggctc
ttcctccatgccacatgactgctcaattttatgtggctaatggagaattgtcgtgccagttgtatcagcgaagcggagatgtcgggttgggcgt
gcccttcaatattgcatcatactctcttctgactcatctgatggccagtatggtgggtctaaaaccgggagagtttatcctcactcttggtgacgc
acacatttataatacccacattgaggtgttaaagaagcagttgtgccgcgtccctagaccattccctaagttgaggattttaatggctccagaaa
aaattgaggactttactatcgacatgttttatcttgaggggtatcaaccacacagtggaaacttgcagatgaaaatggctgtttgaatcatgttaa
ggaatttccttgttactcatttattcctagaaatggtgtaatcgctgttgtgggcggagcatatttgtgtatataagagcccgtgttagctcctcgat
tcagtcacaagagcgcacacacacgcttataactagctctctctctccactcaagatggcctttaattttgaagactctacaaatctctttgccaa
tatggacttgacggctggcacaacaacagaccctacccgccccaatatcatattctttgaaagtctactccccaactctggtattgaggtgatg
aagaggcgtctcgtacggcaaggaaagtgtgggaattttgaagcaagtggaggtgctatgtcgtatttctggctcgaagataatgcagaaga
tatggagaatctcaacagtggttcccatgtcaagacaaactgcttggcattattccttcaagagtttatcagcaactggattgaagagactgatc
gacatggacagtactgtacttttccccaatacatggacggtggggatggttcacgtgggggatattttacttcgctagccatgaaatggatggc
tagggatgtgactttctttgtgtttgttgataggaataatactgtagaaaatgcggcatccatatggatgtaccaaaaactactagcaattggtgc
aaaggtagtaaaggtgattgttgacaatgcatcaaacccaatgttttctgtatgtaatgcgtgtaggtgcaagtacccaggcccagtgtcatac
gttattgaaggccatggagtgggtcattctgatttgacatgtgatgagatttctggattctttgtataataaaaccccataagaaacaataatctttt
ttattcaacacccatgattttagtttttatagtatataaaatcaagaataaagtatgtagatatctacttttcgatagctccttcacctatatagaggag
gtggataaaaacagaagatatggattctaatacttctattttaccgccaagcaaacggccagggttaaatctgttacaggttttagggattataat
aacggtagcattaatagcttccgtttcatcctttatattttatagggtaggtaaacgcaaatattacccttcttcatcctcttcctcagaattatctgat
gtagataatggggtagaaggaggaggaggaacaacaacgacaccaactcaaccttcacctgacggtggagatggatacgtagatctttct
cctcaaaagaaggctgaactaagaactagagttgcaaacgtcatctttcaagaagtgtcaaaggatcaaggagtggcctttagacgggcaat
gaatgattcaactgataagataatggaagaaactgaggcgagaatcaataacttttcagagccattcagagaagcaaccgtagaacgtgaa
gtgtttaaggatgacacagacaaaaactttatcctttcaactctagatttaacagaggaacaatttaaggacattgttatggctgaagtgaaaaat
caattagaaaattttgactatgaagacatgacccgtctcatctttgataacatcccagagactgattatttatggacaactcatttcgatccgaaa
aaatatgacacgtactctgaaaaggtattagggttctcagatataaatagtatagaaagaatatcctctacattttataaaggtaaaaaatatgag
gtaactactggaaatgtagctgtcctcgttgattttgaatctgaaacaataaaagagaaggcaggaaatagtctcatccgtaatgtcgagtttatt
gttgtggacgaacagacctacaaatctttcttccctgcattcaatcaagttttcttctcctttaaagtaaataaggagaaaagggaagttactgtat
ccatcaataatggatgtgtaggtatagtggccaatattactcctctaactacgccagttggagcagcttccggacactacatctatggcactag
cacagcaaaggaaaagacctatctatttgtaatagacaagtacgataccactgaatttgtttgtggtctgagtaacaagtcaactcctctcatgg

ctctaaatattctctttatgagtgatactgttttcccttcatttgacgaagcagaaagacctctgacggatgccaaggcagtagaaattttaggtaa
aagactaggtgtaggaagatacacaaacgccaacatcagaaatactcagtgagatggaaggggttattttggataagatagaaacaattgca
aaaagggcctctcctagttatgggtctattgatgtgggtacggctattttgcgccgtcaattcatggaaaaaattaggggtaaaataaatgaag
aaaccacaatggagaagattatgggcacaaaggaagaaagagaggacactataagaagtatagtggctaatgttatcaaagagaatactgt
taaagaaaatgtaaccgaaaaaattagagcaatgacagataaggaattaaatgacaatagggaatttatgcatgattttggaaaaatttcaact
ggagatggaggaaccttccatctctttgaagatacaccgggttttgaaagtgctttaaaggcagaatataaaaacgttccaggagcaactact
ccaaaatacgtatctatgaacagtttacgtatcgatgcgattaatggaaaaatcgaagaggtttataatccttcacctatcatgggtattagagaa
tacggcaccattcgcaggggcaggtacgaagaaaatgcaggttcgaaagaattggtttttatgaccaagattgaaaaaagacccaataatgt
agctgaaaatctcattattagagttgccaaccagcagtataatgttatgaggatggtgtttttttatagactacgaaacaaagaagggggtgtcca
aggaggaaatgtttataccatataatgttcagaaaacaaaggctcttaaagggcgtagtacttacttttcattcgtaaggaaaattcctgatgaa
ccagaagggagtatcataatacacgcactagggttttattgaggaaataataataataataataatggcattacaggaaaaggatataact
atagggaatgtttctgctgccctacgagagttgatgtactcacccacacatatgcagcatcacgataagctaaacacattcctggacagaaat
gttgaatcatcttcagaagagaaaataagacaaattgtggataaaatacgatcccaaacaacatctgacatatctgaaacagtcaataatgtca
caactaatgggactgcattttccctttcgaagataccttagaaggtatggtgaaaaaaaatataggtgataaccttcagagtggggactttatt
gatggccgtaaaaagctcaatgacatgaagagtctagctactggagccatcttatctagacagcgagattttgttgcagaaagtataacagga
acaaaggactggctcaaggctataatgggttgtggtattataaggtatactgtatttgtcaataaccttgcaagatcaacactcgataatgatgat
gacaaggcagcaacctattataacacccctatatatggcgggtattgtaaaatggctataaaggactatgaaataccagattcgtacagcaag
gtcgaagcggaacatacagttgaaggaagaaagatgacctttaatataaaatggagaggcgataccataaacaacctaataacaatcatcc
cttcagtgacaggttatcttgcttccatctctgaagacgcagatgtgcaggcgccattacttttaaactgcaacaactgttttatagaggcagata
tgagtagcctctacatggatgagaaaaaaacagaggcatcatttaccctcaacttaccggaaatcgaaggagctgatgcgaatgcagtctat
gaaatatgtatagtagtagtttgatggaggacaaataaaaaaattggatacatttcatgtagttttatttaactcttaatcttatatacatttcatgtaca
tttcattaaaaatcactatcagagctatagtacagaaactggtgctggagctgattaatcctcctggccatgacggcagatgtcctccgtgaag
tgttgcgtctgccgtctctgaccctatcaaccatcctacctgaacctgggttcaaattaacatttggtgcatcaaagggggtgtgaagaggtga
agaataaggagaaggtacaggactagcatcatagtccctcacaatccttgttcttccccttctaacaggtgtggatgcgggagggggcatatt
agccgtgtctagcaactccaacatctgtgcctggtattcaagtgcgttagacacaaacctctcagtgtcttctgcttcattgtcattattaggaata
acaccataatcttccatgagggcgtcacgccttcctgcctgatgcctctcaaagagccgttttctttcctcagggtcatcagagaagagtttgtg
aacccagcatggtttcttagcctcccctacagccaagtcaggcacaaaaactggcttattttttacattcttatagggaagaagagggggcaa
caccttgacaggagcaggagtggggggtaaaggagtggtatattcaccccttatatgaactaggtttatggaactcctggcaaggtattttgcc
ctccctgtcctcttttagaggggcggccttacagcctccaaaaaccttcttagctattttagtatgcattttctcaatccttttcataccttcagacaa
catcatgaacccattcttatcaaactgaaaacccctcaagaaaaacttataaaaggacttttcattctttgcaagaagacatcccgccattagcc
taattgcactcaaaacatcccctatctccccttccccatactttatttcaggaaacctattgagtgctgagcaggtactttctatagatgctctcag
agcatcattttccaaggttgatgagctataatcaaccttcttacagctggcacacttcttaaccatatcctttatttccattcttttcttatgtttcttcat
catttctgatgaagaggaagttttgattgatgtcttaaagacgatattcttcccttttattgtctttcctttcttgctgacgatacaccctcccttcttgc
tgctgctgctgccgctgctgctgctgctgatgatgtcagttccggccatttttaatgactctaaaaaacaagggggtaacacatgtaataaaatg
cattataatgtaacattcttttattttaataaaaggcagtagaaaatttgtacatcatttacaggcaggcatatacaaaaacacatcactttatagatt
gggcagggaagatatttgatcagcttccgaagggtcagtatcaaacccattcaacaatgaactgtgtacatgtgcaatttcgtcatccaaccta
gcattattagaggctgacttaatgagacaggaaccaattagctgcttattagcactccactttgaagcgtacttacgcttcaccttcaatttcttag
agcacaaactcttaccaattacatcattgacagttgtatggaacaggttacctctagatatacaatctccaagcttattctcagtttcctgtctccttt
ctctaagagcagaaagttcagccagatacctattggcaatctcattatactcctcatcaatattatcgagtctacacatatacctcttgacatctac
acacagtctcttactctcagaggaaatattagacatcctgtcctcgtcacgcccataccttggcctctttctactgttactactactactactacttc
ctgaggaagtagagggctgattcactgatgacgaagaaggtattacttcttcgtcatcagagtctgaatcaacattaaaacaaaggttaacatt
aatgttaagagaggttgaggtaggggcggatgaggtaggggcggatgacgtagaagcactaggcatattcaagtcaggcaaagacattgt
gtcatcatcatcatcatcatcattaacatcataaagtgaactactaataccactaatagtatcaaaggtaatattgttacttctaggggggaacagtc

aagtcaggaagtgactctgagtcatcattatcatctacctctgggggacaaacactacagtatcctcatcatcttcaatgaaattgaaggatgt
cacaccagacaacaaactaggggagacattattatcaccttcatcatcatcatcatcattactcaagcattgagtaatgtcaatagtattgggag
ctacatacttaagccattcaccttcatattcataagggggggatatccttgaatttacccattaaagctgcctcatgattatcttcaaaggtctcttca
ctgaatgcatgcatttcctgtctagatagagtttcaagcatgaaactcaacttggtataagccctctccatataaacatcctcattctcattaacac
ctttggcattggggacaaagaaagtaggtgctagaatagttgacatcttattaagggtatcctttacaccttcatcattaacatagtcagcagac
aacaaaaatgagtgaaaagaggcatacttgtacacctcttcatacactttacggaagaaccacacagggggtatataggccctacacatggg
acataacaggtgattttcattcaccatatgctcttcaaatctccaggtgaaaagacactccaggtggacacttttcctgcaacacggcatccctc
catttgaccagttatagcaatccgtgtccgaatcgtaatcacccatacagatgggacaggtaggcacctcatcaacggcgacggtttcagcc
atcttgttgttgttgttgtttactttaaagagggagaaaagccaacgctatagcaactcatttctttactttaaacaggttcattacagtccccttaaa
agggaatatgtgcttcatggcaagggcatattctgtgattgaagggcgattttgatagaaatcagtgtctgttccgctttgcaacaacatcaaaa
agtgcatgtttgacatcaaattaaaggcagggccttccatgttagaacaagcaacaacatcttccagtatactggaagccattacactgaaact
gtacgtcattgaataggcacctagacagtgtcccctaaagaactcgggaggtgtgtgaatgtactgcctgattagttcttccgtgttcctaaaat
attcatcgttgtaacaataacctagagggtatgagagtcccaagtcaatcattttaggttggccagatatgctatctataacaaagttatcagcct
ttatatcgacattaataattccctcatttgcaactcttgtaattacattacaagtctcagccaaaacaaaaggcatgttcacctttagtttttcaaaca
tacttcttacaacctttcgggcaatgaaagagtctgttattggtgcgactgtatcaaccccctttacctctggtaaacgtcccatcctttgaagcat
gacaaccccatacaacaagcgtgcgtcccttgactcataatactccccaccatctaaactatccttgacacacctaatgacatcatcaatactg
atacctgcaaatggcatctctaaacataacccctgaggacacacaccagtcaccccaactaccccccttaatattctttctgtatacagaaaggc
cattaacacactcaaaaacaaactcttcatagttcatatccttcataaacttgacaacagtccccctattcacaatgtaatagaccccatatttaat
ctcaaaaaaggtctcccagggacagatgaaagtgttaaagtccttcacatcttcaggcacttcgaccaaatcagacaaacaaccacgagca
gtcttacggaagttttcccagttctcacactggtcagaaacttccttaattagcgtgctagtcttttgtctaattcgctcacaggcacttgaaggca
attggaacacctctacagccatatacattatgtccataatctgggaggacggaatattacgactagacaactccttaagaatgtgtctagtgtca
aaaaatctcatctcttcatcgcaagaagaggtgggccaatagaagccaactttagggtcctttccatagaaacaggtatcatggacatacttgc
tagacttaacaattgacctatggccaattctcaactgttgacaggtatcataattgtcctcaagactacaactcttctcaatgcctttaaagtagta
aggggaatcacgttcaatccactcattatactgcctcttacactgcctaatcaccttggacgcacacacgtttgaaaaggtgcgagtagtggg
aggagagggagcaacttcagacttcttgaaacgtttagggggagggggcgtgatgctaaggttgatgacgtcatcccaagtcttgatagtga
tgttgttgttgttgttgttgtaagcgggaacagttgtggcttcggaggttgaaccagacgcgtagttgttgttgttatattcttccacaggccgttttt
tattcccctgatggtaaacatactgctggtggtggtggtctccaccagtattgatggtagtagtaattacagtgggtccccccattgcctatgga
agaaacaacaacaaatacacgattaattcaacgcccggcacactaataaagtgatgtgtttttatataaaaaatagcccataatatttaacgtta
cgttaatcaaaaaaaacaacaacaataatagtatattattggtacttacttttgatgggtttggagcggagactgaagtaaatccagaaaacaac
agttttactgttgggcactacactgcactggtgacgtagtagtagaagtagtagtggtaatagtccactattgtcactgcactgcactagtagta
gtagtagtagtagtaagagcgttggataagctcgggttggctgctagagctggagctggaactggagcttgtatgcttggctgtgcctctgct
gagactgatgcttctggcctgccccggctcgcttatatacaagttgtccccccctcacccccacttccagaaatttgccgtcgaacgccagttct
ccaacagagtggggtccagatctggctacgggttcactcctggccataccctgtttaggggtctaaatccacccccggcactaaatgggga
ggacctaagggtgttataacaacatgtaagcgttgggtaagaagatctggatctggatgacccaccttgtccttcttatcctatccttatccttgt
cccctgtctaccaccacctcaccccatatctctcaccccatctcacctataccccccatctcacccatcacctctatattcccccaccttattcactc
gctccagtttcaacaccctgttcttgccgagccaacccataaccagatctggaccccagtcttcccttttatccctaaccggcaccatttatgcc
cccaggcgctagcggtgtatataaggcggcgcggccaggccagaagcatcagttctctgcaagccagcagaagagcaacacaacaagc
actctctctccttctacctagaagagacctgccaatactcaagctacaagaatggcctctccagcccccgccgcaccaagtccttacaccatg
ttggactctaagttacttagttctgaggaactaaaggaactaacttcatacgtctcgactagctctcgccggtctgatatgaagaaacacttgct
ccatctattcgaggagcacgagaagatcttccaattcatacaaggtaagcacaagttctcactatacactttggactttgaaattttctatgttatg
ctgaatattttgttggttgaagtgaaaaatattctaagtccaattcctttactctttgacagaaatctccaaccagtacggagactatggatgtttca
caatggccccgcctcacctgaacgctgcagccgatctcttggataaggtgatgtccggacctctatctcccgaaggcgcccaaacttcgtcc
ccggctgcttgtgttggtgcgaaggttgtgaaggcactggtgagcttttgccagaagacccgcttcaccaccaacattgtgatgagagaagtt

aaagccatggagttccaaggagacgattttaactactctgccttgtgtgcaagtatgccccaacgccccgtgactgagaggcagatgttcgc
ccttatgaagagtgaggacgaagaaatgggagtgtctgcaaacttctctccagtctctgatgacgtcatcaacccttcaagcctcccctctgg
acaagaagtcgactcatcaacttccgctcaaatttctggtatgtttcaaaacgtgtggagtttgcttgaagagtgtggtagtggctctaatagtaa
tagttcccctgtctctaggacagtcttagtttgcaccctgtttataatccaagtgttcaagtttttggtgactaaagtgtctaatgtgaacgtacttaa
ccagttgtttggacatgttgtttttggatcacttgatgtggctccaagtaataataatagtgtcccatcaactgttgttaacaacaacaacaaaccc
tcgacctctaataatagtaacaacatcagtaacaagcgtgttggtggtagtaataacagtggcggcggaagatcaaagaaagttacagccac
agccaaaaatccctttaataatgtagatggggacaatcatggcatgtttgccggtgcccctgttgatgttaatttggatgactttgtttttccccaa
gttgaaactcttacaagtaagagcaccatccctaaagaagaggtaaatgtagatgaagatttgagtaaaatgtgccgtaaaactgcccttacc
cccctagaaattcatacctttaatgtgttcatctctgagattaacccctccaaatatgaccgttcaatgttttgcaagggatttttgactgcatggga
taagtttgtagagggggatactgctggcgttaaacgcttccgtaactatatcctcactcgctcaaactatgcctcagccgccagggccgtgtat
gaagcgtcaattaaggggactgtttattataatgacaagtcaaagtttctgttccacgataatgttaaccctgatctggacaagagctggggtaa
caagaatgggaagaaacctagactcccagctaacttgatggcattcatgggtattgacattgtaaaggtgtgcgctaaggggattcaaaagt
atatgtttgcaaagcaattccaacatccggaagtggaagaacttgtgcctcctatggctgtatacgcaaaggttgccgcaggattgaagtcgg
ggactttgtttgatgactgggacctgcctgaatacgaaaattgtcagtttatcaagtatgacacagaagggtgcaaaaagcacagtgagttata
cgccaaacaacttctccgcacaggacttaatcaatacaataaactggaagagggacagagtgcattcccatttgcaaatattgtgacggtaac
atccgcctctagtgatgatattcacggtgacacaatcattgaattgatgtacaagacaaaggatggcgtaaagggagtctcaaaaattgagga
cgaaaacatcatcaaggtgaatccagcagaagaaaagaagaataatagagtacaagccgagaagaccctgtattttgagattgattccgat
gatgaggtgtgtgagagaacagaggaagaattcttcaggcctacatctgttgttgctgccccgacaacacccctcgtaccttctaatgtggag
gaagaggaagaggaagaagagcagatggaagaagaggaggaagaggaagtagaaagggaagaaggatctgataaggaagatgacg
gagacgcaccagcacaggaagaaatggaggaggagaaggaagaagaacaacaacaacagccagaagaagaaagcaatggtaatga
gaaccaagaagaagaacaacaacaacaacaacaaccagaaagagaagaggagaataaggatgcagatagtgacagcgacagtgatag
cagcagcagcagtagtagcagcagtagcagcagtagtagtagtagcagcagtagtagcagcagcagtagtagcagtgaaaatgaagctg
aaaagaagaaagaagaggaagtacctgccaagattcagaagagaaagaggctaagtgaaaggccatcagaagctgcttcctctcccaag
agaatgagagtagaagaagaacaacaacaacaactatcaccatcattggacatactccagactgcagttgatgagatgatggaagaaattc
ctgcgcctgagcctatcgttgctacaacctcacccaaggcagcgacacttgcactcaagacaggatttagttactcttcattcgtaagaggag
atgacctttcagtagctggtaatacttcccctactgaaccagcagctgtgcccgctgctgccacttgcacttccgatgttggaaatgactttttgg
acatgttggacggtttacctggcgatatagtaatgcaacctggcgaatgcgacgtgaccgcaaaattctttgagggcatcaccctaccagatg
gtactgataatgaatgcacaggtttcgatgatcttcttaaagccaccgagactgataacattataaccaccacatgctttacctccccgattcac
ccttctagcaactcagcccccagaaaggatattgataattgcagttctattaagaggtctagggcaggttcactttttgacactgatgatgatagt
gaaacaaatgaggttgaaaaggaagcccctaaacgtaagaagcacttgaaaaagaggcgtaacaagtcccaccgtggttcctctggttctg
cttcttcttctcattgtatgagtagtgatgaagaatcagaggatgaaagggatatgaaatcaacatcaaaggttcacaagtcaccaaaagctca
tgttaaacattcccctaaatatgatgctgtaaatagtgatgtaaataactcatacaacaatgttaatagtacaacatgcatgtcgtcatcagatagt
gatgcagaagcacagcctaaaagccataataaaagccactctcgtaaacactcttcttcctccacaagtgataagaaacagaaccaacaatg
ctcaatcaatactcaaaatgtcaagaagactgttgtacagtctccacctagttttagaagttttagtcctaagaaagatgagcttggtgatttcttg
tcacgcaagcacacaaagccagttaggccctataacaagaagcgtgataatgttaacaccactaataatgtagtacagaggtctgcctgacc
gactcaaatgatactcaatcaatgtacaataataatcttagtacttaacaagaagaactatatttttataatatttttacatgtcttaataacaaacaa
aataaaagaaaaccaatgtattatatgtttaaaatcaaccccatttgcatgattaaactaactatagtgtgtaaggaaagaaaaaaaacatgatta
tttctgccattaaaacaacaacaaaaattctaagcttctcctttctttctgtgtctttgcagattcgggagtctttctaaaaccacagcaatacaaaa
aacaacacaaattctctactgctctccctcaccaatccccctatcgcccttatgctccttctcctctctatacttcctccccattcacttctgactatc
aattttctgattcatttcaacaacaatgccaccaatacaattactaccccacacccccctccatctatgtcctcgacgacctctccatcgtctttgtg
ggatgatgacgatgatgatgacgaagaagacgaaaaagatgtcaagcaagaagtctcgaaccgtccccccatttttcgtacatggaaactg
tatcttttagtgataacgatgaggacgataacaagggagaagaagaatgttttggatcaaactttgatatgtttggtgattcagataacatgccat
caacttctactgcccctttccctcctccctctacaacaacaccacttcctactcctcgatccatcatggatactgattcggatgaatgtgacgaag

aaggagcagcagcagcatcagcaccgtctattgccgcctcttcttctatccctgtcgggatctctgaagctgaattgaaaaaaatggaaaaga
aaaagaggaaggaaattaagaaactcaaaaagatgatgaaagatcctctccctcacctatatgtaggaggagaacctcctgtcgcagcaga
ttataaaacaagggcaaacatttccctttataaagttgaccctagtatcgatatgtgcggtgtcgcccctcctcaattttgcgctgaattgcccac
cccatccatagatgtgtatacttcttcctatgtatttcctcctcccacacctgccatgcataataagaaagggttcaagaaatgtcaattccttaag
gggagaaaggctttgaggaaatggattcacgagaatgtatgcatggcccctcccggtaaaaggggaggtgtatttttggctcacttggaaca
aagattcttggctgaacatggagatgaatacaaggtcccaaggatgtttgtttcaagagtattgaacaaagctttccccaatctgattgctcgtg
cagacacactgtgcagtgatatgacattctatactaacctttgttggatagttaatggagttgtcgtatgctttgataaagatgatggaggaatac
atggcgatgcgtcagagtatgcaacaggagaaaattttgatactgtagtgttccacaagagggaagaacaaaagaccaatgggagtgcca
gtaagaagaggcgtctcacgcctgacactagtaatatgggaacaagcactgatgtgcaagaattccaaacgatgggaacaaatactgatat
gcaagaattccaatcaatgggaacaaataccaaccccatagagacttcatcagtgggtgtgaataccaacccacttcccaaccctcccccaa
gattggtaattactcctttaacgaatgatgtaccagaattggacatgatgtggctttattcgccttccagaggaggtggaaattctagaatgagt
gcaaatacaggaacatctcccctgtctaacaccccaattcctacctgcttcacaggaggtgcgaatgtagtagtgcctaatggatttgtccctc
ccacgttccccttagaatgtgacgaagatgatccaagtattcccaattcttacaattacgaagaggataaagtctttcatccattttatgagtatat
ggccaaatatctatcccctcttgttccatcatataacaagggacagacttgtaatgttgtccaggagtggttcaagggatccttctctcttgcaaa
gcgtagaggaacagtccccaaattctgtagtaacatttcccacgctttcttttgtaatatggatgtatgtactgccatgtgcaaatgggcgaaga
ctgtaattagacatggacaatattgtaatagatgtatcgtaaggaggtcatgtacatccatgctcgcatatcactacattgtttgcagagacgctt
catgtgatgttcccaagtgcagggaaagggttcgcaacgacatggatgactgattgattggttgatatgtgacattttctgtatattgtgtaaata
agataccaataaactaatgttttatatatgattctatttttttaaaaacctttaaaaatatacatataaaatgatgtatttttgaaactacactctggcag
aatcagaccagacccctgacctaaagcagaccacaggggagtcttagagaggggtgtgaatctggctaggggttcatccctcaatgttaat
cacacgcaagtaaaaacaccacttcctagaaagggagggaaggggtgacttggtgatatcataactggggaatttcctctccagatatctgg
ctgtacacgtgtgagcgcttctgggcgcgacaagaaaaaaattagtgatatcataactggggaatttcctctccagatatctggctgtacacgt
gtgagcgcttctgggcgcgacaagaaaaaaattagtgatatcataactggggaatttcctctccagatatctgggtaagaaaaaaaattagtg
atatcataactggggaatttcctctccagatatctgggtaagaaaaaaaattagtgatatcataactggggaatttcctctccagatatctggctg
tacacatgtgagtgttctaatctcatttttatatagaaaatataactgaattagctctctaaacttttcctcatttcttactcctctttgtttgtatgtccta
ctaccagtgtgcatataagaactggagagatgggagttttaaataacaacccttcttaatccctttcatgtttatatttaataaatataatcaccatg
gatatttcaaataagacattatttttagtagtcggtacctttttctgaccacttgtgcatcttgcagtccaactcaaatcgtctggaacctcatggta
gcttcatttgttggatttctaggacataaactacttaaaaacattacacctgtcaatctggatcttgtcggaaaaagcttcgtattctctgcaagttt
aaccatctcagaagaagcccgtttattgaggattggaaacgtattaagggactataatggcaacaattttgaagagtatgaagaagaagaag
atagtggtattgaagaataataataataataatggtttgaataaaatatagagatacatttatattgttttatttgcattatatataaaaaagcactaca
aaatttgtacacatacattgagagaaaaaattgatacaatttcttcctttttttactggtatctgatttcttgatattcgagagagttagtagtagcag
aagaagtagcaccaactccggcagcagaagttgtagggataggggcggtggcaccggtggcagcggcagcggcagcagcttcacggc
gttgttgttgttccaggataatctttgcgcgtttattcatttcagtggtgcgtgtaaagatgcctctgttgtattggaccaccttattcctatattctcc
gatgtccttatcgcgcttctcaaaaaacttggccatttctccacaacgttgacgggctgcccatgcagtcatagccataaaatcaaattggtca
gaggtgatattatggttgtttagaaggtccacagcgtaggccatacagctagcggcacgttgaatattggagtacttgctgtttcttggtttaac
ggtatggatcaccttacttccatcattcaagaattctgcctcttctctgtgcagaagagatgtgcctctgctaactgtcccgtcgtcgaggacaa
aagcaggtccaggtatgggtgtttcgcaatctccgtaagaagtgtcagcaaccgtgaagtgggaaggatatagtccattatctccaactcca
gaagtgagtgcagcgctataaacttctccggtgatttggcgagtgactggattacttagccctgcgtcaatttcagacatatttccttcttcgggt
gtgggaaggaagggcctaggtggggcgagagtgtcgttgtttctttcagccgtagtagtagcaagagatgtgacccaagaatatttcttgttc
ttagactgtcttccagaaccgtcgtcgataacaagagatggttcgtcctcgtcactgctacggttgccattctggttcttcttcctctttgaatcctt
cttcttagacttggacttgttcttggggccaggaacaggtataacatccacattgggtgtagaagacatcctcgttccagtagattcagtcaga
gatgtaagtgctggcacgggaggcatggccgtaacagagataccttgtggtactcggaacattgcagaagaagacgtagaattagagttga
tacttgtaacagcagaagcagaagtccgagaggcgatggcagcagctctggcggcagcaatcataggggctgtagtggtgcaaatcggc
agagtgttggatgaagtcgtggcagcagaggtagagttgatgtgagacatcttgatttcttggttggttggttggttgtggactattgtgttctca

gagaggacttgtagtcgactgtgctctacccagatccacagccccttttatatccgattttcgggggttgccagatactaatatactcctccacg
ccctcttcttgcgccatatctataagggtcgaaagttgtataccttgagttgatccggttataaccgccattgttgggtctttctgagaatcgaga
gttgcgatcacggcatggcctcgtgggggaattttcattgacgtaagcagtaacgtcagaagaattgttggagcggggttggttaaagtgcct
ctggtgttggtggcggtgttggtggtggcggtggtggtgagcagcagcaggggcgtcgtagtgttcgtattgctcacaataatagtccttgg
gcacttgcattcttcttggatgaatatatctttctctttcttcgtatctgggttcgtcctctctttttatgtgttcagtttttatgtgagttactctgtccatct
ccattttctcagaatttcttccccctccattgacgtcatccgaataactcaactggcgccggtgatactgatgacgtcttattcttcggaccttatta
tcccctccttctgaatcatcagatgattcagagtcagatgaatcagacgaatcggatgagtcagaggaagaagaagaagaatcggatgagt
cagaagagtccatggagccactcccaattccactactaaattcagcttccaattctctgacgacatctatatttttagtatctttttcttccacatctt
gattcgtattagggtatgtcacaccagtttcgttgctttgttctgcgaggatattttcgaaaccttcaagaaaaggaatgtcaataaaaccttccat
ttctacgatctccccttcttccaattcttccttttttgttctcttcttgttggtttcttgagacggtggtttcagtaaaatttgggttatctgtagatccaaat
ccaccagttccacgcacgcttcttgcttgatttttgttgtcctttttagaactagagtcaataatcgttctctctcccgtggtttcattaatatacacaa
tctgttcttcctcgacatcacaatatcttaagaaaatcaactgggcaatgctggttccctttctgattggcacactttttttctgcactatggttgcgc
agaatcactttcaattctcccctataatccacatcaatcgttccagtaggtacactagtgttgttcttccatgtcatcccagaacgtgacacaattt
gtccataacagccgtcgggaaacttgtctattccatatccagtagagatcttggccagtcccataggttcgatatccatttcttcagagggaaat
aggtcgtaggcgacagaaccgggcgtggcacgtctgggcggaagtgcagtttcctccccgggagggggcgaatctcataaacacgacag
atgcagatgagtccatggctggatacagggagtacacagtggagacgggatgtaataagtgttcaacgtctggaggacaagtgagtacctt
cctcggttcctcccacccttttataatcgtgaaacccccacacccccgctaaggcagtaaaaaaaattgacccaccatcacccccttgtccttga
aggaggaaactagaagataacttttatcatggcaacatttactgaacaggatcataaaaatgcgtttttatatgctaatgagaagctgaggcag
gaaagaatatacagacttaaaatgtctgagccttcagtttatgcttttattgacataaaagaaatagaaaatggttgggaaaaagaattcgggct
tttagtacaaccaggacagaaattagctcctttcagggatatttcttatgactcaagcaaacttgattgtgacgcattttcttgcataccttcagata
tacttcattctgataatgaaaaaagagtaggagagtgcaactttgccgaacacacctctgtctcatttcctgtcaagaaccctgagggaaaaac
attgcgccatttcacggcatgtggtccagggtgttaccggaggtacaagcaaagagacccccatactggtttgccagtagccagaggcgtc
ctgatgcaagatcacgttgaccatgaaactggaaataaaatgtgtgaatatctgaaccagagcttagtcatgtgggctgctgtcccttggatac
gacctggagaccttactgaaggttacaacacaacacacgtgcctggctttgcattcaaggaggatgacgaaagggattcaaaaagagttaa
atatgaaaatgtggtcatttcaaaggcttattgtgatttctttaaacagtattatgacgcagactctggctcatgttatcgatctggatggatgaaat
ttgtccatttaatgtttgggcagtattttactaatctttcttacaatttagctaacccaaaaccttacaatttaactggaaatacatggtctgatgtagtt
tctgtattgacagataaccctattgtagatgcgggtgctgctccgtccagatctgaaatggatgaaattatcaccaagaagaagttcaacgtatt
cccttctgagcagacatctgctcgccagaaagcagagaatataatacgttctcagtacggagatggtgtagaaatagacccatcttctgtgga
tgctttaatgcagtttgttaatagggaaggtgtagtaggaacagagaaaaaatccgaccgcctcatgcgagtggcagacgctgttatggatg
cggctatgcgccttcaagtcatgggtctggacgacagtcaatctagacgattattgttaaaaaatatgattaaaatgagtagaaacaacccag
aatatgcaagacatttttccagttctctcaaattaattggagtaactttggccataaaaagatctgttttctctaaaggcgcttcagctaaaagaaa
ggaaacggccattaataatggcgaacagcatagaagaagcagatggtcacccgagacggtgacagaagaagatgcacttttatttgcaag
agaaaatatcactgaagacccaaaacaccctgcgccctttgtggacattttacactcaccagatatcaattcatctatcaagagcggttcttctt
cctcaatatggaacgatatactttcaagaatttcttcgaccagaaaactggaagaaaaggcgagtgttttcgttaaaaatcttgtggtgaaagta
gtgagacagttcttggacattttagaaggtaaactgttttcggacggatacgaatgggacgataatattcctctgatgataggcgtggaccaaa
tattgagagaagtcattaaggcggccagtaatatgtgcgctagatttgcctcctcagctctggagtctagtttggttactggctttatagactcgg
cgagtgctatcacttctaggttagccgtacaactggcagccagaacgttctccgtattcttggaggagtctgttatagaatttgtagttgccgca
agccttcggctagcgatacaggcatttgccgatctggcaactcttgccgcgtcagccttaaccgttattggaattgttatattcgtaatacaagta
cttgggttgattttggatcttgcgctagggttaggttggtacgatcacatttttagcccagaggatttaaagaagcaggttctagtgtttaggaga
gagtttgcaaaggcaggaaatgtagatgtgggtgtagctcaaccagtcacccctgaagaaatcgtcgctatcaacgttttccttcaaactgaa
gaaaacggggaagaaaagaaggaagaaggagcacgaaaatcaaagattgattttcttcaaaaatacttccattctactcctttgatgggaaa
gaaaagtaagtttgtctacatacaagaagcagctcaagaatacttgggaggaagaacaatgaacgcttttgggcagcgtataataacagctg
ctgatgatagtgacaccaccaccaccacacaagagggaaggagggatgacgaaacagtgactaagaaaatgaggagtattattctagaaa

caggtcaaactcttaaggattactcgtctgctgttaactataacgcctcccgtctagattacgtgggagaggaatgggtaagaaatactgccct
aaaagaagagacaaggagcaacactactagtgataacctattcaagaaaactgtttctcttgctagtatggccggcgcatttctggttctagga
ataggtgtattggtagcgtcccatattacacttttacgtttcaccaatattggtctagcttttgcgttcgcaggtctcctagcatttattgcacttatga
gtatatcatatataaacatgaatgctatgggtgtagtgaattcggacgcaatatacaggtctactgctctagttggagatatcaaaacagacccc
agaagagtaggaatggtccagcgccacgtaggtgtcggggctaaatacaacatgattacagatttcgtctctccaatgttagacgagatcga
gagtgactaaaaagtgagggttataacccccccccaacaaaggctttaaattagactcaaaccaaattcataatcattgtgaagacatcgac
ggtcgaggaaggtccgtgctcaaaaggtcccaaaaatattcctcaatcaaaaaataccatggattcgttgactaatacagtaacactcctcgt
gaatgaccgtcttggaaatcatcgaacaaacaaaccaatcaccgaacaagatgtggaaaatactcttaaccttaacagtctagaaagggcaa
gtcttttgaagttatattctgttttcatcaaagaaatgcagtcctattctggatgtatacccaagaacaagtacacgaatgtgcaagaaatattcga
agatggactaattactttcgaatggagagatggaacaaaagtacacagatcagtttcaccaagttcccctatacctctttctacaaaaaaatcg
cctcggtcctcaccttcccctcctccatcgatgccttctattaaagaagaagaatttgaagaagaatttgaagatgatgaagaaatatacgaaa
cagatgaaaatgtggaagatttcataaatggtgatggagaagattcagaagaggaagaagaagaagatataattgttgatgacgaagaaga
agagaatgaagagggagaaaacaagtacgttctcgcattctctaatcatctgaggcgccagactgccgccgctgccgccgccgccgcag
ctgctgctgctgctgacattgagaagaaggacaaaaaccacgcagttagtgcgcatgactacacactgtccgccctccagcaacagcaac
aaaaactgctccagcaacagcaacagcaacaacaccagcagcgctcctcatctgagaaggtcacctccacacccaacaaattcaacaagt
ttttactgccgagtaatggcttctcagaacagaccgagctctttgtttgtttcgatgtggataaaattgcgcaatataatggactcgtggagctag
acatcttacccattgttgctgaatacattatcaacggccttggtctgaaatgcagtatggaaactcccccagtgaaaccgtgcaggagaaagg
aagtgaaagatgtgtggtgtcagcctaaaactagctttgaaaatgatgctgtggaagataaacatctcgcattcgcagaatcgcctatacttca
aaggcctagagatttccctatccctaaaaaaatcaccgcctatttttgtttagacgattctgtagacattaaaaacccctggggttcgtgtccgct
tttgaaaagtggatcaaactttcgagtgtccgaatattcgcgtcattttaatgaattctcaggagttaaaaatgacgatgacacgtcttcaaacac
ttgctttatatactctcaaaaaaaccccaacattgaaattgtatcaaaattaaatattgaatttgaggtaatgatggagggaattataacccataga
aaagatttgttcgagacgggcattttgagcgattcttcattagctacggctatggcattttgccacccaaaagctagagttcgaaatgttgcattg
ttttatttttctgtatatttaccattctctaaaataactcgcaaagaaactataaaatgttcagagacggataaggtacatattggttcagacgcgat
cttttctcccccaagtgataatcctaatataagtgctcaccagaacaataacaacaataataacaataataccagtgtgaatatcgaggacaga
cctatccgaaataataatataagcagaaaaatgaccatcacaaactaccaatgtatggcatgcaaggaaagatgcacaaacaattgcactaa
cggtaactatcccgatcgtggtaaccagcacttgtcacatagtgtaaaaggggaagatttctttaagattttaaataatagtaaagtagattcatt
gaaaaaattgagcagagtactgattcccgctcctccctctggaaattatacatctaagttttgtgatagaagctctatgtgccatagcttcttttgt
agagggatcgaaccagtgtctacttctttctcatcggacagttttgaaaagactaaacttgttttgtatggtaaagtcgttgacgttatcaatagtt
attctgccataaaaacttcccataataatagaatcagggtctttttttaactctgaggaaaaagataataagactatcccctctagagccgaaagt
gcaaaaaatgcattcaaggatatacttgttcacgaatgtaataaagaacgagctgtttcatattttgagcaaaacaaattatcctctaaagatgg
gcatctatctaacaagtggtggatcgaacttaatgacttgaacattatgtttgagaaacacgtggaagatttttacaagaaatgttctaaagtaaa
tgatgcagaatctttaaaggatatttttaatgattttgaaaaaacttgtgataaatacaaaactgccaagagggcaattattggagcacaagacc
cttctacttctactccctctaaaaaggagaatggtatcactaggattattagtacattatccgaatttcattcaaaagatgaagctacagtaagtgc
ccttctcgacaaaacaatgctcttgggatcgaggacaataatgtctggtgttagatgtgttatacgtaacaatagtgtgttttcgggctttgaaaa
taagaacactaataataattgggaacttgagattagacactatgtcatctctatgggaggtgctgcagtgacaaagatttccgatgaagatttgg
aacaattcacgcctgtaagaggtgctgtctctgtcactacagcacctaatgataagctacctgtaggggcacatcagacatggaaggatgaa
caaacactaaaaacaaacactaaacgtaatagtctatatgactcttacaattcaaaaaggaataatagggataataataaaataaaaaatcgtt
cattaaaactatcagattttaattggagaacacccaatatctctattcaagaatttaatgcaaataaagatgatgttaacaagaagaggtacgca
gaagtcgtggcgtcagctgctccaaagtcaccttcaccaacgagcagcagcagcagcaacagcaacagcagcagccctcctctttcaccg
ctctcaccaacagtgaagaatagtaataataaaccattgtatattcctccccataaaagaatgaccactactgctgtttgaccaatttatgggttat
tttataatgttgtgtgatattttatacaataaaaatatataagaaaacaaccaaatgattttgtatatccttgttctgtgttgggcggaaattcctccgc
gcccattatgatgaaaaaattgctagaggccttgaatcctggtcacgacaaggttaacaacaaaaaaaattacctgtatgtgcaagtcatatcc
tccagtcccacgcatgagaacctgttattgcaccacgattcgtgtaacaaagtcgtatctgcacatgtgcataaattccttcacgctaaaatgaa

cgacaaaattgccgggagcaatgggcgtgagagccggttcctggaaaaccccacctccgttccacgcactacctcactattgcaccatttttt
ggtgtataaaaaggcacggagaggtagctaaaacatccactcgcagagacatcaccgtcaagatgacgtctcctactagtgctctccgcgtt
gctactactgccctgctgtttgctgtggtggttacggctactgtgatttctgaagctgggggaactaagatagacgaccgctgcgccttcccgc
catgtgactacttcccggaacctgaatgtcgtggaagaagatgtgttagagctgctcgaaggggggaacaacaacaataacaacggaaccc
gtgatggagccaaagtctttggcgaaatcgcaaatggactgattgggcgtaagaggcgtgaagcagtatctctaacaccagtccatgaaga
tatgccagatttcttcccctaccctcaccctgagcatcccattggagggcgtctaccacgtgaagcagctcctccacttaaaggtgcacttgga
cgtaaaaggcgtgaagcagctcctccacttaaaggtgcgcttggacgtaagaggcgcgaagcagaatccttggaggaagaacttgtgtct
gctgaagaagaacgtgaaaagcgcgaagcagctcccccacttaaaggtgcacttggacgtgaaaagcgcgaagcagctcccccacttaa
aggtgcacttggacgtaagaggcgcgaagcagctcccccacttaaaggtgcgcttggacgtgaaaagcgcgaagcagctcccccactta
aaggtgcacttggacgtaagaggcgcgaagcagctcccccacttaaaggtgcgcttggacgtaagaggcgcgaagcagaatccttggag
gaagaacttgtgtctgctgaagaagaacgtgaaaagcgcgaagcagctcccccacttaaaggtgcacttggacgtaaaaggcgtgaagca
gctcctccacttaaaggtgcgcttggacgtaagaggcgcgaagcagctcccccacttaaaggtgctcttggacgtaagaggcgcgaagca
gaatccttggaggaagaacttgtgtctgctgaagaagaacgtgaaaagcgcgaagcagctcccccacttaaaggtgctcttggacgtaaga
ggcgcgaagcagctcccccacttaaaggtgcacttggacgtaagaggcgcgaagcagcagcagcagctatgcctccccctgaagacgat
ctcgacttcttttacgcacctgttgctttgcctctacatggagtatggaaagcaccagaacctacaggttaaaatgtggctgaacatcacatata
caagttaattggataattgaaacaatttgactttttagtacttaatgtttattaattgaaacaattggactctttcctacctaaggtttattaattggtac
attgttaacacttgctctttgctcatagcagacatacctgctatttttaacgctttgtttaacgtgctttttacaattgatattggtacatgtttgtgacct
cttgaataaattttgttagcatcaatatttattttatttttatccttttcttgtgtgttagaattagaaggagatgtggaggttgttgatgtagtagatgct
gtgcagagttgctcctggtgttgctcctcttcttgattcacattgtcctcttcaacatactcctctaatccaaagcgctctcgaattttctcctttaca
ccctcaccacaatctcctatcgaagactgtgctaatataatttttgtagtagaagtagagaataatggtacaacgttattgatgttgttacttctgcc
catagctggtagtatttttgaaggcatgccttaaaaaagggatatatctttgtccttttctatacaaaaggataatcgttcatccccatgaaaaagt
gttagatggggcaatgtacacctatagaacagtgttatttcttctatatcttccagtatttataccagaccctcttgccccatagtttcgtatctccat
tttactattcctgtctctctggtgaacccattctgagccaggtgttctgcccataatttctgatgttttatcaacaacaatatcagctgaaggggca
aaaggccccccatcttccagtgcactattcccattagaaggaagtggactagctggcataggtgacaataatgaagggagtggtgaaaaca
caggataatcatccacaccctctgtcactagagattctagatccgggggaggaggtgtaggcaacataatttccatctgcatttcttcccccga
cgtggacggttgctgttgttctactccttctacctctacttctactcttcctccttctttattattttctgcttctagttccttcattaccatttccacgcct
gttgctagtctcattttcttattgttatttttatgcgttctttagcttgctttgttacttcatatgtaatatcagatgagtagtatttgcctggacaaataa
ggttatcactcatggctgtataattagtatcagaattgccatgtgcattatcttttgttataaactctttgggtggtttatttgcaccagccgtgaaag
tatttgccagaagagccgaggctgatgaagagttttgcttaacatacttttcatcgatagttttattaaataatgtaactaaactggccgaacactt
actatcccggcaattcgcacatgcattattattcttctttttattatccattgaggcctcctttgcagtttcaattaaagctgatatttggcccatgattt
taaggctcgcacacagtgccatttcctgtcctcctttctctagttgaattgatccccattcattaaatttttcttgtagagtcaggagactactcttga
gaaaagagtgtaaaagtgtttgagtagttatacaatttttcaacgtttcagtgattgggtcgaacccgaatttgatctgatttcttacactatccagt
aaatccttgtttaattcttccgtgtcagaggaagggaaagaaattatatcattagctggaattgattcagaaaagtgaatatttttaagtacatttcc
gtctgttgaagaagggtgctttagatcacacgtgctaagcattacatctaatgataagaaaggggtaactctttctcctgtatcgagatttcctac
aacattttctggtgtcaagagatcgtctgtttctgtcactcccaccaatctcttgtaatcttcctcagcaacagtttggttgttgttatagaaagacc
cataagtctgaaaaattctcctcgcagtctcattatctgcatcattagctcctcctcctcctcctcctgccacatttaaaagatctaggtagcta
ctcatctgggagggaaggggagggttaagggttgtatataaagaggcttgtctgaagacgcgagtgatcacaacttcaacgatggacgtcg
agttcggtttcttccacggtctgctctccaaggcccttctcccagatgaaaaacatcaacccgttataaggcgtctttgtgcggatgattctaga
aataagggagaggatggctgctgctcgttctgtggaagaagaggaacaggagagagcaatactgcgtgccttgaacaactaatagacgttt
gttcctttataggaactgtctcatctattggtacaattatcaattctaatctttcaaccagttgttctagactacaaaaaacgtcagacagttatgcg
gcattatcccattctagttttctggatgtggtatatccaagtttgaagaaaacaactgaagacgtattgcctcattctttacgtgccatttggaataa
acaacttccaaagttgtatgaaaaaactcttcaacccatagaagaagaggatattggttataaggattatgttgtttcaattgaagacgacgaca
atgttgatgatggtgaccaacaagaacaaatgattattgatgaagaatcttataaaactattggagaaaaatcaaccattgaactgataggcat

gtataacaataacaagtttggtaatgaatttataaggattcctttaagagaaactgcgttgcacgcacaatctctgaggtacgacactgaagcta
aatttgttaaccacaaggactctatacctctattttatgaaaacagcacgtgcacatgtaaggaacgtcttattgatttttctgagagacaactaca
acaactaaaacaagatggaatggataaaccaacggacaagtagagaggatcttttcaacacgtatacagggaatgcagtaatacgttcggc
agctaagcaagcactggctattgaaaaacacgcagcagaaagaagaggagaaaaggcatggacgacttcagcagcagcagcagcttctt
ctaattttaataatgtacaacaagattatactgatgatgatattacacaagtgtctattgcaaacagtgttttgaataaccccttttttaaagagatatg
caaaacttatagataatttagcaatatcttctttacctcctgatatagaggatgatgtcattatacacactagagatgcctccaactctacagtcag
agtagatggagccaatatctatttcgccataattgacggtgatttatgtgtataccctaaacaatatatatctgataaagtgctgtgtggttctctca
accgggaaaaggcactgttctataatagctccaagaataagtggacgtatggatgtaacctaaactttgatatcgttgacgctgccatcatgaa
acaccccgactacaaggaagagactacatctacaaaacatatacgtaaaatattgggtatcggagcatcggaaaaactgaacattacccact
atttaaactactttatccaataaatcaccatgtctatatacagcccagaaatgtcagccatgatgaatgagaatgaagaaaaaaacgttctcagt
aataggagggctgtaatattgatctttttcataggtgttgtcattttatcactcttgttcttaatcagactagatatagatatcaaatatgggaatgtg
ggcgagagagccactattgtaaagaagactagtcccatttttaaggaacctatccagacagccatcaactttaacaacaaaaagatttcccac
attgaagatggaggagaagaaagtgatgttatagcttcaaggactgttagttctctccagagagatcctcgtgtacttttaaccagttcatttgga
ttgtataataggacagattatatgaaaaaataaaagtaccaaataccatatcaattgttttatttcatcattagttaacatctttttaaatccacaatca
gagctaaaacatagtgcagttatacaatacaataatacatttctccctccaatgatacaatgttgatgttctcctcccatatacttttcaatcaaagc
cgagtcttgttccttggataatttatccatgaccagcttagaatttctccagtacatgggacgaaacactaacctcatactaggattcgtatcattg
tgggaagggtttatacaacttcttgatgtctttacatcctcagtaggcatttcacccaataaattaaacacatctcttataatagaaacagactcag
aatcatactttaaatcttctatgcattcacctcctattttagatttattaaagggtgcaaagaaaggggtttcttccttcttcttcttttggacatcattttt
acctaatacacttaaacctatctgttctcttacaaaggactgaaaattgatatctatataatgtacatcttcaaatttttggatattgcgattaacatttt
tgtatgattctggaacagggatcttctttgcagtcgtcatcattctggaacacgtgcaggcaggttctgggacaacacccatggatagggcga
tagcattggccagtttcactagccacatttttgtttgtggattaaaagtcttcttctttccaaaacgaagaaagaagacgctacatacgtctccttg
ctcgtcaccaattacacgtattgtgtcccttttgtttaatattttgtaggcaaattccagtagtacttcctttccttcaacaagaggtattccaagttctt
caaaatccttttctaaataatccacactctcctctaaagacaaggaagaggatgtgcatttgggcagcaagatttctgtggtgcacatattattat
tcaaaccttgagaagaggacatttttctctttactgttcaggacgaggagaattgggaggagttatgttagttagtgatacgtcttgccgctgctg
atgctggaggtagaatgaagaccacagcgtgaggggcgaccatggcttagatatggccacgccgagcactggcatgaattgcccctgag
agggtagccatttagtgttttcactcggtatttttccctctttattgagagttgccaaaagtttattatagttgtctgtgcatgtcgataccttgtcttta
aaatcgtccaatacttttggaatcctctcccagttcttgcgttttatttgcttgattctgggagtctttttttagttcaggcatatactgcaacattgtgta
ttctagtgcagcagacataaaattagttctatctgtgcatattttatcctcttctgttttaccacaaaattcactatgtattatgggcattaattgctggt
acacaatagtattatcctctagaccgcacatgttcatatacacgataactaaaaggttcatggtttttacactataaggaacaccagagaatcttt
catcgtcgatgccttgaactggttcatagagcaccacgttactattgttcccgtcttctctcttattggcgtccattaaaactccacttagacgtgat
aataaactattgtcattgtacgttatattgtcaagagaaatgcatttgtcgaattcgttaacaatttctgcccacttttcttcatccatcacgggttgtt
tattgttaagttttgatgatacagaacatcttgaatgttcatcctcttctatattatttttagcgagcgatgacgtcatttcgttcgtggctgcagttac
agcttctaaaaagtcagaatctttatactcgaacgctttccttgcctcatctaaccagtcgtcagtttctagatctgcctcctcttcctctttttcacct
tcctccatttttttcctcctcctcctctacatcatccatttcattgaatttttttaaagaaatctaaaaagtcactcgtgctgttattttgttcagtttttaaga
ttgacgtctgctcagtagatagacctggtattgacgatactattcctcctcctcctccattttcttctcccaatgcttctggattcgtatctaaattatt
accaaaaacatcaatcctttctttagtcaatattcccctatcattaattataccaccttcagtgtggctctcaaaaaatccacgaattgattcgaaaa
ctgacgatgataatttttttagtaacaggtgcactttttctggttgcattaaccttacttttcttacgggtagatttttgagggtcattaccacttttacgt
ttattattagaaaatgggataagatcttcagtatcagattctacatccgaccctgtatttttactctcattattattagcaccggaaagatgtatactg
atatctgataaggttgttattgtaccaaaaacacgttgattaaaggaccccattgtcaaatatgacatgacatttgccgctgcatttacagccagg
gctctatgaagtgcgtgtttcaaactaacgtcagtttccaaatcaatcaagtgtgttggaggaaaacacaactgaagcgggtctcgtacaaaat
ctatagcttgagacagatagaaaaatttgattgtccagaaaggaagtactagtttcttatcacattttctacaattacttttagacaaattccccata
gaaatattaccaacactatcatcacagtctttattattcttcctagcttcttgttcactcaaaaaaatctcccatccgcccacacaagtacagtacgt
gttttcaaatactgcaacttctttccttacccatgtgtcgtactcacagttactaacaattgcactattgtagtctgatcctttagttagcataatcaat

agtgtactttcaggtgaaagcatgagcggcgattctgccaaattatatatcgtatttggctttgttcttcttttagctgcaaaaccacccaactcgtt
tgcttgattttcacccgaagaattactgttcatgcacctaaagaacttgacatggccaatctttaagaacgtttcagagtccttaaattgtgtacca
gtaagacgacacacgtttttatgatgtccaaccatcaggttccatttcaccagcacatcagaatcgtgggagaaaagattcatagaaagtgttt
gattttcatcgttggtgtgttgcttcaaaacattcaataaatcttcccacctcttggattcttttctgttaccagacatattataagaatcgtactccaa
tatcttcctcttttctagtagggagaagtgtttcccaggagtctccagattcaaaaggcatgatgttaaacgtacgatatcatcctctgcttcggtg
aatgatgcattagaaaggtacatgcatttgctccccgtagtctttagaacagggatcctcattatgtccagtaatgcagatgccatcagaccttg
aggaagacgcatcatcatccgctcgcacgagtagaatagtagatcgttggtcttcctattgaggcaattttcattccttgattttgccttttccttac
ttctttcttgtctgatctgtcgtttggctctcgtcatcatcttgatggctggagcctttccatccgtaaagagtacagtatacctccctttaaggttttt
agacttgaggagatttttcatcactcggatcatcctaaaagcgtagtacaagatgaccccctccatctggctagcaaaaaatgcatctttgaatc
tggatttcaaaactgctgtcggggaagaattgaatgaagtagtaggcacgttcctgaacaagaaactgttagtcaaaaacttcctcttcttcata
acatctgtgtgtttgttgatccagtaatcgagaccctcatagtcgtccattttgtgattagaaactggaggagccatcatcatgtcttcgattgga
ggttcagttagccaatcaacttcctcttgggatgaagggattataataggcactctttcaattctagccttaattttctcgtgtttatattttctagaaa
agaattcagaaataaccttcttttcatccctgcagtaaaaattttcataaatatgtttcttgttctgcatccgtgaatgcatagtatcttcttccctgat
aacattaaaaccgtcatgtaaaacgtctattatctgttgatttgtagtattgataaacaggatcatctgacccatatttttctgtcccttcaacatggt
ggttgcaatttgcctgcccaacatgaccagatctgatacattataagtgggcaaatcaaccatgtcgagtgtgtagtcacaggtaggtgaaga
agaggaagatttcttgcctcttccaaaatccaggaagttcttggagttgatgttggcaagagaagtggctttgagaagtgagctagttgtgcaa
tcttcttctcctgaacaataatcctcgtcgtcttcgtcatcgtaacaatcttcatatccagttttctccttgatgactctcccattctgtgcaccgtacg
cattttcgtacactttttggcttctgggacacccatttctgttgatgttactcccacgggatgttagacgataatgaatgctcactgatgggcgcc
gtaatttttacacactctgcgtagtctggtgtttttacccctcctagaagaacttcgtggagctgctttaaaggttgtagtgcttggaatctgcagc
aaacgcagctctctacagagacttctcggagacttgccagaagcgagtttcctcttcttaataggggtactttcttcctcttcctgttgttgtttgta
gagtgttatagattcagtagggaagtagtttctagatcgaagaacagtgtagtcttcttcctctttcacagtctcaatagaattggacacctcgttg
aacacttctagggccaattcccacacaccttgttccttaagtgtctcgaccacgtaatcaaattcggattcactcaataactgtaaggaggaag
aatctccatccatgatacgaatctggaacacctgtctcctccttgactgttaactgatgtctgaccccttgtaaccactttactgatgttttataccc
tctcccgacggaagatgtatacgtcacaattttttttaaaaaataacgttatacaccaaaaatgagcgatacagggcagatggaagaaaatag
gcctgctacccagaaaaggagacctggagatgaagaagaggaagaaactggtagtagtaatgttccatattatgccaactttggcgatgac
gccacgtactccatgtacactggagaaggaaaaaggggtaaatttgtattagagccacctaaagaaagaagtgtacaaagggtgcaaaaa
ccacctaaagaaaaggaggaaagggaacaacgttctaatgttcggacacggagacctggtcaagagtttgaacagaaagtgctacaagat
cgatcgcgagaacggtcagaaaaacttgggcaaaatttggcagagaaaggattgcaagaacggcaaaagaaatatactccaaaggtagc
acaaacaatgacaaaaaaaataatcaggtttcgtgaaggaggaagaaaattcaaggcgccgcaacagcagacatctgacaaaggtgcag
caaccaatgttcttgaaagggaagaaattgagatggctgcagaaagagaacaaccagtagaaattacaggagatactatattaggtgggct
aggagaagaagatgacgaagatatgggagaggatgaattaactatacaacattcatctatggctgtatcacaacccgttcaacaaatcgttgt
cagttctcctataccgccaaagcccactaggcccgctcctgatattcccatacaagaagatatagtggggaaaaatattagccagttaccacc
attaccacttgatgattatgaggacgaagaagacgaacatttgtacgaagaagtgaatgatttcttagtggcaccaccaacagcagcagcag
cagcttccacaagacctcccaggcctaatattcctcctccacctcctcctgttgttgctgttgcagacgaaaccttgaagaacttggcttcaatt
gcagccttggaaaaggaggccgaggaacaaagagcggccgcagttgaaagggaaagagaagtagaggaacaaagagcggccgctg
ctgctgctgctgctgctgccgcccaacgggaagcagacgaaaaaagggaaagagaagcagaggaacaaagagcggccgctgctg
ctgctgctgccgctgctgcccaacgggaagcagacgaaaaaaaggaaagagaagtagaggaacaaagagcggccgctgctgctgctg
ccgctgctgcccaacgggaagcagacgaaaaaagggaaagagaagtagaggaacaaagagcggccgctgctgctgaaagggaaata
cttgcccaacaacttcaagaaatgaaagaacaaatgcgcataaaggaagaggagaggcggaaagaactagcagataaggaggaagaaa
aacgtcgagaactagcagccaaggaggaagaaaagcgtcaagaaatattagctaaagaagagcaacttgaaaaattgaatttccagttgg
gtacagaaatcacgtccaaaagagcactcgaacaaatgttagaagaagagaaggcctcacgctcacggtccgcagccagtgcacagtta
gcgatccaagcaatagaatatgaggatgaacttcctcaggcagtcgaacctcaaggacagttagttcctatggatacggatttgtacggaaa
aatgtacgatctcaataagaaattagaagtacagaataatacattaacttctgcatttgaagacgtgaacaaaacaaacgaacagaaccaatt

ggttgctcaatcccttgaaaaatccgctaaagccattgaaaaattaactagtcaaaaacatcttcctgtggatgatcctgcttttatgcagagaat
aataacagagagggatttttctttaaagaatctgggaaatgtttacaaaagagttctcggggtctattttacattaaaaagggacctttttaaatcg
aaggcattaattacagataaagaatcaagggatctggaggtgcgtctaacagatgtatcgacagatctcagggctaatgatctcaatacaata
ctggaaaggttggatgtatccgttaacatacgctctggtggaacattatacactaaatttacagaggcagacacggcattagcagatcaagttc
cttcgaggattgaaataagtaacagatcaagatctgccttattgccattttcatctgcaggtttggatactaattttactaatagttccgacaagtac
aatgaaatagtgaaccaactaagcagtataaatgaggctatgaatattttgaaagaaaatattgtcccaacattgaaccaaatcaaaattgatgt
caccaatctattaacagtttcaagctctcgtcaatatgctattgaagaaagggtgtattctgatgtgtcccgaatggattctgaaataagaaaatt
cctcgctataatgaacagtaaaatatccccttattttaaaggcgattggacggatgaaagacaacgctctattgctgacagtatttcctctcagat
aaaatcaaacgataaaattaaagagagtgttgctacactacacgatatcaatacaacctcaagaatacgtagtaatcccctcctgcacaaatc
ctcagttttatcatctccagactttttaaatgctgttaacgactttagaaattttctcgatatccaaggaggttctcaatttacttatgatgtcctttcag
gccaaaatattgatgacctttcactggcatcaaaaaccactgaaaaggttacagaattgtgcctcgaattatccataattttagacgtgatccata
aaaatgctttgagtttaaatttgcctgcaattacttaccccgcaggggaaacatctatggaagaaagtggttcattggctgttgacattagacaa
gaaattggtaagaacatatcagattctagcgctgaacttagtcgcacactgtcagaggcgttacaaatttttcagcaacaacagcaacaacaa
cagcaacaattccaacagcagctgttacaacaacaacaggaccaacaaaatcaacaacaattattacaacaacaaatagaagaacaacaa
cgggttcaggaacagcagcaacagcagcaaagggaccaacaacaacaggaacaacagcaaagggaacaacagcagcaacagcagc
aaagggaacaacaacaacaaagggaacaacaacagcagcaacagcagcaacagagtgaccagtttcgacaacaattattgcaacaaca
gcaacaatttcagcaattactacaacaacaaggaagaagaagagggggtgacgatggtgatgaagaaagagaggaaagagaagaagg
ggctgaaaaggatgattgtgtgcgtaaggttgcagaatcagtagcgacaaaatatactgctgacttgactaccttattccaacgagaagaaaa
taacttccaatctaaaatagcatcagcaaaattgggaacccttgtctttgccacccctccttcacctatcatgaacttgacaaaattgagagagg
aatattccacattcacaacccagtgtttttcaaaactaacagctgaaaataatagtattatgcgtattttccccgagaggattgtagaagtatgca
agagtaagaatctcaatttaatgggaaaatacttgtacattataactaccgcacaaacagaaatggaagatcgagtgaagaatatattgtctgg
tattttcaatcaaattgaagagttttcaaacaatgtaaaacaacaacaacaacaacaagctgcttcagcttcttctactaatcctcctcctccttcta
ctccttctactactcctcctgttacaagcatgcaagtttgtgagttggatgatcaacgtaccctagaaaaggctgccatagtagaggcaattact
ctggccaatgctgtacttcaaactacaaaatccgcttcagctccttccacggcggctgagcgagaaattgctctaaagctagagaatgggaa
aacatctatccgtatggaaaaagtggatctaagttcaggagctactggtgtttccgaccaacaaaaatggatcgacgaaagtacttccaaaca
agaattggaagatttcattgcagaagaaaactttgtagaaactgcacataatgaaatggatattggattaattttggatgccaagaagaacgat
ccgacccgtgatgccaatcttaggctcgtgaaacctcatggaataaatgtgcagtctttcccatattacgtactccgcacatggctaggagaa
acagatatattggatgaagatactgtacatcctgaatatttccgccaatacattgatcgcaattggaaggtggaagaacatgagcgtgaagata
cattaaaggcactgggtgtttctttatcagatacgttggcacacatcaaggactactattctcccagtgtcaaaaatgatgcatcaaaatcagta
ccatttgcgttgaacactctattgtacaacatatttgctatcgacggaggaatgatttctagcctttcaagaacagcctttatttaccgtaaattttta
aggcaatctatgacagataaagaggttgctcaaggacctgttcggtctcaactgtgtgaagcgacaatagcgtctcttttcacggcatgtagta
accttcttcgatcgtctcccttagccgataaagtagaaccacgccttcaagaaaaattagcggccgctgctgctgtagacacgtcaaccgga
gacatgtttcgtatacgagtttgtcatctcatgtacaattttatagtggcttatgtgaacctatgcaataatcgtataaactatacgttaaatgtattg
agagcgtcgggtctggcaaacaaaaaagtcgtggccggtaaaacaactaaagggcatacatcttcttcccaccggtttggatcttatgatgtc
acatatgatttttcagtattgtacaagattcttcaactacaaaaacagaatatctctttacttttagagaagggtttcaatgcatgggaatcgtgtgtt
gccgccatggcagccttcactgccgacccttctctctccatctcagatgccgaccaatccatactcttcccactagaaggggggagagattgtt
attgaaaaacatgaaaacgatgcagaaaaaaatgtcgatatggttcaagaattgtggaaggaaactgcactcacactcatggcaaaggaact
aaattcatactacaactggttcatatacagtaaggataccgatatggaaaagttggcgagagtgtgcaggatgattatcggtatagttaaggct
gttctcagattaacaaataaagccgagagtttagtagacactaatgctctttctgatattttcaagctacctgtgattccaattgatgataccaaaa
ctctggcaataaatattgtggtgttcactttgaataatgtaattaaaccatggatggtttcgttcaagcaaatgtttagacagaaagacgggggga
gtttcttcagcctatttctcgtttcaaaatatccaacaacaaaaacatcaacaaactgcttccattttagatgcatgggcttgcgcgcccggtaag
ttaacaaaggcagcacacgtgtttatttcaggatatgaaaatcatatcaaattaaaaaaagatgatcttttatggggtgcttcaatgaaattccct
gccgacggacgaggcactgtggtcgaaggatgggcccaacaatacaataacgaaagcgtattggaagattttaccgacttttcaattgaagt

aaacgcacctgcctctggactcttaataccgccagatcctttactttcttccatgttcggtaaaggaaatggtggaagcagcagcagcagcag
caaggataatacaattattggaaagggaggattaattttaaaccgccaagttgttggacaagaacaagcaccaccaataaacacgtcgtcgg
atactaaaaaaataagacgtgatgcaaatattgaaccaataatagggacgccttatagtgtaattaaggcaagtaaaggagtatcaatttcagt
actggatgatttcaatgaggatagtccagaagatttcgcccttaaaacttccatcatcaatgatgccatacgagaaatagggcaacgcatgact
tatacaagacctatatttgatcatcaaacacagaaaaatatacactacagttcacctaaaattattctcgaaggatcagatttaaagaatggaca
acgttcaggacaatcttgggctccttcttcatcatccttgactctggcctccgattggaatctaccttctctggagcttttatatagagaacttgcc
acaaaacaagtagagaaggaagaagaagaaaagagcgaaagggaagaagataaaggacaaaaacttaatgaaaaattatcatttgtcgt
gaataaagctatcggaactatccaacaacaacatcaatattctgaaaggggaggaggaatgaagaggtatcagcaacactctgctgatcaa
gctagtaatggtggcatagatgatatagaacttatgaatagtaaagatgctacttccatgagaaaggcaaaactggcattagccgttactaata
aaattgcagcagcagcagcaagggatggggaaaattcatcagctaaaccgtcaaactttggcaatagattggatgaagcaataaaccctgg
agcacttttattacgtagaggaggaggagtaagaggaggacaaacaccccagagttcaatgctaacaatgttccgtcctggacaaactggt
ggcaatagtagttggtggactactaatacaccccttattcaacgcacaactagtgttgggaataatttagttgtgcttgtacccaacctattggat
tcacaccctcctacatttaattaggaataaggaattatacacatttatttgtatgttttacatgtattttattcaacaataaaggttatgatcatacataa
tcaactggttgttttattacatagtttccattactgacccttccaaaaatgaattttgtacctccgaatacaatttcttccctcggtttacaaaggaaat
tgcacggagcgaatctgctcacccattcaccaacaacaccctcattaatcttcccatgagtgatgtgaggattcagtctgttgtcagcgagttc
aacatttctccctattgcatctgaaatgtgttccattattcgtctacgcaaatcgagtgcatcatctccaacacatactttggatacgagaatgttgt
tgttcatcccaaacaactcaaaattatccccagtgtcgattttgatagctggtgcaaaacaactcccaaaatctgttttcatcagagtattgtattct
tccatatttcgcaaatagaaaacagaaatggtcacatgtggaggcaactgagaaaatgatttggagacgtgtttctcagataataattttcccgt
ttctgaatagaaaggaagtaactcttcgctgtagagagcaaagaacattttcatttcttcacctttaaaatcaacttgatttatagtgaagattctatt
gttgtgtttgattgaaaattgtttcaccacttcaaacatgccattgatttgtttgtacatttgcttgactgcgcatcttctttgcacctgtggttgagta
gatgaaggggagatttggtggccattatctgggtggtatccattcaatatgttgtagccggcacaagggataactgggtccacaaacttgaaa
gcctctctattaaaccctctccctagaggaataagtccacaacgaatttcgtctgcgtcttctgccttgtccattaacatcatatttagatgcacaa
catatctcagaatcccattcaaggtgtgtttttctggatagaggcaaataacatcatccacatattccccatctacataaagtccagtaacaacaa
tctttaacgctttagataaaacaatagtttctctatgagaaaatattgccttgtttactttggtggagagttgttcgtcactgggaatgggcgtaga
ggccaagagtagagccttggaggaaaatttcttgtccataggaagtctgtgccggatattgtctaccatgttagacatagaagtattatcactaa
agtcaaagaagactactccattttctggagagggtttgttgaaggatagatgttcttccatggtggtggtggtggttgtgatggtttgaatgaggt
gtggtacagtcattatatacacgtgggccctttttaaaagggggagggagggggagagggttgccatcaagcaatatcttcgtccagaaaca
cctggtccagaaatggccataagatacttcctctatttctggagctattacatttctggtgtaatctgacattggcccgacccagaggtccaccc
tcggaacttgacattcggtcgagctagcggtccacccccctaaactggagcgaccctaaaaaatttttgaaaagtttttgagatggaggaagag
taaaattctctagtgaaaacagaagggtataccctctcatttctgggtcgaccagtctccagaaacgcctgttccagaaacacacaaaagttaa
tgtacgtttctggagctattacatttctggtgtaatctgacattggcccgacccagcggtccaccctcggaacttgacattcggtcgagctagc
ggtccacccccctaaactggagtgccctgaaaaaatttttgaaaagtttttgagatagaggaagagtaaaattctctagtgaaaacagaaggtta
taccctctcatttctgggtcgaccagtctccagaaacgcctgttccagaaacacacaaaagttaatgtacgtttctggaaccaacaatttctggt
gtaatctgacattggcccgacccagaggtccaccctcggaacttgacattcggtcgagctagcggtccacccccctaaactcgagtgccctg
aaaaaatttctgaaaagtttttgagattgaagaggagtaaaaaactcactatatgaaaggtgtgtagaacaccacatcctttctaggcacggcc
atgtccagaaacgcctgttccagaaacacgcaaaagttaatgtacgtttctggagctattacatttctggtgtaatctgacattggcccgaccca
gaggtccaccctccgaacttgacattcggtcgagctagcggtccacccccctaaactcgagcgaccctaaaaaatttttgaaaagtttttgagat
ggagtaagagtaaaattctctagtgaaaacagaaggttataccccaccttttctggatgcgactagatccagaaacgtatgttccagaaatacc
cataagtcctttcctctatttctggaaccaaccatttctggtgtaatctgacattggcccgacccagaggtccaccctcggaacttgacattcggt
cgagctagcggtccacccccctaaactcgagcgaccctaaaaaatttttgaaaagtttttgagatggaggaagagtaaaatttcttgcctgaaa
ggtgtccagaagtgtaaacacacacacgcattctcgcaccttgtcaccatcattaataaaaaatgtcccctcccgctcccccctcaaaatcgg
gtataaatagagctcaccagcacaccacagacatcattctcaagacatttcaagtactgagaacatcctctcctcttgtgatattattcaagaaa
acctactgaaatcgtccctaaaatggattttgaaggaactaccagttctacccccctcaaaaatgtcccagttgtattcatcagtgaagaaagttg

cagagcattcctttgccaatcttcatgacaaggctactcttgcatcaaaggttattaaggacctggaaggggagaggaagaaaatgtctaccc
caaagtcctcttctgatggacaaaaactggacaaggctatgttggacgatattatcaacgagtatcaggccgttaagagcactgcagataatt
ccattgaatcgaccatcaaggaaattgaaaatgtacttgaaagtgtgcgcagaaccaagattgaaagtgaagccaagaacagtgtaacttcc
agcccagaaaaagtgttttctgtcgaggatttagaaatctactccaaggggcgagtgtgcaaaggtctcaagttaaacgccaactgttcaaga
attggaggcaagtatgcagtgtcaatgagtatcaaaaaacacaacgtctcctcatttgagaacaacaacaaccaagttttctctgaagaaccc
agggattgttttatgcttgaaacaacctatcctcttgttgggttcgaaacttctacagaagatggaaatacatatgcagttttcttgactggtgttgg
gctagaaagatctctacctaaatatgtaccagttttcgacatgaatgcaggtattcaaaccctaaacatgactggtttgaggatggccaagcttc
ctgttctgtgcatgtttggacgtacagaatatgacaacttggaagatttttacatcacttcaattgagacgcagtcttttgacgaagaggaaaatg
atgccagaatgaggtgtcacaccgaagatttggagaggaagaagcgcatgaatgacgcaccagcgattacacctcatgtggccgtgtacg
actacagtggagacgggaaagaacaattgctctatatgataaccgagtatgaaaacacggctagttggtgcaacgcaaacggtgtggtcac
atctgacagtggattttctaacgaatgtgcaattagtgatatgaatgacttgtgctgttttgctgactgcatcgatgttactgttaataatgaagaac
atgaagaacgttctatgaatattgtggtcgaatctgacaggcgtctttttgatgctagtccttcccccatcaagacggaagaagatggagaaaa
ttcatcgtcatcgtcttcctctccaacagttcctcctcctacaccatacgaaggtaacgcagttgtggaggggggaggaagaagaggaagaaa
ttgatgaagacgaaagtagcaagtatgaaggttcagaagatgctcttgttatgaagaaattagccaagctttctactatgaaacaaatgagaag
ggttaagaatgaacctgcactcaaaattacttctgggggtaacaatagtagcagtagtatcaataacgaagatgatggtgatgatgacgatgc
cgttgacgctactgcattatgcccccaaactgaagctacagtgaaaaattccttcatggccccaaacgacgagagaactgaaaatattttgtat
gaaactatgcaaaattctcttgctaaaatttgtaataatccatcatctatgagcagttaccgtgtattcaccaacaaactccaagagtgtttgaata
ccatggacgatagtatccgtcgccgtccaaccatttggactgaagaaagtcaacaatttgctaagggtttgttgtttgatgaggttgtcacatca
attgtggcacatcagatggctcaagatatttgcaagtctgaaatatttggaggaatgtttaacgccaactctaccaacattaagggtaaatatga
aggacaaaagaagagtctgtatggaaacaagcacatttcttcctcgtgcttcaaaaccaacacggaatctaatgtgaataatgcactatttgcg
tgggtgaaatcgaaactccattctggcacagtcatacctaacgtattctccttcaaaatggcatcagaaaagccctcaaaaatgaagcgcaag
cgtacctctagtgcttcatcatctaacgatgaacaccaagaaccatcaacaaaaatgatgaaaaatgatgaaggggaaaaggttgcacaag
aatcatcatctccttcttcatcgtctacaccagaacaacaacaacaagctggtcatgacaaggaaactatcaatttaattcccctcagtttcataa
aaatgccacgcagtaatgtcaatggctcggcttcatatttgtctgaaatattcggtcaacgtctttgtggactgtctgatgcttccagcacatttaa
gagaatgtgcaagacttttgaagatcttgaaaatgaaatcatgaggagctcattcactagactgactagatatgagagggaggtaactcgctt
gtatgagaaatgcaggtctcaagctgtagatattgaggaaaatgaaatggatgttttgtctcaccaaggggaattgtttgccgagttcttggag
gacccgatcgcttactttgaagaagtactggagaatattaagagttggagcctagaaaacgttaacacccctaagcgcaaaaacaagtatgc
aaaggtactggtgagcgttaatgctattcgtaggacatatgaagaataccatgcgtttagcaagtttgtaccaatgttcttgttcaatctgattaag
agagaattggaaggagacaactatacccatgacgttcacttttcttccacttgcctgtggtacctgactgtaatgaccaggaacaggatttgcg
atgtgctccagtacatcaacaacaacaataatgataacgaagaaaccgatattgtggaggaagaggaggaaggagaaggagaggaggat
aaaatggaagaaagtatggacgtagaacaacagaagcaagttcgcaagggagggagaaagaagggtcaaaaattcaacagtattgggg
atcaagtcattagaaaatttgtgaaaagtttgtgtgaaaattcgatggtagtttctattgcaattaatagtttgatctctggaataagctggatgaac
aagaaaatccctcccggtttcttgaaggattctagcacaatcaatacccttgatgaggtctcaaggtttgtgtttagcgatgtcaaaatcaatag
gaaaatcaatggaacagatgataaatatgaaactgtttttggagtcagtacgcgtgtggattcacatattgtaggcccctttagtatacctgttga
tttttcaagcgcaggactagataaggcctcatgtggcaaattgtacgttaacaccatagacggaaagggcattttgacaatttcacccaaatat
gattcattaaacgatgaggatgttgattctactacaacagacaagctagagaaggatattttgcatttgtctaagcatgacaccttttcaatatta
ataagaataaggttcttccattctataatatttctcctagctcttctctcactgaaaagaaaaagacaaaattcaataggaagaagatctcatctgg
tatgagcaataataatggcatgtgtgtacaaactccttctagttcaaattcagtctcttccgtctcgtctattgtagctccttcatcttctgttctggct
ctatcttgctccctttcttctacaaagaaaaagagcatctggaacgagaacatgtttttgacatctaggaacatgtggaggtgtggatttgttgta
ccacccaaactttgcagttttattgttaaccatagacacgctgtaaaacttgtagctgaaactgcacctaaaacaaagttgtgtaggaatattatt
gataggaataggaagattagatttaacggtctaaagaaggtatgcaagagtgttagcgcctttaccggcgagtctacatatttgctcaataaga
atatgactgcaacttcacctagtgatttgaacctatgtatttatacttcatctttaaatgacccattgtatacttgtaaattgacccatgaagagtacc
aggacggaaatgctttggatgactatggtgcagtttttgtaaactatacatttaagagtatcaaatcctgctcgtccaaagatgaaaccgctgac

gacaatgctgctgctgctgatgatgatggttctactacttctacctcttcttcaactgatactgatgctgctgcaatccaagactttatgcatgtaat
gattaagaaaattgatgccatgaaagacattagaggaaagtacaagaaatctttggcaaagaaaacaaagaagcattaaatgtatacctttatt
tttaccaataaaaattatatgttttacatgttgtgttttattcaaattcatatggttcatccccatttaaatcgtctaaccccatatcccagtcaaacgct
gctgctgctgctgctggttcaatgtctgagccatagatggattcattgacatctaaaagttcatctctaaaatttcctggtgtacgcggttgacga
ataattggtgcaataaatgcctcctcttcttcttcttcttctcctctgactcccgtttccttgacagcttgacgttgcgcttttattatctcttcaacgttc
ctccctggagtagcgggaggaagatcaaaatcaatactgccaagctcagagtctatatcagtaaaggacatatcagtagcgttttcttcttctg
ctcttgctgtagacggaggaggaagaggagtaggcttgggtagaactggtgtaggtggaggttgaggggtagtagaacgagagtcaaatc
ttacacgggaaataggtggagaaggagcagtattaggtataggaggaggaggagggggactagaaatgagttgaaccggaaagggctg
ctgctgctgctgctggggaggaggaatattactgggtgattgttgttgaggagtagggggaggaggaatattattgggcggctgtggtggtt
gctgctgctgctgctgctgctgcagctgaggcggcggctgtgattggttgcctaaaaagtacagcgggtactgttgttgtaattgttgaggag
catatgggttataatactggttatattgataataaggatttaagtatccttggtaagtaggttgttgaacctggaaaggtgcctgtggtggaggaa
gaggaaagggaggaggttggaaagggggaaattgttgttgttgttgctgacgttgctgttgttgttgtttgggcacaaacactccagtcgataa
attaaatgcttgacttgtttcctccactgcactctttatcatttttttcaaccggtcttgctgaagctcttgctaaaacaccagctgcacatttctccag
catggagatagaatgctctgatttactcctcatccctcctccaaccaaaagatccatagtcgatgacatttttacccacacgttattatttatagct
ctaaaataacgaggtattgataggccacgtacagtggcgttagagttatcagaagctacagatacaaaattcttatcaaagttcttgttcggtaa
tatacgtgtaccaaacaggttattacacacagtagcaagttgttcaaagtatactcgttctttcacagttcgctgtatacgatcctccacgatactg
ctcattaaattgtttatctctgttgcatttccatcgttaagagatttcaatatttttgaacttaattcttgccactgaatctctcctgaccgtatatctttcc
tatgggaaattgcccaacaactattagccagaagagctttcaccaattcagcacccacatttgtgacaacaaaagtgtttgttctttttcctccga
actttttcttcccttgttcagatgacaataaagatgatgaagtaggaaaagccttggttttattggcctttaaaaggttagctaaaagagaccttatt
ctgtactgtatagccttctccttgaccactgcctcgtctgccaagtatgacacgtaatcgtattgtggcgcattttcactccttacacctaaaatag
agcccgaagttatagcacctctcaatggtcgagacacggtcatcatttttccagcaggcactgctaatgcctttgattctgtgtcacttctctttct
ttttttagagggaggaaatatgtgctttttcatttgttttgcccgcttcgatacagatgtagtaggtgtaacatgttcgtcttcaacattctcctcttctt
cctcctctccatctttttcttctttctcttcactctgttcttccttttcttcccgctgctgttcctccccttccccttcctctttcccttcctcttcttcttcatta
cctcctcctcttccaccaccttcttgatcagtgtctctttttccctcatcattttctaaatctgaacaatcgccagcaacacaatcgccactccttga
cagaattttactagtacgaatcttacgcctttgcctattcttactagaagttggcgcgtccaatttagagacctcctgacccatgtttcgttggaat
ctggattcggcgtcatcctcctttttttcgttgttggttcctaattatatgctttttttaaagcagtattaggtttatgtttaggtggcatatttgtaaggtgg
gataaacattacaagaaaacaaaaatattcaagacattattttattttgtaggcaccgctgttttagtatgtacatcaaattcaaaaccatcttccat
tacatatatctcaatctcttcttcttctcctcctcctctttgtttggacaaataactatcactacattttcttaaagctagacaacatagagtacaattat
acaccttttctaaagcagcctttgtagacttacaatcagaatacaatagtttatggtcaattagtatatggtcagggttagttatttcatcttcacata
tccaacaaaaaactgtagggtacccatctcttctaccgctacaatcataacaagtcatggtaatacatccacctaattttatgttaggtgtcttaca
tttagggcattttacagtcttaatttcctgtccttctttctcaaccttccattctaatgtcctcttttcatccacagtgaacactggatcctttattgattc
agatgtcatgttttgacgtacaatatttacctgtttaacaatttccatttttgcaccttggttaggtctgatacccgataaggcacgcctatctacac
aatgaaggacaaaagaatgggctctatggaacacacattctggccgctccaacaccatttcccctataagattcttgacaaacattttcgccca
taatcctaatgcactctcatggttcaatttttcatagacataaaaataggttgcaattagaggtatactcacacctacaacttgttcaaaatctctat
catcaaaacccccaaaaacactgtttaagtaatggactttaaccctattctgaaacctctccctggtgttgtggattacagatatgtcgttgagaa
gccccttcatattctctggcatacgatcagtcgagtgccagttggctagcgctagaagaaccttgccctccggagactctgggttccatcctc
gaatcttcttttcgggagttctttttagtatagaaacagcctgcctacaatttgtcacctcttcatttggaggtacaaaatttgacgccaagaaacc
gcaggacaatctcatgggccatttatggagtccgggaccagtatcatatcttactagctcatcggtgggagggtcatcaaattcagcatcttct
acatcctggaagtaatagtcttccagttgttcgggaggagggtacggcaatactggaggaggtacatccactgttacaatgtcttccatctcat
taggctggtcacatacattgggtagtaaacactgggtacagatcagggaacatttgctatcaccagtttctccctccaccatcttaaagagttta
acgggcggtctgaatctattatccaccacagtaaattttgaatccttgtctagtgcaaagtccttgtccatcttacacatctccatgaattcggcct
gaagaagttccacagtatagaagcatttctctgctcgcttgatgagttcggtgatgagaagtttgttggctgcaatgtattctgggtcgacatttt
ctcctgctaaaatcgccctctcgaataaagatgaaaattcgtgacatctacctataaactgtaccatatattctccacaacatgggcacctaaaa

acgtcctttagtttttcactaccagccgagaccttttcgtacacatacattaggtgttttatagtacaatcagggcatagttttgcagaacacgtttc
ttgtcgtgcttcaaaagtgtccatgaaagcatgtacggggaaataatcgtaagaggctttcatgttttcatgagaacagttgaagaattttggttt
catgtccaaatccgcgtagcacaacccacatgttttattggtggtattcccactcaatatctttgtaacgtgtgcttgtattacctccacttcgtcag
atggttggtacaccatcgccgcttgactttctccccaagactcttcttgcagtgtctggctaaccatgcatttggcgattgcttcttctagatgcat
attttcagcgttttcatgttcgaggatgaaagatcctagttttctggctgaaggtatgccgtgtctagataccagggacagtggtgtattttctaag
agcgaagcgttcggggagattattagaccagaaggcgacattacggatttctttggctcctctactctttcagtatctgctgccaaaagatagg
gaacattgtagatagagtatctgtacactatatcaatcatcttggctatttttgttccaaaaagaatatggtgtaattcttggggcacatattccata
gcgtcatacacgggcctggcagcgtacatgcgaagatgacccttattgtagtggttgttggtggatggaccacttaaaaatgcttctgttacgg
agaaggccatattttcccaaatcctgtcattgtttcttccttcatgcatatcagaaccaaataaaacactaaacgttctagcgtctcttatttttgtcc
ccacttgcaaagcagagaagggccacatgtaccactccttatgaaacacatccttcaaagtgttgagagaaactaccacatcttcataaatga
cacttttcatgtggcggtgcttcttgttctgatctcgtttgatggcaacaaggagtaagaaaatgcgtatcaaatccatatctgatatcatctcggc
gtacgcctcacacaacgcattcttctcctccactgcatcgcctgcatttccacgtataatttcatccaacatcaagcgcgtatttttcatgacaaa
catgccactataaatacgatgaattagatccttgtatgctggcccatcttgtttttttgcaactctttaaatgtccaatgaattcttgttcgttttggtca
agagttgtaaatggaactgggtcgtcgttacttttcatcttgatgttcatgtcgagggctttcttcattaatagagatatctctttcaaaagaccggt
tggattaatcacaactgtttctccctccaaaagttcatccctataagaattgcggttgcagatgtccagtatggcctccagaagatcaccactttc
tcccactatacccacaattcgttcataggtggacctcttattatttctgcctcccattaggtcggtgatcaagattgcaacttctgggacagcagc
tgcaactatggatgttacaccttccatgttgaacgagggaatgaatgcgataaagtctggggcctccatcttatataccccttcagatgtgaaaa
aaaacaacatgaggaaaaaagtaaaaaattcaatatacatgtatgtctttattggctaggaagaacacatttcacaagcaccagggttattgat
agaacataaaagagcagcagcaggatctggaacaacaatagaggacttcttctcctcttcttcttctcggacaggagcagaaaaagctgcaa
caggagagggagcttccttcttgacttcttggagtacattcttgtcgactgtgaactggacagctctagctgcgccctttgtgcgtagataatag
agagtcttgatccccttctcccatgcatacatagtcattgaccgcaccttgctgagttcgggttcttccacaaacaagttgagggattgagcttg
gtcaacaaacatacctctctgaatagccatgtccaaagtagtacgaggattaatttcccatacagttttgaatagttccttggtcgatttagggat
attaggaagcgtctgaatagatccaccacttgccataatcctctgtttagttactgaattccattctcccagttttatgagctctctaatcacatattc
gttcaccacttggaatgatcctgaaagtacattacgattatacatgttggacgttaaaggctcaaaggattcagagttgcccaggatctgtgca
gtggatgcagtaggcatgggagcaacaaacattgaattgtgaacaccatacttcataatgtcccttctcaattgctcccaatcgtgaattggca
aagagttgaaatatatgtccctattcttaatattttccttccccatgtcaaattggaaaatccctttgctcaaaggactaccctcaaacagctcatat
gtttctcccttttctttggcaatttcacatgaagcttccaaggcaccatagtatatagtttcaaaaatcctcttgttaattagtgccgcttcttcagatt
cgaaggggattctgagtttgaagaacaaatctgctagtccctgcacacccaatcccattggcctagttttcatattggaaatgcgggtcttgtca
actgcatagaaattgacatcaatcaccttgtcgagatttctggtcatgatttttacaaccctcttcatctcccggtaatcaacatagggccttagg
gaagggatgggagaatacttcacaaacttgttgactgcgatagaagccaaattgcacactgcagtttcctccgaatcactgtactggacaattt
cagtgcacaaatttgaagacttgatgatgccgacattttcttggttagactttctattgatggtatccttaaagcacacataaggtgttccagtttcg
atacgtgcagaattaatttggtcgaataatgcacgtgccttcaccacctctttacctttgccttcagcctcatatttctcgtacaacgccttaaactc
ttcgccatggacgtcggaaaggccagggcactcgtgggggcacatcagggaccaattcttcccagccttcactctctccatgaagagatca
gatacccagatagctgggaaaagatccctcgtcctcaaatcttcattaccggcattctttctgcagtcaataaagtccttcacgtccagatgcca
atcagaaatatagatggctgcagctcctctcctcttgtctcctccctggctaacctttttcacagagacgttaaagatttggaggaacgccatga
gaccagggtgggtaccactccatgacgaaatggggcttccttttgccctcaaatcatgaaagtggatgccgagtcctccagcagtcttggag
atgattgccgcctccttaagagtatcataaataccctcaatgctatcatcttgaaggcccagaaggaagcacgaggaaagttggggtgtgact
gttccacaattaaacagtgtgggggaagcatgggtaaaatagtgcctcgacatgagatcatatgtttcaatgacagatttgatgtctgatccgt
gaatgccgacagccacacgcataatcatgtcctgagggcgctcaaccaagattctcttctttctatcagtgggggaaccaatcttgatcaaata
agagtattctagcgtcctcagtccgaaacaggtgaagagataatccattttgtaatcgataacagcatcgagaatttcagcattggccataaca
ttttcatagtaggtatcattaactactgatgctggtttaccagttccagggtggattgcgtgcctcaatttctgagttgttgcactaaaactattcca
ctctttggtgtttttgtggatgttcgaacagatgaatcttcctgccagttttccaaaatcagggtggtcaacaatttttgtctttgcataatcggcca
gaaaatcgtccatttcttggaaggagatggtagcgggaagacggtccatgatgtgagatgcaagttcttgagggttaattgcgttcttgtcaag

cttgggcacatactggttgactggaagacaggcattttcaatcctcttgattatcttttcaagactaatttcttgcttagtgccattcctctttgagat
gaatgattgttgctggttagaacccattttatgttattaaattgggatgttgttatttggagtattttcctatacgtcttctcggctcgaggagggtga
aagatgtgagtgaagaggcttggacggtagctgtatttataccggcccggtgaaggtcgctggatatttttgaaggaggggaagaaatatat
agtattatacgtagttctttgatgtagaaagaaatcagaaggtgaggtgacagctgctgctgctgcagtattagtagtgcaggtgcaaaactgt
tcctcaaaaactttttaaaaatttttctggatcactcgagtttagggggtggaccgctgtctaggcctgatgtcaagttccgagggtggaccgct
gggtcgacccaatgtcagattgcaccagaaccgactgttgctccagaaatgtgcattaacctttgtgtgtttctggaacagacgtttctgggca
tagctgcgcccagaaagggaaggtgcaacatactgttttcgctatagaattttactcttcctcactctcacaaaattttgaaaaatttttctgggtc
actccagtttagggggtggaccgctgactaggcctgatgtcaagttcggagggtggaccgctgggtcgggccaatgtcagattgcaccag
aaacgactgttgctccagaaacgtacattaacctttgcgtgtttctggaacaggcgtttctgggcatagctgcgcccagaaagggaaggctgt
atcttactgttttcgctagagaattttactctttctccctctcacaaaattttgaaaaatttttctgggtcactccagtttagggggtggaccgctggg
tcgacccaatgtcaagttcggagggtggaccgctgggtcgggccaatgtcagattgcaccagaaacggaatagctccagaaatgtgcatta
acttttgtgtgtttctggacaagtcgtttctggtctaggaggcaactagaaatataagcttgtagcctagaatttagtacaccaaggataaagaa
atataacaaataccatactatgttttatttgagtaatatagtatccaagttaaaataacaatcacgcacttgtacattaatagtctgtaaaagatggg
caaacgtacacctgagagtattgtcccgacaatcgtacatgaaccctctagtgcacgtatcacgttttcgccctattaccattgatgcacaaattt
cctccttcattctaatagaggcagattgtttgtcaaaataacactccctatagtaacaaccaggatttccattggattgacactgtagttgttcgaa
cggttttcctgtataacatttcccagctgcgtggtagtgacgcacaaaactcatgcaagaatgtgcgtgggtagtttcatttctaagaatctccca
cgcactagaggcacagaaaaaatcaactgctgcgtggcagtgacgcacggccacacacccgcattcttcacaccataaaaggacatgatt
cgtgttagtcgtcacatctctcagaaacccccatgtgcattgacgtcatggggaattgctattgcaccaccttatggtgtataaaagtggccctg
ggtggcatcggttagtagacagaaacaaaccgtcaagatggtgtcgtctattacccacctctctctgttgttcgtcgtggctgtagtagcttccg
tcgtttttacaactgaaggagctagtgtgagagtgaaacggtgtgctgttagcccgtgccccgacgttattgaccccgaccaccgctgccaa
gggcgactgtgccgcaggtctactcgaggaggtgacgacgacgacgatgacgatggaggaactttcgatacagtagggtctggtata
cttggacgcaaaaagcgtgccgcacctccacctgaggatgaagaagaggatgatttctaccgcaaaaagcgtgccgcacctccacctgag
gatgaagaagaggatgatttctaccgcaaaaagcgtgccgcacctccacctgaggatgaagaagaggatgatttctaccgcaaaaagcgt
gccgcacctccacctgaggatgaagaagaggatgatttctaccgcaaaaagcgtgccgcacctccacctgaggatgaagaagaggatga
tttctaccgcaaaaagcgtgccgcacctccacctgaggatgaagaagaggatgatttctaccgcaaaaagcgttaaactacgcacgaaagt
gacggtggttgaagaatagactaatattgttgatatgttaacccctttttttcatgaaatgtgtacacacctgctatatatacgtgcatatttgaataa
ggaataaagtttatctgcatctgtatttttttattagtatcctttttgtgggataaataatcggtatttgtgtccaaagatagtggtgccatgatgccat
atacaaagcacgtgatactttccaagaagcagagtcgtatgaccattgacgtcaatgcgtgggcagaggcggaggtggtgataaagcgttt
ctgagaaacattgggcgtatgacgtcaactacattattcttcctcctcctcctcctattgcctctgccagttcagtattttatttttcttcctatacaat
aaaaagtatcagatgaatatttttactgtttttctgttcatccctctttcctattgtaaaaaaaccaataactaactaatcatggataacttgaaaggg
gaatttgttgcgcttaaaacagacctcacccattacaaaacacagttggatagatctatattggtatttgttgatgttgttggtagattatatgttata
gtaaatagtgaacaaacagctaaaaaggaaggtctagcaactagagtggcaaagcaagccacagagatacaacaattcaaggacgaaat
aaacaacaaatataatgctctaacaaatactttggatgatatcatctacatttttgatcatggagggagtttcaaaagagcaaaacataaggcca
taattgaagcgagggaatactctaaaccgctgagggaattagagtgcatgtttacgcgtatagcggacatgttaaccttgacttttatgactgtg
tacaccaatatcattactgaatttagacactctagtgaacaagccactaatagtataaatgtcaccctcggacgtcttttcttgtgtgacgacttgt
gcaatcaattaccaaaagaagaggaagaagaggaagatttgaaacagaaattcattactttccatgcgaacctatacatgctggacacacgc
ctaaagaaagatttgataattttcaaagatgtcatacaacaacttcacgtgattttgcaaaaggatacctatgctgtaaaagaaggtgtggccatt
agatgtgcgaaacagatgaacgaaataagtcaatacagggacaacctcaaggataattacaatacattttcaaacattttgaatgaaattgtct
acatttttgatcacgggggacattttgaagaagtaaaacacaaagccataactctgactagaaattacttgaaaacactcatgggattaaaatg
catgttcaaacgcatatccgaaatgttgtcattgacttttctaacagtgtacactaatgttatagcagaatttataaacgctagcaatatttctgata
gagagatcaataattatcttgtccaacttgtaacatgtaacgaattgtgcaaccaactccccaaacctaaacaataccgtcccctcagtttgata
gataacatagcttatttttctctttctgtccaaaaacatctgagtgggtttctttagtatgtggagtattattgtcttgaagctcataagtattcaacctc
tcctactggtaacatctcttcctctctataatcccaatatggatagttgttgcctgatatcgaggataacaccagaactggctggcaagttaacct

ggatcttcataccagaaaacaacttcaagattgtccagaactcactcccagacgaccaagttatctcccaattcagatatttcgaccatagaca
ttgctatacgtttatggagattttgatggcaaacattaaaatccaagacaggaaacaaaacaccacagccatatgtgaattgacaactggaag
agaaggacttttatgtagaagaaccatacctgtatttttgggttcagaggaaaaacgagaagagttattggggaatctccctgaaggtgcaga
aattttcaggcctagagaagttatgcaagtaattggtactctcttggacaagaaactagaaattgacgacggtatagcttctgtaaaggctgcc
ctctgtgctggttcatcatcgttatacctaatcatgagccacatagtgaaaatgaccttttctgctatcacaaacatgaaggatataaacgaaga
atatttcgtagactttatatttcgtcataaacaattcctcaaccctgaattcttcaagcaccttatatctttgctcaagaattccaggaaggaacatgt
tgcccatctagtaagacgtctagaacactttctcatgctatggaccctttccaagatgaggttcacagaaatggaagaaaactacttcccaatct
ccagcgatagtgattacggcatctgtgaaaaatgtgcacgaaaaactcccaaatacaagctccgtatttttagggaacgaaaatgctgcgata
gatgttgccgtctttatcaccaacaaccgcctccggaggtgtataattgggatggaaaaataacccaacaatccaataaaggctacattaatg
caggcgatgaaattatcggcatgctaaactcaaatgataagggaaaaacattccctcctatacctaagatggttgtacgaagagtggtggac
ggtgtctacgggcaaggaactatcctgtcaaagattttgaagttcaggcaggcaaatatccccacgtgtctattcgtgacatgcaataaatgca
ataggattttcaggctcactatcttagggcctacaagaaacatcctttgcccaccttgcagaaagaaaagtgttgcagtaaatacacaacaga
aaggagaaaataaaccttcgtttgtgcaaaaaggaacaaaacgtctacgagtggataccggtagcaacaagaacacgttagaaaaattctgt
tcctgggaaagattcaatactgaagttttgctcccttggcttggctacactattgagtctaagtggcagaactgggaatcttttctgggttattcga
gtaccagatataaggaactgtgggcctttgtgaacaaacaggaaatatcttccatgaaagactcctacataaaaattgaagacatcgaccagt
tattgaggagtatcttgcaagaccagaagggtgtatttgagaccgtctgcaaaataaagagcagagatggtttgtgaattggccacactgatt
ccgataccgactaaaaaggctgattgatggcaacccccccccctccagactcagccgcatgagtataaatatggccacttctcacaccaca
gcatcattccctcgtcatcggtcctaccgtcaacttccattattactccaataataccaacaaccccagaaatggagtcaatcaaactgttcacc
gttgctggtctgaatatggagcaagccaaccaagtggctgaagaaatcaagtcagaatataaaaccgaggaggaaaagaggattgcccag
gaagtgtttgacaaattcaccaaaaaactcattatgcaagtagatacgtctaaacacttacttacaagagaaaaccccaaccgttttgtatcccg
ccccattgtccatgaagatctctgggaaatgtacaaaaaagaggttgcctgtttttggacattggaagagattgatttcgaaagggatcctaaa
gattgggagaaactcactcaagatgagaaggatttcattctccagattctggcgttctttgcatcctctgacggaattgtaattgaaaatcttaca
acacgtcttcgtcaagtggcgcagattccagaagcgaggagtttctttgacttccaagttggaatggagagtattcatggcaacgtctacgga
gaactgattgatagactggtgcccgacgaaaaagacaaggctatcttgtttaacgctgcacaacacttccccgccatcaagaagaaggagc
agtgggctattaattggatgcaaagcaataacgatttggcggaactaattgttgcctttgctgcagttgaaggaatcttctttagtggtgcattcg
catccattttctggatcaagaacaggggtattttgcctggtctcacctcctccaatgagttcatttctagggacgaaggtcttcatcgcgactttg
catgcatgctgttgaaaaagggttttgttgataccccatcaagagaaaggattcttgaaattgtcactgaagccgtccgaattgaacaagaattt
ctcacagtttccctgcctgttaaattagtgggaatgaactgcaagttgatgagccagtacattgaatttgtggcagataaactattggttgaaatg
ggactagaaaagcactataatgttaccaaccccttcccattcatggacaatatttccctcgagaataagaccaactttttgaaaagagagtcg
ccgagtatcaacgtgcccaggtcatggcttctatcaataagatcaagaaggaccaacaaacccaagaaactggttctcctctcccaattctga
ctgcacctcctccagtctcttcctcatcatccgaacaagaagatgttgaagacggcgtcggggactacatcagttatgacgatttttagttccac
tattgtgtcaataggttgtgtattgtattattattgttataatatttttaaaaaataaatgttctataagactaaaacaatgaatttacttccaatattcctg
acaaccttttttgttgcggtagatgcatgctcttgctctaccatctgccttttacctgatgggaagaaacaacccttggtttttgattctgtattagaa
gaggtggtataccctacagatgtgtgtgggccaaagggagctggcgaattattcactggtgtggatcttttgaccctctgtataggaggtaaa
aacaatggaggtgaatggtcaggaaaaggtccttgtccaaggatcaataacgctgtcgttgaacgagattactcccttgacgaggaggattg
taaagggtttagaaaggggttccgaattcctggcactgaccattttcatactgtcttttccctttgttgggtagacagagatatgcacgccaagtg
ggtgcgcaacaaaataaaccctggtatagtaactgatgatgaagatttggtagattctggtattaggactaaatttaaatactcttctaaaattttt
ggtaaaggattcaatccgagacctctttactccctcgactatcaagagaggattaagatattaaagtctcattttaacaagaggacgggtaattt
ctttgctcgaggccacttggctccggctggagatttttttcctcgcttcagagagatgggcaacttttgctctagagaatgcagtacctcaaatac
agaaccataacaatggtgaatggaaagatattgaaaatcgtgcaagaactacgccaggtgccgcgtgggctgagactggaccaatattta
ccaacacaagaagaaggaatatctagacaagaagaagaagtacatccctatccctcatgccctctacaagattgtgtacgacaagaataac
aaggaattgttccgtgtacagagtgatatgtcttggaaataaaatacataattaaagaatttatattgttttattttgtcatttatttgatacatttgttgt
atgttttaaagacaatacagataattttcctccatgacgaaacactggtgatatgtctggagctctgatgaatgtcttgccagatgccatcctagtt

tttctcctggggaaggtgacgtccgtttgagagagggttcgctgggtttcaactgcagccaagaaaatatccctttgtccagtgacggtaaatt
gtgtaccgttatttagtgtgatgacattatccctctctgattgtttcaaagcttcataatcttcctcctcatcttcatagtcatctgcaacattctcagtg
tctaatagaacaataacagagttgggggaaaaatcagtcgtgctacgtccagactggccagtatttttactgctccatggtgtaggatcatgctt
tctaggcttgtctgaatgtagaccattatagttggctctccccatcatatcatacagtgtacacactagtgtattaccaatgcgttttccactcttatt
gttatcacaataatcctcatcttctccaacaagagttgagtacacactggaatctgttagaattgtatgagaaactaatgcctctttaaaaatatcg
tcactctcttctaaccagaaatctgatacactatcattacatttatttgcagtattatcaataaatgaattgagtaaagggtgtaaaggttccacgcc
aaataaaggtataggcacaaactttgcctcatccgtgcgaggtaaaaatggcctgtcttcgttgtatgattttgaaaaggttgttgcaagaacac
aatttcttgcatttttaatgcctcagctgatatacgtagacgtatactgtttaacttttgctctgttgtctggttattaaatcctccacacaacgagta
catggcattatcgatattctccctcgcatgatcctctatgaccaaaccagccttggtcattagaaaatggtctgggtctttaccgagaggaagaa
tatctgcattgaattttgaaattatatcattgacaacacgttcacagtgcacagatctctttatgaatttagaggctagagatatgaaatgtctattta
aattggaacctagcctattccaaggaagtgaattttgatgtcttttattatccctctcgtcagaagggaacaaaaattgaagtaatctggtccaatt
aaccgcagctgcagtttctctctggacgaatgtttgacgttgttttagtagctgaagagacacttgggttggcttgtcactactagctggaagag
ggtgttgaaagccacagacgccaaatctgaacgtcatctttagattggtgtgtagagattttgtgtattcctcttcgtggatttttcttctcctttcatt
gggcacaaactcctttgtctttaattcttccaagaagtcttgtacattgtcctcattcgcgaacgttaccttcccatcttgccctcgtttcatcacgtc
gttgttcttaaacaggcgcgcagaaagctgcatcagggacatttcttccaactttttagggtcttcatggatgtccttcttcctcttccccgaaga
gataaaagaccctataacattctcaatgcaatgaactgcatactgtctcaattttctgtttgtcatccctgaaccaaagcctattactggaacaac
catgccgcacttgtggttgtatgtgagcccacaagcaactgtcaatttcttcagaaggatctcgtatttttcacttccattaacatcgttcgtaaaa
ctctgtttatacttgtgagcaacccagttcactataaataacaaggcataagaagcagacacctttcgtcccaaaagtttagaaatttgaaggga
gagccagtctgccatgtagccatctgattcgtgttcggcgataggtgtcctcatcattgtgacggatgctttctccactctatctgctacaaatttc
actgctttaacaatactttttctgcacgcatttttgtcattagagtttattcttgcatattggagttcacaggggtaatattcttccccgtcattatcttca
ttggaggatgaatttttagttttgttgtcgttctttatatactgtttcaattttgagaagggagtatccagtacgttttctctacatgcactagccgcag
aaagagtgaaaccgagcaggtaaagttgtacacatgaaaaggcacctccaaaaatattaccaatgaaataagcaaatattaatcccctgatg
tggttgctagggaatctaaagttgtgctccagactataattcgtttcaacgttaaagtacctttccaatctacgaggcactttaatgtttatgtctgg
aaggatttcaaaattctcactaaacatacccgacaacttgtttgattttcccaatacaatatctgaccttatgtatttttcataaaagtcaaaatccgt
catcgttaaatcagttgggggctttttgcatttttaagagtatccgaaagatgcaccaaaccagttctgtagttgtagtttgccattgcttcatcgg
gcaataatttgctcacttcattgtatgcatcgacaactttttctggagcaatttttttcagcatataaacgttgccatattttcttatcaggatctagtag
ttcaacgtgtctagattctattgtcttcacatcgacaagaaagtatccttcacgtatactggtgcaggccaatctctccatcttcttgttgtagtagt
ctatcctcttcctgaagcaagatattatggaagcaactccatccaacatgtggtacagggctgattgggtacaccactcgtcactatttactcca
tcacgagcagtgaagaaatcttgaaggaaactgtatatagagatgaacgaaaacagtcttctggatgatgttttcagtctcatgtcagttctctg
cataagtttctcgtgccgtataactctgtctacgaattcccatgtcaggttttcatttacgtattcttttaggagtgccttttctgcttctatagttgtagt
agaagttgaatccataatgtaggaatgagggaggtactttggcacgcacgtgtgcactgaccgagacacaacggtgtactgaacaagggtt
aaagacgttggctgttttacacctctcagaggaatccctctaccagaataagtcctttcatttttttaattaacacacgcaaaactgacgctgaaa
tcgtttcgcaataatgggagaggaagtgggaggagaactttcgcctgtggtcgtagggatactttcggtggtgttaaccgccataattatctct
actatatttgccatctactacttttatttccgtccaaaggaagggttaaagaaatcgcgagtcagtaaagaccccttggctatacgcgtagatgc
taaccatgtcctctccaataattctccctacatgatgttcaatacaaacattaaggctagaaatacactaacagaatttgatcccaaaaatatgaa
tgaagaggaacgagcagaatttttcaagatccattattttactgggttccctaaagatgtggaaaaggaacaggtgcagaagaagaaggag
gaagaagaagaagacgaagaggaacagacgacagatactaaccccaaaaataagaaagtggaatatgtaatgctcgaatctctgagagg
ggataaggctggatttttgatagatgagaataagaccatgtctatacaaggttctcctggtacaaaaactgtctgtaagaaaacgaccctacca
aagtttgtcccaacccctaaaacggaaaagtatgttccttcagatgaaagatttttcaggatggaagaaggaacaggagaaaaggcagaac
attattccttcacgtctgttaagacaggaagaaccttaatcatgggctcttgtggctaccctgtgatgaaggatataagcaaaggtacaagcca
acagtacatgtacacatttgaaccaataaaggaaaaaaagtaatgtctttgtttatttccccctttttttagaaaaattggaacaacaacaatgtcgt
tggctgtgacagaagattacgggcacaatgaaaagttgatcaaacggttacaaacctctgtatatcacacccctctattaggtgcagaccatg
taatgaaatccatatcagactacataatttctcgtcgattcatgaactacacaaatttattaaaacaagttgaatatgttttcgatgaagaaacagg

agcagttatagctaatatctgtctgttaaaaatcctagaaagatgcgcacagaaaggaggaatatatgatgcaccagaagatgttgcattcttc
aattctaagatgggggaagtaacgcgcctatttactattataggaggtaggcccaatatgacggtgcgggttaattttaaacatgggcagaca
aataatcctgcctatggttatctcacagatgataatgatactactactgttactcctcctgttactcctcctccatctccagctgcaagaagatccc
ctttttcacacgcactctcatatccgagtcgtcttcagttgaccattatgtattgatgcatgataacccaaaaagatcttcatttaaggtgtatgata
ttcacgcagaaacctttccccataaagctccttctgttcctaccttccccctaaaacctcgtttgaaatttctgacgtgactctcgattgttcaatg
gagattttttcacgagacagggatgttttagacaatgttcacgactatattgctaacgaccccgtaccatttttagtggatgttgtgcaccgtgga
tctagtctccgttgagtaagtactttagttgcagccggtctcattagacctgaaaaatacccttcatttgttagaaaagtgtgcttagatcaaaatg
gcatccacttcttcatcaactaagaaaagagtacatgaagaagatgaaaatctcatcccacaacccaaaaagaaaaaatcaaagaaagtact
accatttcctgttgacaagtatagagctgtggataaaaaggtggtaaatctcatacacaagatattagatcaagaaaaggaccacctttctagt
accgaactgcaaatgataactgaatgtaatggtgcgcgagaagatctgcttaaacatcttctagacgaaggagaatttaaccctactataattg
aagtagtatcatccatgcctattgaaacaatatacgaaatactctcttcttctgctgacgacaagaagtttgtacagatatcattatcaatgttgatc
cacatacttttcttcgctgataagggtactatgtgggtatccaacgcgtgcgttcaaaatgttttggggaacgactataaagtggaatttgaaaat
atacgtaaaaagtatctgatattggaagacttactgaacggcgtttcaaatcattggtctgaacatggtcctctttctcacatgctccattcttcaat
ccctattgtacaagacatgttattgaacaggctggtgcgttactttagcacgtatgatggagatgctcaattcgatatatcattcataattaatagt
gtcttgtggggaattgataaaagtgttctcaacgaattgacacaattgatatcgaggggtgttttcattgtgtcgtacgtaccgatgcgtgtacga
acaccttcaaaggacagtaatcggccacaaaatactccttcacaaaatatgtcagcactaggtatgaaactcaatacattttcatccagaatctc
agtgtacagaaacaatacctttaaaaaactaaccgagttagtgcataactttgattacggttccaaagatgcatcatcatcatctcctcctcctcc
ttcattatcggacagcgtcaacacttttgtgaggttgtacaccaactatgacatattcttaaaggtgatttccgactggaaaatgccttatgggttc
tttaagaaaacttttgacgtcctttattctaaggggttgatgacattatcagtgtctgaatatacactcaaaaaagagttggttacgtttttgcgcgc
cttgaaggaaagggaaattttaatctataaaatggagaagagagacattatatgtatactgaaaaagtctttgttcggatttaatttcaggtgttta
aaacaattactccctctcttcaaacactttttaaaaattgaagaggttaaacatatagcacgttttgtctttagagattacagtctcatgtgcaaaac
tcaaaaagatttgcagagtttccctgccatacagtctgcttcacttttcatggaagaattcccttggcttgcaaaaacttggatcgacgacgatg
atgatgaaggaggaaagggacataccctattaacatttgctatagtgcacagatatcccttaataagccaacttatttcacacccaattttaaaat
cgttagtgaatactacatgtagagacaagcactttactcccctcatgcacctcgccaacacgtctataatgtaccaatgcaatacactcttgtgc
cttataataaatggagctaaaccagagttcataaacaagttcaacgagaatgttttgcatatagcgattgaaaatgttaactatggagtcatcact
gaattgagaggaacattatccagcgaacaaattgaaaaaatggtcaatgtaagaagaatgatggataatacaacacctttaatgatcgccttg
gcgagggagaatattgtactcgctcagctttttgacggtctttacaagcccaaaataaaggtccgtttcggttcttcaaagaggctaaggatac
cagagtttgtcctcttaaagggcctaaaggaatcagttgcatatttggaaacgaggaatatatcctacgatattaacatcataaaggatgcagta
atggacaacagtctttttgaagaggagtacgaaatagcagcagcaggactgcgaggcaataactgcgaccctgaagcagacgagaagac
tatgaacacgtggaactttttcaccaaaaattcaaccaaatgggcaagctctattttccaaaagaataggcagaaatttgtaaagattgtggatg
gtatgaataggacatatgaagactctgaatgtgcaatatgcttggatagtctggacggggatcttccttcagggagaacaacgtgcggtcatt
gcttccacaacgtctgttggttatccttgataaggatgagcgggccgaataatggcagccgcgcaagaggaggaggaataaaatgcccgtc
ctgcagacaagtcacctgcctcggaaaaagactaggggttgccgactatgatattgaaacagaggaagaacgtgacacgaaaaatgtcgt
gccttcggtagaagaaggaagaagggaatggaggaagattggtgttgacagatatgaatttcttgtaggtggagtgtggacaaatgaaataa
aactataaaatgaactagaatattggtattatttttacaccttacacatgttccagaaacgtctgttccagaaatgtgttttgagatttctggacaag
ccatttctggagtaatctgacattgggtcgacccagcggtccaccctcggaacttgacattgggtcgaccgagcggtccaccctctaaactc
gagcgacccagaaaaatttttaaaaaatttttgagatggagaatgagtaaaatctcttccctgaaagggaggtctgaagaggctatgacggtt
ctaggtgcccgccctctccagaaacgtctgttccagaaatgtgttttgagatttctggacactccatttctggagtaatctgacattgggtcgac
ccagcggtccaccctcggaacttgacatcaggccgacccagcggtccaccctctaaactcgagtgacccagaaaaatttttaaaaattttttg
agatggagaatgagtaaaatctcttccctgaaagggagatctgaagaggctataccggttctaggtgcccaccctctccagaaacgtctgttc
cagaaacttactaaagttaaagtacgtttctggacaagccatttctggagtaatctgacattgggtcgacccagcggtccaccctcggaacttg
acatcaggccgaccgagcggtccaccctctaaactcgagcgacacagaaaaatttttaaaaaatttttgagatgtagaacgagtaaaacact
ctagtggattagggtgttaacagcctaccctttctggtcgcagctagacccagaaacatctgttccagaaacttaataaagttaaagtacgtttc

tggacaagccatttctggagtaatctgacattgggtcgacccagcggtccaccctcggaacttgacatcaggccgacccagaggtccaccc
tctaaactcgagcgacacagaaaaatttttaaaaaatttttgagatgtagaacgagtaaaactctctagtggattagggtgttaacagcctaccc
tttctgggcgcagctatacccagaaacatctgttccagaaacttactaaagttaaagtacatttctggagatgtgcgcacgtatgcatgcatgca
tacgtacctatctgtatttttttaaaagtaggggaaaatgctatttaataaaacaaccaccattctccgctcactatacagctgaccttcgttgagta
accatgtctagcggaaaagtaacctacgaaatcgttgaaggggggattgttgaacaacaagtaccttctagatggaggtgcagcaatctgtct
gcagtctaattgtgttgcaagaaaacgtcacgccggttccctccacgataacctcttcaagatgctaggatttggcgacccctataaacagag
acggggaaaaacaaacagcaaaaatctggccataattgaagatagacctcaactcgggtcagtatcagttgtccaacacccgacagaacc
agaaaggttttgctccatgacattcttatttgctcagtacaatatgggtaatggaagaaaatgttacttccctaacgacaaagagtatgttgagag
ctgcaagaagcacgaaagggtccacaaatcttccacagaaatgaaaagattgcgcttgtattactttaacaagtgtcttcacgcgatcgccaa
atcacctgcaatgaagaagtacaacaagataatcttccctgccagaattgggtgcgcggcagctggaggagattgggagaagtaccatgct
tctattcgagatttctccacaatcattgataaggaagtgataatagtgtctcaaaggatgtaattaaaaaataaaaccgtcgctggcaacccccg
ccaccaccaccatcgaggcgggtataaataaggggcgctggcacatgggtggcacactcgcatcatgtcttccaaccgattcagtcagctg
aggggcaacgaggagatggttggggactattcaagatggacaactgtcaagaacaggaggaacagacagcaacagtattcccatagtttc
cgtccccaacaacaacaacaacatcaaaaaagaacatcaaccaattctcctcctgctccacctcctccattccccatcattagttggggagcc
ctcggcagctactcaatgtatcgactggatgaccagtgcagaaattgcgatgaaactggctattacaatttccactcttatgatagaaagaggg
aaagagttcgctcattaaacaacactccaagtgaaggcatgtggcggcgcacaagtagatcttccccccttccttaataagaagaaggacgtt
gacgaagctccacctcctcaatcaaaccaacacatgtaccccctcaacaagtacagtttccgtgaatatactccttcatcaaagcttgtgaatt
ggcgagacccttcacaagaaaaacaggacaagatcttacaagaggaagaagctcgcgcccctacacccactccccaagaaaaggaacc
agaagtagaaactaaagatgatgttgtcatcgaggaagaaactgcaccagaaccagaaccagaaccagccccagttccagacccagatat
tcccgcaataactgcaactactactactactacagttgcaacacgtcacgacgattcttctacagtatttctcagaaatgttattctgagtatcgtg
ttttggtttctgggtgtttattctgcattatttgcaaaatgtattagatctaagaaggaataaataaaatggtatatgaaatttaaatctttattgtttcttt
ccaattactccattgaaattgtcctccttgtgagtgcgccttctggcaacactcccatgtccacaagtgctaaaggttgtatagttccatccctctt
tggtgggaatggtcctctatctgattgacaaacaaaggggaatctatccttcaaatcaccctctgaaacagaaacagtttttctcttgtataata
atatctgctcattaatccttcatatccaaaatggtgatggtgtgcaaatttccttagattaggtccccctacgcttctaaagtagagggttgaaaga
tcatcaaatggttctgggcctttaactcttaccccctccaatgttgtttttgttccactaagatctgaaggtctcatgaaatctgcataggttatttcc
acacctcttgtacaagcactctgagcagctgtaggtgcgcaagattcgaatcctccatggtagggtgtcaatacaggaagtacatggactctt
ctcatgactctagggactcttttactcgcttcatagaaaccgaccatgtctttttgtcctccctctacatactttgatgatgatgatgttctaacgcg
gagattttctgtagcctttccttgcttattgattgtgaaaacttccgcatcttcttccaaatccacataatctgaaatagataatctcctcttctttgtac
taccggcagtgcttcctcctaacatttccatattgctgttaactacttccaccacaacttcactcccatcagttggttcaatgcttgcgccttcattta
ttgtcgcagaatcggccgatattgttggcgaatctatgtctgtaaagcaaggcgttgcatacaggctcactatatccttgacaccctcaacgta
gggggaaaatgttcccacgttaaatgtagaggcagtaactccatgagaagactcgaaaaagtccactatccttctagcctttgcaccctcttg
acccgtttttgctgcatcccttataatgtggaaaatttcagcctccgcaggaggcacgcccggcaagcgttcataatcctccattaaaatgtcat
tagatacaatatctgtcataccaacatcccctaacaattctgcgttcctatcgaacgtacccttcctttcgtccaaaacagcagcaggtacctttt
cttgtcttgtggaataaggagtgtaagccgtactcctacgcatggcacaaaatttattaggggccaacataacattggaagatatatcccatata
ttttctgttgcagcttttaggtttatctttcttccattagttttagtttctccgatagtatttagtattctagcgtctccggattgagctctcaggagtgct
gcagagagatgaacatcgtttttagggagggtagtcatgaggggaaaacgtgaatgcctatttcctctgctaggtgttgccgctatagggtca
gttttgttcctaccccctccattcactttgggacagaataaaccgttggtgcaattacagttctccttttctcccaattccccgattttatcgaatatg
tttttactattaggatcttgcatctcaaaaatactcttttctccaatattggcaggaagtagccgtgtttgttcctctacataactctttatagaaaattt
agataacgaagaatgtgattgcgggcccaggggtttggggaccatcattgtagtgtactgagagagggcacccgagcacgggtatccgaa
agattggccagtttcttccctgacacaaggcggggagtacacgccgttcaagtagtggccggccgtgtgggggctcagaccccaatctgct
ctgaaccgattagggatcaattctggcctaataaacgtttcagaggaaagagtcctaaccctgagtagcgactcaaagagagccataggag
gagggatgaaattgagggctccttcagtttccttgacttcacattgttgccataattcatccccttcatggaattttagagggactattccatgcat
ggacccgaccgtgccgttcgagactgattcttggacactcttttgaatgagaaaactggaaggtatgggtaaaggaaatccccctaaaactc

ccaaaaatgatgtagcttgagcagccgcagacactcccgcaggaggagcgagaggcttagaaaagggtcgtgctggagcgtacgaattg
aaggaggttttaatgtctgttgtcgaatttagaatagctctaatgtctccctttgttactcgatcataagtgttgttgatggctgtcaagttgtcggcc
gaaccagggtctcctctcgcgttgaccggcgatctgcgttcgaaccaggtgtgttcggggagagaagatgcaggaatttccataggtcttac
aagaggtgaagatgcaccattcattaaaactccatctccagtgaggattgctcctgcaaaaatatcgggctttttgataacgtttttgcgaccct
ttcggcctcgcgcatacccgtaccttctattccattatttagtcctcttcccgagaaaaatgtcccttcattagaagctatgcgccacgcagttgg
agggtgagcaaacatttcatatgcttctttttcccccactttcttaaacactgcagatgaaccagccttgtaatgatcaattattttgggagaatttg
tagacaatattttctcactgaatcttgcaaaggtggaacttttcttagcaccctgcgctgtaacaggttgcagaataatacctccaatttctttgaat
ttggcattgtttggttcattgtatattgatcttgtattgatgggcaggccagacctctcttttccttcgccgttcaagttggggaaaaaatagtccag
aagtaaatttgtaacgtcgtcaaaactattactagaatttgtaaccatcatagaatttaatgtactgttcgcaaaagattggtctctttcttgtgccgt
caaaagagagtgcttgaaaattgtctcgatgggagtgtgggccattcgcttagcggtaggtctatagtgtaccaagggcaatatcttatttact
gttttatcatttttagcagaaggtttttcctttattactctatggagtgaaggaacgacgtatgctgttgcgcatatgccagagtcgtgagattttctc
agtttctgtaaggggttatctttgtacgaatcgtaccctgccttgtcagaatctttaaggaaggatgtttctatggtcgtatcgttgtgcgcatccct
tctgtcagtctttactcctgaaggatcgcctgctctcatgtgaaccgtctcggggaagattttaccctgggacgctattatgcccacagcatgat
ctatgctatcgattcctaaaccgggaatattgctatggggtaaaggcacgaccctgagcgatgcgtttggatgttcagtaaaatttccccctcc
agctccaaccagcccattatcgtttagtgcatctccgttccagatgtgagcggccacacttcctgttccactgtagggcactacacctcctaca
gattcactgactgtgagttttccttgttcgtcagaatcttcaattcttgagtgccaattgccggcggcgtccatggcagattcgcgcaataatcct
cccgtaatcagatggcctccgttaaacgctgtaatttcaggtgctcttccaaaaagagaagcggcataagaatagccgagagccgacatcct
cgtatcgtttcctggtctagggaggcacgcgtccttagtactaccacctaaagtttcgtacctttctgttatttgcagcaataattttgcaccctcta
caaagggttgagcataatctgtttttgccgcatcgggaagatctataccggccacttcttccagtctgatttgtgtactttcatcacaaagacagc
cctttgttccagtatctcctacaaccccatacctgaacatgacagggcagcctagaccttgttccttttcctgtataccaccttctttcttctcacatt
ttggggcgttggtggcagggtctctctgctgggagtagccaggttcgcacaaacaatccatagtagaagtcatactgcttatgtcgaaatcga
tatcgtccacatcatcaataattgagaatgttgtagggtgtataggtcgccccactccttttcctccacacagtacaatatctgtgcacccaaga
atgggctctaatcttggatctttattgagaagggaaggatcctttcgcaccacagacgaatctacacaccctctttttttggtgatataagtcgttc
cctttttttcctcctccagttatgactcctccagattcgtagtgcaaataagcctcctcgacagtagtaaaagggttgacgtttctgtaggctgcatc
tgtagcaacatcatcgtcgtgcttgtcgaacgcagaagaggctgaaaaaatagtatcggcgagagggacacaaactccgtcatattcatcat
catcttctataccgtattcagttttcaattctctactggatttgacgcaagttcctcccccgagagaacgacagctaatgcattcaaaatcgttgcc
tcctacgcgacatttttaggagttttaaaacagtcaaatgtgttataatgggaactaatttcccctggtaatagagattctacaaaacctttcccta
tgatgtctgacttgactccagccataccgaagggggcgtccgtatcgacatataaggggtcgaaagggttgggaataattcccccattttcttt
taatcctttgacgcgtaaagagtttcgagacatggctcctgtatctacaaaagagctcgcttaaccacagttgcactcacccgggtacaataag
acaaaaattataaattgggaagaccgatacagtctttcatgacaaaataccccgagaacaaaagattgttgtctaggaacaaagaaacattaa
aaatggttgcccgaagctccaagaccaaatcccgccgtggaagcaagaagaggtccaccactgctggacgcatctccaagcggaggag
cccatcaatgaagaagcgtgcaggaaagaagagctccactgtccgtcgccgttcctcaaagagcggaaagaagtctggagcccgcaagt
caaggcgttaattcttccctgtacaacaactatgttatttaattgattttttttcttctgaataattggaaataataaaacatccattgaaacttatgca
gtatttttattcaatttttaaccaactataaatccacatgtggtataaaagtttaagggtacagatatattataagatgatgtaacagatgcacggt
caataacaacagggcctgaagaagtgtctatgaatttcttgtagaattcttccctgtctgtacttgacgctacattggcaaataccagagggtct
actagtacacctgagctagtgccgggttctattttagacatgacattttgagggattttaacacataatacaatatcatcatggtcgaatgtcgtcc
tgttccagagacggcgatgtagtgctgaatttctatccactaaaactagaccatcgtcatttttcaaatgcttctttccttcatttctagctacaatttt
catgtagaatcttactctttctgcatacgacaccaccttgtcttgattcccaactatagcaaatagtaacgaggaagtaaattcattgtttgcacca
ttattattgtagaacccgtttgcactaccaattcctccacatctagggtctagacgtaccagaaattgtccatcggctaaccgagtaacagggtc
gccgctaggccatgtggctagccgagtctgtagaggggcaactggttgtctcaattgagaataagaacccactgcgcgtaggcggccagt
gtttctccagttcgtgtatctgtctgcaggttcgtcagggtatttatttgctccaaaaacaacaataccccattttcctaaaatagccgcaagtttct
tgtcttctttatttaccagaagattttcctttttggttccagactcgtctaccacgtatgtagtatttgaagtgtgcacaatttggtctaatgtttcttcag
aaacgccgtctcccatgtacattttagcgtacactttctccccctgtgtatacgcttcactaggtgctggcaataactgtcgatgacgttcttgttc

cttatcgatttgttccctcaaccagttttctattgaaaattcagtttgagtatctgtattgacaatgtttgtagtatcattcgttcttatggaagaaatga
aaatgtgccgagggtactttatttctcccccgtaatcagcgaatgcccattcaggtatgcttgtaggtaggggacaagactagtctgtttcatg
aaagggtcccttatccctctattagaaatgggaggaagtgtagtgcgacagtagactatgcccttgttggttatgtcggtggcgggtgtgtccc
ttacaggtgccgtaaaggttgagggatcattccatatagccaattcgttagatggtacagacatgtctgaaaactgtccttcaaaatgcccgtc
catcgtaaagggcatctgtatgcctttatcgtcacctccaaatacttccatctttacactgaaaaccctatcgtctcgtgcaccgtcaataaatata
gggcggctggttattctcgaagaaagaacgttttcttttgcaagtatagaaccagacacgggatcggagactgtgatgtggtagcttttatcttc
aaggacaacatcatcatcatcatcattgacgataacatcagatacggacaagacagtagttttgttccttgacgtgatactgtgaatgggttgg
aatgggttatcactcgctttcactaggccgttatgcgaatctaaaacagggaaaacagcgtacagtttagtaggttctgcgggtgccttaaatg
taggctgttcacctcccagtaatgcagtttgaacgggggtaataatatctccacctcgtacttcagtcaatggaccaactgaaacgtttaaagc
acgaggcagtagagatgttgtgcgttcaccggcatattctgtggcgggggaagaaagcacagccgttattcgaggttctctattatcattatca
gttttcaagatgatacatggtcttagttggtaggagaagaaatctggcccgttcgtgggtccaggccaatctggtgcagttgcagcatctactc
taaagtttacgctagaaccgaagcacccgactctcgaaggtgtattatgtacaacaactttgttcgtgttagcattcacatcagtacttataaccg
agaacaatttatcagattgagtaacttttgatgctggaaatacaatattgtacccagaagatgtggttaaaacgtcagacgacgttttgaagccc
acaatccagcccaacattttccattgcgcttctatttcactttcaggtacagacacacatataaaggtccgagctgggttagatggatttctaagg
tgggttatctctgtggcgtttagaatcttgtaaccctcggattcatttccctccatattctttaacgtttcattgtatagtactttgtttaatactggtgtg
ttaacagtgtttgttttcgagtacacgaacgacactgaaggtttgttaatatctgaaagttgattttcgcgaggtagtttagctgctaaattggacg
ctatttcagcttgtcgggaaagacggggaaaattagcagggtaatctgaaacatatttttcattcaatgtatcaggagtgtattccacattatcat
agtttattgctttaggcatatcaagcctcatgagggctacatttctacaagcaaaaacttccgttatttttggtatgggtgaagatattgttacctga
gaatcgaataacacggtcttgtttttatcctctgttacaatggaaattttcagtctattcccagtgttgaaaatacttccctttgttgatagtgttatac
cgtcagctagagaatttacttccacgtgtttgaatacggcttgttgtttattgccattgttgcgagagaacgcattcgggtcattggtgataggag
gccaacctgctgggacaaattcagggaaagtttcgaatagagacggaccactatcgctgctgccacccccgtcttctccttcatatttttcatc
cactaaatcaacttttatctctgccttaaagggaagagggacattatctggtaatattccgtcaagaatatgtatcctcacgatggtatcttttcca
cctttttctaataattcttctgtacggatattaatcattcgaattggcagcaatgatgatgatgatgaaccatctctatcaggagccgattgaggag
aaatatcatcaccaatatcataaccgatctgccgtgcagtgcgtttcattatactgtgtaacaattttggatcctgcacatcagtgacacgttttaa
ttcctcagaggatatcctttcaatatttttagcgagtttttttgacgccggttctaacccattcttcaagttccttttctttttcgggtcttttctaacagc
agcggaatcatcgatagccgctgctatattcccgaatacagctttttctattttcgcaggttttcttatgtgtatcacaccagatctttcatcgggcg
taaaattgttctggacaattttcataccaatataggaaataatagaaataaacacaacgagaagcgtgaattcaaaggcaccccaccacgagc
caaaaatacccgtgactgctgtttctccaaacattgttattgctgttcttaacggcagggttggttgttcggcaaatcaaggaaacaaaaaagtt
acccgggcatggcagggaatagaacccagttcgtatcgtccctgattgctaaatgtatatcagacgtggaacaaggcatggagtgttgtggc
agacaagcacaggatgcattaatgactcgcctagccaacttaaaattgggcgattctcttaaagaaactgatgttaatttggaatacttgagata
cgcgtctacgcccctccttggggaattaaactacgacaaacaacaatatgcggcaacagttgacatcaacctaatggctcatttctcctacgct
gctttgggtatagaaagtatactgaattctatacggagagttgtagtggctaatcatcaacgtagaaataatggaaaaaaaccttctgaaccaat
ctcacgccctcacccgctgggaggggtagaacctcctctatcgtcagagttggcaaatgcaataagggacaagttcatcagcatgggggc
gttggacagattgaattcagcaatagtgacagcggccttgggggctattgccagtgaacgtgaactattcttacgtgaaaatgctgtaaactac
atgtacgatgtagaatttgcagaaagagatgctgctactacagatacagggaatgtagtctatctttccaccaaaatggacgaagatgaagat
gacataataaagcgttcagaaatattagataaggtatcaaaacgacccgcaaaggaaggtatagactggcgccccacccctgacaattcgtt
cccttaccaattgatttggggcgatgattctgtagatgatactgttcttatagatctcatcaccaatgcgatcgtgcctaatatttttatggcaaaatt
tatcctgttcatatgtaaccatttaagggcagttattaggagtatgagggaaattttatacggtaacatttcttcttcatccgataattattttgaggat
ggacgtaaatggtgcttctggttgaacctgtacaatagactggaatggttcatgttagtagttagatttgtaattttcctccactcaaaaaaggag
tccttttcaggagctgacaatgttaacgtgaaaagacttctggtggtagttgtggagagttttcctcccgttctcttggacactgaatgggtcaag
actaatataacgtcatggcctgttattaataacagcaataataatagtacactccctgtgacagaagacaccttaatgagactagcgataagga
cgagtagcggtgcccgacatcctattttcgacgaaattaactccttgacaacagcagtgaccaaccgtattaccttccagtctgcagaattctg
cacaaagattttgctcgggcgagctctggacgaagaagaagctggaacaaaaatgctagtaaaatcagtcaaagagacgggagaagaaa

aggataagaacaatacgttctcttcatttggtttattactgaagaacacaaaaaatgaagaattggaaataaacataggcgataacgatgatga
gactacagatgtggcttgttgggcacgtacttcctcgacatcctttatccgtaataggacatatgcgtttaaaaaaatatggggccttgaggatg
caagtgatgtagtcgagctgaagcgagagagtgacgccattacatcctttgtcaccgataagagcagtcctctcctatttccgtatgtgtccga
ctggagttgcttactattacatccctgttgtaaagcaccggccataattaaaagtgtgtggttacaaatcctgaaagatttttcccaggaaaatat
aaaaactataaatgaaaaggtacaatctctttcatctgagatttgtcagaaatcaaacgaccgttttaaaaataaaaaaattgctgccgaacacg
ttcgcagtgtaaaaaagttattaaatacgataagcaacagggagcaagaagcagcactatctacagaacactgtatttggttaacgattttgtg
gaaacaagtcgttcagaacactctcaaccttctggagaattttcccgtataaaaggggaaagtactcaacatgttcactcacttgacccgagc
cttccgcaagatgaacaacctagtcaatcgcagcttcattgacgttcaccgagtggtggccgagttatcctatccagaattcgaagaggatgt
aaagaatccggaatcatccatatatagaactccgatatccctcttccaaaacaaggatattgttacaatagttggtgattacatcctctctccgaa
gacggactcattccaagttctatacccaatcaagaaggtcatcgaacacttcccagtaatcttccactgcacccacaataatgcccctctctgg
gtacaccttctggacgaacgccatcatcgcctgctccagagcctgctgacgtacgagattgtgaatgccaagtacaggggtattgttgtcatc
ccatactacaggcgccccatcaactatcaaactgggaagagtctactgatgagcaaactggcgtccgtaaaagttctggacattttaatgaga
tgtggatcatacaaattcatctcattaatgtgcatgatcaacaagaagaacaacaccaactttcttcactgctgtgcaagtaaatggggagaag
ttggaagcaagatgatgctccacattgctgaaatgttctttgccaaccctactactagccaacacctatccgacgctagtagtttccctgatgct
gcagcagaggacgacaaggggaaaacacctgcccatctagcaatccaagaagataatgctgatgcactcctgttcctgatctccctctacg
gcgcaccctggtttcaagataacaactcgtacatgaaatctgcccttgaactcaagtctaacaagtgtgtcaaggtactatcctttgcagctga
caagtacgagattttacccaacattaacaacaatcaactagaaccagataccatgtgtggagtgtgtgcaacatctgtggaagaagatgaaa
atgaagggaaaacaacaagtctttcctggtaccagatgaattgcaagcattacatccattgcgaatgcctgatgggaatgtgtgctgctgctg
gcaatgtacaatgccccatgtgccgtgaggatgtgggcgacgaagtactggaaagatgccctcctacaatatttagatggttaaaactggct
gagagatctgaacacaatcgtgtactttttgaagcaaaaaagcaagaattctataagcagatggaagcaatgaaacctcccagagttgttgtt
cctcctcgcaggacatttctcaccccagccagaagaggcgaacgagccatcagaatcgcaagagaaattgccaccaacgccatcgctga
agccacagctcaaggagatgtcaactcctacttccctgttctcattgacgggagcggagaagaatatgaagaagagggagaagaattcttc
aattctgaagaggaggcgcttgcttttggaagaccatttctggaagatgaggaagaagccagacaaatacagatgcgccagtttgctgaact
gtctagacgaggcgtttctgtcaatattattaacaatgataatcctcatcgacacatctctacagtaaatattgtgcaaccagtttatggagttgaa
aagtcacctgctgcttccttcatctacaacatgctcaagaatgacgtctttgagtctatacgctcaagagatactcgagttggaggagaaagag
tgcccgtcatgaacctgtccaatgacaagagggcattattccacgcagcttcttccatgctttgtgactttgccacagaaacaaactctcaaatt
gttggattggactttcaagcagtgtatgatccccatcacatatccaactatatcgagacgtttggtagtcctcttcacgcctacccaggagccgt
cacttttctggacggggcccaggactattatgcagagagtatcagatacgacaatgatattgtctcattctcagaaatggcaagtgagttgcac
atcaccgaagcattagatgtctttgagggtagtttattatccccactgttcaagaaaatcaggactggaaaatcttactctaactggaacgacca
tttgaggcgtagaaattatgctcgagatattgctgaggaatttgttagggtatgtgaaaactctctagcttcacgcgaacacccccctgttcatgt
acatccctttagagatggagcaatccccattctcattgaatatatagtagatttcatccaccactgcatcacctggtctatgcaagttaatgcact
ccattgtatgagaaagtacattgaacacgagaatacaaatgtgcacctgttaaacttgcgtcctactgatgaaagggtggaagttttaagggta
tctcaactcagatggagccgcttgttcaatgaacaatacaacactagaatgtccctcagcaccaaaagattgagcctcatgaagatcttcaac
catgatttgggtgtgtctaaatttggtgtatacaaactcctagatattattgaaatgtactgttttactttaatctaaacaataaaaataatgtaaaga
attccatgtttttcatccctcctcctctgtggtgtgtatatataaggcgggtagcctcctccacaggcacacttgtccagtagccgttgtcgagtg
aacatcatggcagcagacctcctagagttggctatccaggaaacaatccagtctgaattggaagaaattgccgatactgaattcctcaattatc
ttccccataaaactggcatctgcgaagaagctgcagctaatggacggccatatcttcctacactagaaatgaggaacgaagttgaccatttct
ggtcccaagataacaggaagctgaaactcttggggcatttttgtggcaacttgtatgtggaggcatttatagctggttctatagatgctgaaac
gtgcgtgggctttttgaggtcgcaagcaactggactaggataccctctattgaaaaaactagccctgattgcccgtgaggataaatcaaatac
aactaattacaacttgtacattgatagaaattcaatgatgaaacaagtttttagtgctgaaattgataagcgtccttcatcaatacagaacacttct
cacacaaaatcttcccccgtgtacttgaaactgatcgacagaagaaccgagtgtcttgccctggattggctggacgcatccaagaggacgg
ccaaggaaattggagcagccagaaaggtttgtttcctccagaacctgattgttgccatattaatacctgcatacacagaaacgtttgttcttgat
acaggcaacgaactagaacagcaagtattggatgatgcatattttaatgcggagaataaagataaagtggatgaaatgtgtgtagtggccat

attgagtactttgcacaatttatttgttaggaaaagtcttccccatcatttgtacaatgcacctttccgtcttcctccctttggacaacaccctatcat
caacattgaaaattcctcattctttaatgaagatacgacacctattctagcgtcaatttctataccatcaagtatggtaataaaacaccatacgcg
aaaaaatagtagatggagatgtcccaataacttgatgactgcagcagaacggtctatatttttacgtggagttttaacagtcagtggagattacg
gatggttttctgtaattgtaggatctactataatgcctagtgtacttttctatggcgaccgaaaacatctaataaatacagtcaaatctaataactttt
ctgccataacctgttcttattggaacaagtatatggattgcagatcttatgggtttgagattatagacacacctgaaaataactgtggttttcgtata
agggctgcaattgattgctcgaacacagatttccattcaccggtaacgcgagtcaacaagaagaaaacgagcattattaatgcggtaaagaa
ccctttttttattagacacacagaacctaagtggtacaacaaaaatgccatgtgtggtgaagtattggaaaatgttggcgtgaccctcgaacaa
cacgtccgtgttagtgatgagtatatggacagatttggtagtctattacttggacgagaaaagaaatggacgtgtaattatctagatagaataaa
gtctctagaaactatttctaacaatctcaagggtaaaattgacaccatgtgtaaaattctagaaaccaagtataactacaaatcttcttccttatact
ataagcaaataactgctacgtctgatgatcctataaagatgaagattatcgcctctataaacaagaggaggtatttgtgtaatattctagaatttg
ctataatttcatcagagaaaaaggatgaggtggaggaagatcacactaaaaccggcaatggtggttgtgcgttttcaaagtataagaaaaaa
caactcgaacctaaacaacatttaattgttaaagtgaataaatacattgaggctttttcgttaattaagatgctgagaaatgattgcgaacgcaac
aagtgtaggtttaaagaggctgaaattagagagtgcgccaacgaactggtacgagaactgtatagggcttcggccagaagttatgtccacg
atctggtattgaagcgaactaatgtccacttgacatggcaacgcccttacgatgaaaacgctaacactatcatgtctttaatacctaaatgtaag
ctacacacagtattgtacgataaagattcgcgcgatgttaagttgttgaattttctgagaacgagggatggaaattataacccaataagacactc
tatgctggagttagtatacggagaggagtacgctaaagatgtcagtactgttacctgttttgaatggttaaaatggtgctcgaaaaaaggtgtg
attaaatatgaagactttttggatcgttacgagaaaacgggggaggaagataaagacgaaagggaattctttagactaaaaaaatgtagtaga
gatcacactaaggatataaaaaaaatagaaaatgtactaaattctgatacactttattcttattctctcgataaaaatgtgcaaacccacgcatctt
ctagtacagttgtaaaaaatgacactgacggaaaaacgtctatggtgggctgggattatattttttcaatcggtaaaggagaaaaaacaacca
aaaaacgaaaactggaaacgatagatatatcgagtagtgacgacgacgatgaagaagaagaggaagaggaagaagaagaggaagaag
atgaaggaaaaagaatgaaaatgaataactgcagcagcagcatcaagaacaagagcaagaacaagaatgggagaatgtgttgcacagat
attctcaatgttgtagaaccttctctacctaatactttatcgttcaattgtgtaaaaagtatggatgtgttgaatttgttatgatgttaaaaataataaa
aatattgtacattttattacatttgtttgttttgacacctgagcggtttgggcctgttgacgtgcctaactttttgtcaccctcatgaatacaatttgtaa
aggtgctgaaatgtacttgtttttttatccaaatttcctgtactgaagaatattgaaagaagacttcttgaagaggaccgataaaaaaatggccac
cttccagactgacgccgatttcttgctggtgggggatgatactagtagatatgaagaagtgatgaagacttttgatactgttgaggcagtcagg
aagagtgatctagatgaccgtgtttacatggtgtgcctaaagcagggatctacttttgtcctcaatggaggcatcgaagaattgcgtcttttgac
tggagattcaacgctggagattcaacccatgattgtgccaacaacagaataaaataaagacggtgacgggagactaatatctttcttagtttcc
cgtcacggtgaaaatgttggttatttcttccctatgtttaaaaatttgtcttggttaaaaaaataaaacgaaaactgtcaatatattgttttattgatat
acaatatccctttttacacagaaatggcatgtctctttacacgctgttgttcttgagtgttgtttccatcttcttcttcttctacgtcgctatcaaattcat
tgtcgttttcccaggtcctttttactccccttttgctcctgttcactggtagactgttctccattttctcgcttcttttgttcctccctcttctttaccttctca
aaccacttttcggtcttgtttctcgtacgagcttcctttgtggtcaaatgtctttccttgttgtacctagtgacaatttttgatgtgttctccatcatgga
atggagcttcctttgttgttcaatttcaaattgggtcatattttcttcatcaagttcttgaggcgaagagatgtctatcctctgcccatcttgatagac
ggtgccatcattcataaagattcggatatcagaacctgtcactttttgtacaatacccttcaaatcttcatcacttggaggggtctggaaaactat
acttgatcggttacattttaataccttgaacttgttttggtgtttgatactgtcgttgcaagcgataggtgcgtcacagccagtcacggccatgatg
agtctgggcctcatacctgcagtatcccttttcttgtttatgggggagaaaatgtacctcaaaccattttctgttgcagtttcaaaggaaccactttt
gtctgtcctgcacaacaagtagaattctgtagaggatatcgagaagacgagacaagagccgctgtcgtatctgaaggtcgcattggaagatt
tcttgtgtacatttctgagagattctttttgttcctcagagactttgaaaagttgttcagtttcgatatacgagtttttgtactcctctttctttactacttgt
gtttctagttccttttcaacttttgtggtctttactacttgtgtttctagttccttttcaacttttgtggtcttttcgacctggttttctagttcctcttttatttgt
aaatttttcctagccaactcttcgcgtatctggcgttgcttttcatcactgtataagataacagcgtaggctagtgtcgaaaggtacatttcaatgc
ttgcagcagaagggtgtgcgctccttgcaaaaacgcccgccaccataaacaagggaacaatcaagcaatgtgaagaatggctcttgaaaat
aggattggctatatattctcctttgtaatgggtacggtgacaaatttcaaccaatgggaccatcatatccttgaaaccatgtttctgcatagttctct
ccattgactttaaagcgtgcgtgacgttcattgtactgaaacctaaaccgattaagaaatcggaacttattatcatacaatcgtgttttttttctact
gcttcagaaagactattgctcatgtacctgtataattgactgaatttaaagtccttgtcttttcgttttaatttttccttgactattgtattgattttacgca

cgtcttcttcttgcgttgcaagccattcctcgtcagtaacgtgcccagcatcatctttgaataaatgtccaggttcaggaggggcacaacttatc
cgttcaattcttaaagagtcatcttctaccaattcttgtaatctttttaaccctcttgcgaatttgagttgtttatctttcttcacatgtacaacaaatgtg
tcttcggacaattcgctttgtcttatagattcgatagcttcataaacacccatgacttgctcatatttacttgttttcttcccgatcatcaggaatgac
gatcttagttttaactccttttctgatacactttcattactaacagggttattgtttttctctccaatgtattccatatttatcttcttcaaaacaacaacgg
aaaatacagttttcttttaagaaatataggttacaatactgtggattgataattttactcacagctttgttcaaaacttaactacgtggttcggcccat
tttgcaccagatctgttttaaaggaactgggtaaaatacacatgaaaacactccttttgaacaaattcagatttatatgtatttgtcgatttttattaat
attaataaagtatatttttaaaaattaggatggggacagatggtgtagattgatacagtgaccgtccctgttattttaccttaatttttttccctataat
acaaataattggtatcagtttaggaaaacatatttttaaaatgaaggttatcatatcaactaaacacttttatttacccccttttcacctcagaaatat
cctcacataatcgcatgcaagacttgcgcatttcttccctgccgtacaaatcaatagcaaacgctaacgccaacagatacgtctccacttctag
agatgggggttgaacattacgggaaaacatgtaccctagatacaccagtggaataactaggcaatgggaatgataactcttgaaaatggggt
ttacaacaatatcacccttgtatttttgagatggcacacttcggccacaggcacccatgtatatttcagaccacttttccgcatatttccctctatc
gctattaatttttccatgacatcatcagcactaaaaccgagaccaatcaaaaaatctgtgctaacaacaaaacgttccttttccccgaaaggtatt
tgaggaagactttttgttatatacctgtaagttggactgaaataaaaccctctaaacaacatccgtgtcttttcctttacgagcttatttacttcctgt
acgtctttgtcttgagaagcaagccactcgtcatcagtagttgtagacgtgtttttaagtataacgtgtcctttctcaggcttaatagaggataata
atgattctatcttgagggaattgtcccctgtgagctctcttaattctttcagtccccccttcaaaagtaatgtgtttattttcctttaattgtacaatgaaa
acgccgtctctaaattcactctttcggatagtttcaactgcttcaaatgtactcattacttcttcatacttttcatattgtttttctccaatcattagaaaa
gaagagcgtagctgtaactcgtcttcttcaggtgtctcatccaggagatttatcagtttttgttcccctatatactccatggtttatttagtcagttga
tttatttttctctttgacacagtttttatggtactctgggattactttactcaaaaatttactcacatgttgttaaaagtttgagtgtattcttttgtggtaaa
ctaacaatgaagtacatttcttagtaaatgaggtaggtactgattcgtgatggataacgatggcgtcccatgtctagtgatgtgagtcacataat
aaaaagggaaataaaaataagaactagtaacatatttttttattttatccttctcaatatgttgtttgatttctttctaaaaatggctgctgcaaaaatg
gacgctatccttgcggatattaatgggaatgatacagatttatccaagctaatcacagacgtgattcaaaagagagccaaggctgtcatggat
agaaatagggctaaaatggacatgaatagaagagtagatgaggctattcaggaagccgtagcggccaagaaacaaaaagcattagtggta
tttgataaactcgtggaagaaactgacagcggacaaagtgtccctccaacattatcgggatccgattacgacgcgtgggtagacagagcca
tgccttcgcatattgaacttgtagagagtgttgagggagattctttgtatgataaactccctccttttaacgtacaagacatagacgaccaaatcg
gtgatgagatagatacaccaatatcttaccttgccatggtagtggtaaaagtcgactgtgaaactggggatatcgaagaagagtacaatcttg
ctcctacctttggtgtgacacaaaataataaaatatacagagatgaaagagaccagatttttacaaaggctgataaatctgtgcgtatttttaaac
ttgctaaattggatagtatatcaggtaaaagtagacaactgacgtatgcggtaaaaaataacaatgaatatacagagtttgtctgtagcgtctttg
cagagtttgaatctgacagtgacactactaaatcaggtataggtattcgtgaatatgacaaacctaaaaatgaattcgaatatgaggagcgag
agatctttacatttttttattccaatacaacccgcagggacgaaattattgttatactttttggtggacgtcaggtcccgtattatataggactataag
gcacaacactctgtttgaagagaaaaaatgttcgtcataagcatagctacgtcattggtgttattcttcttccttcttttcgtctcaataacaattttta
gatggtgcaaaaacaatcgactctcaacctttttaggaaaaggaggaagaggaagagatatcgtaccagtgagagtggtgacggcatagac
ggaggaactggaacaacaaacggaggaggaggaggtggaggagaaggaggtggtggtggaacaaatggaaatggaactggaacaac
aaacggaggaggaggtggaggagaaggaggtggtggtggaacaaatggaaatggttctggaacaacaaacggaggaggaggtggag
gagaaggaggtggtggtggaacaaatggaggaggaaatggaaatggaggaggaaatggaaatggaaatggaaatggaggggatacag
acacagacgattttgagcctacgccagcccttctaaaagaacgtctacttaattccatatcttcaaaacctaaagaatactatgaagcattcgtat
ctgcagaggtagaaactgccttacaattatctcgagatgattctacccaaactataataattgatgacgaccaattagaattggacgcctcaga
cactctacaaggaaaaccaagggattatttgttcaagctcgcaggtgtttcatcggcctttttagaaggtaccactatccgcaaagcggaaga
ccgtgcccgtaatataaatgaggaagaaattgcacaaacaatattaagtcaactaagagaaaaacacatcaacgatgaatacgatggaaaat
atgccacacccgaggaaagagctgatttttccaatagtcttaatcttgttactaaatacacaaaccatgaagtgggtcttttagtaggagaaact
attgaaaaggctttccctcatgagattgagtttgagagatgcataattctagtagaggattttaatagtggaactattacttcaaacactatgcagt
acaggtccaacgcttacaaaatcagagtagtagaaggatcaacaacagatccaggggaagttgtccctgatgattgtttggtttttgccgtagt
agtaaataaggaacaacattctctagaaatatctgcaactaacagatgccaagacatatgctttgtaattattcctcgtttatccgcaataggaaa
aaatgctaccatggtaataaggaaaggtgatgaaattaaacaagaaacctatctgtttgtggccaataagaatgacactactcatttttcaatca

tcacagacaaggatgaatctgttggaatagaattaaacatgctcattttctcagaaagaattctccccactttgagtgaccctgcaaccgttcca
agacctttgactgacgccaacgtactgtcagcctacggaaagcgcctaggtgttggtgcctttacagacaaaaatttattgtccagccaataa
aaaaaatataaaaaaagattgattgttttattttcaaaaaaaaataaaacaccatgggagataagcaaaaggtggaacagcttttaagggaatta
aaggcagaggctaatgatgattggctatcggctaatgttgatccaattgttgagagatttgtcacaactaaatctgatgagacagcccaagttg
ttaaacaggccgttgatgaaaaatacgatgaattattagaagataaggttgaagaaatgagaccggatataatcaatgaagcatccgaaacat
atgataaacttgctgctgatatgataagagaggtagacactagtagtgttattgctcctgcaatagctggcacagtggcaagaactatcaataa
tttaagagataaaaggaaagaatatgaaaagaggctatggacattagcctacaaaccatggagaagatatgtacaagcaattacagtgatgg
aatttcgtttatcatataaagacctgactgtccatgccaattccgatacatatttaacattcccttttttaagaataaaaaagatcgcctatattaata
acgaccgcgccagccccgtgaactgctccttgtcagtgtcttatcccaacaagtccgagtggggaaatgataacggagttggacgtaaggtt
gacatacacatcagaagaaatgatttacaggaaaaggatctttacctgtctgttatttgtatgttagatactgatttttcaggatatgataaagctgt
ggaagttgatgcccataaatttcattttgaagcagggaacaggactatgtttttacccaaaacctctaatctatttaatcgatcgcatatagtaaac
tctaaaatatgcactatagtgttccctcccgcttctgccagttctgcttctacaacagaactggataatgtctattacaggatcacatgcacatgct
cctagagggggggggtggctgtgtccttgtataaaaagcgaggttggcacagactatcaaacattcagccgtaaaccatggaggaagaaa
gtcaacgagtacagaggaggatcggagtcatcccagaagaggctgcctcccagatcctgaaagacaccaagttaagagtatcatatctgg
gcgtgggacattggggatactctgtttcagttataaaaagtgctctccagaaaggatgcaggaggaacgatgaggatataactgcgtggtca
atcagggaggcttatctctattaccatcttggattgaattacattgaaaacgtaaaacctgctgcaaaatcattaaatacaaatatggtaaacaga
attaaaatcatagctgtggaggataccagtcctagaagtatggtggcttctaatgagtgtgtgagaactctagaaaaatatgaaaaggggaatt
ttaggcaacccagttacttgatggatgctgccatgaggctagttcacgcctccagttctagagtgtgctcccacatgagagctttgtgttgcaa
ggaagaggacagtgataaattaggggggtatttattatgctaattttaatgaacttgaaactcagtgtgttagtgcagttaatttttctcccatagaa
agaattaaacacgtcttcagggagattgaaagtgtaaaattggggaagaaaagtgtacagttattaaatttaagaagtgtagcagcttaccatg
tgttaagatattatggggataaagtgaaagatacaaataagaaacatagtggaccattcaagcgaaaagagtttgagcagttttgggggttat
gctttaaatttgttactcagcacgtaaaaacagaccctgaattacgttgttattttaatgagttgacatatgccataaattggagaagggattttttct
gctcaaaagggttctttagagaagaaagtctattccttacatctattgtggaattaattatagccatgtgcataggtgaccgtaaacagtttgcca
agattcagaaaagggatttaaaacgtttcaataaaggagaagaagggagaaaagaagaggctgctacatttgattggatagaaggacatgtt
aaacggatgcctcaaatgcctgtatgggttctggacaaacatacgaacaaaaacacacatggagtatcttttgccctagagagtagtatggttt
ctggaggagacaagcgctggtccccaggggtgtggcttcattcttatacaaagatgcgtctagattctcccccctcccccagaagtgggccaa
tttctggatcaggcttttaatacattgaagcgagaagctgctagtcattgcgtaacgaggaatatttgtactactacaggatttataaaggcttcta
gttttactgctaatattaattctgaaccgatggaaattaaggaagaaataaagaaaagaaaaattgagattaaggatgataatactactgctact
gttactgttagtgctactactagtagttctattacttctactccacctcctacaaagaaacaaaaaacaactccaagtggaagcaataaagtaga
ctctatacaattgaataatctaccaactttaaatatggaggatttagatagagtactagaagtacacaaccaaaactctaaaaagggtgtagctg
ctacagttttaatgaaggatggaaataaagttgtgtttaaggagatgagaaaaagctttggttggggttctcatcagaattttgttcaagtactaa
aggatgaagatgtgtgtaaattggactatttgttgccatgccctgatagcggaccatacagaggattatacagatgctatttcaaaatagtaaag
gatgaaatttcgagtacagctgctaggatagaaaaagtgaaatggggagaaaatgccatgtgttatttcatctctgggtgtgtgactagacaa
gaaggaattgggaaaattataacagatgtgcgcctttcacatatgggaccaaataaacaatatgtgtatgataactataggcaattaatacaca
ttttaatttttagattgttgactggtgtgagtgataccaacacatcaaacattctcgtaggtgatggaggaaacttgttctctgtggacgagaattat
gtgggcgcaaaggatccgaggacagctctcgaaaatagaaagataaaggaactacaattgctactgaagacttcctttaaggtaaacaagg
taacaaaggaagatatagatagctgtcttccatcttggctatttgatacatcaaaaagtgataaaataatgaatggtgtatgtaatattggtaaaa
atatgggaattggtcccactactttggatattgtaaagaataattgtacttgtattttaggggtcgtgaatgacctgctttatgacaacaaataaag
gaataatactttatacaatttttcatttccttccttttttaaagtgtataaaatttttctaagacaaaagtgagtttgagggttggtcataatctagcacat
ttcatccctttttccccttcttcttcttcttcttcttcttcttatttgaaagaccctggttaggtccctgctaagggtaggctgtaacctaccatatacac
tagagaattttactcttcctccatctcaaaaacttttaaaaaatttttctgggtcactcgagtttagagggtggaccgctgggtcggcctgatgtca
agttcggagggtggaccgctgggtcgggccaatgtcagattgcaccagaaacgtctgttgctccagaaacgtacattaaatttctactgtttct
ggaacaggcgtttctggacacaccagcacccagaaagggtaggctgtaacctaccatatacactagagaattttactcttcctccatctcaa

aaacttttaaaaaatttttctgggtcactcgagtttagagggtggaccgctggctaggcctgatgtcaagttcgaagggtggaccgctggctc
gggccaatgtcagattgcaccagaaacgtctgttgctccagaaacgtacattaaatttctactgtttctggaacaggcgtttctggacacacca
gcacccagaaagggtaggctgtaacctaccatatacactagagaatttttactcttcctccatctcaaaaacttttaaaaaatttttctgggtcact
cgagtttagagggtggaccgctggctaggcctgatgtcaagttcggagggtggacctctgggtcgggccaatgtcagattgcaccagaaa
cattgcgtactactcccatgtccagaaatatctagatcctttctggttctcgcccaccaccagatatctgggaagccttaggcgagtcatgtttct
tgcaccatgtatttctaggaatttcgctgacgtcaataaactggtttgtcatcctgttttatccagtttgctgacgtcaatagaccatatttgggcat
ctcgcccacacagaaacaggatatgatatcatagatttctgggaagagggtgtttttgtgccgatctgggtgtataaaagagcgctgcggag
aggcagaaacatcagacagacttgatctgtacagctagcagcagcagtagcagcagccaagagaagatcggacgcaaaccattctcgca
gccatggcagcagcagcagtctcaggagaggggagaatctctgcagatctactcctgttggaacaactcaccccggacggagacgtgata
agatacgactctgagcagtacaccaaacctaggaagatctttggtgacaaaagtgtgatagagactattggacattttctcatccacaaccac
aaccaaggtgagagttaccaaatcgcatcttctgtgttggaaaaattccccgctctactcaattgcatatggaatggagagtcgggaggaatg
gctctatggaaggcattgtacagggctaaaaagtatagacttctcaactcacttttggtccacaagataaaaaattggccttcggttgccgtgat
ccctatctacgggtcagtgtgtgacagagaagaaaggcccatcatcatgagtgagattattgacaaggaaactcttcagaccatatgtaaga
gtgatatacgttctcttcttggaatgatgaacgccaagcatggcacattgggaggtaatttttacacttctatgcccgctcaactaaaccgtttg
aaaatttccaatatgaagcaatgggagctaatgcagtgctaatggcagctgaagctatttatgatggattcagagaccatggcttaaacccatc
agaatatacttttcctggtttggaatctgctgatgtgtacggaaacaatccagtggaaattgcaatatcaggagatgatgacaatatgttgttgaa
cctcatctgcaactatggtgtatcttatgaaaaaactcgcggtcgggtaaatagatccttgttagattttttaaaaatgaacacagcttcaaagtgt
ctcagtgttttaaagtttgttgaaaaacactttaaaattgagtcaaatacaccaaaaggagaatttgaagaaaaggccgaaacatgtgttaattgt
cttgatagaaataatgtgttgactaaagggtctgaacaagaatcttacaagctttcttgtggacactttcttcatgtaaaatgtttgaggaatatttg
tatagtatcacaacacctaagatgtgaaaaatgtctaaaaagatttgatgagagtattttgagaaagtgtacacctaacttgaattggtggttgac
tatgccggcaggtgctggaaatgaagaagaaatatgtttcatgagaaataagaaactggttgatgatttcagaaaattgttgtcccctgtctcaa
ttcctcatttcttcaaaaatagtagacagcgaaatcttgatatgttgtgcccctacagtgaccacacgataataccaaataaagaagatccaaag
aaaaacgaagatggaaacagagtgagggtcaaccacacagccatcagtgaaaagcagaacaaggaggaagaagacgcgaggataaa
gcgtgtagccgtcaggacatttacagccatcagagaaaagcagaacaaggaggaagaagacgcgaggatcaagcgtgcagtcgacatg
gctgtcgcagccatcaacgaaaagaacaaggaggaagaagacgcgaggatcaagcgtgcagtcgacatggctgtcgcagccatcaacg
aaaataacaaggaggaagaagacgcgaggatcaagcgtgcagtcgacatggcggttgcagccaccaacgaaaagaacaagaaggaag
aagacgcgaggatcaagcgtgcagtcgacatggcggttgcagccaccaacgaaaagaacaagaaggaagaagacgcgaggatcaag
cgtataattgacttgactgttgatatgaggattcaacgtatagtcgacatggcaattgcagctgccactaaaaaggacaagaaagaagaaga
gaaaaggacaaaaagggaacaagagttaagggctgatctgagaagggcaatggatatggtgaacgaagtacagaagaaacttgaagac
atggaactagaaaaggggtgtaataaggatgaagccaagaatactagtaatgttgttagtagcagtagtgttgttgcctattctaaagaaattgt
accttgtttaggaaataataataatgctgtcattggtatgactagcaccaactattctgccaacaatactaagaataatgtatttggttcacctcata
aattttccttcaacgatgcatctagattctccaatattgtagaaactcccaaaatgtctttcaatttctcgttcaagacataaaataaaatgtgtaac
atgcaatactgtttttttgtaaaaccccttgtgtctggataaaataccacacatctgtgaacatttcttgcacacgtaacactacgacttttcttcaactc
acacatctattttttgcacatgcaagtaaaacatttcttcaaacctctacagagagatggtatataaaggctttgtatctccatcatttatcactattc
gtacgatgacgtccaacagaccaacaacatcccctctttcgttctctgaaggcttctcgctttcaggggataagtatgatacgtatgaggatatc
ttacttgaacaattcaactgtttcaagacatcttctccttcttctgctcgtaaaagtgaaatagaggataaaactttaatctttcaacttaaagaagg
agaaaaattccatcttgcaaagggcatagaagagctccgtgagattctagacgataattctgcaacaattgaacctattatttctccaacaacat
tcaatgacagaaacgaattactaaaccacgagggagatatatcctcaagtcccctatatactcagataatgaagcatatttcaccagagcatg
atatttatgaattggaccttattgttggcactgatttgctttttggtctaggtgtgaatctacgcaacgtttctaaactgatgaagaaaatatcgtatg
gtactttaaatgtagttgatgtgtgccacagaaaatttttcaacaataggattatagttaatcccatttcttcgtcattctcaaaaaaatgtgtgtattat
tcctctattttctgcagctgaagaattttcgtctctgggggaatgcagggatttattcaacggtatttgtgatgacgtagagagatatatcaactctt
attttttctaccctgaaaatactactactactactactgctccttcgtcgcccgaaatggaaattgcagatgaggaagaacaatccccaaaaact
ataaagagagaaatgacaacgcaagtagaaactggtctggtgtctgtttgatttttgaagtatttaaaaacacgtactacattattaatagaggagat

agaggaggttcttttgaaaaggctgtgaagagtgcaatttcttctatcaaggaaaagagatgtaaaatcacagatattaatggtaataaacctc
gattggttatggtgataactgggtgttatacagaattgtacttcaaagatgcacttaaacagattggagaaaacaggcgcaaatttttgaaaatg
aatgggaattacttttctctgattgatgaacaagcagatctaatcgaattcgcgatgagtgtttctggtgccggggagaggatttttgttaacggt
ttggggatgttccagaaccgtaaaatgatacctgtaattgatcctctcacatatgaaaatgttgtatgtggtgagcatgatatacaaaaagaaga
tgctattctttctgtaaggagagctattgcagactataatgactttgtaagtaagaacaagagagggaagaaacgcagcgcagaagaagaaa
atgaagatgaagatgcagacgctagcagcagcagcagcagcagtcctcctccttcttctcctcctgcacataaaaaatcacgtcttccggat
gaaggcgaaaaatgtacactctgttaatttttcaaacaataaactaaccaccttgtatatttttgtttgtttgcaacacccttttatgataccattaa
gacacgatggcggtaaacttggataatgttcttgtgaatatcaacaacaaggatgaagatcttacaaaactcgtatccgaggcaataaagcg
gcgagctaaaactgtatttgacactaaaaatcaagcagggtttgacatgagacgtcaagttgaagctgcattatatgaagcaatatccaaaaa
gaaagaaaaggccataaaggcattcgatgagctcatacaagaaagaggtgatgaaattacacctttgactacaatgcagtatgaagagtgg
gtaaaccgtacaataactccctcattgacgactgaaaatttattaggtgatgttgagcacgccgatttttttactggaccgaatgacacccgtaag
cgaggaagatattgaaggtttcgctgcttctacttttaaggaggtatcagattcaaaaactgcaacagtcatagttaaggcagattgtgaaacg
ggggatatcgatgaagtgtataatcttgcaccatcattcggcgtcactcaagaaattaaaatatataggtcaaacaattcctcggaattggataa
tgtcgcagattctttccatatttataaaatttctgcaacagatagcgacagtggcaatactaaaaaattgttgtatgggttaaggaataaaaaagc
aggttatacgtgtttgtgtagaatttttgcagaaattgaatcagatgggattatggccaatacaaatatcggtgtcgctgaaaacaacagagatg
aaattgatgaaaacgaagaaggtaaatatggtttttaatacccaaacaaccagctggtgcaaaattgatcatctacttctttttaaattgttggac
ataaaaaatgatattttatacaatgcaacccttccttgggtttctggtcttttctgtgcttattgtaattgtaatgactgtattagctgtatacactgcac
ctcaaataaagaaatcgaagaagagaaaaattgaagatgaaaatgaggaagaacccgtaaagactttggaagattttgtaaagggtcggct
ccttaacgctgtcaaggaaaaacctgcagagtactttgagttgctaatatctgcagacactgaagcagcattaaaaactgccgaagaaacag
cccttcgagattttgttattgagaacgactctgtcgaaatagatgtggaggaagtacttgaagagaaaccaagagaatatgtcttcaaattggc
aggcgcaacaagcgaaacgctaacaaacacaatcatcgcagaggtacaaaaaaaggcagcattaataacagaagaagatatcactattaa
aatgttaaaacaattcagggctgcgaacaaagataataaagacggggaagcaactcctgaagaaaaggaagattttaccaataattcagatc
ttgtggggttgtacttgaacgaagtagtagaaaaaacaacaaatattgtcattaacaaaatattccctcatgagatggtttttgaaagatgtgcta
ttttaattgaagattttgatactggtgttgtgactgatcaagccattcagataccctccaacaaatacaaaatcagattagtcgaaggggatgaa
cctgaagtattccctggtgactgcttggatcttgcagtttcagttgataaaataaaccacgtcttgaaaatttctgcaaagaacggatgtgaaaa
caactgcttcgttattattccacggttctctcctgtaggaagtgtttcttccatgatattgggcagcactgaccaagtcaaacctaaaacattcttat
ttttagccaacaaaaatgacagtacacattttcaattcacaatggataagcaacattctgtagggtgtgagttggacatgttaatttttcagaaag
gaacttgaggaatttacccgattcaaaacctagacctctaagtgatgcagacatattggcctcatatgggaagcgtctaggaactggtgttttc
acaacagaaaatttggtagacgattaaataaaatttataaccactttatctgtagacttttatgtcctttgtgcaacaaaaataaacatgtcttcttcg
tcttctgaaactcctaagacttccaccgatactggggaagaaaggattaaagacattgtaaatgctctagataataatggcgagtggttgtcttc
ctatattgatccgattatcaataattacatctcacgaaaaacggcagaaactgtccaaaaaatcaaccaagaagttgatgaacggtacgatag
aaaaatagccgacaaaatcaacgaaataaaatcatccatctttacaagtgctcagactatgtatgaccaatatgcaatagacacatttcaagaa
ggaaaaggagccaacgggactggaccagtcatggggccagtgaacacggttatcgatacaactttaaataaaatgaggggaaatatgctc
gaatacgctgaagatatgtgggacggagatgactggaaacgattttccagttctatgacaacgcttgaatttgatctaagttactctgatttaact
atgatgcgtggttctgacgggtattttgcattccctttccgtggaacaaaaaagataaagatggacggttcaagaaagaaagaagacccaatt
aattgtatcatttcagtaacatatccaaacaaagtaggggatgagtgggaagagggtaaagaacgtgaagtgaattttaacctagaaagagta
gacgactatgaaagagatatccatgtttcaattttgtgcatgttacatgcacaacttgataatttcgaacaagcattaggagaaaatgcaaactct
ttttattttaaaaaggggcaaagagtcatgttcttacccaagaaatctaaactgttcaatagacctactgtagaagattctgatatgttttctataata
ttcccacctgcatctgaccaagattttgcagatgatatttattatcgaataattgtaacatgttcataataaaatttgttgaaataaaatacagtaata
cattcaaaatcttttattatttttcctttctaatccaacatatataatggtgatacatttacaggtgatacctgaacatatttataaaaatcaggaatata
cataaacatgctattgtaagttgtattcatatccctggtgatcatctcgtgccagatgcccctcagcgagatggagcttgctgtaatctttgccct
gtggagtgcttgtttgtatgcgtcttgagccctcttcatctcatgtgctgtagtttcatccccctcacaaggctccatgataatattctttctggccg
aattatacttgtaatcaatgaggtattgcttgaaaggcccagtcaagatgtcgttggttgtagtctggatgaggtcacatttcttggcaacattctt

aaaggcggcagcagcagtcttggcagacattgaattcgcttcaatgcttccaaaaatatcctcattattctcgtgcaccttcaagtagttgccac
tattttctcccacgcctccgtcaacaataaggttgcgaacgtatctgcggctgtgtgtcttgttctcaacaacttccctctttagaagattttcaacc
atcttagtgtacatgtcagatctccaattgttgttgatgtcgttatgttgcccagaaaagttccctggcaaactaattctgaaggtcttgtagcttcc
ataaggagtagtgttattaatgtggttaatgggcatagattcacagaatggaatagccagtgtatggacagtagcgtataccttggcagtcaca
ttatcttcacatagagcattaataagagcactcttcttgctctcacaataaggcttcacagacttgtcgaaagaggaccagaaactcttatcataa
tctacctccaccattgaagattcgtatgtagtggaacccatagaaaagataataagattgtccacatcgacaagtactgagtagtactggctgtt
ttgcagatcaagtccctgatcctcctcatcttcatcctcatccttcttctttgtagactcgatgtgcacacaatcctcagcagcctcagttccggca
gggaacaacttaagagacttgaggatagtggacatgttgaatgagttggtgatgcatccagaggagtgtctgcatagagatagacagacag
gagttgatctgttctttgataggagagaaacatccatctcttcctcagattgaggatatttggtcactgaagcagatggtctgaagctgttgtctg
aagccaggtagagagggttcccatcgctctcgtcgaattcgcactctgttacatccccaaacacaacttctgaacactgggaagttgcaccat
catcatcctcatcttcttctgaatcttcactgctactgctgataacactttccttgtcatcttcatcctctactactgttgctgtatccttatccttatcct
cctcatcctcttcctcttcttcttcagcagcagctacaacagtgacagatgaagacgaagaggaagaagaacacactgatgaggcgtcgtca
tcatcatcatcggagtcttcttcttcttcctctgactgctcaccctcttccttttcctcctctacatcactgagaccgatagtttttctaataagattctt
gatgacatttccaactgaactttcaatagtttcattgttgtaatcgtaatcaacttcctcctcgttttcatcttcgtcttcatcctcactgtcgctgtatg
catatgctgacttttcgtcatcctcttctccaaatacagtattgttcagtgacttgaggttactctccctcttggccaaccacttgtcgttcttgaagc
aacaacatccatttctcatagattcagtgacactttcttccacaatttggttagtggtcttgaaatgtgttcctgccttcttggtagacgaagaactg
ggagttgacatgaccaatccagttgacttgattgcactatcaatatagtttcctccaagttccttaaacagtgacttatctccattacttccaaatat
ctgcctcttcttctccaaatccatctcgtccacatatacagcatcagagagcttcaatgtgaagagagggcctcccttatcaagggccttcttgtt
gagtgtttcaatagtcacgtgctcatggaactggagattcatacaaccaaggaatagattggccagttttgtagtatcagtggcgattgcagttt
tcttcacagcttctccctcaaccttgacaatgttgctaaccagatcctttccttcaaatgccgacttgacctttgagacacacccatcacaaaagg
tggaaggggacagttctgtgcgtccagagttcatcacactattgaacagtaccgatttctgcgccttcttattcttctccaattcttcgtcctcatct
tcgtcattctccagaaattcagcaacagatttggccagaacactcacgtcaagatgagctggcaccttgcccataagagcgtccagttcttga
agagattgctgcttcttctcggcgggtgtgatttctggaggagtctgctgctgctgttcttcttcctctcctacaacttcttccacagtttccgtgtgt
ggactcctgatagtgttcctcctctgcttagagaccattcttcttccagcagcaaacttggccttggaaggagttggtttcttggcagaagtagta
gtagtatcctcaaattcataaacttgatctggttcctcagagaaagtgacagttggctgctgctgctgctgttccttgattgactccatttttagaat
attgggttattaggattcagtagttctgttattcctagacaacacgtcttgtctgtacgattagtcgagatgatgtgatgtagagcccttgcccgtg
cctcaatatataccccctcagtagaccttgataatcacggttgccaaacaacaacggttggaaatcttagtagaaagaaccaatggaacaagtc
aggataataatatacagtataagcttttggggaacaacattcatttattgtcatagcgctaacattagaagtacaagaacaggtgattctgcatcc
agctcattttgttcaacccacttcatttgcaggggacaacactcttctttcttgaaaaatatggatgaagacacttggaattcttcaataggccctg
gggcgttgctggaggaggacgcagaagaagaagaagaacagctgccgttgaacgagtttatagtttccagaagacaacataatttagctgt
aattggacgaggagacttcttgtaatgagagaaccacaaaattcttcttcccgtcttttctggtataaaagtagtcccataatatgttcctttctgg
actaccacattatcttcgtttaagggaacatatctgaccgactttctcatcaagggctgatctccatttcttgttgtgcttagagccgatttgtattcc
ccgtcgttcacgttatagttgcatatagaacagtgggatgaaaggaacttgtcggttacgtgttcagtttgtacaagataactggacgcatttgc
tagttctggagggacgtcctggtttttgtttattgttgaagtaggttttatctttttcctatttaagcacacaccctttgaattaatgtgtatgcgattcg
cattgatgtatatgcagattgtgagtgctggagaaaatggtatctcaaaatcggtagctggttcaataatttctggttcaacttcctgttgttcttctt
cctcatctttctcttccttatttttgacacctagaactctaaggttggcagtactaattattctggctgcagccgtgctggtagcagaaggggaag
attgttgggagataacaggagacatgttgaccgtgccgtgtcgtgaggataaatggctaaaggtatatacggcatgcccttttttccgttactgt
gtgcgaggtcgttttctaggaccgtagatgaccatctgccttcctctctttttctggacaacctttaattgccatggtgttatgagaggtaaactcg
cctcatagaatccgcaaacttcctctaaagggaacaagtgtttccccataggggacaattcaaaaatataattccaacatattaagggtgttgtt
ttattttccttttcatcgtactccgttattgttttactaaaatttggaaggtgatttttttgatagactatcggacattatatccgctgcaagttgacgttta
tgttcaatgacataatcattcgaatcaaagttgtcgtgtgttgacaccataccattccatttccatttggctagtttggagagggatttattgttgcg
caattttagacgtgcagaaacgttttgccatattattttagaaagtacaggcaaaacaacatccctcacgtcaatactgtccattaatatatcagta
gaatcattgaaccttataacatctctctgtatatcttcgtttgatggggccaatcgaactagaggaactaaatccccagttatgagatcggtgaa

aataaagctctcgtcatcaacaatttttttggagcctatgtacatctcacacttactataaaagtcatttttataccctggtttcttttcccacctagaa
cataagagtattaaaactctcgacaaattatacatggcacctactcttccttccatattactgatagctctatacatattaaccacgtattgtatagg
tgcttctacgtacaaatctgaaaatccactagcggcagttccttcactaaaggctagaacttgttccatagtcttctcttctgaaaacatttgctga
attattttagtggatagtttgtgccgcttattgaattttgtgccagggtactgggacccgaattcgttgaaatgttgttgttgtttttccctacttcttta
gtttcatgccaggcggggtgttcttgaagagcagccacctccaggaacaaatcacagatggctcgtactccatcacatgtgcattttacagttt
cagggaaatttaatcctgttattttataaaacaactctctgtctattatttttcccattcgtaggcgattattgacattaaacccacaacactcgcctc
tgattcccattataaaggctaaagaatatactataatatttcccgcacttgaacccacgagtctaagagttgtatattctccatcgcagttatagag
acatcttgccaattcgtccattgatttcctgcccgtgaagcgcacatccatgagtcttgctcctccatatccaatttcagcgtttattgaagtcctc
caacctttatccttgtttaggcgctgtatctttttcgccttctctgccaatacattcataaaagtgtaaaatttaacgtctcttacagattccataattttt
tcctcaagtgttagcggcggcggtggcggcgatccttctttgtagccttcaaagataaaagagtgttattcaacttacagtgtttattggcccag
caattcttttctttcatcaactgcatagtagaaggtcgatcggaaagaatgaccaatttggaatccaaatccctaggtttatctagtccagcttcta
tgtccatactctttcctcctattatcatatagtcttcaaacagtgttttattagcagacatgttaatcatttcaattacagtctggaggcatgtttcatgt
ctattgaccagatcttcatcatcttcatcccatgtcacacccaccagaatattttctcctttcttgagtttggatacattttctgtaatctttttcattatc
cgttgtttcttggacgccacactttcgtagcatagcgtcggtaggggagagtcgcaatccacaaccaagtatttttcatgattaatttgttgagg
attgtgttctttgaattgaagaaaaaaatcattcgttctgtaccatggagacaactatggataatgtcgttcagaataacgacgtaacaaaacca
acaccagatgttgctactgttacaacagcaactgagaaacgtcagtcatgcaaagagaaaaaggatcaacttaaggccgaatgtcctcaagt
actgagagcactaaaattgtccaatactttaaaggcaaattttggaaaatccatgtcggctatttttgctcaacatttagtggacatgacaaacgc
taaacactttaaggacccaaagacaaagaagattttagaactggatggaagtagtagcagtgacagtgaagaagaggaagaaactagttctt
catccaaacggaaaagaggtagtggtgctagaagtgcttcttcaaagaaagaaaaatgccccaatactatcaaaaattggctcaatgatgct
caaggtgtattccgccagtttgcagatatcatcattaatcttccctcttttgatgatcttagagacgaagtaaaggatgaacaaactgagctaaa
gaccatatatgacttgtatagacaggacatggaaaaggtggtggaagaagttttagggcgccaagacctgtttgatcacaagtcagaaatag
ccaaaggtttggcccgtttcgatacccacgtctcgttgctcccttcggataggtcggctgttctagactcgtccatatccaaagagttggaaaa
aaatagcaagggcccgaacagtaatatttttgacacactaaacacactcaaggaagaaatcaaagaacttttatgtcatcatgtcaaatatttat
tgcaaaatcttacaccggaggatgcaaattttgtgttcaatagttctgtaaagtatgtaaagaaatcatatcaatactacatacaaacatcagaga
tggaaagtgatgaatttaagtcccttctcactggagtcaacattaaaatattggagaagataatctcgtccgataataatgttgctactccttaca
aacacatcactaatcccaggaacattatttcgtctttacaaaaagtacgtgaaactaaacctgtttcaaaggattatccgttcagagtggatacg
gccagagatattgtactacttccagagacgggtggcatttctgatctccctatcaagcccgttacattattgcagttggtgtcctacatcaacgct
ctcttttccctcgagcgccgaaatgttttcaccgacggctttttaatgcagcgtgcgtcctaatttcccagtgcctaacgaatgcgaacctattat
ctaacgactttcctaaacccattgaactggcagctaatgttactcgccataatcttctgagtatgaaaatgcttcaagaaggttcatccagtgaa
aagaagagtaaaaagaaggagaaaaagaaggataaaaagaagggcggtggtggtggtgacgattctgattcagaaacagattcttcctca
tcatcatcatcttcttcttcttcttcatcctcctcctcctcatccgaagacgaagaagaagaaaaaggagaagcagtagaaaaaggcaagaaa
actaaacgcaaaacaaaaaagaaaccatcaaaggacgacgatttagatacaattagtaaactgattctaaaaacaggaggttacttccacga
cacgagtgaactcggcaataaaattagaaatttaatagacaaggatgattttgcgggcgtagcccaatatgcagtaacaatcactgagatgca
atctacgccaatgaatcaaagattagtatctagtcttttagatttgataatgagactaaaagaacaagtaaaatatagtgttgatactgaaagtact
tcctccaccgccaaatctaataatgctttagatagtgctaaattgacatctcaacaagtggttacaatgatggtggattctggagctgaattggc
aaggctagctgccttctttttcgtggtggtggataatactgtagttaatcgccatgaagcattcattctaacatcaaaacttttaccctcgaatgaa
aatagaggacttaaaactgttgtagagtcattctttaaaaatttaacaattagtaacaaagtctctacctcgaatgaagaaatgatgtcggtgatg
ccgttcgaagacgaacaacaacaacaacaatgccctcaacatgaacaacaaccggatttgaaaagagtggtgggagaagtatttctagaaa
tgggaaaatcaatagtgaactcattcccttccaataagagtgtacaattaacggctgacgcgttcaagcaaaactactcgcctatgggaagac
gcataaatttggcggccaagataaaaacggctatatctatcggatctaatatctcgcccaatatactattttctaacctcccagaatctgtaggg
aataatactgtaactggtctaaggctaaccaacctattgaaaaacatatcccaaagtgcccaagctaataatataatcaaaaatgccaacacg
ctcgtaaacaatacgatggaccaacaaaactcagcagccatgtccatactactctttcctccaacatcaaaagaaacttctatctttcccggaa
acgatccttcatctataaaattacaggatatgactacaatgtcaaatctggcacgaggattttattccatcgcagaaggatgtatcggggtcgta

cgctcgagggaatttgatgaaggaggagttaaagcgtacactttactagtggactcaaatactatggacatggcagttaattttgccgctcagt
ctctggaaaaatcaatgtctgaagccttaacaaataatgcgaatatgaacccttctaatgtattagaaggaggatcttttgtagacggtgctcttt
cttacatgtttgaaaaaaatggcagtgactgtgaacccacccctctagcaaaatatacaatgaaagatgtatcaaatcgatacttgaaaaaattc
aacaatgataaaaatacacaagatttatataaaaatagagcagaaagagcactcgtcgaacaggttacaaataagcctacaagtgttgtgcat
tcccagctagcaaacgccatgggggtagctgttataggcgcagcatcaatcaagctcatggaagcagaagcagcggaatctgaaatgaga
gcagcaaattatcaagcaacatcaaaatctacaaatgctattaatataacaaatacaattggaatgatacgtaatactactcacctgtgtacgac
catcgccgtgagtgcagccgccgacatgtcaaagctcgccaacaaccattttatgagtgtattaaacactgcaaataacagtcatagcagca
gacgaggagacagatcgtccctgatgttgcagcaacaacaaccaacacattcagcatttctggaacagacaagaggaagaggaggagga
gtattaggatcaggaactgaacagacaaaggatcatgtggaacgtatgaagagagattggatattaaacatgatatctccagaagacaaaaa
tactactactactacacccagtaatgccggccggacattaggatatggatctaatataactggcataaacacgataaaacaagacgacaaga
gtatgatggataaactttctgagatgtctagcttcagaacttaaagtgccaattttcatataaaaatgtagtattatgtggtcgtttttaaactttaag
cgtttgtggttttgggtcgttcacggtccaagatgtaagagtccccttgtacccaaattataacaaagtgtcagtatggtcaatggaaagtatcta
aaaatccacccgtctatcattcaatatcccaactgaaccaaccaaccccagaaccaacaacatgtctcttgttgagaataatacccaagaaga
aatgattctggaaactactgttgaaggtgttgtggagggagcagaagttgcccctagaggtgtcaagagacccccttccttcttcctcatcttctt
cttcagcttcggatagtgaagatgatgaaggaggagagcaaccacaaacgaaacccccaaagaagaaacgtaatatcaatagcggaaag
tactggaaaatcgaaacaatcgaaccagcatctccagaaatgctcagtgctgtaaatgatattgataacgtgtccaaaactattcctctaatcg
ataatagttttggcgttcagtttaaaaagagtgtatctgaagaacagatcaaaacgctactcactgaaactattgccgtggaatatggaacaatc
acaaatgttaaatattcaacctttaatcaactcgaaaggaccggagagcctcttaagaagaagcgctcaaacaatggaaataacaactacag
atactggcaaattcgaattgaagccgctgcagcagagaatgttacacaagctgttttggacgcaatcgttgaaggaaatgacaccgtgatca
aggcaatcctcctccctgaaggggagggaatcgggcttcaatttaacaagagtgttagctctcaacaagctaaaaatattgtccaggctgcc
gatattgaatttggacaagttgcgcacatgaagtgtaatttgttccacaagatggagaaggccgatgaatcttctaattcttctggtgaatcacc
aaaggttaagaaagtaaggaggaacaagtcccagcctacaaattcttactacactttcactatgatcggggattctcttcaagagcgtattgat
aacgcaatcaaagttatcgaaatgtctccagttaagaggccattctccaattctgctgctgctgctgaagaagacaccactactacgaccacta
cttccactggtgttgttaatccacgagggattaaggatattcactttttcgattcatccatttctaagggatgtttcactgtgaggaatattgttgca
gcaaatggtgaagttccacaagaagaattcgtttctgagttgtacactaaccttctaaaggttgaagagaaggtggatcatcctactttcaaga
agcttattcatgaccgtacaatgaatcgacacattaaggcatggtactgcatctgcccctactacaccacaggaggcgtccccccctgcagct
gataaggtttccgctaagggaattgctacctacagaatatacgaggacaggactggagtgttccaattcgatggggcacatacatctactacc
ccagcacaggcagccgaggcaactggtgctattcacaagtctatgctcttccagagtcctggaactgatattcaaaagttcctcgatgctaag
aaggcagaaggtttggagcctatttcatcgggcgaaattgtgtaccgttcaaagtggagccccaatgatagcagggcaacacgctgcttcaa
attttattcctcttcagacgagaagatgaacattgctgatgttttgtctattgttcatactgatggacttttcagtagtgtacactttagaaaggatact
atggaatatggtgttgcaaagagcaagagtaaaattatccccaaaactatcaagattaagaagggaggagatactttccatagcgaggagg
atattgaagtgcctgtaaaattcactgcaatcacttcggaggaacttaatagggaatgtaacaccaagggaatgaacagcctccgagcacac
aagaaacgaaagagcaactcttctactactactacttccaccacttccacttcaacaacagcaaatactccaaagaagacgaagaagagcg
cttctgctgccagtgacccatttgcaaagcttactttagattatgttgatagtacatcatttgtattttacaatatcagtaaggaaatggtgcagaga
attttggctcaggagagagtaaagacgctaaaggcagtcaagaacgaagagaagatggaaattgtagagggagaagaagcacaagaaa
cttatagaggaatcgtaaagatcaagacgaatgcaaaggcatacaatcttgccaacaaaacatctggagtacttttccctgctgataaggtgt
gtctaaagcacacattggaagatttgggggatgtacttgattttgatgttgtaagggaggataatgttaataagacggtcgcttctactactacta
catcatctgaaaataaggcaagcggaggagatgatgaagaaactccaatggaatttgaaactgacggagagaagctgttgcacgaattgtt
gaatgaataatgttgttgtgttgtattttgtatatagagaaaagaataaaataaaaggttaatataatgttgtcttttacattttcctaaacttgatataa
taaactatatagcaaatattaatgctattgacacatgttactagtccctagtcaacgtgtaattgtacatagtaacatgccttttcttaggcatttgag
gcaaataggaaaagagtatttacgatgggtgaccgtcgtgataacgcctacaccaggtgacagtatagagggtttcagcagactgcaaata
ctcacctaacaggccttggtatagaggaatcaataaaaaatcaacaaaaaccttaaaagtttggtgtttcattttttttacatcggggacagaaat
ggggttaaaaacaaatatccgagctgcaatctgacattggctccctacccagaggtccaccctccgaacttgacattcagtcgacccagcgg

tccaccctctaaactcgatcgagctgaaaaattttttgaaaagtttttgagatggagaatgagtaaaattctctagcggaaacagtaggttacac
actaccctttctgggtacgttcatatccagaaacgcctgttccagaaacagtaaagagttaatgtatgtttctggagcaacagacgtttctggtgt
aatctgacatcgagtcgacccagaggtccaccctccgaacttgacattcggtcgacccagcggtccaccctctaaactcgatcgagctgaa
aaatttttttgaaaagtttttgaggaggagagaggaggtaaattgatacagatagaggtatctttagtagaggtggtggtccttctaggagctggt
gtctccagaaacgcctgttccagaaacagtcaaaagttaatgcacatttctggagctatcccatttctggtgcaatctgacattggccctaccca
gcggtccaccctccgaacttgacattcggtcgacccagcggtccaccctctaaactcgagtgacccagaaaaaattttataaaattttgagtc
ggagatagaagggaaattctctagtgaatacagtaggttacacactaccctttcggggcgcagcaagtctccagaaacgcctgttccagaaa
cagtcaaaagttaatgtgcgtttctggagcaacaaccatttctggtgtaatctgacatcgagtcgacccagaggtccaccatccgaacttgac
attcggtcgacccagcggtccaccctctaaactcgagtgacccagaaaaatttttataaaattttcgtcaagggaaaacaacactaagttgctt
gtcaattcatacactagtgcaggttggttgagatagagagtgaaggaggctatccagatatctggatcaaccagaatccagaaacgtttctat
ccatttctggaacagccatttctggaaagggttacattttattatatttggtatattatttctggtaccacttctggcacattcgtgcaatctgacattg
gccctacccagcggtccacccttcgaacttgacatcaggccgacccagcggtccaccctctaaactcgagtgagctgaaaaaatttttgaaa
agtttttgagatggagaacgagtaaaattctctagcgaaaacagaaggttacacatacccttctggttacggtcatgtccagaaacgtctgttc
cagaaacagtcaaaagttaatgtacgtttctggagcaacagacgtttctggtgcaatctgacattggcccgacccagcggtccacccttcgaa
cttgacatcaggccgacccagcggtccaccctctaaactcgagtgagctgaaaaaatttttgaaaagtttttgagatggaggacgagtaaaat
tctctagcgaaaacagaaggttacacctaccctttctgggcgcagctatatccagaaacgtctgttccagaaacacacaaaggttgatatacg
tttctggagcaacagcgtgacagtagtttctactaagccttcacttgtacacatttcacttagtataatttgtcatcacatggtcgacaaaaattttat
aaaaatttttctgggtcgctcgagtttagagggtggaccgctcgctcggtctgatgtcaagttcggagggtggaccgctgggtcgactcgatg
tcagattgcaccagaagttgttgttgctccagaaatggtgggggagtattcagagtgtgtttctggaacaggtacacatacccaactgatcag
ataaaaattgggtgtatacattttctgatccgagaaagtaaaccaactgaacggtgtaaaacgtactgaaacattacatcacccacagtcgtcc
tcctcccttctaagagagaacatatcccgtacccaactacctcaaacaacaacaacaacaacaacaaccatgacatccattatggacgatctc
tacatgagctgttcttcttcttcatcgtgctcaagttcatctgaagaggagaatgaggtaggagtagaaggaggaggaggcagaatcggacc
aacagaggccaagaaaaagatcctccgtaaacgaaagagatcttctgttaaaagtacatcatcttcttcttcctcatcatcatcatcagacgatt
ctgactctgatagagaagaaaaagaaggaagaaaactatacgtggatattgcagatacaaggaaaccgcccaaagtaagaaaactggata
ctccttcacaaactttagagaacgacctctacatgtctagttcatcttcttcttcctcctcatcttccgactcatcatcgtcttctggtgaagaagaa
agtgacgatgatgatgatgacgattatgacccagataatgtccatgttcttggttgtaagaaggaaaaatctccccaagacatagaagctgaa
aaggaaaaggaggaagagtatgaagaagaattcaagagaatggcattaccttcacggataaacacatctgtagatgattgtgttatacctgat
cggattttaaccctcttttctacacttttgaagaaaaatagtttccagttctctcagccagtttctttcctccggttggtgatgaagcaagtaaacga
ggccatgaattcagcattctcttccatgttatccagttctggtatgaggttggtggaggactctttgggtgacacatctaaaatttcctcctttataa
cacctcaaacggatactagtaattcatcttcatcatcgacgttcgtcaataattgtacagatgaagatatcaagaagcgtaacattgcaatggga
agagtagcagaattgctgtcaaatattgcagccagttctaatgaggagaataatttccgccctgtagtttccttaatgcgaggaccaacttgtgg
aggttctaatgcatccaacaagaaactcaatagtaacaggcaaactattccccaagtattaaacaaggttatattttttagagaaatacacagtg
ttatcgcgctatatttatcgtccgtctgtgtgcagcgagcaatgaataatgacaacacaaattctagcggatacgctgaagggatggttactaa
aatcttgaatattattggtaaaattccttataatgaaatgagtagagaaaaattcatatccgttggaagagatgcactatatttgtaccagaatgtg
atcacggatatgactggccccaaacataacaagagactccgtatccctcaacaacaagctgatttttgttacattatagccatgttggttaatga
tgttcccattacctcagatttacttttaactggaaaggcgacaaatttagtacaatttgcttctgctatggtggatcctgcatatcgcctggctgtcc
ataaaatggcgtctgttttcaatagtagttattccgtatataaagtcctagatcttgaccataaaatgttattaagggctaatttaatcctatctattta
tcagctagaaataagtgtctcagtgaaagaaaacctagaacattaactcagagcgtgtatttgttcctaaatcatcttttgcggaacaaattgag
gtctagtggtttgaccagtgaagagagttctctaggaacagccgtaaaattggtgtcacagcaattaatgtatgagggtgtgactcgtcaaac
catcgaggacggatgtagcatgattagcggaaactttgaggatgaagacggtgtaacactgaaatgtttgggagccgatgttaaggacgtg
aagactgttggactatctgcattgctatctgataggcttagaaaaaacattagaagaaacgtccctttctactgaaaaaatatggaaactactac
gctcgctttcattgcaatgggtttgcttttcgtggtgttgatggtgtacatctttatacagtacacgtcatcaaataaaaacaagacagaggaaag
aaagggtagtaaattatcatcatcatcaacttcttctacttctggtccaacatttaagagagaagcgtctcctgaattaagagaagacataccact

ttctccacgatcttctggtccagaactgaatctaactgacgctgaatttatagaccctgaactcttttctgaagccagggtcagtcccgaccgtta
acggaaaaaaggtgcaaaccggcaactcttctccctcctttcgagtataagtagaggcgttgcaccgagccaagcatcattcgggctcattc
aactgtgctgacaacaactaagaatatacattgaaataacccaaataatcaacttttcaagatgtttggaagttctgctaataactttaatggtga
caagaaatcttcctcatcatcatcagctgccgcatcatctgacgatcagcagttaggtccccttggactgtctactgctgatttcaagaaagttg
ctgccattcttgccaacagaacagagtccctatatctgttgccagattctcccaactttaagaatgtgattaacaaccctaaccagatatctattgt
gccgtttctgggttcatcgaaggcagctgaaagtggtagtgcaaacaagaatgaaaaccaggctgaaaattcttctaaaggaggaagtgat
ggaaagaaatcatcgcagcagaacaagtttaacctattgaacaaggtagaggctgaagaaatggcctttaaacgtgtggctgaactcattgc
ggatactcctccctctaaagataatccattgagggatgaccctgatgctattccatcacgcaacccatgggtcaaattgactcagaagaatctg
gaatatcttttctgggaagcagtaacaattgaagtctccaatgataggagtatccgtagtggaagatatcttcaagccagtgaagtgggggag
aatccattcctaatgaccatcagtgttgacattagaatcctgcaaagaatggcacttaatgtcgtgtggttcttcaacagattcttccgcatggttt
ctggacttggagtagaaaacagggccaattccacctatgtagcaactagcgatgctattgcccagatctgggtagagatgctcctcaagaac
tttatttctggagaaaatgtgccccaggcactgaagtatttgaaggaacattatgagcatgtttataacaagatcagtaaatgtggacgtcagcc
atcctattttgttgttgaatttgaaagagtggacaacacgattggatttgtcaattctgatactgaacataatggatcatcatacatggaatacagg
tgctttgacacaatcaggaagaatgcatcatctgggccaagtggaggcggaaagagtggtgtcctgtcttctggaacattcttcattgataac
gaaatggggaataacaatagtagtgcagctgcagcttctgcccctgctgtttctgctggagtttctccatcactttctccatttagcagtgatgga
gatgatgatgacgacgattgtagtggtgatgacgtgtgggggaagaagatgatattcaacacatcaggagacggatcaggagaatcttctg
ggcagaacggtggtggtgcatcaacttacaagagatttaggtgtggagaaaatactgcttctctttctcagaaggaaaatgttcgtctcatggc
gatgccaaagggaaatgaagataaacaactactcaagaacattatcaactttctgaatagtgcacttaactctgttgaaaaccatgtaatgtgta
cagatgaaaatatcttcgatgaggatcaagccgagcattatacgtcaaacaaggaactatacaaggctattgtttgctccaacccagccaatg
tgtacagagtaatggttgaactgtttgtcaaccttattcttccccgtcttaggaacccaattgtgagtgacattgaaactgtacaaaatcttccttca
aacaatggatcggtgagaacaaagaagatggttgaacatggatgtacagatatgcgctatgacatccctccatatgcaaagggaaagattc
gtcttagtgccaagagagcttgtgagtgcagaaaactgtgcaaggacgtgaggtgctttgacaagtctagagaggcaaatctcacccctag
ccagaaggcaggaagggaggtggaggaaccttttccccgcaaccacaattctcacaggagtaacgctcacgactttactttctatgacaagt
acagggcaaggatgaacaagctcaagaaggattcaaagaagaaggtaaagaagattgacacctttacaacaacagacgattttctcctgca
agataggaacgcttttgatctacttagaaagtgcttcctttctgcctctcttcatcacattttctgtcctgatgtgcttatggtgcatagaggagaca
gcttcaatattaactttgcaaacaacaaactcgagtgctataatgaacgtaacggaattgaagaagttacttcctcacaaacggtcaacgccaa
ggaagcacttgaggatattacaaaaattaaaatgaagagaggggatgatattatagatgttgtgaagagtaaaggactttctttgagggaattc
tctaagaaggttagtaagattgtgagaaggtttaatgagatcacaaaccaactctgcaacaactgcaacgttaactcttctaatggagatgtgg
atttccacgtctttacttctgtgtgtgtctacatccacaacattattcctgtgctcgaagatatctccatttttgcagaattgggtgaagaattgacca
agcttgttaaggagtgtagagacgtggctggagaggacaagacatatgatgatattatccgcaattacgaaattactgtaaagtactttaagct
ctttaatgcactcgttaaattctgtcacaggaattataatgtggcagtaacctctgccattaacaggagagggtacatgtgcatggtgagcaac
cttgtcgggtattattgtaagctgtctgataacgctatccagtatcacgaatcactatgctctttgcactctagcatctcttatgcagactattatac
gtctcgcaataacaattctgaagatggaggaggaaactcttcttcagaaaagagcaatgcagatgtagccaagactatggcctctttctatga
ccagttcgataagagtgaagacagcaagaaaaataagaacaaaacttcaaatgagatccttataaaaatgttccaaatggatagggttttgga
tggcatggatgatgatgatgatgaagatagtgatagtagtagcagtgagaatgaagaggaggaggaagaggaggaaattgtaaagaaacc
agcaaagaagaggaaagtggaagatgttgatagcaataagaagacactgccaaaggaacctgccgttaagaaggtgaagcaggaagaa
gatgtggagatggaggaagtgaaggaagcagcagcagaagaagaaaagaaagaggaacaggaggcgaaggaggaagacgctactg
agtatgacgacgatacagaagaggacgagaaagcagtagcatctgatgaagatgaagatgatgaagattctaaagctattttctaaatatcat
gtgtataaaatgtagtattttaaaggtataaataaacacaatataaagttaaacattgtttttttatttgacacatgtacatcatcaatttcacatggtta
tatttcttgtagcttgctcggcagcaataagagcaagagaagagtcatataaaacagatgtctgtgaagcagccatgatacccctcgatatatt
attaataaggggagttgctgctatattaacagcataataaatcaggttcttccaatccctagaagcccacacattgggcatttttatataaaaatc
aacgtctactattgacctggcaacagccactatacctagctcatcaatgttaacaatcttcccctgaacacccattacaatttttgatttttctgaat
tctctcgtatagcagcaatacggctagcttcattatttaaccattccaattttacacttgaaacataagctggggatttttcatcagttacttgagata

caatattattccattgagataaaagttcctccaattgagttgtagtcttgtccgcttgctcttggaaactttgcatgacgatagccgaatctcttgtt
gcagtttcgatgatggaaatacagttgagccattgcatgtcagtaaggatatggcgcggattaactggaacaaacagattctgtgcaggaga
aaagttcaagtcttctggaatcttgttgaggttgatattgtccttggtgtgtttccatgggataggtttctgtgcctttgaagcaataaagtatgatg
ggattctactgtatttctttagtgtttctgaaatgtttgtctttatacggtcctttcttccatcatccctaatgtaattcaaaagtttagatgtgaatagag
aagagtctctaatttttgcatttgttcattgtgaacctttatatcctcgggaacagtattgttattattagcctgcctctgttgacgctgctgctgttgt
tgttgttgagtagtacttccaatggtatttaatttgtccaatattgaacgtaattctacaggcacccttctaccagttccaagagcacctccaccgc
cgatggccgctgcagcttcttcactagcttcatcatttataaaaagtctccccgaagaagcatagcgtaaagatccatacccttcatttccagtat
caatagtaatcccattttgttcagctgttatcctttccagtatttcttcaatggcctttctaggatctagtgcaacatataatccatcgaccataatatg
agacacgtgctcgcacacttctttaataattttcatctcttcgttgctgacaagcatcatcattctagatataatagtagcaataacattcctaaattt
gttctgaagaaatagttcgatctctgtgaacgtaggctcgaaacgggcaggcacattaactggaacaaccgcttgtgtttcttgttgttgctgct
gttgttgttgttggatcgctcttcttgttccaccacctactatattattactgcctctgagttggtcggctgcaatagcaatgatgttttgtacaccttt
accagattctgcaattacttccatttttgagtcttcgcctgaaattggtacagaaggaatcttgggtgacaacctttctagagttgtgggtaatcca
gtatcatactgaagaactttccaaataaattcgaatccagttcctgccactgtgcccactttactggggaaaatatcgatcctccctgtagatag
aggactaacacattccttatagatcctcattgggcttaaatgttgtgtcgatttagtttttgaggcttcataaagcacagcagacgagagaaggg
gttcaaagaatacagtttccatgcccaattcctttaccgaatatttgacttgtttagatgcatgaacaggaagatttgtgggctgagaagtcgtac
cagggaggaatgcgctaacatgttcaattttcccatcaaccgaaaaacttgtccgtacaccatcaacgaatgaagaagaagaagaagaaag
ggatccagtgtcgtcttgatggttattaatatcaccaatttccatatcctttataccatcatcgcgttcagtattctttacaactttaaagtacgacac
caattcacctttatttttcaatccaaggaatccatcttctgcgatattctgaccatatgtcacatttcttgttagagatttaggtggtacataatttaag
aacgattttgaaatggtttcagaagaaaatggagcagtggcagttgtttgctgctgttgttgtttttgttccttagaggatattagaggtacattaat
gaaaagttccagatccttattcatctttaaaaaccgtcctgttggtacaggaagattaactaccctaataggggtttctgatgaatatctggagga
ggatgatggagaagaagcgactgaagtggcagactttaagttttcggctacattatctccattcaaaatactctgaggtgtatcggtcagtgtat
agtatctgagagacgtatcattttctacaataaagggaggatctctagggaaagcaaggcacacgcctccattatcggctgccgaatgaata
actgcataatgtgtaacaatctttgcaattagatccatcacgggcttcttgaataataaaaatttgtgttgatcaggatcggctgtatattttagtag
cacttcacgtactatataatctgttagttgtctagcctcctttacagcccatacgcccgatttcaaaacgcaatcaaaaaatggaagaagaacaa
cttgaagtacctgctggtgcatgtacatccaagcggtctcatccactttcactggctgtcctctcataatatttcttgtcattgtttctaaacacgac
cgataaagacccaccacgcagttaatgggatcgtaccatctgtactgccgtccataacttcttcttccggccagtataggctcccctgggaata
gtatggaatcgcagttagattctcttccaatattgtatatggatgcaatctctcgagcaatatctcctaccgtaatagcagatccgcccgagaaa
agtcccacaacagcatcagagacatttgagaattccctcccttctagatttacattattataatatatcagagatgctattatagcagcagaagat
tccccagaatcgtctctcagaattaattttaacgtaggaatatcaacatctggtataaaggctgtagtattattttgaccgaatgtaagaaaccttc
cagaatcgtgtatttccgacacgttactttttctgtcgacaacgagtgttttattatgttcagcgtcgttaagagaggttaaaaaaaggtgcatttct
atccagtctgttttctccgatatacagtccatgattgaagtgtattgaggcccaaagaagcaagtaactgggtttttcatggaaatggaagtagg
tgcccctgaagggacaatgttggtatcggtagaagtttttttgcctctttgtacgttttctattttcatcgtattccttagaaagggtggaagatacat
ctcctaaagattttcctttagacgataaaatcgattgtgcgatttttttccttttgttcggtcgtcaatgaggcggttgaggaggaggaggaggctg
ctgttgctgttgctgttggttcataaagtttgggttgaggtttatttttaaatgatagttgataataccgtgtaaccaaatcctttaaagaattggtattt
tttccaaaaaagcactgggccgtaaagtagtgcataccaaattcagtcgagttaaattttgacaactctggtatgataatggaagaaactgactt
ggagtttgaagccatgtcatttaaatcacgtagccttgtaagagagggatgttgatagagaattttctttgcatttgggttcttactggtgtcagga
aatgctatctggagggggggtgtttatccacaccttcacatccattagacgaaatatcacagggtgttgtaataatttccctgtgaatctagaggt
cgtaagaacaatatcaagtatgccgtgatgggcataaaacacaccataggcgtctttcaatactattgaggaagaaagtctttccatttcccatt
cttcttgattcagatgggatatattgacaagttcttccatgccgtcgtttcctttaagatctagccgtatttctttctcatcatgcattgtccagtcctta
atattaatgacaacaggcaaaagccctgccaatctctctaaagacgaagcagtttcttccatgttcagcttgatgttgttaatttcgccttcgataa
acctctcatttttacgagaataacatcctcctcctcctcgaggaaatttttgttctgtcagagatgaaaatcgtcgtgcttctcgtgctctttttttcca
ctaatgcatttagagattttgtagatatttcttcaatagtcatctctttattcgtctctataataaacacagacgggtgtatttttacccttatttttgcatt
ttctgtttcacttcctaaacgcccctcggtgaggtttgcaagggtagaatcataacacgcaaaagttcccgcattaacaagcggtagagaaatt

tcgctatcgccatcttgttgttgttgttgttgctgtagcccttcagccaataggcaagctaaacggctattgtcttcgttgttcttaccaggcaggg
gccgttctattgtttgcgtttgcgttgttgacataatttcctcctcgacgtttctcgtcagcctgtaattcgagtagagtttgtttccttatacgggtgt
gaaccaatagtcggtaaaacaccaatccaccggttacgatgatggatatcaacaatatatagaatagggccttgtcactagccttggcagcat
tgtctctttctgcagctgcagtcaaatctttgtagtctgggcctaattcgctctgtacgtggttccgtattctctcagacacgtccatgttgttgggt
ggataaagtataaatactcaggtgtacataaaaacctctcacatgatggcaggaatcggtagaagagataatagacccgtattacatctggac
attgatccaaataaggaaataccctataacgttccaccaacccctattatctgtgaaaaaaacccccttcgtgtttaacatgcagaagtgctcaga
ctgtgctcctttccctccctaccccggcactgagaagcctttccctccataccctggtactgcagtagaagaggaggagaagcaaaaggaa
attgaggagcttctggttgaccaatctttccctcccccattccctggaaataagctgagagatatccccagaacctaccctctcgaatttcccga
gaagaaggagaaggatttcccttgcgttgacactaccggtcacagcgatatccccttcatcgatctggagaaaaccccaccccgtagtgac
gttaggcacggttaccactacttaatcaaccccaacaaggttggggagcttaaccatatcgttggtaagctcactgaaaagcaagaaaacct
gaacaaattggtgttggatgttgatgacgttgtgattaatctgtcaagcactttgaaggaacttgagaagctgcgagctggcctgtgcaagttct
caaaaaactagacggtaaaggtatgaccaaaccggtataaccaactttttatataagcctaatattataaatcaacaactaacaaacgcgaaa
actaacaataggaaccatggcctcatcatcatcatcccctgttgccctctcctctgtcgcatcatccgtgatgatggaaagagatgaagaaaat
acactgtccctcaggaacaggaatgtgaacaaaccaacacctgttagcgccgcctgggtgcctgttgatgaagaggatgaagatagggag
gaaatgagaagacttgaagatttttcttcagatgaagaagacgatgataataaatcatgtcattgtgaccatagcgatgacgatgacgatgac
gaggaggatccttcatgctttaagggtttttcagctggcctgtgctcttttgtgaggggtttctttggcttcctcaggaagtcacttaccaagaaa
caggtgttccttcttacaagcgcagccgttgctgctatttttaagactagagatgtggctaaaactgaagaaggcgcagcaaccatggaagaa
aattcaacagatgtgattactggaggagatggagatagtggtattgctgctgatgttgtctctctcgctagtgagggagagggagaaaatgga
tctcttttggaatctattgcaacaacactcatcaagactacaattgaaaatcttgtagatggtggagaagaaaccacagaattgtaaaatgtttttt
ggtatacaaataaaagaatacataatttatataatattttattgtttcctcttaaatgacaatttccttatgatacttttaggtgaaggaataaaggtag
gagaaggtgtggaagacacaattgtagtgtttccatacacatgattctcccacatgttattattgttaggtgggaatttcttaaacgtaacttgggt
tatacgtggagcattgttgttgttgttactactgctcctggttagggtgtttttcttgtgtttcatctccttcaattcctggatacacttctggatagatg
gagattgttgaggtacttctggcacgtattcgtagacatgttcttccacattatcatcctctccctcctcattcttgatattgaagatgggagaagg
tgggtttgaaggaagcttcagatgtcgtccgtttacatatgacacgtaatcaccgccattgcttctaggtggagatggcggcggtgttggcag
aggaagactattgtcagaagaaaatgacgacgagcagctcgacattggcatatagacactttcttcctctttatttgtacttgttggttgttgttga
ggtgtattggcaactggcctgggtgtaggcggactagcataggttggaccagaaaatgttaccagttttgtgacttgaggggtaacattggcc
agatcgtaagtggtagatttgagcatgggttcagagatgtccaggtccttatctggagtgttgaaccggacgtggctgatgaaactggattcg
gtgttgctcccgttcctgttcttcttattcttattcttctcccacggtgagcaagaaatgcaagtagtagttctgtcgatgcacttcacgagcatgtt
catggccagtgcgagcccaatagtcatggagcagccgactgccaatccactcaggaaagtgacgaggataatgttgtcgttggtgataaga
ttgtccatctcgagagatgtttcttcctcgactaccagacgagatatgatgctctggggccctgcttcgtccttttataccgtcacttggttagaaa
aaacgatagtgtagttcacagtatttgtcccttctggtcttgggattggggttctgatactctgcccagcctccagatataggtggatagaccga
gattcttcccttgatgcagtttctattccatccaagatatcttcaccaaataagtccttatgcaaggacaccagaatattgaactcgtgaacgcttg
acttgttgatgatgcataattcaccttgttcgactgttatacatactccttcaatagaatcgtatagcccggggttattggcacggtcctcttcaggt
aagaaaactgcaaatattgtattaggcacaacatctggcacgtttacagaatgacagtaatgtttatctataatgtcaccagaaaagtgaaaatc
tatcatccgacactggagaaaggttgaaggaaaatctttccgcatattcttgagttcccgcaataacttcttgtttctggagtttgactttttaggat
tatttgttgccagacttacagcatcgaaaatactttctcccattccttcagcgggattttccatcttacttttgtagtggttttcagttcattaattgtgtt
tcagggaggctataaaacaaaactcttacgcatctgttcattcagtagtggaccagacgtcgagatgacgagacatggtgtgcttgttccaaa
aggccgttctaggcatgttatcctaggaaatgtcgactacactttctgtactactgacaataattgtgtcagtctagacatcgatttcaaggacaa
tatcacagaccaaaacattcagttattgaacaagaaattgggtaagaaaacagcaaagaaaataaagaaggaagatgcacctgaaacaaa
ggaaaatagtgacgaagacatatacgccaccaaggaattcgaacagacaataaaaggtctacagacaaaaaaaggtgccaccgaggaaa
acgccatcgcggccgcagctgccgctgccactgctgctgcggtagaaaaggctatgctatcagaaagtgaaggaaaatcaatggtcatca
acagagctagaatggtgctttctaagcgagacacgtcccagaaacagttcactgcattgaagaacagggaatctttcttcagtgttttgatattt
gaaactggatcagtgatagttgtcgggcttcaagatccttcgcttacaaaattgtgtgtgattaaagccacgactgatattgctgatattctacag

aaaaacatcagtgtggctaacgtgtctatagtgaatacagtgtccacttttaatagattccacttgaactttattcgactcgggaaattcttcgaaa
gaaattgcatctcttacagttataacccagaaacgttccccggtatgtttttcaagctgcgagtgcccgcaaagcctctcttgcctggagagact
atagggggaatactacacaaaggttgcaatgatgcgcgatagtaaggatcccaattttaaaatgtctgactggttgaggataaaaactgcattaa
catttaaagttgggaaaattactgtgctcggagaaggagagagtgggtgcggtgatgtttctgtcgtatccaaattactatttggtttattccatta
ctttatggacaacaacattaaaatgtcccccaaagaagcacaaagagtcagagaaaaatacggcatcccgcatctagaatggtacttgtaca
ttgacatgttgctccactcctacccgtacgtcaaaccatcggccgagcaagtgaaaagggcgatggtggaccaacaacatatttctgaagtg
gataggacatactatggaacaaagaacagtatggacgctgccatgtctgcaaatttagtgccttcaaaagaagagagtatctccttcattaaaa
aaataagatcacaacaactttttggacatttgtgtaaaccttcaaaagaaactactcgacgtgctatagacacactttctttcgatcctataaacc
aagacaggtggtggaataaaaatgaccaatattacggtaaagagagatgtgacccgttttctgttgcacgtttagtgtctgtttctgaaaataca
aacagtatgatgaatagtcgcatttcttgtcaagggaaatggtggctagatgaaaacgaatacaaggataaacttgatcatattgtggatttgtg
tacagaagaaatagtggaggaatgtgaatcaaagggttttattgcctccccatttttgaggaagcaccagaaggaaaaaaataccaacgcctta
tgttttattagcgagagcctgtaatcaaaaaaatggtaacaaaatgagtattaacaataatagtaactatttgtcgggttcaagtagggcgaaga
ggaatgcaaaactacaggaaaaacaccgtgtaactttagccaggttgaacacgatgatggcgtcgtaccgatttttgaataattacatctcaac
agacattgcacctgattttgccaagttatttggtaatgatgtatatagtttattacatttaatgacaaacttgcctaaatcccgtggacatgctctaa
catacaacgaaagggccctttcaagtaatgaaagtacatataaaacacctggaaatgcatactttagtactctatttgaaaaatccattataaata
accaagaaactgctaataaaggtaacaatagaaaacgtaaattttctcgaatcggacaagaaaagagctcttttctgtgcaacgcgtgtggtg
tcaatttgaacaagggtagtgatgaaatcataaagggtatttgtacaagttgcgatcaaaatagtaccagttacatagagaatgcattatctgac
attaacagagacaagaagattaaacgttttaaagcagctgcaacccatccgccagtgaagcaagaattggtagattctttatcctcctcttcatc
tccttcttcttcttcttctcagacgtctaacaagaacaatagatgcacccctagtgattttatagattatgtgtacaaattcactgacgaaacaaca
ggtgctccaaaggtgggcttagtgtttaaaatgtgtgatattcttgcatccttagcaagcaggagagggatggaagatcgtcccacagccaac
tatagaacctccttacattcagctactcaaaataaaaccaatttgaataaactattagtttctgctatcaaggaaacaggagccactgaaactga
agcacagatattcaacaagattattggtagtgaaaagggactatcaattctctgtcaacttgtggaaaggaggaacaaagacaataatgtcttc
gactgatttgtctaagaatgccttccatgactgggtggtatcaaagacagattgtgaggtgtttgatgtacactgtgagacggacagagattgt
ggcgctgcttgcgagaacacgtactctgttgacggaaaggaggttacaaaattctcttgtaaccaacagtcgggaagatgtgccaggagtgt
ttatagtgcgtcttctctagaaagagcagccaatgatcttggccacattataggtatcatcaagaaaaatccaaaattggaggaagaactccct
gaatcatttttgtggtttatcaatcacaatggaggagatttgtttgtgaataagcgagccgcgtactacgacacgatgcatctaagcatagga
aactggataatgtggacactcttgcccagggtttagataaacggatggcctcatcattgagagagcacctactgaggaagttggactctatac
ttttacaaattgataaagttaaatatgaaaaggcaaagaaatggatattggatataacacaggaggctggcaccgaagaggacaataaagaa
gaagaagatgcgaaaaaagaggatcaatctcttagcgttagtgaaattgtggatgttttaacgggcacacatgaccctatgcccctgagggc
tagagggtttatccagaaaaaaatatatcctttgtcaagaaacgaattgagagaattagcccttaaggaacttttccctgaagaaactacatctc
ctcaggttttgagtaggcaacatgatgtatctacgcgtgaagatttatgcaatgaaagtatgaatgcagggagggcagaatccatttttagcga
ccctgattctggagagtacgtggctacttgtgcatgtctttactcggaatatttaacagggcctgcgtgtaagcacaaaacatacaggtatgtta
tagactacgacaaatggaaaaggactggaagacctgaatttctaactgatcctgtacttcattttaaaaaggcagaagctgtgtgtaaatcgac
aaatccaaacttgagggcaatttatagtccagataataaaggtttcttgtgtgcgcctgtagctgaacttgtaaagacggcattaacttttaggg
gttcacacgaaccgtctctcattgtcgagagagatataaatcaagctgaaaatctaccatccaattcatttggtgtaaactggccctatgtgaat
ctcctaaatcgcattcaagaccagtacacgtaatttgataaaaaaaagatcggggaaaatgtctgcatctttaatattggacgaatacctcaaga
agactgcttcagccgttctggatgtagctgactcatttgagaaaatcaagggagaaatccaatcacctgaggaggctgcggctctttctgttgc
tctctatggagcacctccaaaaccttcagcttcggctgtggcctctatcatcactggagaaagaacatctttaaacgacaaatatctatcggata
atgtcctattgaaaatgtctgttgctcgcgttggacaagaaaataatcgcaagagagccgaccaggcagctgatgaaattagaaccatcatg
gaagatattacagggagtttgtccggtgcgtacaggcaatatagcccgctcgaggaagaaaataaggtgcatataggcatcatgaataaca
aaacgcctagcattgtttgtggatattatacaatggacacatctatttcttccgaacctctttctctaacagattttcaaaaccccactgtcattgcc
aatgtgactaagcggatggagagcatttttttcaaaggtcgactctgctaggtctacaagattcgacgcttttgttaatggtgttgcgaataatatg
gatataaagtcatcaatagattgggcaaatatggtagaaaatgtgatcaaattaccagattctacacctaacccttgttcagttgacactattgtg

tccagagacgcaagtgtagttaaaacagcagttaatgatatatacgcttctgttggaaaatcttattgtcgtcctgcaacacagctaacctttatg
agcgagattgaaaaactgcgaaaggctgcagttgtatgttttgaggcactcatgtccgatactagggagagggcattcgtagagttcctatttt
acgttagctttaaggaagatgcatcaaataccaattcaaaattgtttgttcagaataagctatcttccatgtctggaaaccccagacagcccata
aaattggtacgccgttctgctgaggaaacactattcgggctctgtttcatgtttaaggtaatgcctccagaattcatgaactgtatatttaacttcc
ctaccattccccattcaacacaataccatggtctatatggtacatgtttaacccctctacttagaaaatacggttcttcattcgaaaagtcctgggc
tcattttgaggaaattttaagcgaaagagccaatgcagtgaaaaaatttggtgtaaacgatacgaggatagattgtctagatgcagtagcaaat
ctcaccggacctgtgtatgttctcattttagatcttgtacgtactctaagtgcgcagagatcgtgttcaactaaatttctccgtgaaattaaggaaa
actatcttttgtggaataggtttgtgtcataataaaaatggctcagacatcaaagatgggaactaacaagaggtgttttgaggaggaagtggag
gaagaaaggcaacaacctttcacaaagaaatctaaatcggaaccacccagttttgaagacaagagttcatccacatcttctaagaagaagag
caaatccaataaacacaccaagaccaaggaagaacaacttctagaattcgtgaaggatctggagcggagcgaccccactgttcctgatga
gaaggtcaagcaagaagttgaagaaaagtcccctgaagctattgctgaaatttttcaatgtttgggatcgctcaagacagcaagttcaagag
ccttcttcccattgaacgcataaagagcatcactactaaaattgttatcgatgcaattaatcagcctgtgcgcaagatgttggttgaccacctcta
tcattttaaggagatgcagaatgttgtggagaaatataaggacgatagcgacgaaaaactgagcgtcattcttaagagtaagaaatcccccca
aagaatttgacctctccttttccgattacgttgatcgccttaacaggattctggttggtgtaattaagagggtggccggagctattgaaagtaag
gaattgttgcagagtaacagcatgatcatgaacagtgttctgggtactgttgtgtccaacattccttacaacatgaagattaatatttgtgtgttttt
gactaactttatttgtacatttgctaatgatgatttgtacacattctttagggatgatgagaaatttgtaatgagtcaggtaacaagatacatttcaaa
ggattagaaaaataaaagatggtataaaattacttgtatttttattcaaacaaaacctttatatttgtacaattcccactttacttcttcttgatttcgtc
cttgatatcgatcacattcttggaggtgaatccaatgttcattgtagcttcgaaggctgcgccgtcttcaaatttgctgcacacgtcaatgagggt
aatttttgaagatgaagatttaatgtccttgctcgagaagtaagtgttcgacatagtatttcccttaatgtttcctgcaataacagttggccgaaca
gtaatgtcgttgatcaagagatcgacgccagtgttgttgaggatcttcaaagtcatgttagagttctcggccccagtgccagcgggatcagtc
cttggggctgtgacggtagagatgacaagatcagcagaaacatccttcatgtccttatcggagagaccattcttcatggccacctttccaaga
ggagtattgtaggacctctctccacgaattgacattacctttgcccttctttgaattgggactcgcatcatctgatcataattagcgacgacgcttc
ttccaacacgtgtgttgaatacaatcattataaccatgataacgattagagcaatgattgcaatggacaagattgcaataattgcaacgtccag
gtttgttaggttgccaaattccatttttctttgttttagatggaagttcttcctttacgtgttggttggatccagctagttagttcttctgtagcaccatat
acccagaaagggtttgtttttaaaatttaaccctacatcgaaccggatgaagtgtaccttgccgtgatctaatttcagtaacgatacctcccacta
cccctactttcatctccatctctaccccctccctactcctcccttcacctcctcctctccccctttactcgtattggtcgtcttcatcgtcgtcttctga
attggatgaagaattgtaatattttgtgtgcttcattcttttcacttgctttgctagagaattgacgctagatttgttattcaaaaggtccataaaagtc
ctccgtctagaaggtgtggaagattcttggtcaatatacacactgctagcgctactcatgttggaattgatgtcatttgaaaaagaaaacgttctt
ttcctaccaacatgagaagatacactgctaggcacgtccattgctgttgctgctgctggggtggagggagaaacatcttgatttttagaagga
cggtacattaagtttcgcataggcacaacagatgtagattttccgtgcttacccttttttattgtatttctcagatgcttgatttagcttatattttatgca
gtacacttcaatggacaatttagctttctttaaaagtttgatcactactgcgcttagcaagtgtttcctcttgtccatgaaattgactagtgtatttaat
gcattatcaaaaataggggatctgtgttcgatgattgatgagcctgcattgtatgctacaccaaaccttcccccgccttgtgtctggagatcaaa
catactattcttgctgttattgttactcactccagaatacgcgtcaccgaaaaagtggtttaatgatgtacggtcattactgtccccgtatcttatatt
atcgtatactcgacgaattgcttccttcttgtccttctctgaaaacatgttgccgtcttcacttcttttctctgcctcacttccattaccatcttcatcct
cttcatcgactgccttatcaaattgtatactgtttaagacgtaaatgggagggacagccattttagaagaacaactagaaatactatccaaaaga
agaactttataatcttttgacaaagtgaactcttttggccaatttaatagtacaattggtatgttcaagaaaattgttaaccttcccacgtccatacat
agtttgttgatgtgtttctctatgattgtagaaattgccccgaaacaaatttctgcccctatagaacctaaccccatttcttccattatgtttgcagatt
ctttactcatctgttttagtttagatgaggtagacaaatctgatacatcatttgtttcagtttgatgtttttcaaaatcctcttgggctttgttggtgtctc
cgaacagtgccactgatgccttggtcactatgtccgaaagcatattgactaatctagaccaagaaacagcacttgatgttttaaaggatatgag
gttgtttaggaacgccagtctttgaccaagcttatacgtttgacatggttgattatcgtcgtcttctttcagtatatttatcagttccatgtgcaaattt
gcttgagtttgattactataaaatgaagaatgctccaccacctctcctcctcctcctcctcctcctcctcctccagcaatagactcgtcgcctt
cagcacatactccaccagcagatgaagacatggaagtttcatcctcgtttaattcagatatactttcaacctttttaagagaaggagcaggagt
cgatggtcttattttgctagctctttcccccattatataagttgacgcgtatgctgctgaaaaaagagggttggacatgatttgcttcctctgaaac

aactccaatttctcatttgttttcctgaacagctcggcagatttaatgtctgtcaccgcagcagaatctaaaaatgccttcatggttaattctgcag
cgatgatatttttaaggcgctccattactttctcggatggttttactgggttttctggatcttggaacacaactttcaccagttcagagcattgaagg
ttctttgttgtttctagaagagtttttacccttgtttttaagattatagactccatagtcttgtgtaatctttcttcggcctctttcccacctaatgtgttgtt
aacctttccaaccactgtactcaaaaaggatggattgttgttgttgtttgaaaataaggactgatttgtaggtggaggtggagggggaggaag
aagaatattggtaggtttgagttgaggaggttgatgaggtggtggtggaggtggaggaggaggaggaggcatcagcatcgaatcttggtcc
atgatgtctctactttcttctcccatttcattgtctggaggaaaagtcacgatagatggaacaggcggtggtgctatatcgtttagttggtcatatt
gttcctcacctccactaccaagaatagaaaacattctaggtgcttttggtgaaggttcctttatctgggggagaatatctgtctctttcatgattgg
agggaacacaactgcatcctcgcactcttgtatatctggagggggagatcttcgaaccctgggcgcattgttaaattgatgcatagtacttattt
cttcatcctcgagtgtgtgccgagattcatcttcatattcttcatattcttcctcttcttcaatatattccgtgtgagattgaacagactctttagtttcta
catgctcttcagatttttgtgcaggttcctcctcctcctcttcctctccctcttcttcatcctcttcttcatcctcttcattatcattgccgccacctgcta
ttgaagagaagaaataatccattccatcatcattacctccttcactcatagggctggattcaactctatgagagtgcacgctttcattttctattgc
atcttcatccttctcgggttcatttttttcttgttctataggttctttttcttcctcttcctcctcctgttcttcttcctcttcctcttcctcctcctgttcttcctc
ttcttcatcctcttcctcttctacgggttctatttgttggttatcatcttctttgtttttctgttcttcacctgctgatccgcccatggcgttcacattttcaa
gtaaaaagttcaacatgggaagaacttgcctatttggccctactggcgagggggtgttcactatacttttccactggttttcaggtataattgtcg
catcttcttgttgtgccgctagcgccaataacgcagggtcagcctcatcttcttccccttcaccttcattgtcatcttcatattcatctccttcttcttc
ctcgtcttctagatttccgtcatattcacccccttcttctccctcttcatcttcctcttcctcctcttcctcttcttcttcatcatcatcatcgtacatgtcat
tagtgacatctctcacaaatttactttgtttttttagaagatggaggatctttatttcttccatttctaactcccgtgtcctcatcttcatcttcgtaatcct
cctcctcctcctcctcctcatcctcgttcatttcctcctcatcctcttcgtcttcatacaagttactataatcaacattattattgatatcgtcctcgttgt
cgatatccaacactcccgcaccttcagttatacgcagatttgatgccgcttccatgatgaaattttagtatacaatattgtgttaaccgtggaaaat
caaacactacatttactaagttcctttcctgaaaaaacgagttacctctatagggtaattctcagggtgtatattctatagtttggtcttgaacgtgc
ttttggtttttgttataactttctctaaaatctactaacggcaagagcattgacagttgagtgggaaggggtggttcagagtaagaccaaaaataa
aatattttttgattgtgtgaatttcgaaccgtttatattactattttagggtactagaaaacgttgtcagaaccaatcatgaggtttatcccgatgttga
taattgcgcttatcgctgcatttgttattgcagctcttcttactgccaattcaaattacctggatcacaatatcaataaggaactaaatttgacccgtt
cccttcaacttcggggaacattcacaccagaagatatagctcataacaacagaattctcccttccaagctgagtgttttagaaagagggtcaat
cattttagcagaaatggacaagtacaaaaatgcacaacctacagtaaataattctcaagagaggaggaacatttctactcctcaacaacaaca
acagaccaccccctcatcccaatcatcttcccaagttgaactttaaattctataagaaaatgctgatttatgttcctgtatgtgacgagtctcgcac
cgacgagaaagaagattttaacaacgatgaggaagaaaatatattaaaggaggaagaagttagtggaggagaaaaggttgtaaatatcgta
gtagggagtattaagatgatgaaacctggttcaaaaatcaaaacaaagtacaggctctactccctccttgcaagtgcgatagggagtagaaa
gaaagctgaagaatatattgaacgtctatacaattccttttcttccatgacgttgaatgaaggaggacgtctggcttcagcgatttattgtccatc
ctacaacaataaaaggataaaaaacaatcgttctaggcctgtgaagctgatccatgcctccagggaattgttatctgaaactacggtgaggga
ggaaatactacggaaatctcccgcttcttcttcttcttcttcaacgtcatcatccacttcttcgttttcttcaatttttttgtttgtaccttctaattgtactt
ctaaaactgtatgcgattttgtaaaacacatacagtatgaagaagatattaaccgtttaaggtataacataatacatatatctgaagaaaactatg
cttcaagattttctaaaattaatagattaactacatgtatacaaggtataagtaagacaataaagattctataaaaatcaacatttatttgtatcattta
taaaattccatatttttgtatccatttcgtagtcgacgggtagagatggtctgtagaaagaagatgctgctccttttaaaaggttattggacatggc
tatcacatacattttcttattgttttccctattgatgatatttgtcatgacgttgggagtggcatttttttgttgctggtgtatgacaaacggcaacattac
tgttgttcttaaccacctgagcaaacatggctgtaggggattatctttctatgtcctctgtaggggaagcaacgctcgtgggttttatgattttaaa
ttttataaactttgtcacaatattatctttaataatatatgccgtgacagacgtttatagaagatgcaagcggccgtctaccaatggatattctggtt
gtacaaccaatgtggtttcttcgactttacaagaagctaatctcgttaccactgaaaaagataaacctgttcaatttgttagaggtttagtcccca
gaaaaatgatggaaaaatatagatcggacttgtctcctaagaacgtgggggaatatattttaccttcagaaaaagaaacagacaaattgaaaa
gtgattataaaaagggtaagaaggttggtcttttaactgccctgagtaatggtcatgacagtaacaagaggattatagggccaagggatctga
ttagtagagatgatgtgaaggacaaaagttatgtctttaagagattgagcaaagatccgctcgtctactactcttctgcaacctctaaatacgtta
gaaaattttcccctttcagagcaaaaaaattcatgacatcaacacagttggggagtaagctcgtgtatcctcaccctatacggtatggtactgct
tttgtactacccacgggatacgtgatcaacaaagcatacggaatggataatgaggatctacacacttggaacccaccctcttcttctgtgctcg

tcccagactctaataatgatagattaacagtagaatgtgctaaaacggacccaacacatagaatcggcatctacggctttggagggtctgatg
ataatagacgtgcaaaagaagaaggttatgtagaaatgttgttatgtaattgtgacaaccacaaggacttgcttaaggctcctctaattacaga
gtattctacaaatccaactgaaattcaagtagatgttgctgcaaaacgtgttttattccctgcccctggttccgagcctgtaaaatcttcccaagt
gacatctgctgctcatcaactagacggagctactggcgagcacgatatttcccatgagcccgtgaagctatcagatacgggtgactatgcag
ttggatcacccattgtattcaagccagtttatggtacatctttagtaaatcttccagaaacaggatctcctctggcattgaactgcccctgcaccg
acaaggctgatggaatatatcaagtcaatcaaaagggagggatattatatagagatatggtggggtatcttaacgccaaccctgtggaagct
gcatcactttcttcctcggactcttcttcgtggttgacaactggtaacaaaatatcttctgttacatgtgaaggagaaaaaataaagaaaattgtgt
aatcatatgtgtagttttattaccactaaatttattggtcttcttggtactgtttacactagggaattttactcttcctccatctcaaaaacttttcaaaaa
tttttctgggtcgctcgagtttagagggtggaccgctgggtcggtctgatgtcaagtttggagggtggaccgctgggtcgagccaatgtcaga
ttgcaccagaaatgacagagctccagaaacgtacattaactttatactgtttctggaacatgtatttctggagactgacacgtccagaaagggt
agggcatagtctactgttttcactagggaattttactctgcctccaccttaaaaacttttcaaaaatttttctgggtcgctcgagtttagagggtgg
accgctgggtcggtctaatgtcaagttcggagggtggaccgctgggtcgagccaatgtcagattacaccagaaacaacagagctccagaa
acgtacattaactctaaactgtttctggaacatgtatttctggacacggacgcacccagaaagggaaggctataacacactaaatacactaga
gaattttactcttcctccatctcaaaaacttttcaaaaatttttctgggtcgctcgagtttagagggtggaccgctgggtcggtctgatgtcaagtt
cggagggtggacctctgggtcgacccgatgtcagattacaccagaaatagcagagctacagaaacgtacattaacttttacgtgtttctggaa
tagacgtttctggtcatggacgcacccagaaagggaagggtgtaacctactgttttcgctagagaattttactcttcctccatctcaaaaactttt
caaaaatttttctgggtcgctcgagtttagggggtggaccgctcactcggtctaatgtcaagttcggagggtggaccgctgggtcgagccaa
tgtcagattacaccagaaacgtttgttgctccagaaacgataacacacatttctggagcagtagttagttatttctggagcacgttaaaataagt
agggagaataatacataacattatattttcaattccattttattggttctatccatctaaaggtgttagaggaaatactttcattgtagtataagaatc
cttactgaaatttttcaaatcatccaaagagaatacaatctttccagtataacttagagggttccttggtgtagctgttttcgtgtccctggcgaga
cggtacttattattgcttcccatctttaaatcttcgcttctactcctcttcatagacctgaataaggctatggatgaaataggggtctttcctacaggt
tcgacaataagtcttcctatagatgcgttccgtatcaagtcttgcttggccttcttcacattcctgtcggatgcggctttactcccactagttttccat
atctctgattccgtttgcatctctctaaatgctttccagatcaatttttctctgttaacattttccactttgacataattattcttcaacatgttatccacta
acatcttggatgaaaatgcagataagagaggtaagggcctccctggtacaaaactagtgtttgacgcatcaatcacatttcttcgtgctctaac
agcattccttaaagcccaaaagcttgcaaaggataataatgatgatgtattttcatctgttggaacatttccccataagttatgtgtaggaaacttc
aaatctccccatcttgccaatagtaaatatactgctctgtcgaaattgttcgttattgaaggattggaggtggctgcagccgcgtacaacaggta
gaaagaaagggcagcaaaccctttagatacatggacatattttgcagcttcaaacacataccctttcttatccactaggtgcatgttacgcactt
cttcgtcctctctttttttgtttcatttttccttcaaaacctatatcaccacctaaccctccccctcttaatagacccgaatctgcagagagtatctcaa
ccatcaactcccctaacccattgacattttctgctcctttattattaccaccaccaccacctaaagcatcgagtactgctttccagtacgtctctag
ttcctgttctcccatagacggtttactttgttcctttaaactgatggtaaaagtgtctagagattttgttatgaaaccagattcacaacgctggcaga
cgaataaagaatgcatatctttcaaaaacacttgaagaattaacctgcttttcactgcagagctattagttttctgtgttagccctccagagcataa
aaataatgcattccatagcacatacattgaaaatcgggcagctgagtcgggatcaagaacgcgggtaacagagtagttggagaacataactt
gtctattattgacaagaaaatactccaaagtttctggtgaaacaatatcgttaaacttgttacatctcttgtattcacgttgaagagcttgaagggc
ggtaggtgtgcgtttaggccctaaaattccgtctatcgcctttggaagatatgggtggtccctaaagagattctcagcagtcatatccgtagtttt
aattcgttctctttccctctcgacccattctttttccctttctgtgacactttcggatggcatctgcagatggggattgttcacatatttatccagtttag
ttgtcaaaggatttggtttagcgtttctggtcctggacagcactgccctgtactctgcgtcgatggctgctgataatagtaccatgatttccctcc
ctaaaagagcttcatggaaagaacctaccttgggggggtagaatccttcctgtattacagaaaccatgttggttaaaacagtcgatatagttgg
aacaacataagattgtccatcttcttcactgactgtaactccatcactaaaggacccttcaataaagtcttttatcctttttttcaatcatgtcagcag
cgcgtccgtactctaggtgaggaggcatggcagcaaagggatctaaacccctctctttcattctagaggcgatagattgcttccatcttagaat
atcacaaagagtatatcctagaaatagagcaatggctctttttgctcgtcccattgccagatcggccaccgttgaattgtttctgttggccaacat
ttctgaatcaattacccatgaaggttcattagtatcacctctagtggaataaagaactagattagctgctgatctaaacacaccgtccataatatt
ctcaaaggcattcaaaaggttcaataatggtgcttgtttcaaaaaaagtaaagtcacatagttcttccgctgcagtttcggtcaatctccaagagg
ggtcaattatatgctctctaacaccttctatatctctgtagacactcctcatgacggcatctttagataggtgtcctttggctaatcttagtcccgaa

ttaaaaaattctgtcgttttattgaaccacgtgggtccgaatacgtcagaagaaataacaaccttcctttgtttttcaggtgtattttgtaaatttgaa
atggcagcaactagcccttccgatacataatttagtcgtttttcatcggttttagtttctgctactaatgccagaacatagtcctttgactctgtcatg
atttttgtggtgctggtaaagtagtttttgtacacgctgcaatatcttcgaccttaaatgtcttcggttaaagggatatttatttgttggtactgttttcg
tcaaggctgaaaaataatttcaaatatacaatggagtacatggaagaaggagacatcgctgaaaggcgctcagaaggtgtcgactatattct
ggacgaaaactctgcttgtgtagttaatgtgaagagtatccgtaacaggctcggtgccatggacgccgaggaggcacagtacgcacagga
catttccgcccaacttgtcacccatattatccgtctggcccactgctccgaatccaacaagattaaggataccattgccagtattgcgggtctttt
catcaacaacatctttgacaacaattcaacaaagaacaaacttaaaacgtataatcaattcaaggcagagtcacaaaacaagtctagcgttctc
aatatctttggctctctagatcctctgagtatgctttctagcttcatgggttctgatccagcaaagagtggaggggaaaatttggacaaatctttg
ggtgtgctctttgaggtgcttcaaaattacaacccttgcaagattgacgatattgtccttctggaaatgtgcccatccaagtgcgccgcctgcac
cggtctcaaggaagccatccgccaggaacaacccatggaagcaatgttgttgtttttcaaatgtatcaaccataataggttcaattttggaagc
gacataaagtcagcatacgcctctgaaacatgcatgagatactctcaggacgaacgcgcagtcgttgtgcctctgaggagtatcctcctcgg
ctgcctcgacagggatgatccagctcatactctttcctctttcggggatactatcgagtatgctgattcagataacgcttgggtttcttccctgttt
gcagccgtctctagaatgcctatggtagacagagccgttattgctcacttttacgtgtacacaatgctcagccgacataggcgagtatctgga
gacagcttcaagcagtttgtctataccgtatttgttcgtatgatttactctgcgattgaaattttgttctgtgatactgaaaattcgtctgtagaatgtg
atggaaagcactttttgagctatgttaatgccatggttaacgtatccgtgctgggttctacgtttaacgtactaaaagcctaccgttcatgggtgg
tggatcaagcatccgtcgcaccggttctagacattatttccggaggatggaagaagaactacccctcacctgaccacatcaagagggtggc
gtacgacatctctcaagtcatcaatcatcttgcatcaccttctagaatggttaaaggtaacaacaaggctagcaacgttacatctggcctggata
gtatcaggtctgttcgtcaagcagaaaaatatatcccgtttggaatacttgaaaataaagcagggtatggtgtaataaacattgccaagcacaa
catcagccgtccagcaagagaacaatccaacggaaggaactttaattgcaacgctttacacattctaccttcaattaagggctgtgaggcact
tggggcacaaaaggggagcgcagatcaaactgtcaatgtttttgataattttgtcgcatctcatatggatattgccatgaaaaagcaggggtc
ggggaagattcttggactgctcactagcatgattgacaggcaaggtctgactacttcattccctagtagtgaagcggaatacaagaagagaa
tccatgatttcacaagatacgtcatcttctcttcaacacccatcaacgacgaactagtcaattctcgctgtattcttccccattctaatgttctgaac
tcccctatcagcttgagaaatattgacccagaatcagtccccgacactcgattccactttctgctcatgatgtggcagcggccaaatatcgatg
aacctaatctttctgctctaactacaagtcagctagaattgttgcttagcaagaaccaaaaaatgggacaaactcaccaccagagcgttcttcaat
atcgacaggatcaatttccagatggcagacgctatcattaagaacgtttctggaagcggcttcctagatgggagtaaaactgcctcttcttcttc
ctcagcgcctaactttttccaaatcttcagtggtgctgaatgcactgcaaagcagctccaaagtattcgcaaattcattggagaatctatgcagc
atgtacaaaaggaatggagtagtgcagtaaacaatgggaacagaggagtagaaaattatgacggactcaatgctcagttctctgaagaact
gttcgagctgctctacaaattgatcatcgaggaggatatgcggccatccagcctgatcgcctcatctgaattcttgagcaactacgtcaacgc
catggatgaacttcttatcagagctaatgcttcttagatagggtttttatgaaataaaacaaatataaattatatatcatgcaattttatttatacctttttt
ttgtatgtgatacaatatttttagtgttatgacagtgcaaccattatattattatctgtaataggtcttggtaaatttttaatgaaaggaagattaggtg
cacaatctattgtcataaaaatgtccaggtcaatttcttcaacaccttcctcaaagttatagaagaaacttgtttcatcatttttattagcaataaaca
agtacgtttttttccttagctgtgcttgcagagcttatccaatagtctgatgcctttttaacattataacagttggggagtaaactacaaaaacagtc
ctctgttttgtttgtagcgtatacccgtacacggttaccttctactacaaatgaaaactctatccgggctttttcaatacgtcctcccacagctgctc
taccttgtttttgtagcacgggttctaatccaacttgtagagaatccagattcgttgctgcactaataacattaaatgtcctctgtttcatgttgacat
gtacagacattcgtccttgccaagtgcgcatatcacgatcactgataaaacctcctcgtatattatttatcaccaattcctttgagtaccgggcaa
ggtaatcaggtgtaggttgaggaggttgttgttgtctagctagttgtgaagaaggtgttgttgtcaacatcaccatttctgcgtctgggtctgttg
atctctgtaggctttcatccccattttcatcagaaggatcaactgttactgtaccaaacaataaaatattggtcaagtctattacgagttgttcgtaa
gtgatattggtagatttttgtgcaacaacttcatccatcgtttccttgacttcgtcatggttcgtccctatgagatcgaacaagttttcatcagcaatt
gcctgtccgacatcggggacgtttatgtcaccttcagggattgaaacttttccatctgttattttatcctttattaaccatacgttttctgatttgtatgt
gtcaaattcctgatctgctaatttggtgaagaattcatcctcagctttttcaagagttcttgcaaaatgctctctcgtttgacgttgaagcgctgtct
cattgataacatcaactgttcctccccctcctcctccattattattattatcgtcgtcagtttttttcatcatcgtcagtgtatttagagtctaatttatattt
cttactcttttttgttttggcatttgaatgtagagttcccttccctacgcccgagatagcaaagaagagggcaatagaagctcctattaccaaaac
aataaccgcaaaaacaatgtatattcctccattgtttgaaagtgccatttttaaacccttcaattcttatctaaaaaacaaacagattgaaatctatg

tataaatcatcactcgcgttttcgatttcaggcatgtgaattgtaatgagagaggccgaatcctcggatgggttgattatgccatttgtatcatga
ctaatatatataggatcatttctctttagataatcacctatgttatcagaagaatcactatttcttagatttgataatatcactacactttttggtctgttt
acaataccagcgaacgtggatttgcggttaaattctaacgtgagagttttggctgaagaatctattttgaacgataaacttaaatcaattaaatcc
ccttgagcagctccaggggggagtgattgataacatgtgtttatcgtttattggtacactaaatgaagttgaagagtcggtaaaggcaatagtatt
gtacttgaaagaaaatgatgtatgataaaaaccaggtatatatcttctccaccgctcctcttttaaaatttcagttatgaaatcttccctgggttcca
tgatccgtctattcacatcgtatactttttccataaccactggtattcctttctcaaaaattgcttcacagataacaggattgaaagcgtcttcgtaa
agatgttcaaaagtcattacattagttaaatggctattgatttcttgccttacttgatcctctatatattttatagaggcatttttgctaccgaataggc
aattgcacctaagcgatactttaattggtcactctttcttgtgatccaagaattgtcatggtgagcagctcccaattgagcctccactgaagcgat
ttcagcttcagtcacagcaggctcatctccgttagacgacattatttactcctccttaaaagcagtgattgcaaacaatatatcgtcccttgtatcg
cctctttttatacgtttaacagcgtatgttgcagattctccgtttagtatatgggtgtgtttatatctagatttacatacaatatcaatatctttactcttc
ccttcaacaatcacattaaagaatatgactacgcgttctcctgttcgattaaatgcttcaaagcttagcacgtcattaactacgtcactacgagac
acaagaatcactgcatctacatcctcatcatttatggaacgaaacaattttgccgttattttaaaatctcccaagggaatggagaaatcgtacctt
gtaaattctccatcgtcaccaccaattctcaaattgtacgtagcaacactacaccgccctacttgaccagtttcttgttgaatttttcgtctatatct
gtgcaagatattgggggccgataccaatgtacaagattttgcgttagttttcgtttagccaagtaatcagttattatttgtctaacatctattccttgt
ttttcaattgcctttaaaacacaatcctttttatgtcttctaggcgagatctcacatccagttctgcatccttcatgtaaactctgtctccatgtcttac
agtgttcttcaacaattcgtttatgtgtttggttatggggtgtatcaacaattcagatagattagaaacaccacgtctttctacaattttatcaaggat
gaggatcaacctctcatcgcttatagttgcgccattagacatgctgttgctacttgcttttggtatacaacatcaacaggtgtaaaaaacatacaa
aagaaataataaaaatactttaggggaagaaaatgtttagaagtattttcaacctttattgttctggggcgaagattatatacattcaatcttttaa
cgtagacctttcttcgtcttcttcttactcctacttctttttctcctcctttctggtcctggaactggaactggggctcggtaactggagaagttgtgtt
ggtggtagtgacagatattacctcctctatttttggtatcctgagtacaatttttggttttggttttaattgtactgtagagggagttttctgtgttaccc
ttatagcctgtttcaggaacaagggtaggtgtctatacccaagggtagccattgctcaagtgatgctccaccccgtgcttttccccccttatatac
aatttctcacgtggtcaataaaggaaacacaaaaaatattcatcaattcttttatttgatcaaagaacatgttttacacagtttgggcacagataat
cagggttaagtctcaaaccccattggttcattacaatgtacttgacaatatcagaatggtcagatttttgagaagcattaggaagagctaaattg
gtagagtgtctaagagcatgcaacggtttattcacaaccattgccagaaacatgggaacagatagtgaagaaacttgattctcatccgcaata
tcctctgtatttgccctcttgtgggatgaagaggacaaaaacacagggtatttcagattactaaccatcaattggtacaccatacaggctaacat
tacagcaacataatcaacaccagttttatcactctttagtgtttttaatgaggaaagtctcctcttacattcagtatactggagagccatttcaaaca
acatgtgtaaatggtggcaagaatacatcaaatccctaacatccattgaaccccaggcatcacacaaatgttttgttagtgaagaaccaactcc
tattaattccccttccttatcatatcctacaacatccatatcaggatttggtaaactcataccaatacaattgtcaaaattgatggctggcctattcttt
gtggcaggagagtgcatcttgatgttaaaatctctcccaaagaatcttgtggtcttaacaaagggcatttctacaaataaaactgaattgagcat
gctatttagttgagcccatgcgccttgtctagaacgagcaatattcatagccattgtattcaaagatttaggattaatctggcacatttgaagagg
tgatggtaatttattgcacatggactcgtaatttgctggtttcttacggcgcaattgttgttcctcagcatcacaactattgtcttcttctacttcagtg
atagtagcagaagaagaagaagattctccgtaatttgttgcagccttagcacaaagactcattgaagtgtacaaatggaaaaaggtgttagtat
tctcctctaaacgggaaactgattttacgtctctcaattctaggacctggtttctgatcaagtttttcattaaaactgatggaagaacggatgatac
gtcactaattctatccttctttttgttgtcactggaatgcccactaaagtcattccatgctttcttgtgatgattaaattcgtgttgacgctgttttattcc
atgctttccaaatttgatacatctttccatcattaaagatgaatggatatacttcattcctctatccaaaagcccgataattttcttaacatcctcattct
tattctcaaagacaaatgttgtatcacaaggaaaaacaataagtcttttagacatagcttcgtcgacaaagaatgagaagatgtaactgttgga
catcataataggagacacagtaatgctgctttttgcgccctgttggaaaatgcctctaatcatgacgctgttttggggtgaagtcatattctttaat
aaactggagggaaattggcccggaagcatatgtttggttatagaactggacacatcattcttttgggggatggccttagcgatttcggtaataa
tggaagaagatccgtcactgttatgtttttggagtgcagacaccaattgccttcttttaggcgtatcgacattgaaacaagcctctttattactgtct
cctccaccactagagttagaagcaaaggaaaattgtgaactttgttgcctattttctccaaaaacacttaaaaagtttgcaatatcactgctgttat
cgtctgttttaggtaaagaagacgtattttttgatgaattagaggaagagtcgtccttttcgtcagaaagcaaatttttatccatacacatgcgaat
aaaggatccagaattcttagccgacatcttatccttctcaatacgcttctttaagcaagaaaggacattttcgcagtcagaagtggaagttgaca
gaaccactgaaccatcagaagtaaatccaggaaaactatattttgacggtttcttaaccacatcgaataaagatgcgtcataacgttcgtacat

gttcataaagtccagatctctatccactaatagttttgcagcttctgggtcctttattatggacgataacttgcgagaaatagtcttgatgttacgaa
tactgtcaatattaatattcgacagagtagggcccagaagagccttctcggcagcatcatcgacagctacaaagtgttgtttcttgggagggg
gaagtaaaactgggtagtttgtttttgaaagatcatcagaaatttttccagtgtaaacgcaatggtacgcaatcaatacaccaggggcggatgt
attaccatcaataactgactttttaacgcccccgttagtcgacaggcaattatcccccaggtatttttttaatttcgagcatgtcctgggcagacaa
agatgacgtgatcatctcggaaaacctttcagtggcagggttgcagactaaaacaggcgaaccgtcgcgatcaaatgatgtattccatttcaa
aaggccattgtttttctgtattatgttcataatcgcatcaactttgtcttcaattttcatattctgcttatttttcttgttcttgctcataaatttctcattccat
tcatctatatcttctttacaatcgttccaggaatgaagattattaacctggcagttttcatcaataatggattcaaatgattcgttatcgcatattttgg
atccaaatttacctaatatgcacttcaaggcagtttttacttcggatttagagaatttgccttcatattcgataggtttcttacagtgcctgtacacag
catcggtttcgttaagccaaggagaatctgtgcagtctttaatagagaaaggcatggcgcctctatggagcatattatacacaatatttttctcttt
aaaggtagttgaaacaacaccagaagaacaattccaccagtgagcttttctggcaaagaaagaaatattgataggatcctttatgcggacac
cattttcgtcatttttaccgtcgtttctctcaatttcctcacattctaccccttcatcttcaatttcgccacctcttttacgtttttttgtatcattacaagca
tttttcctattcttttcgtgtttgagttcagaaataaaatcgtactcttggtcgcatacttttagttttgattcggtcattaaacggttaatattactaaaa
acgatcttagagtttcctccgtcaccgtacgcgtttcttgcttcattgatgtcaacactattcatttctcccccactgccgccaaaatcgacattac
cacttctggaagcagactcgtccttatcgtcttcgctcgatgacttctttgagccaaatgaaccttggtggccaacttcgtcaataatcacgatac
cctttttgccagcaaaggaagccaaatttgcttgagtagatgaagaagaatccccagcattattgagagccgtaacgttaatatttgtaaacctt
tcgaaccaagtcctcttacacaaatcgacaagagtggtctttaaattgtttccctttgttcccacaaatacaacagcagtttttgtggcaatattca
aggcaacggaacaaagagtttgaaccatgtggtgggctttctctccatctcctgaagcaatggctgtgcacatacctccaaagttcattaggc
cacgtaaaagatacacgacatccattctgggcatgccagtttttttggtcaacacatgagatgtaacattttcttagcgcatgaagggctgtagtt
gtagaattaacatcatccttttgcttcctcttcttattactactactactactactactatgttttgaagaagaagaggaggaggaggaggaggag
gaggaagaagaagccgtttcacaatcgctagtggctttagcaagttcgtataaaattgataaaaatatataatcttcatcagcaataaaaattga
tggaggttttagagaagaactgaatgctgtttttgatacaggttcgtactttacaacgctagtattttccgtgttttcttcgttctcatcctcatcttctt
cattttcgtcatcatcatcctcattttcatcggtttcttcctgatcttcagccatagaccaaaagtttttgttcatattactgacagagtcggtatttttc
atataagatacgtgttcgattaaacaaagtcctctacccattgtagtgccttccatggacgaagctagagcagcagaaataaattcggttgaga
ataaagaattcaaaataacgtgtttttctgcctttaatttacaccttgtatacatgtccgtgaccaaatttatcccttcaaggcgtcctcctagaaga
taagtaccatcaatcaaaaagtacctgttattatcaacgctaatacaggcagtgttttcagctaacctgaagggaattgctgtaccattactaata
aacgttctagcatttcccatttttttcaacttcagccataattttagaaatgagagcagttcctgcatttgtaatacaaagacgtatatttttggctctttt
gagcatattcttggcaggacacactggattagctaaaaatgagcatttgtatgcttgttccgaaaaaatattctgcttgttgtagtctgacaaaaa
atcgggacagtatctatcgtggaacaactcgccttgttctgactcattgtcaaacattgaaggagttaatcgctcgctgaaacatcccccgtttc
taatttggagagtagactcgatcgccttcttgttattcttaccctttgatgatgaaacaatacaggcccaatcagtaaagaatatgcgggcaaat
actcctcctgtgcctttgttcaataagaaatcgcaacacttcttaaagatttttgttcctctaggagcgacaaatttaccctgcacaggggagtta
aagaaggaattatccaagtagaaacccttatctgttcctgctaaaaagttaaaattatccttcttaatggccagactagattcgtccagacttttcc
cattaatgacagaccgcatagaaattaacatcctcttagcatccattgaatcttcttgcagaaggcaagttttatctgtaccaccttgatcgtcatc
atcatcttcatcgtctgcgcagccagtatcaatgccagcatatagaccagaatggtccttattgtaattgatgacacgaagcacgcgcttatgg
ccatcagagttggtacttaaaaacacggcagttttaaggatactgacaatttcaatttctgatgaaacaaagttgggacatttcttagtttcttcttt
accaatgaaacagtaatcattgtcttgttgccgataacaagatttgttgaaaaaatctcctcctcgtcttatatttgtccctggtgtaaacaacact
gaacaactattagtgatccaagacgtacagaaattaccctcgggatctaaacatgaagacaaaacttcacgggcgtaaacagccttagccaa
actagctgcagaaatttttttcgggagaaggatcagaggaattattgtctttagatttagcagagaaagtgcggtgcagattcataagagggaa
agtgtccttcagtagttggccatagtcttccacactcaggaaaccagaatggtatactgaaagtggagggtttgaatcggtaacgccacatgg
agcattattctttgtcatttcaaagtggctattctttgaactattgggatgaatacatccccagttacacaaaacgctaatgtgacgctcgtacaatt
taatccacaaaatgtttagatttttaaacagcaattttttaagcaaaactatctggttcttaggggacataccattgttacagttccacatcttaccca
aacttttataggtgttgatattggatatttttttggtgggtaaagtattgctgggagaaagtctggtcaaatgaagattggttagaagagcccaaga
taacgtacgcgtctttagaatcgttaacaatcttaatgttttctaatttttcagtggaagtagttagagaatcatcataaagagacaataaatcttcat
cataacccataggaacattctttaaagagtctttttctccaaatcttgaattgaaacatatttcaccgatgaattactactgctattagagttattgtt

aaagttcttccagggtctctttccaaacttcttttttgacttgtcctcatctcccagggccatattcttaatgtaacaaattggcataacttgagacca
gtacgagtcagaaatatgattggagaaatagcgtgtcaggacctttgtttcagcaaaacctttccccttgtactggataaaagttctcaaggga
cggccaactattctaccctcatggcctgtatttgccaaaacactgccctttccttgcttcttttccgaagaattgacaactttcttgtagaagttaaa
ccaaaacgagcaatcacttccttcatgtttcaattggtcttcagaaatttcattggtctggttagcttcattattattattagagagaaggtgacctg
atgaggatgatccacccccacacccaccatttcctccctcgccaacagaagacatgtaatcgccaggacatatgatgttcatatcataaacgc
ttccagtatcagtcaaatcgcggcactcttttcttatcttgttagtttcttcctcaactgccttattccacctttcaatggcatccattccagcacttaa
cgcagtttcagtgacaccgtacaaagcatactctttttttcgagaaaggaacacaaatcttaaaaaagttctcaatgtcagaaattactttcacaat
gtcattatttatctgattttgtgtgatcatgttcttaattcggttaagtacatagccatcgtacgaaaacaagccactttctctgtcttcttcgtttctgg
cgtccatgtgtatctggctgaaacccgtgcgggatctcgagtcctcatcaactacggatgatgatccccgactagaatcatcattgtaaaaac
cacccgcgttattaaaacctcctccgccagatttctttgatcgtcccctattacttccaccctttcctgtcatttgttttgcaccactattatcctcctt
gttaatctcttctagagaaggatcgataaatgcttcttcgttagcgtccattactttagcttgaccggcaggaatgttgtggaaagaaatgttagt
gttgatgattctagaggacttttctacagcagcgagttgggcagttgtacaggaattactaaccagattgtagttagtatcgattctagtctcttgc
agttcttggcgagtcttggcgcggaactcgtccaaagatttaacactgacatcattaacgtccaaaatacaacaatccaacaaatcatttgaag
aagcacgcacagagtctgccgcttttactaaaacaagccttgcaatacgtttcattaccttcccaacacgtttttcatacatatcattgctttttttcg
ctatttttgtcggccaaagaggaaacactaaccaaattcttaactagtgaaaggggcctagaaagaagcctgcttcgccgtcccattaggacg
tgagaaaatctataggtgataaattgcaaccaattactagtatcaaatccattaaaatcaaacgacatgagaatctttcgggtcatttcttcaatg
acaatatcattacttgcaacttgtccaataacatatgcggcaaaacggtaaaagtggagaggatcttcttcaatatcgtaaaatctaccttctttta
tgctagcgtcggccttttttgggagaatctcccttgcctgtacagctgcaccacttgctagtgattttaaagcatcgaatgagtggtaccattttatc
cacagatagacgcaaaggagcagttgcaatttctcgcacactttctccctgcatcatgatacgatacatagcgaccattgctagcacacaaga
gagggacatttctgtcttgagaacctcttctgttgcagccatccactctgcccgactgcgcaaagtagacttgtttcgtggataatattcgtcttc
aacccaaatttgatgcctaacatcattgctaggggtgggttttatcttcaatttcttaattacgttgttatttgttagcaagttgcttgaatttatgaaat
tttcaacagatctaatgtcgttgtctttgagagcaaactcataggccttggcgacctgcgctttccagtccttattgtcgtcattcaaaagacattc
ccacgtatcagcatccaaatttgtgtccagatctttaccaaaaaatgccctcgcgctaacgcccagaaattctgatagctcaaatacggcgcc
agattcgtccacatttgacacacctgcctttaaatttagcatgtcactaccgtcccaatccttaactaaacttccccaatcaataggaaggctgtt
attaaagcactccctaacagtctctagaattgtccacagtctagaagctggactattggttccagaaccgtatcttgttatcaaatttgaaaagaa
ctcagaatcattacttgtttttccattcaaagctgtacgcatgatacattcttccatcattttaaagcactcgtatatgttaagttttgaaacgttatggt
ccttgaaataatttgctgcatctaaacacatctgttgacgtttttctttaagttccttctcagaaagcgattgattaaacactttagttggttttagagg
atcaacaactgtagaacccagcacaacagactctaaatcttgattcgcacacaatgcccaaataccagaaaagaaaccattattaacactctt
atctgatacaccatcacctcccgatgaacatgccaatctggctaaacttctaggtctaaacgtgttaatgagtttgttatcttcactgatcacatttt
catagatacccaaaggtaatgctgaagatagacattttcctgccaatttacactgttttatggcatcccacctcttattatctaaaatagacctaat
attaccattaactttgtcctttttcacaaatcctaaagctgcagtaacatcactaaagtatctcctttcacatgccattgtgttgaaacggagagcta
cagtcagatctttcagatgtttagaactaggaattgggagtagttttatgcacttttcgttagacatggtattgttactactaaggggagaatcgct
cttgaagttattaagagaagcacaagaacaaaagggagggatattgttgttgctgttagcggttccgtcttcactattgacattaggcttgacaa
tttctcgtacaggcatagtaaaaggcactacccaattcctcattctcaaagatcttctattaatttctttcaaaatatcaccagattcagctgcagaa
ttgtccaatgttagtacatcacacaaatttttctccatttcctccacaatgacagttgaaacatacttttggcgatgacataaactggatctaaagtt
gaatgttttgatgaaattaacatccctaaacacatcatcttgtacgttattaatactactagcattgtcagatggtacattttcatcatcactagaacc
aaattcgcctagaaaaccataacaatcatcttcatcatcaccaccataataatcacctgttgcacctcccttggcagctccataattctcctcatc
gtcactctcttcatcactttcttcatcgctttcgttactggccacattgttcaggatctggcgagcagaaacgtcgtcgttgtcacattttcttagata
ttcttgtttttgttgatagagtttctccctcttctccgacttcttgtctgccaaaacagcttccttatcagacttgacgctggcaaaaagtcgttggaa
agtctccatgggggcgcattcatacattggtccaaaagacttgtcagattcatcataatcaaacgttttaaatgtatcgttgttcaattcgtagacc
attgtgctttcatcatacttaattttgcacaatttgctgtttgtcttgttcttgatcctctccatggaattttcgtatgctgcaactcccattctgcagaca
gattgcttttcaagcacacttttgggtacatatatgctaagttctcggttaattgcgtcgcacatgtgtctaaacttctcaaaacctccttcaaattca
aaatcttcacgaattatttgctgatagtactcttctggatgagaagtgatgttgtacagcgcacaaaaccgctcagcaagagtaacgactgtcc

acgcgaatatggtgtcgttcttgtcagcacccgtggcgaatgttgacgtgatgatggacaagagcttcttgatattttccaaagaagctatctgg
ttttcatcagatgacgaagtggatgttgcaccttcttctagactagaaaccatatctggtgttgttgatgatgttgatgcagcagcagcagcagc
agcttcttcttcactttgagtaggtggaggaggaatactctccaattcttcaattgtaggaggttttagtgattctttgtgggtaaattgccacatttc
tgcaaatttgccctttgatagtgcatcccatgtgttattcttgagtctttgttgtagaatgtccgtatctctgggcaatttcaggtttctgttctcatatc
tcacaacaatttgaaggatggcgaaaggcatgattgtgtcaaacttgatggtgttaacgccagtcaaattcctgataatgttctttgtttctggttt
cacgcgttcaacacaattcttcttgctcatggctttctggggaagaagaaactccattgccagaagaggagagttgtcaatgtcactcagaaa
gaaaccgtgttttctcttgaagaaacacacaatctggtcaagcttatcatcgtcattgatgatgctattatagacattggcagcaaggggaggg
atttgtttaggggctggaggaggcttgttgatgttgttctcgtctccagaaccgccgccagatttccctccagatcctccttcctcattatcttcttc
ctcgtcgtctccatctccagaatcgtccaatcctttacacgccatcttctcctttaatagggtagcatttgtcttggtcaataacaagccacctcctt
ctgccagaaacttgtaacacatgccaattgtacagaaatcttgatccataatcaaacttttgtttctgtaaggcatttctggcccaagttcagaaa
aaaggatctttcttctccaatcgtcggtattggccatagaacccatgaggttttcgagatatagacactttttctcgaaactttccagtgttttgcac
atccactcttcggctctatctgccttcacgacacccagagaagtcaccatgttttcgttgctgatggagagaatgttcttcttttctcccaccaact
gcgcatggtgataggggaacgcagtgtcgctcactttacaatgtccgggaagattccagttatcactcccgccaatatcaatcttgaaacagg
caacagttttcttgatcatactttcaacacgttcgacgattctgggcatacattcactcagtaattcaagttgtacctttaagattgactttaacaac
accacattttctgtgtctgaaacagctgcaagacgttcctcttcaaggatggttgggtagaatattaccttgagccccatcaaagaggcattggt
gatgaaggcaaccttaaacactggcacgtctgttgctgttcttccacaacccagaaactctctgaaggcaggagtcacacatagtttctctgca
ctagaagtttcttcctcataaagcctcatcataataataagttccctgagaagctttccagttccaacgtggggtccttcttcatccatgcgcctat
tttcattctggcaagaacgctcattcagcttgtccagaaattgatcaccagtaatttctggaacctttttttactttgtacatgcctttcttcttggatttt
gcaacatcctttaaagttgcaggttgactgtacattatcaattttccatcgtttcctctgttcactgtgtcatattcaacaacattacatccagcatac
agaagattgttggggttgacactgccgtctccgccgactctatcttgtacagctagatgccccatttgtttgaccagagggttgccattcttggt
caagttatcagtgactagaatttgttccccttcgttgagaaggagaagaggtagacgtctgttgtcgtactcgggcagtttgttggagagacag
atgcgtagagtcgttggttcctctttcatctggttggcacattttattaagtgatctcgcacatcactttcaagaaacttgcctcctcctaggttgag
cacatttgccttaaacatattgacgagtgcagtgtgctggttataagttttgtttttttcttggttagttatttttctgaatgtaggataaaataacgtaac
ttttcttgtgtcgttgttcaccaaggaatcttttgatatcaattacagctacaactggtaaggggggttgtccgacgaccgtctgtatcctacggtat
ccagtgtcggactgtgaggcagagagcttgtgagggcgtatttatacaagaatagcagctggtgccaactagtgattttatagaacagaaatc
atggaccagtacccagaagtgagggatactcctcagacagaacaggaacaggcggcggcgcaacaacaacaagcagcaacaacaaca
gctgccgctgctgctgccgctgctcctacgcagtacagtaacactgtttctgcagaaactttatccgccatttctgaagatggaaaattggaga
ggtcaatcgcagcttcgtgctggatcaataaccttaaccctgatgaaaaaatggctcaacgtgtccaattccatccacttagttccacgaccac
atacgattcagaaaatgtgaaccctggtagttctgttgtgtttttgaagcctagagccctccccaccgggggcacgtgtttagcccccaactac
attgcagtgcctactcttcgtgctgcgtcagaaattatcgattctattgcatcaactagtctataccaatgttcaatgttcaattcgtggaatcttatt
cccatttttatgagtaatagcaaacatagtcaattcggtgatcgtgtcatcaagagatcaatgatccgtaattgtttttccaaacaaaaaaatgtag
aaaatcttttaaaggaattgcgcaggagaaaggttaacgccgccaaggcattttctcatgcagttcaacagaaatcagcagtaaatactgccc
ttgcagcatggaacgcgggaagtgctgcaaacctcgaaaaattagtagatttttgcaaattgaagtacagtcccgatagaaaatacaaagca
ggaggtctttttagtgcgtcagcaacagctcagtcacaatcaggtacttcttcttcctcagttgaacatacttctaatgatttcttactcgatatcct
aaaacgtcacaagggaacgtcccttgatttagattcagcaacaaacacatttgatactgctctttctagggtttttaccgagtttaaggaacagg
ctagagcagccgtagatgcagccgcagattcagatcacctatccgcgtcggaccctattttctctattgttcgacacaatagcaggcgtgagg
gtattctagattcagtgccaaatatcggtatgttggctccacgatcaaaatattctgtcgcagagtacttgatggcagacagagacgagtctgc
agatatcgctgctaagatagggacaaagattgcaacagattttgaagcattgagaggagataataataagagaagagcagatacttcagtcg
atgatctcaaagaatcactagcggattctattgaaaaaacttccattaaaaataccggcgatattaattcggtcacaaatattcccacagataca
gaggagtatgaattttctctccacatcacacagcttttcgcgcaggcatttttggaaactatgggaagtttattgagctgtgctttcggtgtacagt
tccccttctcagatgaaggttttgcggctatcgaaaggattatacgtaaaacagatcctgatactggaaaggtttcagaaatggacccttcctct
ctttcagatcaatatctcttgttggtaggaaatttccaggtttctcccttccacgtttcagatcctaaggatatcgttttcggtagacaagttacgcct
aatacgcctattttgagtatcattactagatctaaaaatgataagaatgaaacgtctactattattaatttcagggataggcttcttgttaacgatac

cgtgttgagagacgccacgcaaaacgtctcaacttccaccccttctcaaagaagagtccccaccgccgcaggtgaaccaaaaaagccaat
gctctctggttgcctacctatcatcaggggaccccaggtagtgaccagggaaagtgatgatatgattagtggtcttgtgggtgattggtatatct
ctttgggggtctattatgccatgggttcatcggctgcagctattgccgctggccaccagagagctctcgcttcagccgagtcaattaattcacc
catgatgaaaaaattctcaaagaagggaggaaagtatacagaggaggaaaagcgaatcaagaaggctatgcgtaggaatgcagatcgttc
tgctaggattttggctttgttggggcaaactgatgcccagtatgggtatgtcgaacataattctaccctagattcgttctggtcttcgaatgccgct
attcgtgcaaaggcaaaggaggatgctcttagccgtgcagaaatcttggcagttaggaaacaattggacggaaaatgttcctcttcaaggga
tgaatattccatggtagagagatatcttagagactcatttttcaggtcagttaataggtcaggaggaggatatgaaatgtttgatcaaggttttgat
atgggaagatttgccgactttttgagtgacaactctgctgccaggaacgcttggcaacagtacgcagaagtaatgagaggactttctaagcat
gagaaacgcgtattcaacattgaaggtcttttcagtgctctaaattctttcaagttccccccttgttccagaacaggggcgtaaaaagactgtagg
tggaaggcataggcttaacaatttgaaggcggccaataagatcattaacggcatcacagagatgactctccagtcagccatcgatggtactg
gaatctctgatattattggttcagttagcgatgggtggggaaatactacagctcagccgtctcgcgtcaaagctcttaaaacattgtccaatttca
gcggaaacggaaatgttgtgtcaattccagttagtcgagcagtaaagtgtgccgcgggaagtcgaggtggggaaactctcaagtgtgtgga
cattccttcagtcattattgcaaacctaatctctgataagagaattctagatcaactttgcggaggagggatgaacctcgctcacgaaatcacaa
actttatcgagacgattgcaggtaaagaacatacaggaaaggaatccgttttcttgtctcctagattgtctgtcattcttttgcgttatatttggttca
acgcagcggttgtttcccttacggatagcaacattaaaatgcccctcaacacaatgtccgagggcactggcgatgacatttatagggactattt
ggccatcagaggcatggttaataattacaacagtagtctctcctctatttcagtcaaggctatttccgataggtacaattgcggtagtggaaata
cttcaaccagtaataagaatgtgaccattaagactcaaggtgaattgttaactgtccttcaacagactgctaatgccttgtccgcctttaccaaca
agggaggcgtgggtgcaacccccgatgctgccaatatggccaacgttatttccccaattgctaatgcggatgtagttaagaacaccaacgtg
gttgtttcagggctagataggatcactgagaccatcaacttcttttcatttttgtctcagatcaaaacaatgaacgagaacattgaagagtatctta
ggagatataggctaggagaaggactagataagaaagaattggataattttgtgtatccaaatattgcagctattgttaagcgagaattgggggt
aagtggttccgcattgtccagtaatctcgatactgatcgtccaattactatcgacctgaacactgaacagcctttgatcgtaaaggctagcaag
ggttatgcctctaaccgctacgctaaattattcaacaaaacaacaagaacagcagcagaacaagctcagatggagcagtataatgcacaaat
ggctgccaatactattcctcaattagtaaacaggttgaccatccctggatccatcacggcagacactgccatcaatgtcgttaaagctttcaca
gaaaatggagaatttagtaacgcagaaacacacttgggggttatgggtaacgcgattaatgaaatgcaacctcttttcacggacggattcaac
gttgcgaacaagcgtttaacagttaacgtgggttcagttagtaagctgattcagaatgggttaaccgtatctctcattcttgctcactcaaaggct
agcccctatgtctttaagcctctcgtgcaagatttcgctaagcttttactggcagtcactgcagagacttctctggttgtctctaggtcccagaag
agtttcttccccattcctccttcagtattttcttcaggtggtcttttcaaaattgatagggaaatgttcgataatatgaagacagattatgtagtggaa
gtaattagacagctatctaagaatgctaccgccgccatcgaaaggtgcaatgattccgattcagctgctaggattgccaagtcaggtgaaattt
ataacaaggatgttgcatcaaccactgcagctcccggaacttcttcctccgctttaaccttgttcgccaataatctccagaaccctgcaaaggt
atggtctatgggagctctcccccatttcgatatggccgtggtaccaaaacttcatggcatttctcacgatcaaatgttccgcctatctacatattat
cagggtattcataagatggaacttaacagcgattgcaaaccagaagaatgggataattctcttcctggaaatagggctagcaaattctttggcc
tttcttcggtgagcgataacaaccgttcattcaatttggcattggatactcttttggcttcacctgcagagatttgcgatctggtgacgagggaaa
tggtaaagaccagtaacgatattgtgcataatattggatcgaattccaacacggacgcgcttcaaaagagccttcaagttggtgcctcagcag
tagaaaaatacgacgagtctactctttctactaaagaaactgacgtatattcccttgtttctgctctggctaagagcaaatctcctctatcttcttcc
tcatctttgtcgtctgagggacatctcacctctaaggagattgataggacatggaacacccccgctcttctcggtaccgctaaaactacatcct
attctgtttctgaagacgctctcaatgctcctctttcagccgtgttggactttagaaggaatgttgtggatgctactaaatctctgtacgaagttgc
agctgtttgtagtgtcatgagtaaagaggaggatgtgcgttcttcgagtagaaagattatgggaatggtggaacaagaatcgcccgttatgca
agacattggcattgaccgcattgctagtcttgttagtacagttgctacccccaaacagcatcgcagattcttacagacagtaaacgattacaaa
aattatctcattagaaaagttgcatcgaatccccttctctcttcaagattgggaggaatatcccctactagtggtaacaccgattacaaccttaaa
gctgtatatgatggtgttgtttcttcctcatcatcaatgaccccctcgtccatgtctgtctctgacagattctggtcgggagtattttctcagtgccta
gagactggcccttcaatgtttgccgatgctggtcatggaggtagtaacatgttccaaatcactgcacctaaactttacggttctagagtcaaca
cctacgcagctctgagctctggcgttgagcggctgagagactctatttcttctgcgactcaggaaagaaagaataggattgcaaagagcatc
gaagctctggaaacgttcgtaaccgatgtggtgggggagatactttggatcaattgcgtaaggcccagaacatgtacaacaaactgtcag

atattacttccaactctatctatagtgatttcggaaacattgactgcgctaaaatcatgaagaatgtgacgagcaagaaaatgaccgctagaca
acaatcagatactattcttagctctctcttgcacgaactcgctggcctggtacacaaacaacaacctcaattggctactcaatttgctctggcga
gccatgttatcaaggcaaagtatgtcactaatgacctcaataatatccacgagaaggaaacattcagtcaattgatggccgtggccggtgttg
ccgattactataatgtgtcggcagctgccatgtgtcagcgtctagttgcttccgacgtaacaatgttcctcggcggaaccatgctccaacaag
gcctgttcgtttcattccttcttaacaacgtacttttctcccaggtttctgataatattaaaatgaacgaattgaacgatgaaacaaagtctcttttgg
ttaaactggtaggattttgcggtacagtttcagatgcgctaggatctaggcacgtgtcttcaattagacgtgtacagaacgaagaggataaga
aattagacaggagttttgttacatcactttattcagcatacagagatttgaggaagaagactgaactatacagggaaactgatactattaacaaa
cttttcggacatcaaaactttatgtcttacgaatcttccatgctcaagaggacttccttggtacatgacgctgtttccggccctaggccaagaag
gtacagcacccttgaggatgtacttgaggctccttccacggttcacaaatcgttcatggtttcttacccagagagggcagctgcttctaggcga
gtgaagagggctggactcagggctctggctgataacaggatggaatctctttacggggaagaagtcttgaacgatatgaggtcttcggcgg
tctcttccgaaatgatggatatagagtatggtgagggaggattcatgatgatgattagtgatgatgaggatgatattgcctttattgattccgaag
aagagtctgaatcatctactgatttctcctcatcagatgaatattccgattcatccgatgagtatgattttgatgatgataataatggccagtctcct
tattcaactacatcttattcgtatgatgctctagaccgtctgaattctgccgctaagcctcttactgccatctacgggtgcaggggagaaggtga
agacgatgaggaaaatgacctctatgaagaagaacaagaaaggaggagacgtcgctcatcaaagatggggaagatccttagagatcttca
tgagagtgatgatgacgacgatgactactttgatgacgaatttgatggcgaacgttcaatgtcagaaactattgcaaccagaagagctggcc
gtattcaatatggtccaggtttcctatctcattctaatattcttaaccgtccggctaaagcacgcgctttcttgacacgaggcaagaaattcaggc
cttctgcgtacgatagattctttatggaggatgacgattccctcctcttctctgacgaatctaccacttcttcttcctcttccgattctccattctcctc
cttcagcaaggggagaaaatgcaagcgccgaacaagcgaggaccaatgtgcctttgttaagagagttgtacgtgcttttgtgcccaccaga
gtaacaatgatcaatggtcgagttagcatgatcaccccagtgactagtgaaaatacagtaggattctatgaaaattaccagaaggccaacaa
gagggaaagggcgcgtctgatcgaagaatacaaaattgttaagggtgcttcagctaccttgcccgacgaatacgtagagggtagagcatct
aaacaagtgtctcccagggaactgaggaggtctcttatcaaggcagctgcttatgttgcccgcacccaagaaagtaacttgaatattatctttg
acgctctcaccacaacatcaaacgccactctagttaatgacccatctactcttttgggtgatacacttttgttcgccaaacaactagaggcaatt
accgagaggaggaataggctaatgaaagacctaactgaaatctctccttcacttttcacatcattcggtgatgcaagtaaagacacccaaatg
atggccgatgccaaacagatcgtttcaggaggaaatttcaagtctgccggctatctaggtgtccctctcagaactcttgcttcatgtattaaggg
cactaatacagttgatcgtcttttggctaccaaaaataagaaccatctcgaatggatgaccacagccgctattgtttttgcacgttcattcaacga
tactactttccatgcactcgaagatacactaaaaatgacctccgctttgacagacatgtacagcgctttcaccaacctcgtcggatcggaacat
tctcagcgtctaaaagtaaagagtactcttttagattctattttcaacactaggatggctcacactgaagcagtcatgggtctcgtataccctaca
gcgttcatcaaccatgaaatgccctctgattacacacagcgcagagagatgcaatcactcgctcttaacattcttaggggagttaattgtagcc
aattgccacgaaaggatattggagacactgctggcctgttgacctttattacatcacgtaaatttgcaggttatggaggagaaaggggaggttt
gtctctatacagaatgtccattgttgatgctctttcttgcccctctgacaatcggctcaagggagcagtctctctagaggtaggaaagtggcag
gatatgggagaggaaatcttctacaagaggagcaacgatctggtcgatttttgttcaaagaacaatatttctctggaaaatgccgtaggtcctat
tgctaggtttgttcccaatggaactaacatggctgatattggcatgaccgatatcatttctagaacagtcaaggatgacgcttcaatgatcaggc
ttaggcgcgcagaagagggcgctggcgcagcaggaaaattcattacagcctcagccatgggtaatttgtacggaggtattgataccgttgt
gaacctaactgaaaaactatacgactcgttcgttctgctccaagattcagactcgttcaatacaccaacagaaatggccactgctattatcaac
cgtatgaagtcgaggaaacataaggctctcaaaacaccattcgggggagatattgccacctataagaacttcccatcctcctctgaagcaatt
gtagttagagccaaggaaatgcgtaactctattagcactatcgtgatggacatttccaagtcaaggggaatcaactcgttctcatcccgcagtg
gttctactttggccaagatttccacatctgaatttgaaaggatactagaaacatctgctgttctttcaaatacaaaggccaatctgagaaccattg
agaatagactcgccgaacactacaacaaactaaaacaattcagccatattagtaatgatggactttccgagacacgcgcagtcgttgccgta
attgctgaatctttaaccccccgtgtatgcggatgacaccagcgagagaggagcatctgttagtgaactattgacagacaatactctcctcaaat
ttattgttcaaaatgaactgaaaaacattgaagaggcaaaacgtcacgtgaccgccgcaattgaaggttcatcccaactgcacgaaaaaatgt
tgagcctgctcgttgcctcagccgacatcaaccgtatgtccgcccaaaataacctcgaatgtaagaaattgactgaaggaaatagtaactttgt
accaatgactaacgaccaaggtggtacattcataaagcacaaagaaacaggtatctggctgaagaccgatgaagaaaataacaccagttct
atcaaggacaatgatcagcgtagagtagctaaaaccatcctcgcaattgtagaggacaatagaaatgcaaccatccgttctcgtctacagtct

ctttgctttggaaaatatgccatgaacgacatttttgcacttgatgatgccgatattaagaatatggacaaactcattgaaaaactaggcgaagc
actcgcagagaaggcatctccttctagctcggccatttcttcttcctcatcatctaacacaacatcctcctcttcttctcccagttcttccccatcat
catcatcttcctctttctcaatggactattcaaacaaccttgccaaaactatcccctacatgcctatcgtcttccaaaacaaacaatctaatgtcaa
ttcttctgacgcatcatcctcatcaccatcatcatcttcttcttcatctgccaatattgataatgttgagcacaaaaaagtggctctccaacaacttc
aaacacaagaatctaacgatttgagtaacgtactttctgttaccaccaagcacagatttgcgtctcataatcaagctgcaactgttggcatcttca
acggaaggcaacacgcagagacagttgttgctataccaaatgcaaacaaggctaataataatgccaccgtttccgcaggccaaggaattctt
acccgcttctcagcccctgaaaatgtttcctccaccagcatgcaattgcctccatcatcatcatcatcatcaaatggagatgataataaggtacc
agtaactgtcaggcttaaccagtacgccaactcaatcttatcatctattgaaaacgcatcagaatttaaggacttgaaggaagcagaaaggaa
aatcgatctggccatccaggcagcttccaccacagaaacaaaggaaatggtcaccgtgtctaagtgcccctctgctaaccagactgccatc
actgccatctctcaagctaaatcccttaagaaaagtgccctcgaattattggaaagagttatcaaggcagtcgaggtttacaccccagattcat
ctattgcagccgtttctcttcccgttaatggagattctatggtttcttcttcctcgggatcaggatctgctccctcttcatcatcctcctcctcatcctc
atcctcttcctctaatgtgacagactatttcaactatgcttacggaaaattgaagaacattgatgaaaatactgaagaaggggcagaaactgtc
cagaaaaacatggtcgaacaagatgctgccgttcgcatccctcttctagtatcatatgctccattcagcgaaatgatgagacgtgctattgaca
agttgaacgaatactaccaactgattgatgccatcaaaacaaagatcgtgtcagacactaaacaggcttcctcatgggccatcaaggaaacg
gacaaggagcttgatatggacaaagaacaggtgatttcaaagattaataacttgcaacaaaacttttcaaacgaatcagacaagataaagatg
gctattagtgttttggacaacaaaaggaacgaattagagcttcagaacaacaaaactaggagctttattgaaactacaaagagccgtatcgag
gctggtggaggagatgtagcaaacttcaaggagattatcgattacgaaaacacatctgaaaatgacaacaatctcttccagagcctgaaagc
attcgctgctgataactcggggacagtttacacccccactgacatgagcaatggaagagacacaaaatcagacagtaaatttgtcgacatgt
acaacaaacagattctcgagggaggaatcaaactcatcaatgagggacaaaatactgtaaaggtagacttttcaaaggctttggaggctttcc
ctagacaatccaacggtgcttcagagcctgtatcttcttcagttgtggagaggagacagcgagaacgtcttcaggctgtcgagatgtttatgg
caataatgatggagcgcaccgagcgtctgaggaagaggttggcagattcggctgctcagtggaatactgttaataatgtagaagaaactgtt
aatagtggtatggttaacatcaagagtgaaaggctcacagagattaggaatcaagcacaaatcgctgaaagcactgcactaaactccatcaa
cgacgagattgtagagtctcctctcaccctctctttgggagcacgagtcgaccagctcttgatcaaggtagatagagtaggaagtatccaaca
gcagcaacagcaacagcagcaacagcagcagcttcccaaattgacagctacagaacagagaaaggaacaacaatacgctgcagatagg
gttgtttacgatccttcatacacctgcttcctgcaacctcttcacgagacaattaaacgtatttcttctgtctataattcaaagaacaagggtcctct
cagtaacacacgtggtgttcccactagcgatgccgacttgcaactgatgaccatcactgacttgtctaggtctgtactcgactcttcttccacttc
ctccaagaaaatgctgtacgaaaatgttccctcatcaattgttcctggactctgccagcaatgcgcaatgatgatcaccaacgtccacgaagc
cactcatacttctcctcattcattcaatttcgagaacaaaagatccctgaagcagctgacagaaatgttgaacgctgccacttcatccagtgac
ggtcctgccgtgagacacgatgtactaacaatgttagagtccaacaatggttacgtcaaagattttggattcactcaccgccaaaaggttgcgt
gtatcacccctgttaatacacttctgggaggtactttcagtggaaatgttgcacctaatactgttatccttcctacttctgagttgtttaactgccca
ggagttgaaaatgacaaatttagatccatggttaacaggacaaccgacaagaatgtggctgacgcacccaagtcatctgcaagcatcgtgg
agactcttgctcgcacgtctcccaacgccgagcacctttacttccccttcaaggaccagaggcgacacttcaactccatcaccgacgccatc
atttctggtatgagcggcgaatcttcatctcaattgaacactacttgtgatcaaaatctggtaaacattgatcaaactactggcttcccagtgttta
caggaagaaagcagggcgaaagaaggattgtgcacactgaaaacactatggaaggagctcgcaaggacaagaacagtggcatcccttc
atgtacaaaggaccgtcaaacttatatcgatatgggcacaaaattcatggttgctccaggctctcttctgaatgctaacaaggaagaaactctc
cgtctaaacaggctttcagacattaacaacgtgagacattatggcactgatgttcatgtggcaggcgcaaactctgcatggagaattggtgag
gtggtgagagccgcctcctcattccctgacggagataaggaatcggctatgaaaaagatgcttcttctaggatctgtatctgccatctctgctc
aaaaatctgccagtcacattaacgatcctactgctttgttgagcaccaacacgtctatccagaatctggtcaaggaagctttcccagaccctgtt
tgttcctctaattacttggggtctgctgaatctacgttcgccactcaactcgcctaccgccagcgcctgttccctaacggagatgacgaaaacg
ttacaaccgtttcaaatatctgccctatggatttgatgggaagtacaaagcgctacaatgacgctttcaacaacatctttggctctaaaatgacat
ctactaacaaaaaggggtcaaattgtgaaaatctactgaaatctgccatgtctaacgttcctgctatcaacactgcctttggagcctttgaagaa
gcttcatcttctgtcaggaataggctttctcccctttatgaagacagcaccaaatattcgtccaaccaacttgctgtacaggccatgaccgatac
tgctgtggatgctttgtccgctgtttctactgttgtcggtcgccagaatggcagaaatactcttctttctctccctacttctattacttctatcgcaac

cagtggccgtccatcactctcttattcttcggacatgaaatctaacctcatcaagacaatttcccgcatcaatagagacgctagcctcctgtcaa
tgggagacagccaagtagctgcaggttcttccttctttaactctttccttcgttcttcttccatccctgtcaccaccagccaggatggaaatgttgc
agcagcggaaattgttctggggactattctcgacaagactgtggagatcaacaagagattcgagatgcttggaggaggaaaaatggtcgcc
gggagtcctgaagctcgtgccatccagcgcaatacaatgtcctctattctccagatgaacgaaaatgaactcgctcgtgacttgtgcgaaatt
gaaaataaaattgagactaggcaactgagggatgctttccaggatttgaagaggtctatgctgatgactccaggaggcgtgggagccatttc
ttctggagcaagtaccaacaatgttcccctttctcttctcatgtcacgtgtcgatgcatccagcggtcttctgatgaacaacaacagtgccaatgt
aatggaagctgtggatagtttcaatactactcctttgctcgttaggcacatgatgttggatagtggaaagtcccccgttcccatggccaaggaa
attaggagcatgctgacccaaccaagagctctcaccgcccgcgctctactgagcgaatcttcccctcttctcactgaaatctgcctctacaac
acccgcgacactcaaccagaaagggcagtcgacagactactaacttcagcctatctagtaaaacaagctaaaagattcgacggagttgacc
cagccttccctgccgccctcacctgcgcttctcacctcatgctttcttccatggattcccatacaaagtcatctttcatggacaacatcaaattgc
acatgactgatactcaatgcttcttcaagaacattgaacgatttgagaaattcttgggaagatatggggacgaatacgccatgtcccacaagc
aaaattgtaactgccccttccatctccaccacacttttactccctcagataacgagcatctggtatcctctttcgcattcgcccgcccagaagtct
ccatggaagaaattagagccacaccctatcaggccaacaagcttattagtgacaaacattacgtgatgaacatgtccaagatcgattctagag
taacaggatcttccctccttaagaaggttagcgaatggactgaaatgagaatgaactccaactttaatggaacatttgaaccatcaagactcgc
cctctccaactctggcatgacaacggcaggagtcaacctcgacgttattgtcaaaccaaataatgcaagaagtgtactaggaatattggaatg
tcatcgccagcacgtgtgcaccgccgacgccaagggaactgtcgcttcagccatgccagccgtcttccaggcaaccgatggaaacggtaa
cgaatctgaactgatccagaatgctctgccaaggaacagatacatccaaaagagcacaatgaacgctcaaactgtcgtgtttgctaatgttttg
gaacaacttatcgccgatcttggaaaggttatcgtgaacgaactggccggcaccatcgctgaatctgtaccagaaagcgtatatgaaaacac
caaggaaatgattgatagactaggctctgacgacctcttcaaatctaataataatggaggagtagaatcaatggattatgaagatagcgaaac
aacatccaacaatggtcccgtcctcatctcagaagccatgaagaatgccgtctatcacacactaatttccggcaaggcagctcgcccggaaa
atgtaccattcgcctcatgcgccagcggccctctcgcctttgatttccttctgtcaaagggagatacattcgaagaaaagaacgccgaacaag
gtgcagcagctgccgtatcctctacctattcttcctcttctaacactactcttcgtaagcatttggctcgagttttcgaagccatctctaagcaagt
aactgatgctgaattcaaggatatcctcaacgatatcgaacgtaatatttcttctgactatactaactgtccaccaaatactaaccaaaatgccttt
gctctagctatcaagagagaattcagcagaattgtttccttcttaaccattcttcgtaagaacattacacccgcattagtcgaccctaaaggcgc
gttacacgagaaagtagccatctatttgacccttctttcaaccaaatcaaaactagaaaactttttccaatacggtctcagtaattcgtcctcagtt
gatcttagccatctaaaacccattaattgtagcaacaatgtcaagaatattgaagacacattcatgtacagaaatgtccaccctattcttattatgg
ccctcccagaaaatttcacagctctcttgcaacaggaacaaatggaccccgatactgccattgaaagcagacgctcccttaccaccttcctta
atcatcccaacactgcttcaatggcgaacggtgcaagagccgctgtgggtgcaggaggaggaaacccaatgggcttgtatctttcttcccac
attcttcacgagtctaccgtcacaacatcaaaccccgtcacagacaccacagaaaacgtcaactatcattcctctgttacacaagatcctgttat
ggtagtgaaccccttcaaggattctgctaggttgatcgttaacaacaacaatactggaattgatgtcttgaatgataagtcgtgcaactacttgc
aagtatccatgccatctgaatcatctggcctcgtcaccaatactggatgctcttcttcttcttcctcatcttcgtctgataccttcaagtacgtcagg
agagacaatacgcctgtgaatcttccccgtgtcacaccagccgttctctgttctgatgcttcctctaatctcttggacgtgttctccagggcagat
attgtcctcgaaaacatgaacgtgagatttggtttcatgcccgagattattgctgccgtctccaaattcaaggggctgaccaaggaagaggtta
ttaagcaaatggtttctcagaacaacatcaacaacaacagcaacaacaacaacggaaatgggaagaaaacaaccgtcgatccagtcactg
gggatattgttatcaccaatgccacattccccgacactcgtcctctatacactgcagcaaatggaggaacatcatcattcaaatggggagatat
caacgacagaaaaatgcacgccaaggctttccccaccttctttattggtaacccaaccgccgccgcaacagctaacggagtgcctcttacat
ctgagggaatttccctcactgaagaaaaacgcaagaaaatcgcaggcatctctgaaggatcaattggcacgggggctctgcgtgcagccg
ccaacacccgcctctcatccgacatggaacctgtcatgaagggatggaacaacattgttcagcttcaacaaacattcaagaaagcttcagat
aaactcactcatcttttgagatcgggaggaattccacccagaagccaagaaacaaacgctattattaacaagatgcacgacagcttcaagac
attggaggaatgtcgtagggtgatccaagacgaggctgctctgctcgttgccaccagcgatcttttgaccggtggctacggaggagatgctg
ctctggccatggtttctccagtacgtccagaaatgactggtcttattggtgcaatctccgcgccagttagaggtattagccacttgttgaaactg
ggaggtgtttctgctgctaacgcagctatccgcaagcgcctcaacctacctacatccaacgggaaaacactaccagaacatggaatcgtac
acaaatcagccaagacacttttgcttgattcagactctattagcaacctatacaacactgatcttcaagacgttgtctctaacgctagggataac

aacaatttgggaagaattatgcaatctttgggacttaaggggaataatgcaggggatttggtttattctgctagacaactgacggaccttattact
gtaccagaatatggaaacaatcgcgatcttaccaagcgtcaagctatccttaaaatgctcatttctaaccctgaaattctagaaaatgttgcaga
taccatttaccttacaacaggtaaaaatgctctcgcaccggtatctgctcaggaaatggcttgtgcgtctctgacagtcggaggaagtggagg
aggaaaactgtcatcagacgacaatgttcaatccctgaaccgcctttattttcgggtctagactagtgtaatattgtatgtttaatttattgtgtaac
aattggtttatatttgtatgtgatttttcggacaaataaaagaattggaataaaagactttgtattatttaccaaatttattgatttttttaaacaccttttt
tatgccacggggttgattgtagcatttcctccccctggagatttcatagtgggacatccaaagagtgtaactcctagtgctgatacacccaaac
ctaggaggaatccggtaagtgcaggagtaacagaaagcatgaaagcgaccatgctcaatgcgatgacgacaacggccaagccgacgat
gatgtttctggtcatatcagacatgatttctggtatctgtttatagggtttcacccagaagtgttttaaacccaaaagtataaattttattctatactgc
tcatctcactcttccgacctcacaaaggagaaaagagtcagtcttgctccagccaccaccgaggcaggagctcctcggacgctcctgtatttt
tgtccagaaatggataaagtttgtgttatatcaaatactagggagcgcacgtttaaggtacctgccgatctactgtgcgttgcaacagaaccgg
aaatctctaccaaggaagaagatgcaggtatagaaatcgagacaagagtggtggtgttttcaaggtgcgtctcggtccaggaactacataca
ataaatccaaatgatgaaggattttctgtccaacttttcaaggactacctgaaattgcaatctgcacaaggaaaaaaacccattggtttgtacatc
caaataaaggctggagaggatcttgaaaggagattaatcagtggaggaactgcatacctggatccggcaacacaccttttctatcttgatttct
ccctttaccctaattattcaatattcaatgacatttcatcccgtctgaaaatcattgatgaagacacgtacaatggtgttgttttctctaacagtgaa
gaaaaagagaaggatgcactagtgctgataagggtgactttttctacgcatgaaaaggcaattgaagcagccataaaaaaaataatgctaag
gaaagtgtttttcaaggatggagatcttgatttcgggtacttacgtataccaaaatctaaactggacaaatttactccctattttcggagtcaatac
gggatagtaaatgttgaaaaaaatatccctggttacatatggggagaaattatgaagcaacgagtgcgatgttccagatggtacctttacaac
accgactcggaatgggaatataaaaatgtggccgaagaaagagttggacctcgccagttagtgaaaaaatatggtgccaagtgtgaaaattt
atgttttagggacatagacctcagaaaaaaggaagcaaaggaaaaaagggatatagaaagagaaactgaaagcagatatgtggtcgtaac
actaacccataagcatgaaatgcctgaaaatatgccctattttggaccaaagtgttcagtggtgaggttggatgaaactagaatacttttatgttt
tgtggatgaaatttcttataatgatgaagatgtagacgaaattttgtctgagaatagatcactaagaaatgtttctattagacataaggaaaatgta
cctgtacacacgttattaaaaaaaggtgtgtctattcatgctagatttacccttaatggtttggatgatgctttaataattttaaagagaataccaaa
aacttattttgaagatgaggaactacaagccgcttgtgcgcatgttaaccttgaacagtacgaatggctttgttctaataatagagggaataaag
tagaacatgtaaagtcgcgggtagtgactcgagcagttaagcgtaggagaaaatgtagacactggatttattttgataaagacactttaaattta
aactacaaatactttgataaaaaagttactgctagtatggcatctaaaatatgtaatgcaaaacacgactgtttagttttccatagaaaaatggaa
ttggaagatttgactgagagcgcatatttcaaggtagaaccttccccaataaattttgccaagttaaaatcttgcccggatgttaaatatgtgcag
aaaaaaacagatggtacattttctgttataagattctttagaaacatgacaaagggtgatcttattcaaaggatggatctttttttgtaggtttattccc
gactcacacactattacacttttgagtagggcggatttttatgcatgtaaaagaggagaatctatgcatatgtgcacaaacaaacaccgtattctt
cactacaaattctccaacgctccccatgcggccatcgaacaaataaccaatatcatcagtgatacaagaggacgtaagggtatacacataga
atacgcgatcgaaaatgtacaagaaatgtacgaagaagatggaagaagatatgaagctaaatacactggaactttaaccgagtacaaaaga
aatgaggacaaaaccttcaaatctcttcttgctcctcatttaacacctgtcaacaaaccatataatattaaccatttgtatgagcaatatggaaattt
tgatgaagaattagaagacaagttgaggagtggtttcatttcttatgacacgtatgttactgcaaaagacaactggggcaggtgtgcaactgg
aaagggggcgtgcatctaggacccataactcaaagtcaatactggtatgtgtttttttgagaaggtgtcagaatgtgcacattaaaaacatacaa
aatgactacttcaacagaaatatcaaagaacctttcagatgtgttatccatcaaggcaactggagattggtgcagtaatatcaagacggtatttt
caccccttcacagaaggcaagggaaatttaccaaacagtctcccgtttacgagaagtcccaatacaacatgtggttcaagagaggcggcaaa
cgccacagagcattttatcaccgtctttgcaaaggacaaatataagcggaaaagagtaaaacgtacaatcggattcaccctcgacaacacaa
aggagttgacgcccaacagatacttggtagcagatgtatactcttggcaagaagagaaaatggtgtttgaaggattttgtgtcccaccaggaa
agtcgggaacatttgtacgctactctaatgaagataaaagttttctactagcagataccggaagatatatgaaaaagaagtacgatgatccag
aaaataagaccagtagtgggggtgatgatgacgatgacgacgatgatgatgatgacgacaacaacaatgttgacgtgtatgaagaaaacg
accccagaaatgtattcgaggtcgaaaaggatgaaaaatatgcctgtactttttcaattttggtctatagagcaatgaaaaagtctcctcctgtat
gtagagggttattagtagagacagatggaccctcatctcaccctaaacgggccccgtcagcatttaatccattcggaggaagttctatgttga
acggttatggtgcaggtgcagatgcactagaagaagaggatgaagttgatggagttcctgaaagagagaggattacaaattttgctctcaag
agaggacctgcaactggccagaactttgtatctgttaaactggaacatgatggatctaaagcagacctgtacaacgtcacgtgcttctccaag

cagcgtggagtataaaaggcgggacgcatcagcaaatgcaactcattctttctcatcatctaaccatggctggtcgtgtagagctcgtcactg
gacccatgtttgcgggcaagtctacctacctgaaaaacatataccaacaagaaaatggaggcaataaacattgcctgtttgtcaaacactccc
tagaaactaggtacggttgtggaactggaacaatagtcactcatgccggagaagtgattgaaggttgtactacagtttcttctatcaaggaact
aatcagtgtgttaccagaagttgtggatgtgattctcattgacgaagggcaattcttcacggatttggtgctagtcaatagactggctgacaag
gggaaaaggattgtgattgcagcacttgatggaacttctgaccagcaaatgttcagtcctattcataagctattgccttatacaaattccattgtta
agctagcatctaaatgtatgatttgtaaaattgataccaaagaagctccttttactgtaaggtttggtaatgacaatgataataatgttatatgtgta
ggaggagctgaaatgtacgctgctgcctgccgggactgttacaaaaaaattaacaagaaaaagaacaaggggaaacttgttgtacttgaag
gaggtgacaggtgcggtaagagtacccaagccaaactcttgttgaccaataaaaactcgcctctttatggaggagaatacatgtgctttcccg
acaggagcagccatacgggtaaactcatcaatgattatttaactaagaaaattgaactagatgatcatgcagctcacttgttattttctgcaaata
gatgggaagtttgtagtaaaattaagcagttgttagacgatggaatccatgttgtgatggatagatattactactcggggattgttttctctttagct
agaggagtggataccgttgagtggtgctctgctagcgatgagggacttcctcagcccgatcttgtattgttgatgcttttagatgttgaaaagtg
ttcaaatagggatacttttggtgtcgaaagatttgagacaaattccattcaagaacgtgctagagccctattcctagacctcgcaaataaggac
gaaaagaatgtatggattaaggtagacgctcgcggcaccattgaggaggtgcaaactaaaattataaatattgtatataatattgttgaagaat
aaagaattgtaaactgtttccatgatgtgtttctttcctagtgattttcacatgttacctagaaagactttgcccgacactgaaaatgattattttgtct
tggacgagtctcttctggagaaggtgtactatgataacaacaatgaactgattgtaagagttggtgggatttatatgcagatatgcaagtcaaa
atacatcttccatcacgatgatccagagaggttcttttatagtgtgttggaggattatcaccccatcaaagagattgttgaacgactagcagaag
aggatggggtatttttaggaccgtgggagtttttatcgcgcaaacaagtgaacctccaacacgggtgctacaaagctcttttgtcattgccaga
ggacaaatattgtaacctattattaccccagcaaatgaaaaccaacctggaaaaaatggaagaaatacagcgtactagactcattcactctag
aacgtacaatacaccccagatagaattgtctgaccagctagatggatgtgttatatgttaacactttttcagtaccatgtaatatttttgatatcaaa
ttgaaaggaaaatggttcagaaattggtagtaaaaactttattgtttctttcctctttcttttgtctgtcactttattcgtcagacataacaaccataata
ctgagtatgggtcttggatctttgttgggacattttgggtgtgttaaaagtatcacaggaaaaatcgtctcaagaaaggcaatggataattatag
gctcgtgtcggtcctcggctctgtatgtggagcaggaattttctgtggataccagcaatattctacacctcagtctgcacccatggcgttgctgt
ctgtcttctgctgcactgctactgctgctacatcaactattgtaggattctggatatcgatgattatccaggcattgaaaaggagtaataaagata
taatataaaatttgtgtgttttatttgtttgataatacaattttcaccttgtaccgaaacccattctctacaagcactgtgtctttggacacggggatca
atatcttggattttggatgaatgttcctgaccctcatatcttcttcattttcaaatacgaggagggatgttttctcagtattcctgttgtgtatttggag
gtgaacagggaaggagaatagttgattgttctttggtgttagatggaactcgtataccatgaattgtactcctccaatatcttcaagatagatgca
tgatgtatttttgtcataccatctcgtatggtcgtcagtgctcaggtacaatctcttgtggtggtggttacacatctttccactggtgggagaggaa
gtattatggttcaatattgatggaggatggatttttataccctcatgcgaatgcgatgtggaaaattttatatgtttccgacaaggggaaaaatatg
aataaactgagcatacacatgtcgtatccagatatctaggtcgggaagaatccagaaacgattcaaaccatttctggaacattcatttctggaa
agggttacattttattgtatctggtgcattatttctggtaccatttttggcacctctgtgtaatctgacattgggtcgacccagaggtccaccctccg
aacttgacattaggccgacccagcggtccaccctctaaactcgagtgagctgaaaaaatttttataaaatttttgatgaggaaaaatagaagta
agatgcttgtcaatgcatgctccctgtggaaggtggtggatgagataggagggagaggctatccagatatctaggtcgggaagaatccaga
aacgattctagccatttctggaacaatcatttctggaaagggttacaatttattgtatctggtgcattatttctggtaccatttctggcacctctgtgt
aatctgacattgggtcgacccagaggtccaccctccgaacttgacattaggccgacccagcggtccaccctctaaactcgagtgagcagaa
aaaatttttataaaatttttgatgaggaaaaacactaccagtggagcgaccgtacacactcctctagctaactactaaccgagagagtgaggc
agagtctgtccagatatctaggtcgggaagaatccagaaacgattcaaaccatttctggaacaatcatttctggaaagggttacaatttattgta
tctggtgcattatttctggtaccatttctggcacctctgtgtaatctgacattggcctgacccagcggtccaccctccgaacttgacatcaggcct
agccagcggtccaccctctaaactcgagtgagcagaaaaatttttaaaaatttttttgagatggagatggagtgaaacttcctaggtgtaaggtg
cactacagaggcttgtgatcctttctgggcggctcactctctagaaacgtctgctccagaaacacataaaaaagataaagtacatttctggacc
actcccatttctggtgtaatctgacattgggtcgacccagcggtccaccctcggaacttgacatcgggcctacccagcggtccaccctctaaa
ctcgagtgacgcagaaaaatttttttaaaaatttttgatgagggaatatagtactccgtagccaacatatacacatgaacacatgaggcggtcta
caacagaaagagaactgatagctgttccagatatctgggtcggccagaacccagaaacgtttcaactcatttctggacaagccatttctggaa
aggggtacaatttcttataactggtatatcatttctggtataatttctggcaccttcgtgcaatctgacattgggtcgacccagcgatccaccctcc

gaacttgacatcgggcctacctagcggtccaccctctaaactcgagtgagctgaaaaaatttttgaaaaatttttgatgaaggaatatagtagta
tagtgcccgccaaagcatacacacactcgcccatgctcgtctagctgatagctgttccagatatctgggtcggccagaacccagaaacgttt
caactcatttctggacaagtcatttctggaaagggggtacaatttcttataactggtatattatttctggtataatttcaggcaccttcgtgcaatct
gacattgggtcgacccagcggtccaccctccgaacttgacatcgggcctacccagcggtccaccctctaaactcgagtgacgcagaaaaa
tttttgaaaaatttttgatgaagagattgagtaaaatttcttgacgataagaggaggcagtaggtgaggctgcttgtttgatgtgtcagccacatc
tgcgtcatacattatatttccaagaattttgctgacgtcaatggaccataaaaggctttgtacgtccagagacaagttttagtctgatagatttctta
aaaaaaaagagggtgggaggtttgtttttgtgggttctgtgtgtgtataaaagataggtgcaaaggtagagaatcatcatatggacaaaatctg
tccatgagacctcagtagagaacgcaccatggttgcttcaactccgtgtccaggcccaggaccagttccaacccaagaacttctttctacaaa
ctttcttgaagctcacaagcttgtcgtggaacttcttctcccgtcctacagtagtgatgtagtttattgtgactctgagacgtacaccaaacctata
ccgatttttgggaacaagagtatagtttctaccattggagactatgtcttatcaaaccccaatgaagatgtgagttaccaaatggtttcttccgtct
tagaaaaatttcccttgctattccactgcacttataagacgaatgaagaagataaaggtattcctctgtggaagaagttgtacaacaaaagaaa
attcaaactcctcaactcattgttggttcataacaacaagaactggactcctgttccagctatcccgtttgacagggagaatatatgtgatgcttc
aggaaggagtgttcttatgagtgaaataatgtccacgtcaacttttcagacaatttgcaaaaacaacacacattacttgtttgatatgttaaatatg
gaacgtggcaaacaaggagggagttttcttcacttctttgcatctaggaagaattcttttactaactttgaaaatgaagaaatggactctcatgtg
ctcagtaacatagcgaaattcatatgcaatgaaaaggaaaaactagactctttcatacctgccaacggaaaaataccatgccctgataaaact
aatgatgaagggtacatcccgctggaaatagcaattatggaagacaattaccctgcattgctatatctcgtttgtaggtatggagcatcttggg
caaacacatacggggatcataatgaatctctcaaagcgtttgcaataagaaatgatgcaaaagattgtctggaaattatagagtttataagtgat
cactacagtttcaacaaaaatgtgacgaaggaagaatttgttaaagagaagactgtagaatgtgttggatgtttatatgatattgaagacgaga
aacgttgttacaaactcccatgtggacatttcatgcatacattttgcttgtctaataagtgttctaaagctaactttagatgtgttaaatgtttccaaa
cctttgatgacacaatttttagaaaatgtcccccaactatacaatggaaaatgggtataaaccaaacgactaaccataaggaaatggatttgttc
aatcgtgcatttgacacatatttagattttatttgctcatataacgtcaaattagacaaaaaatcaaaacctaaacacaaacctgaaaacaaaaag
gtggaagaagaactagcaaaaaggacagcagaaattgaagaggccataaagaaaaaggaagaagaactagcaaaaaggacagcagaa
attgaagaggccatgaagaaaaaggaagaagaactaacaaaaaggacagcagaaattgaagaggccatgaagaaaaaggaagaagaa
gaactctcaaaatataataaaataattgaaaagggaaaaagacgactgaatgaagaatgtgtcaagctgagagatatttcaactgcagccata
aacatgtacaaagagaaagtgagaattaatggtgtattactaaaagattccgatcaggagttggctgaggcgaaagagaggttgaggaaaa
ttttattgctagaagaagaaacaaaacttgacagatttttgtttagaccgaaacgagtagaagaacgtatattcctaactaaagatgatgaaacg
ttagccttcaagttagccctagaaaagaaaacggaggacataattgcgaagaaaaacaaccaaaaaggcagtgaaagaagagatggaga
atatactataacttctcatattgagaaactacctcaatccactgctttggctagtgtgtgtgtgttaaacgaataaataaaagtataaaatgtaataa
tattgttttaatatcattaatattttaccacaatgtctacttgttcgaatttgttgtcagtatttggtggaggagattggacaacaacattcccattcga
cctcgtccatacacgtcaagagtgtgataaaaagagagagcaagactactcatttttcattactgaaacgtgtaaaggagagaatattggtata
cattcgtatgaacacacgtcaaagattattgacacgggtaataatgattctacctcaatagaggaactagaagtactgaatatatacaaagctat
aaaccatttagaaaatatcctaaaactcaacaaaggagaaaaaattatactgatggatgtagaaacaatgatactggaaactcataaaatttta
atgaaagggattcttcccaagggtaaaaatggaagtttcagtacatgcgtacgctttgctgtaaataagaacaatgaacggcattactaccctg
tatttgaaacagagaaagaagcgttcaattctatacaaaatctagtagattattataatgaaattgtagctcacaccaatgaccaaattaaaataa
taaaagcgtgcgcatatttcatgtacaactttctaactctccaccctttcaatgatggtaatggaagaacagctagattattgtatagttttctattg
aaaggtaatggtatcgtacctcatttttcacccataacacaccctagggatcaatttgttgatactttagtgtattttagagaacatggagatggac
gacctttattgtatgttttgctggaatcaataaaaaataagtaaaattcattttgaggcatttatttcttcatcaattcatatggcatcttgccagttttct
gtattgttatctcagcaataatgcgtaacacatcagcagatgtgcagattttatccccttccccatccctcatggtatttcctcctgcaaagtaatg
gaggaaagaatctccagtaatggggttagttacattcaatacagcaaagaattggccaatatctttgtcggccaacttgtacaaaagctccttg
gtcatgacttcactcataatgatagattttccagtggtgtcacaaatgggggatttatcaaacggcatggcggggatattggggtggaatgggt
tggagaataacaccaggtcgccgaggagggcaaacttcccgtgcttgtaaagctccttccacaggggcgtattcgtctgcttgtagttgcaat
gaagtaggacagggaatcgcctgagtacgtgggacagcactttgaagctcatcttttcgtccgaagaagaagacagtacactctctccaaca
tcggtgacaatattcttccctcccgacttgttcttgtacaatggagactcgcaatcggtgaacatggcgtcggtagggaggagcatggtttgca

tgaacatgctgtccgcgttgaaaaagtcctgctcggcaatggagaatatgttttccaatttgcttataagtgagtccattgctgctgcttgctgctt
gctgctgttaggcgtccagaaagcgtgtgttaatttctcggcggaggactgttcttatatacaaattcgttaaagggtgacgcccacagataag
aatgtggttagacttgaatctatttgaaaccatgacaaggcattttcttttgtaaggggtgaataaggactagactcctctaccagaattcatgtg
cgtcattgcaagcaaaaactccctcttgttccaagaattggctagtgggcggctgcatgccagtctgtatgacatcatttccttatgtgcaatgg
ataattataatttgtaatatttgaagtcgtcgggtccggtccacccagaaagggggcgaatacacataacatttctgggggtgtatttagtgcatt
atatctggcatccagaaacggatgcaagcagtatctgaaactctctctcacttttgacatcaatgaaaaattggacatcaacacgcattggtgt
ggtggataaaagggggcaaccggctatgactaacaagcatttcctgcctaacctgcagccaagcaacagcatattacacaatggaggacct
aaaatccactatcgagagagtatatgaagaaagagtggagaatctagaacaatggacaaatactgtagaggaagaagaaaggactgtctc
agcaatcgattctgtcctggaggaacaaaaaagggccctggacgcatgggaagcagcgataaaggaacgagaaaacgacctcgcagta
aaagaagggatatctgcactcgttttcaacgcagcagacgccaaaacacgtaaagaattgataaatacgtggatagccgaaagggaaacgt
cagaaaaaagaagaaaggaagcaacctctaccaataatcaactgaagaaccagatgtcatctctagtcaacacaaccaaaacactcaaag
aaaagtacaacaaatattacagaagaagtgccatactcaacatgcaatacatcaataacaaaagggattatgaagcaagtcaattttgggtgt
atacaaacaatgcataagttttcaaataaattttattttataataaaaggtgttttaattataatttttgtccctgatgttgtgttttctatatttttactgcct
cctcttggggtaagacataaaaacggtggcggcggaaacgaggaacagaagagcggacccagccagaagcatcatatccctggtcctgt
tcttatatttgtcctcatcatcgttatcgttggcagtgtcgtcatcatcatcggtgtccttatcagtgtcagaatcgctgtccttctttggtcccatcca
tacattcatgacggccaggacgaggataccaattgagatagcaacgatcaaaaagagacccatgcgagccatagacatggagccttcaag
atcttccatggtgtaagaagggccagcggcctgggctccggtcctgccgaaaggaagagtgttagtcgtggtggccattttttttctgtttgat
gaagaggtacgtcaatgtaacgtaaagacctgttgttttatcgtgtacggtttttgtcaggacaccttacgaacctatttgtctttttagacttactc
agcgttagttgtcgcagttggtcatgttccgtcaattctgttcactatatcttcttcaacgccgggttaacgacaatcttagatccacggcgtccg
cttcggcagcggcctcattaaaaggagatggaactgaatttataaccggagaaccaccttctcataaaatgaggggaccttcttatagcgtttt
aggacctgatccgtgcgaggacccagaaagggtatatgttgatattgtagtgtctattttgcagacaaataatatacaggtaacaaaagaatg
ggaattgttttccgataagttgagaaaattgggtccatggattgataggagcggaattgagaataatggcgaaggagaagaagatggagatg
aaaatgaagacgggggtggaaatgggggaagaattgaagacagagaagcacatcgacgaaaaatgatgaagaaattgtcctttgttggaa
gagaagatccagtcgctgtagatttacccacgtggcgagaaaacagtacagaatttgcacgtcgtttaacactcaaggaattgtgcgatttaat
agttgaatgtggatgcatcaaatcaaaagaggaactctttgacttcatttttgaagaaccgtgggagattaaagaggctgctgacgttaggggt
atggcaaacaggagtaaattcaccaaggaatcattaattgactggtttttgagttcgacacatatagtaaatgtgtagtatttttgaagcagtca
actggtacttgaaatctcaagcgtctccaatttcattggtactagatgatatatattgttgtgtcttttcctacataagacgccaaacctttttaacta
gggcaaaaaacccatctttaacagtggcttcatccttttctcccacgcccgacacaaagcttttggctatcgacgagtgcgtgcaacactttta
aaatcagacattaatattagccagatggcattaactgaaagggactgcttcttccctcttttaactgaaatgccccgccaacaaaaaaaagtaa
acaccttcctggacacaatgaagagacctaccttatctcttctaccttccacctcctcctcctcttcttccaacaacaagagaaagagaaatact
gccgctgccaatattcttcttccagtgtacaggagtaacttttctacagcatccaataacaagagactgaaaactgatgatggggaaaatgcat
cagcctgtattcttatcgaagggtatgcgaatggaaaaataagccctataaggattatggtaagaaaatcaactattattccagaagtgtttaac
catcttttgttccctgtctttgcctctaaagacactggtgcgaatatcttattttttatcaaaatgaaatcctttgcaagtgcatctttactcctccctgg
actttttagacaccccaaacaatttctcaacgggccgtgcaaatggatgactctagcagaaaacaacatcaacgacaacaacataaactcttc
cacgatgtggagttacacgctagcagattattgtcctctgggctattacacccaagagagccctcaaccctatcagacatgcggcaattttact
tcgactacaaacaagagactacaaaacgtgcagccattatacttttaaacactcttttggaatactacaggacaccttcagaagagtgggaaat
tccgtttaatctcttgcttaatgtgatgaataacaagtggagtacactcattccaggtgtcaaaataagtgcaggtatcatatcgaaactcccatg
gaccatgaaaacaatgtacgagattgtttcttcgcccaataataataataacaacggagactactattctacatgcaggcgaatggtaatggaa
tatcctatcgggggtttattgcacacgcctgccataactaataagtatccacgctccagaatggtcacctgtacaaagggcaaagaccacca
gaagctatatgacatctctagacaaatgtttgatataatagaagcaaatggacaactctgattattattattacggttacgaaaattccccaaaga
ctaaaaaattcaatactgcactgatgattgggttcattttatttattttattaatcatcggaatttggggtttatttagatctcgaaatgcaaccaccca
agagagcaaaacttcttccccaacaatctcctcgaccccaactacatattctggcagctcaaccagcaggggtccaggttctggatctggaa
acaaacccaaagatgacacatccgttgaaggaatagaccctggcttactgtaacagaaaaaagagtaaaaggcgacagctcgcttgccaat

tgtcctgttacgtactctgtggtttcacgaggttgtcatcaccaaaggtaaccttttttttgtcctcgccgacaaaacgacatcttaataaccaag
caacgttcgataaagaaaaaaactcgtcatggatctttctttcactctttcggtcgtgtcggccatcctcgccatcactgctgtgattgctgtattt
attgtgatttttaggtatcacaacactgtgaccaagaccatcgaaacccacacagacaatatcgagacaaacatggatgaaaacctccgcatt
cctgtgactgctgaggttggatcaggctacttcaagatgactgatgtgtcctttgacagcgacaccttgggcaaaatcaagatccgcaatgga
aagtctgatgcacagatgaaggaagaagatgcggatcttgtcatcactcccgtggagggccgagcactcgaagtgactgtggggcagaat
ctcacctttgagggaacattcaaggtgtggaacaacacatcaagaaagatcaacatcactggtatgcagatggtgccaaagattaacccatc
aaaggcctttgtcggtagctccaacacctcctccttcacccccgtctctattgatgaggatgaagttggcacctttgtgtgtggtaccacctttg
gcgcaccaattgcagctaccgccggtggaaatcttttcgacatgtacgtgcacgtcacctactctggcactgagaccgagtaaataaatcgt
gctttttatatagatagggaattttaatattacaacaataagaaaataaaacaattgaggaaatttataccatattttattgacctacttaaccttctt
gctatacaatgaatgtttaggtgactggaaaagtttagcaatattatccttgaacgggaaacatgcaccaattacaggcgcaatttcatacgctc
tcggcctattggtcttttcctggtcatacattttagatacaatagacaaaaatggaatgtttgtatagatagaattggcagacaaatctgcagttct
cttaatcaaaatggacaacatgtctattaacaaataagccaacccaaaagtcatggcagtttctgaacacaactcactgttaataaattcagga
gctgtatgaggatggttactaaagaacctctcatcagttccccaacatttaaaattgtagtactttttacatggtacaattaaaccaaaatcaatca
tcttaggttgaccagttattccatcaattactatattgtcactttttatgtccggattcactaatccttgttgagacaaccgagtgacaatatttacaat
ttctaccaaaacaaagggcaattggtttaacattctctccctcatttttccaacgatagctatggctgaaattgaatccgtaatgggtttcttgcatt
tagattgtagaccttcaggcaggcggccagtagcttgaagcatcctaacaccgtacagagtatccctcatcctacacaagctgccatgatctt
gcatgagctcagcggctggcttggaggaagtagtcgaggggttaggttgctgtacaacattctcgtccattttacatttcaggtctccatttaat
acatcaatcacagaaatgcctgcaaaatccatctcgacacaaaaaccttcaacacacatctttctcacacctactgcgcctttcattcttccctca
agagcgtgactaaaaacagtttcaaatacaaattcacagtacatttccttgtccgtcataaacttgataacttttccagtttcaagcatgtaatata
ccccgactttaacttcgagatataattcagatggtccaatcattacgacttgagaagcatattctggttggacgtgtgctggtacaggaatatttc
tgcggtaactttcccatttcttacattgggcagtgacctcccccaagagtgtagttttcttctccaatatagattcacatgctccaggggaaagctt
aaaaacattctttgccatgtgatcgatgctcatggtttttatccaccctttaccattattcttgcataacctgttccagttcaagagagggctagtgt
ctaaaacggataggtttttactaccaaacccaacaggccaccaagatgaaggggggttgtttagtacggttgttagaccacaaaaacttggac
gctttttcctttgcgtcaatacagctctgttttatggtctcgtagttggacgacacatcttgcacgtcagaaatgaggcctaattttgcggcctgtct
aatcaaatggaacggagattctagaccggctgcaatattggtatcagcctctgccactgcaaagtaacacgggggtttttctagcttgaccca
cttgtcaatggaagaaaagttcctctttactacatttgtttcagtcacggtatcacaagaagatttcaagtgttttggccccttgacatgggaacg
gaggacagacagtggagtatagtctctagccttcaaattgacagtagagaattttgcacaatttggatgagagaatctttttctgtactgggtgt
cggcattggcgcggtgcttcctcaacacagacgcggcactaacagaaggcaattgctgcttcttgataatgtctgtcgcatacgttggatttaa
tgttgcaaattgattttcatcaatagtaataggcctcttcttcttatttctattaaagttgtaaggtgtgctctcgtagtcctcgcggagcctcaaagt
cacaggaggcggtgcttttcctcccacctctccattattcttctttagtctcttcatgggcctaagagaataaggagaatctgagggaggggat
cggcgagtagcgtttttagagcaacaagatttcacagctatggctgaaggtatcttgaattgccctcctccttctcctccttctccttctcctggc
gttttcgattggtaaagtcccatcacggttgctggcgtaagttttgtccgttggtccccaccctccattttaacgacgtttttttatactctgctcggg
ctctgggtcagagggggctatagaaatcactcgtggatagttcctttatatatttaaccaatttttcaaacaggggttattaaaactactcttctttc
tgtatatattaggtacaaacacacaatacaggtcaggcctgcaaaggactgggtatttaccagtccgaatataacagtcaaaaggcttattttc
cttgatctagagctggaaatgagtgagtaaaaacttgctgagacatatactgtctggccgccccttccttattagcctgaatagaatcctcgaa
gcttacaactacattaaatttccccataccgttctgggttttcaacatatttgcacatccaaacgttacagttttcatggatactacaccctcgtctc
ctaacaaatcacagttcctactatttttgtcagccgcattcctttgcctcttgacagattctgttacagcgtcctggattttcttggcccacagagct
ttaatagatctctcggacaattgttcaccataccctaaaacagcacctctggcagtaatggggcagtaattaaaacagacaacagaacctcctt
ctctccaaattctggtagatagattgtccgcttctttattcaacaataaacttcttagacagtttgaaataatgtccaattctttactcttgtgggaaa
agggtcctccaataaatcgaccaaacacacccgcaccagaagaaagggtctcgttagaagatggttcattaaacatcatacaaaatttgaca
tcagaaggaggtttcccgctcttacaaaaatcccagaagttagcagcgatgggtaaaatctgtacagggttcacctttgaagctctaacttctat
tacagcatttgatatgtcagtttccatggccttcattaatttggggatattcctattaacaaaatcacacataggtattttccattcagggtggatgt
ggggtgaaagaacaagctctagaagcactttaaagatgtaatcgccagtgattgttgtgactggagtgttgattgggtatagagattcttggtg

cttcttttccctttctgatgcttcatcacaaacatccatgccttcacagtcatcgtcatcatcatcatcatcatcttcactgtccgacatttggtgggc
actattattattagtgttcttgacactgttaccaattttcttgtacatttttcaatagttaaaacattagcattatcagcactctcattcttcattatgttctt
cagtgaaagatccctctccacgttgttgtgtttggacacgtaagccttgacgatcttcctccatgtccttataaagtgtatgactgatgcatatttct
tacccgtaacaggtttatgagtcagcgaggcttggtagagcttgaaggcgtttgtccagtgattatatacagtcgtgtacaaggcctcgatggt
ttcctttgccagctcaggggatgtagttttactcccatatttcgaatccatatgtagtagttcttgtttgttgtcgagaatgtccttcaataaagggtc
caacgtactcttatttaaaattttgtgttcgaattcatatatcagcgaacaagttttcatgtctttgtatgacacacaacatctcacaagacagatga
ctgcagtcctcagaaacgtatcacttttcccacttctttctttcttcgtgtcaagaagaggtataacgttcttgtaccagtaatcgagtatagattct
gcattggctgcgatggtttctttaacagcttggggtaaattagatacagttctgtcctctccttcaaaaatcactttgtcttcttctacctggcggca
ttgtcttttgtttgtgtttttgtttccagagagagtaattgcagaatttttctgttcaactaccagaactctataaagattttccactttttggaagagag
atttctggcggtcatcttcaacaactggagccatgtctgtaccggagccaactttgttcacaactaaccgctcggtgaatgcgtcttttaaagcc
attacggtccatgccatatctcaaaggttgccaatgaccaacataaatatcggcacagcatacattgaattgcctaatactataaaggaaggac
tcgataacccgaaacctatatactttgatcctttatctaaaaaatataaaccggatacttgcactgacgtgctggactcgttggacgtgtcgctag
aggactatagaaaaagtataggatggcgtcctcgggggggattctttacaggaatagatgaccttttcaagacagtgattcaacaagaaaaac
aagagaaaaataaaccaactcaagcaccagaaacagaaccaaaaccagggccatctcaagctccagatccagtcccagacccagttcct
aaaacaccaaccaatttctgtcctcctccacctaatcctctccctcctcctcctcctcctcctaagccctcaagagaagaacggctaaaga
cgtcaaaaatacgtttaaacaaagctcttagtgatattgttgaagccacaaacgagcgtgttgatgcgttgaaagagaaccaagcattaaatac
agaatatgacaagaaggataattacttccaggttttaaagtgctcgataacaccttctgtaccaacagctattataggcgcacacgtgaaacag
gtggccaaaagtagcgaaatcgaactggccgtgaacgaactcgatataaaaaataagtgctctttagtgtacaacgaaaatgagtcgttaaa
atttttcagggaccatgagaaccttatactacaaattgccgtccagttattctctaggcacgataacaccaaatgcgtgggggcagaaatatgt
gttaaaggcaacgaaaaaaacaagtttgttaacaaactggtggtaaaaaaactccccaatgcaccatcctcatcttcaactgtgctagaaatta
gaggcgctaccagaaatttactggagaataatttcaacaagggagaaaataacactgtcaatgaaaacaaggacattcctccttcagaacga
gccaacctggacacgaccaaggcagaaatatcgcacgtcttttccactctacacagactggacactaaaaggaagcttttctttaaaggcaa
cactttttatcaacgaaaaccaacattcgataataaattcaggtggacagaagttatagggtggacagaaagtgaagcatcaaaacaaacca
ctaaatcgctagacaagccaacggacgacaatttattcgtgctaccccattctttcaataatttggcagaccacttacgtttgaaatttaaaaacg
tcctctataaaaatagtaccgcacatcccggcaaacgaaattactacaagactcaagagacgctaataaatccccagattgattcggcgaaa
gagtacaagatggtctttgcagaaatcgacaagtgtttggatgttcttttggccatagggaagaatgacaaatacacaaaaagcactgtcatac
aatatagaggaaagtttagaaggtatttaatattctgctacgccttttatgctctaaataaggcaaaacattctcgcgcagtatcccctctaccatt
taatttctttaaccttttctccttcatgtattgtcatggtccgtttctccattccgccagtttttttgtccacattgacgttcgtctatcaacacatgtttttc
cccatgggcacagccgccccatccgtctcagccaagcggctcatggatatcgattccgccctaatgaaaggaggaaagggggtgggtgt
gagggattttggttcaccttcaaaaacaagtctccatacaagaacattggtgtctttcttaggttttgctgaaatggctatgggaacaatgacgg
ctctcttatctggtgtagaagtgcgtgtatctccagctctccaacaaaggatatctaaatccctagaaagatggtgtgattcagtcatctttatata
tttcacctttgttttattccacagattcagtggtgcgaaaaaagtatcactcgaatcggcgcttcgcctcatcatggggcagacgcacgcccac
acaaataaggtgagggccgccaagagatgccgaatagaagcagcggaaatggaaggtgtggaagaagaagaggcgggcctgacactc
tcttatgcccatctattgggtcttccttactctatacaaaaagccctcggattacctgtccctaagataaaccctctcatgacagcatcttcttctca
atacaatttaggggattttgtaggcgtggaacaacttctaaaggctaagagagagtttccagccgaaggagaaaccgcaggatttctcggca
tgtttgataatctagtgaaagattctattgacaaatactacggcgaaggagccttttcagacgtggttgaaaatgtaaaacaaggcatggaaca
aaacacaccgtatgacacatcttcagcgttgatgacacctatccctaaagcattctacgaagaagaaaaggatgttccacagcaggaagaaa
attctacacaacaaagatatagattgaatagagacgtggaggaatatttaatggcttctcctatgaagatggtgtttgtgtctatactcgataaaa
ctaaccaaaaagaacgtttcatgtctgttggggatattgcccttctggccgtgtggtgcaaaaggaacgtactgaaaaaggattggaacgaat
acgctatcgctaaaggcaactacgaatggcttggtgctaaaatgtgcaaccatttacttttagctgatttagtgaattttggaatattaggtgactt
gaaaataaccaataaacttgacacaaataccgacacctttcacagagacagtgatagattaccctcagttgcagatcagaaaaaatttataaa
aaacacatccctatctgatcgaaaacaattggcccttgttcactcgtgcgttaacgtgagcacccgaacccacgtaggaagagtgactgcaa
catcatgggctgtcgatgcgcttcgtacctatacaagaggtgataaagacatgtttgccgccctatcttcatcgctggatatgtaccatcttggg

cacacgaattcagctaattttgttccatattttagtagaaattacctatgtaacgaacaagagaatggattgtgggggtatactcgcagaacctct
gaaaaattggccaaagaagaattgggaagaggacgtttagggggcctgaataaggtaggggtggctaaaacagaactggctgctgcagc
cattgcaatttcttctgccttagatatgggggaagtagaagctgtaatggacgactcttctaaagtgagaaaaatagcctccacctgcttaaatg
ttaatgcagccaaggtctcggccgccagagaaaaggcgagagaagctagtattaaacgtcttcttctggccactaatgcaccagcagctgg
ttcatccagaaacagtaacaggtttctcctcaaagatttgtgggggttcttttctgacccagacaagcgccagaagcttataaagggtgaagca
gtttctgtactatgtcccaatacaggatttcttcatgctgctgttcctgattttgttattgagtattccttcgaaagtgaaacctctatagtgagattac
gtttgagactgattaaacctgaaaaacaagacgaaatggtatgcccttcaacagctcccgaagctaataagaagaggaaattagtaaggaat
aatcaagacgctgtactgacgttggatgatgaagataacatcgttaaatacaacaaatatgatatggttgaagacgaggaagcgcgtgaaag
attacgccaccaggacaaacaatcggttattgcagcccgtatcagtaaagtgtgtgagcggaaaaatccaaagaaaaaacgtcgtttagaa
gaccctgaattgcaaagtgtggatgaacaattgatacgggaactggctgccattgcctactgacgagatctaagattttgtaatatattatgtgt
cttgcagacctaatatcctgaggattaataataataacaacaactgttcaaaaatgagagacgatacttttaaccaagaaactgcggtgaaact
tgtacgatggtatacagagtacgattgttgttgcccattggttaaccgcgtggagcgccttctaggatcgttcggaggaggcgtggacgcca
cgtctgtgcgaagccgaccggctctttatgaagaagataagaagggagataaatgcataccctttaggataacgtcccttattgagggtatac
ttttggaaagggctctaactaaacccgatttagctgctgcagcttttgatgtatcagaaaagctggtgtattgtagttgtaataacactcaaggca
attttgatgtctcttcaatgaccatatggattgatggcaataatagtaaaaagtatgaagttacatgcccgtcatgcactgtcgagaaaattagtg
gaggtgccgaatctattcacaagaaacccatgtctcttcttgccttctttaacaatctggtagagaaagaagccttcgccgaaagaattgaact
caagaaattgtacctctccttactaacgggctcggcagccggaggaggaggtatgtacaaggacagctcccaacaatcttccttcaacggct
cttggacgtcgttgttgttccacacatctaaaaaggacaagactcgtttagaggctgaagttttagtcagtaacaagataaaacacacatcaag
attgcagcctaggtgcgtctgttccgatctgttatatgccctatgctccaccactaacaactctgcatcttacgcgtacaaggcaagaaatttgt
gtgttattgaaggtggggaatttttatattttaaatacacaatctttgaagagaatggacctttcgactccaaaacagaccttcaatcattagttaat
aacgagcctgtttctgagacaaattcatcggcactggccgcttcttcttcctctttagaagacgacgatgattgttgtgatgatgatgacgatgat
gatgatgatgaagacgaaaaaactaagaagaaacaacccaagaaacaaaccaagaaacaaaaaacaacaacatcaacacttccacctat
cagcaaaaccaatcacgacaacatgttgatgaatgtacttaaaaaaggagctgttaatggaaaacggaaaatgatggattctttgtcaggaaa
aaagggccaacactctaaaaaattgaaaacctccgctgctgctggtggtggtgcttcatccgacgttgttgcaggagaaaatgaggaagag
aacaacccctcttcagtgagtcctactaacaatagggatagaaaagactatgtgcttccatgccctcaaatagaagaagtcactatttttcaca
acacaggatgaacaataacaagttggcagaaagtgtagtcaaacattctgttgttattaatggaaattgtttaaacttgtttgttactcaacacag
aaaaaagtatatcctgcctcacgaaaatattctttttgcccacctttagtccagcatgtaggatttaacaaatttcgcattttgactggcgtttcttg
ctttttgatagaattgaaattgtttttccgaccaatctgactctgtggtgttgagtaataatgctgcccattcggctattctaaggctattgtcatac
ataagagaaaactcattgaagcgaagtgtgaggactgcttcggtgaaaggaatcgattttgtcgtgaaatcacaggacactaatataggcata
cctttaagcaacaaggaaataagagaacggcaattatgctcagcttcaaccctgagtatgctagctggtttgggaaaatgatcaccgacccg
ggggtcattttacccgtttccaaggatgttgtactctttggatcacgaggacaatcagatgtgggaataatgacactggaccctcacgatttgg
atataaaaatcacatctaaacgcataggtgtggaagaaagactagctcaatacaatactctacctatggattttacacgggcaatggaaaagg
aactaaataatagtagaaatatgaaagagtcaatattcacgggaatatttttagacaccggttcggcaatcttcgaagacaacatgttcaacgg
aggaggttcagctttgcgcttaattagatcccccgctttgaattctgctgtattttcaagcaagaactacatcatcaaacaattgcccaccataac
caaatctctaaggagaagtcaagctagagataagcaagtggataaaacaagagagaagatagtggtggattctttcagcatacttagtgctat
agctgctcaagtaatgcacctcacagacggagagatgacgtacgtccccgatgggcactgcgttaatgttgtcatgtcagagaccaatgctt
cgtccatctacttgatcataaacgaccccactggttcgggatggaaaattatgcctaacaatttcaataaaacacttgaaatgagagacggtgt
aatagatagagtagaaacattagtggagtttgcgtgcaagtgcgtcgcatcatccttgattaaaggggcatggatttagtggatatgcaaag
aactataaggtctatggatttcctccctccagcttcttctacttccaataatactcctagagtagcgataatgacatctggaagtagtactactacg
ggcattggatccttgtccattcttgcagaagatggatcaacacaccaccaaatcaagctatcggaatataggactggattatccattactgaaa
ataatagagaggtgtcttttacggtagaaccttcaatagacggcgttcaagcagagcatcctctatccccttctattcttcagtggttacctcctct
agttaaaaggccagaagtggtagcagcagcagcagcagcagtagtagaagaagaaaacggggacaataaaccttctgataaagataacg
aagacaagtacagtgatactgatttttggtctaatgtccccgtcacacctctaattacacccaagaaatggagagcgtgcaaaataaacgatc

gggcaatgattagtagttggaaaaataatctagtgaaactccacaaatatgattggacaaataaaactacaaaggttgattattttgataagatg
gctgcctttgttgccctcatgacctttagaaaattccaagacatactagcggataactatgttcctcctcaaacccccttctcagggaagtgaata
cgcagtgaccatgtctaacgtggctacactctttactgacgtgtacggttttgaatcgaatggaaataagccattgtttgccctagaacagctag
aaaatgaaaccgggattgaaagcatatacgtcctaaatatcataggaaattcccctgatggtaattctgtcagggtcgtcagactggaaaagg
aaatgagtttcctcttgaaggcgaagcagtactttacagaaatggccatacctcctattaatgaaaaatgcaaatggacagataaggccccgt
catctgtaaaggagtacaagtatttttgtgatctaacagcacccatttcaaagagacctagaaaagataacaacgacggcggtgtggagcatt
ctgcgttgacttatacacctaggtgcatataccacactgaacgttgtttagtccatctttactctgagccagaaaaaataacagaacacgtatctt
tcaacaaggatttgaacatattagaaattggaaaaaatattaccaaccaataccaaacaaactacaaaagcatattcgaaattgtggacgttcc
cataattgtcgcctctatgtcatcaacaaaaacaatgactgtaaacaactacataatttcaacaccttctgccacgaccaagtttgttcaggatcc
gccaaaaacagggaaacaacttctggcagttgaagaggtcagaaactttaaactcaaatctgttcttgttcctcctccttattttagggacaata
agcgcaacacaactctttgttcacaaataactgaacaaaattgcccgtcatcttctgaaggtgggcgtttttcatgtccatcagagtcacttattct
caagtactctaatctctctaaaaagcgcgcactggaagaaattgccccagagactgagactagcattttgtcactagccatgtaacgtgtctcg
atatatgacaatttaaacaacctcttaataaagaggataattatcataatagaaaattaggtaaaatgggagtacagaagaatatcctggtcggt
ggtggtggtggtgtatcattgttattgggtgttgttacacttctaggaacagtaacagaaggagcaccggcagtcccaccttttttcatcgtcttcc
tattcctttactcctgaatcgagtgttttctgggtcgaaggaaatcgtgttttaagtggaaccaagaaggacaccttaatcaacgttctggggaa
gaagattccttattatgctaattccatattcagacatgactgttctgaaactcgttctattcaatggccagaaacttcccctttgggcttgaaccttat
tttctgttcatgtgcgagtcatgaacatcaacaccgtactcatgaaacaacagaacctgacgatttattgtgggacggatcaagaaaaactacc
accataattctacctaaaaaatggtggtctgatgttgtgtggacatctttatggagggataacgaccagaagtgtgggtgcgggcaggcattt
gtttcttcctttacttctacgcagaaagaagtccaaggggaatggcttgctgctcacactaacggaaagacatctgagggtgatactaattcag
cctaccttttcataagccttcaacgaactacactcaagcctatcatcactgacgtcacagaagataatatgatgatgggaagaatgtcaggcac
acccatgaaccctaaggatatgacatattttgtcaacgatttttcagacgatataggaagtactcctcagtgtcttgtatcgaattcggacatcct
gaacaagagggaagaatggatagctgtttggggtgttgcagactctaaagacctcctaactaaacatcaactgggggagcgggaatatgg
gagcgaaggaagaagaagaaatcccggtgttgaggaggaagaagaagagagagtggaagaagaagaagaagtagaagttgcgctacc
ttacattaagaaaagtggaaaacttatcggacctcgtagaagacctttgacaacaacaacaaccactactactactactactactaatcctattg
ttagagaggttgtggaagattttgattacgagtcttttaatgaaccagaaatctttggcagtaactcaaaacttcccttcattagatttcttgatcaa
aaaaattggagacttggtatcatgagcagagtttcttcttccatcgccaactttaaaattgaacaagagtcatcaaaagccttattttgtttggcag
tctgggttggggatgaacatacccctaaattcagacttagtgtatggaagaactggaagccttttacttctgcacctattattgtccagaatgtag
gttattcctctgatgttttctggcatgaaactcttagaagcaaaattgttgatcggtcaagggacctgatagaaacaaaagtgacaaagaaaatt
ggggaagattgggctaataaaaaacaaactgtagttgctatgtttatttcaggtattgtttgtataacagtaacagttatttctatattttcaattgtaa
tatattacaaaataaaaatgcctaaattctaatacaatggtttattcattccttattcctactactagtgtatagcacatgaaacaaagaaagagac
acgcttcccttgttccttaaacaatatgtttgatacatttttcacaataaatttgtcaactttatccattacttcattaccttctcctgatatcttttccaaa
tccaccacagttccaacaccagtaaacatgctctttctgatatccaatgttgaatacttggcaaattcatgcccttgcatttcagtcatggcaatg
attatagcaccatacattttacatttttttgaatagaggatatgtggcaaagttgggagagaatatctttgtgggaagaccttcttcttcctcaaaagt
acttttcagtgctcctccagttctctttacatgcaaagaaaattgtaccctttttcattggatgaacttgtccacagaacaggacatcttcagacatt
gaaaagtggagattgttagtaggtgaaattgtctttcttggtgccaaaagactaatcgtatcagatggtgtttctgggttactttcaagaatactttt
gtccttaatggggaattcactaacaccagaagctgctgctatttttgacatgaacattaacatacaatatacattagcttcgtctgacatacacag
ccatcccttttctgcatctacatggatattttctcccttctcttgagctgtgcatgtggacgttacaatagacccatgtttcgatagaccacaagttt
ccgtaggacgacaaaatttaccagtcctacgcaagttgtgctgcatactcttacatacaacatccatgtatttattaaccttattttgattagatgtg
tcactaacttttttcaaaataccttcaatgttatttgaaaactcatccgcgcttgttataggaagacgcaatttgacatctctttctcccccgtccttg
caatgggcaaaatctttagactcgtaacccagatttggtacatgagtagaattggtggtaattgattttgggcgtacatactgattcaggcacgg
attgggagatggcaaaaaggcgaatgaagaattcaaaacgttcaatttttcatacactaaaaatatataatggacataaaattggaacacaatt
gtagccttgttatggggtgcaatatcaaacaacaacatttcaatagtggactcctttggtgccctattcacaaaaactgggtgtattttgatatagt
ctttcttgttcttttgtttcgtctcatattcttcacacaactttgaagagggctgttttacaataccactggctatagagtagtgtgtgattgaacgaac

gatagcagtgtatagctcaatcattttcttgttacataatttctcataatgtttagccacatctccattattaacaccaaagaaaaagtctctaatgttt
ttaagttctgtttctagcccatgttgctgaataaccttagcagtgttggccatccaagacgcattaaacgataaagattttcccgatctacttagga
tgaattcttcagcggcagctaccacatctttagaagggtcataggcagaatacattgttactcctccatctccattatagtcatcatccattgcag
cggaagattctgccacctctctagagacactaactgtatcgtggttaataaagtcgctcccattaaaatcagcaatgcctccaataggcatagt
ataagccttgtctttactcatggcgcatttaatgttacacaagttaccccgttcaacagaatgattttgaataatgaacaagttttgaggattactat
tgcgggttacagcaacatacgcatcctgggtagatgcaccagaaaggtcaacaatagtgtctctaaagaaagtgtgcccttggcaagaatat
atcgtcatggcttgactagattcaaatggaagatattgaacgtatacattgttcccaaatcctccattcttgatatttggattaccgataatttgtcgt
ccttctttaatttctaggcatttttgtcctagccgttccacgtacacaaatactgtcaattcaccattcttgtacattaaattagtaaccactccagtgt
ccttagtcacgaacctttcctgagtgccatggatcattctattagatgtagtgaaaatcacattttggccctgataaagagtaatagatttagccct
agttaccttctttttctcatccaaaagaatcgtagttaatgccctaggagattcggctgccctcaaaattttcctgtctgtaaacttgtcaaaaagtg
gtgtactgcgagttttaatggcgttaattaattgttgccggtacttgtgagagatggttttctcgtggctaatgtcagacagatccatattcaaaac
tacaaattttacttccactcttaatgcgggcacagaattttcaactgccgcagccgctgcaagtgaagtagaagaggaggatgaggaattgtt
attgttctttttagtttctttgatgatagaagtatccactaaattcgagactgcattctttaaaacgtgtttttttagtgagtgtgattgcacccatcatctt
gtcagttttggaagaaattgcatccttttcttgtttggctttagagttggccatgaggcgttcacacatatccttaggctgcccttgttcttttgttata
aactcttcaatcttcttcctcctctcatttccctttgaaatactagtcaccatttgcatggccaacactatagatttttcagagtatttaagtgagttatt
gtgcctattgttaatgttgttattttcagtgtgagcagtttgaagcactctggaaatctttaaactacccttaacaccctccatcatttgaccattattg
aaaacatcttcctttctgtacataagaaattcgttaaccattgaagggaagatgttatcaaaatattgcactgggtcaaagttgtccatgaaaaca
ctcccatttctggatgatctaccggccattcgcgtatctaccaagtcttgaaggttggcagtcatatcaaacaatgaaataccgtacattaacga
ataagcgtgcttaataatgtctattctctttaacgacgacatgatatatgttctcgaaagggcagaaagataggaagaatatgcaggtaataaca
tactcttccctacatctcctcctgttgatatgggcatagataataccctcaaccaatattcgtcatggagaggcttatctgcccctgctcctcctcc
tcctcctactcctgtgcttgtggcccattctgagggcgagtggaacaagcgtccagacagtgcagatttatcaagagaagggagttttccatta
tcaaatttaacctgttgcattctcactagtgtatacaaataccacacacgggctcgtaccaaatctgaaggcttattttcagtgagccatttttcca
cacaacatgaaagtcaactaaatgacctacacgatctagagtagatgcgtcacggtgcaatttatctggtatatcagttctatatatcatttctcc
actttcttcatcatcgtcatcacctcccccaccactacttttcataccagaacctccacttcctccgtcttccaaatccattgcttcttcttcatcgaa
ataacaatagtcctcgtcgtcctcataataagctgctgctgcagctgtacgtgatgatgcactattattattaccagtttctttcttcaagagaatag
atcttatattttcagatttcgctagtagttctttgtatgttaaagtttcattctctaaaaatgaagcaactgtacgcaaattatctttaaagtaattagcg
ataggagtttcatccacaaagtcttttccatctctaaagaaggcgttcctcacacacgctgccacagcagacgcacggttatgtgcagaattga
atggtactgcatctatacacttgggatcagaaccgtgcacatacttgacgttcaatatctcttcatccattaaggaggtcagatgtgtatccttaa
atctctctgataggctagaaacaatatacaatttttctcggccactgtaatcaatttctctcatggctttggccatcatgtcgaacacaatactgcat
acattttcagtgtacttttctagatttgcattgctcattttcttgagttcagatatgtgagctagctctgaacatattccttcttctattagagcggaaa
gggggagagatttcatcatatccaaagaagacttcttgtctgatacagaagttgttccacttcttatcttttcgtttgcatcagaaatgctcttgcag
cgtaaagagaatccattcaaagctcccgatgctctgtccttttcaagtgcatttctttccatcagtttcaaaaatgcttcattgttaaagaaatcgtc
atcttcttcgtcatcaaattcgtcacaattttcttctgccgtggattgggccggggaagaacttggtgccgtcacttcctttttattagaagcattac
cacagtcttcaactttaatagcacttgagactaaattgttgttaactgacttgacgttattactcttcattaaaattgcttcatggcgtttattgacaat
attccgcaggttacttgggttgccaaaaaagtctctagagcaaaacattttcagtatctttataaaaacttcagtgtatagttttgcattatcttcttg
gccaaaattatcttgttcgactgacccttggaagaaggtactcttttcttcctctttcttcgtctcagcttcatcgtcactttcttcttcttcttcctcctc
ctccccctcttcttcttcgttttcgtctccctcttcttcttcaccacctttctctttcttctctttcttcttcttcttgttcttgttgttcttgttgttcttctgaggt
ttttcttcctcactcgggttctttattttcttctggatatggttagtgttatcttccactactgcagaattttgcctgttatctccaagtagtataacaaac
actcctccgaaaggtatatctggcctgtgtttcgctatacgtaaaacagcatcaataaacaccaattccctgcaactggccattgtatactcatc
gataacaaggaaccttaaatttgccaacacttgtatcactttccattttagtttatcggccacaaattggatttttttctagagttatttcatctttcctaa
atataactggaatctcaaatcctttgttgagggtagaaaacacactttctgggtacatcaaggaagaacctcctggtttcttgcatgcacacatg
agactctcctttgcagaattagatttgcaacacaggagcatgtctgcagatccagaatcaatcaagccgtggtcgttaagaactgcaattagtt
ccttgaccattgtagttttcccagctccaggaagaccacccagatggaacacagatattcctcctccagcataactgacagaaatcgagttgc

acagatacctcactgccagagactgttgcaaggaaacagtgctcaatacctttctggcctccttgcacgtatctgggagcttgaccgcagaa
gcagaggaggaggaggaagaagaagacccagaaggggatgccgctgcagctgcaactgcacttctgagcacctgggttctggtagcca
gaaaggggtatgatttgatgttgataactggaactggatttgaattattgttgttgttcttcttttgtttttttggtaactggctcttggccgctggagca
cggacgtttactccccatgatgcactctcggtgaatgtcggacacgggggtaaaaccaatccttttatactagttctacctccacgaggttgcc
agatggtacgatactttattaccaaaaatggatcgtcctcttttgcttaagattttgtgtgtaacccttatcgttacgattgttacaatatgtcttatgg
catttttttatacatagcaattcctcatcttcttcaggcgccactgccgccgctaccgttgcagcggcaggagggatatcttcatctgctttatttgg
tacctacatagggtccttggtgcagtcagaattttttttcagaagatatcgatttaccagaaaaacaaaagaaaaaggctgataaggaaaaggc
atcaaaaaccgataacaataacaacaacaacaagcgtcaacgccgccagacaaggaagcagagcccgcgcgccagaaacaggacttta
agaaggaagaggaaattatccaaaagaaaaatacacgtcaagagaaagaagactctaattaatcggcgcagtctcagaggaggaagagg
attaaagcgagcatctagaggcaaaggactgcttagagggattggaaaaaggacatttggattgtttttcctccccttttagaaaggatgtttctt
ttaattatcatgttgattatattagccccattaatatttgttggggtgaatttcttcgtatacataaagaagaataactggttctctggggccattagta
atgacactactattaactccacaaattctggaggagtaggaggattatgggataatgtttctgattcagagtcagggacgtttccatctagggta
caatattccacatcaacgaaaaatttagtcaatatagccatggatggagattatgcgacattagtaaggaacggaatgtccacaaatcaaagg
ccatatgaaacctacaaagacgtggaggatagccagttctatttacatttttttccacgttagaaattttaaacctttaaacggtgatgaaaataaa
gaccaccttgaaagggatgaaagttttgtgttgatcgaatcaccatattataatggagggtttttatcatacaatatcaacaacccaaatcccattt
acaattctactgaaaagccgtatattaacacggagataacttccatcgtcagcaccactggtacagatgaaaggttcttctgcctcgagaagg
aatacgtcgaagatggtgaagaaggagttacagaaaataggtactttttacgccacatggcaagtaattatgttgtaaaggctagtttcaaatc
agtcatgcccactattgaaattagtgatctgactggcacatacaataataagaatagtgttaaaattaagcctgtaactggtacttctataatttat
gaaaaattacaaggaacaaaataaaatacattttaaagtattatgtttattgttttatttcttctaaaattttattttaggtatcctcttgaatatggaacg
acgggtgggggggttttttcttacccttctttggtttttggttagttttcttggccccaccatctggtgtttttggtcctcctttcccctttccttttccaaa
ggctttcttgaatgcaccgcccactcctggtattgatcccagcgcagaagcgatcgtgccggccagtgctgtgcccgccaggagagatgct
aagagaggagccttgattattattacagcaataacaatgacaattattgttattaaaagtaaaccaaaaagtatatttttttcctcctcctaaagacg
ccaccaatggcgaagccatcttggctatcttattgttattattgttattgttattatccccttccatggttgttgcacataataatagtgtgtattataac
attaaaagaaacatataaattttggctcaaggcaccttcaaggctgtgaaaatattacatgcagataatgttgattctgaattcagggttgtgattt
ttggtcgcacgtttactgacgtccaacataggcatatggatagtacaaattgtcttgaattcctcatcttctttgtcctctgtaccatcctggggtttt
gctgggaacaatgtcctatatgttccctttttttatatagtaattattcgggtttgaatacaagtagttatggaaatggagatgcgaaccactaaaag
cttgtatatttatcaaaaaaattcaagtcaactttctcgcactcttctcattttgggtaatatctctcccgttaatgacgtagataattgttgcattcaaa
gcactgatgtgcttttcatgatattcgtttaaaacaattttataatgatttttaaagggataaaaatttggttacaaatagttaaaaatatgtaaccatg
cagtacaataaaaggtgatcaactggtttttcatttctatatcgtaatgaaaattttgcattacgcgtattctttgtaactatatcggaatatacttcat
cccatacagcagatgaacctctttttttcaggcttaattgcgatatccttttcagattccatttcgtggaagatctttacaaagcaagatttatccgttt
gattattgaacaaaaaaatcgagccgtttttttcccattatctacttcaaaattctcaattgttagagaaaaaatccctcttccagactacaatatcttgg
gccttagccagtgttctgattagcccgtgatttatttttcctcccatagaaggagataaattgtagctggtttcatacctcccttggtcatcacatttt
actctaaaaattgccactgtaaaatcactcatgtgaagattttcaatcaattgaggaattctcttccctatatcaatttcaacgctagctggcgttttt
agtacgtcccatgcaaacgcattgaggctcttgttcggtacaaaaaaaagttaatgccgatttcggatcagggaaaacatccgcagccacctc
agttacccaggcactaatgttgtcttcagacattggtaaagttgctgctgctttttcatttattagtgctctaattttccctctgttgccctgcattttatt
attgaatacagtctctccgtagggtttaatatgtcctccatacaaatctaccccagccatggaagcagttgcagtatctgacgacctcgtgtcca
aaacatttgtcaccctagaggaaatttccccatctttaaatagttctgcaagtctcgcctccattattattatttttaatgtgtacaataaaaacaatta
atttgaaacaaaaggtaataacaaaataaaatacataagaaagattcaaaaagttttattcattatccttatcacacacatattttttgacggacaa
gtacattaacattatcaacattcaacagaccactaccatcattcattacaacaacagaaaatggggaagttttgttgcacaagaacttccataca
tcattcttctttacatgaatcaagaaagtattccaatcatcccattgctcatcgagcctgtaccccaaccatgggtttacaaatagactattaaattt
atcccatgaacagtattcctggagagaaatgaaggggtctttcccgtccaggtcatttggtccaacattttccactacctttacttccccgagtcc
aattagattgactgtcgtcttttcttttgtctcgatcgcacactcgccaatggcagccaacaaggaatcgggaagatcactcctcattttctttggc
ggtgaagaaggagaagtgacctgaagaggcgctggtctcttattattattttgttgcagtttatttccttcggcaattgcagtgttcttgagtttcaa

gaaatagtcgataatgtaaaactcttcaccctctcccagttcttcatacatcacatcaccagatgtcctgatggcttcttccaccatagccaaaga
aggtgctccacattcaatcaaatccatcagcagtttcttccctcctctccccattgcgacgagtttgttggcgtacctgcaccagggacattttct
gttctgcggcttgtccggaccaagagtagacatcatgtacttggatttgcagcaatcacattctacacatacagcaactggaatatcaatattgc
cctcattcaatagcttgtagagaagacaagtcttcccatacaaaatccccacagccttcgctgcagctgaaacagcaatttcattgtctttatctg
ttgggggaagattgagagctccaatgatatcactagccatctcctttactttgtcatcaattttcttggcattcaaaagattgagggttgtgggtga
tgcagttttggcacgaggatctctggtcaatttattgtcccagtgatagaccattggtttgcccatcctgtagtttcccttgcatctcaagcacaatt
tttcatcattatcatattccatctttagagattttacaccacatcttttgcatatatatttcaagtccttctctgataggtcaagtcgacgtacagggaa
cgtctcatatttctccccaatctttgagtcgccagcctttgtcaagattctcagagccaaaaagtgttctagtttattggtaaagttcttgtcaactc
caatcttattggcccatgtaattaagcacctcaaaaatactacaccggaaagattattctcgatctctccatcggcagttccttcaagaatagag
ggcaaaataatatccaaatcagtctcggaactgctattctgctcttggtaatactttgtgaatgccatttttgcaatagagaggatcatgtttgaag
tgttggagggtacttcacaggaggcagcccaagaggcggctgcacacacagatttgattgctgcataacctccttctggcagactcagacgt
acatcatacaaagttcccaatttttgaatcacagtctccaagtcgataaacatcatttctgccacatcttctcgtcctccaaatagatcgctggcc
aattcttctcgaagcacttccttctggtcatctgatccacaaaagactgggatggcaaaatatctgttggaggtttttgacgtctccaaaagcac
atcttcaacatttctagtaatctttagacatgataccaatttcttgatgaaatcaaagcactgtccaaactcgaacctcttgtattcgtccagtgata
ataatcctcctcctgtatatgaattggagaaactaaatgactggtggaagattttgccagcttcttccaaggcatcattgggcacatagaaccac
ataaacatggaggattgttcaggtgtgagatctgtattgttaatcacttggagtgcatcatagctagcagatttcatgtgctttccttcttttaggaa
ggcatattctgccttgtttccgcattcagttcccagatcagaaaggatggtagagaccctcgagcacactcctgcattattgcctgtaacaaag
gcataaatagccccaatttcatcaggcacagcactgcaagcgtacatcattgattgttccttgatcatcattttgaagagtttttcagcaaacatct
tggcccagtgtacgcgagcatacttgcgagacaacttggtttttcttccttgatcagatgtagcttcaacacttgccatcaattttgtacaacccctt
acaatcacagaggtttgtatggtcaattccaaaatcgtccaacgactggaaacacgcgttcatgtgcagtgccagagtagaggcggcatttc
ctccccctgtaattgactgggccctttgagtcactacagccagtttgctgataccaccaccaccgctagaagactgttgttgttgttgctgctgt
ggattaaaggtttgcttagagcaacaaaaggcactatcatatgcacaattcaagaacacgctacgcttgtggttgttatatcttcctcctccctgt
acaactcttccaaaatctaaaggtgccatatttaaaaccctagacttgaaagattcacgttgagcgtcgccatacttttcaaatcctgcaggctct
gatcttccttcactttcaataaaaacattcaattcttcttggttgatttccctcaagatggggaaggcactaaattgatcattcaggttcttgaagag
ggaaatagattgcctcataataaaggttgcgatattcttgcgcacattgcacgatcttctgtcaaaggattcaaaaagttgccatctgttgttgtc
gtatccagaaaaattgatcacatcttcttggcacaattcactctggttactgccactacatctagagaaattgaggatgcgtagccaacaatcat
tctccactagcgccttcctcatgtcaagaatagtcaagggtttttgtccccaagtcttcatgatgaattcttcttctggtgcagtttccgaatcgtac
gagttggcacataattttccagtgtcactattggcaatacgcatatcctcatcggtgaacacaaaaatgaggatggggaatttttcatccctggt
aatgccatgcattttcttgaatgcaggaatcatgttgttaaacacgtacatcagggaagaatgaagagcttcactgccaccgccgccgcctcc
gcacacttcggtcttgcgcatatgcttcacaaaatctccccccttcccactctgaacagtctggtgcacgcacaacaggtccactttcaatggag
tgtttgtcgaaatcattgtagtaaactatagcattttgaatgcctggagcaatcacattgagtgtagccatgatgccagaacaactatcagccac
agccgatttcacacttcccattgaaccactaatatccatcaacgagatacttagaccagtaggttttgcagttggagtatctgtatcagcagcag
cagctttcttagtccatccatcttctaaaagttcaatcttggaggaaggtgcattttccatcaaagagttcaaaaaagatgcagatttcacagtag
aactagaactcttgctcatggtggaagaagcgcttgaacggatacggcccaacattgtagtatatctttatggtagaagttgatgttgacggta
ggaaacgatctcgacaactgtatgctgaggtgcctccgcaggccgcttttataccccttcgtgtgacgtcaattgacgctcgtacgtataaac
agggtctatttttagagatgacatttttcaaaaacgtttatataatggtgcaatttcctattactcaaactagtcatatccggctaagacaggatgttt
ttttggccaatacgtttctggcgcaatctgacattggcccgacccagcggtccacccttcgaacttgacatcaggccgacccagcggtccac
cccctaaactggagtgagctgaaaaatttttcaaaagtttttgagatgaagaagagggtgaaaagtgggtacgtactatacactcagagggtg
gcagatcggccctgttccagaaatggctgtccagaaatctgggtcggacagattccagaaacgtttctaaccatttctggaacagtcatttctg
gaaagggttacaatttcttataactggtgcattatttctagcatatattctagccccttttgtgcaatctgacattgggtcgacccagcggtccaccc
ttcgaacttgacatcaggccgacccagcggtccacccccctaaactggagtgagctcaaaaaatttttgaaaagtttttgaaacgaggatgag
ggtgaaaacacctgttagaaagtggttgctggtcggtagcatccgttcactgctgttccagaaatggctgtccagaaatcagattccagaaac
gtctataaccatttctggaacagtcatttctggaaagggttacaatttcttagatgtggtacaataacttaccatacatttctggtatcttcgtgcaat

ctgacattgggccgacccagcggtccacccttcgaacttgacatcaggccgacccagcggtccaccctctaaactggagtgagctcaaaa
aatttttcaaaagtttttgaatcgagtaagagggtgaaaacaccctcttgaaaggcacgatctggatggtgtcgtccgttcactgctgttccaga
aatggctgtccagaaatctgggtcggccagattccagaaacgtttctaaccatttctggaacagtcatttctggaaagagttacaatttcttagat
ttggtacaataacttaccatacatttctggcaccttcgtgcaatctgacattgggtcgacccagcggtccacccttcgaacttgacatcaggcc
gacccagcggtccaccctctaaactggagtgagctcaaaaaatttttcaaaagtttttgaatcgagaaagagggtgaaaacctggtaggaga
aggtacgcacgcatactggcggcacatgcctccagaaatggctgttccagaaatctaggtcgaccagaatccagaaacgtttatagccgttt
ctggaacagccatttctggaaagtgttatattttctcgcatcttgtacattaacttgattatggtatagtacacctctctgcaatctgacactgggtc
gacccagaggtccaccctcggaacttgacatcaggccgacccagcggtccaccctctaaactcgagcgaccctaaaaaatttttcaaaagtt
tttgagacggaggaagagggtgaaaacccttgcaacagagggggtataaagggaggtaccctcgcacaccttagacacacaacctcatc
accctccgtccaatcaacatcatcaccccatctctaaataatccatcatgtatatcttcgtcgaaggttccccctcacagggaagagttcatgg
atgtccaagttgatagatacaggatcatgtggaatgtctttcctcaattttcttcgtatgaacacttctgactactacaactggcctgccgaaatcg
ggacagaacatctccagttaggtttcagagaaaccagagtggtggatggaatgtttgaacctgtcctaaagacctttgtcgactcgtggaaga
aagagcaaggaaaagagagtttgaaggaatatctggactacaacggccaagtcatggagatctacatcgcagaatggttgagacaaaggc
cactagccttccacgtgtttacctatacagatgaagctgtcaagagtggattcttgaacgaggaggatctagatatggatactgcaaccaagt
ggatggctgaaattattagagagaagaggggcaatattcaagaaataaaagtgacccctagagtagtcttcaatggcaatgtttgtagtgcat
gtttctctaacactaagagaaacttgtataactttggaacaaactataacaatgttgtacattgtgatttgttgtgcccttttgcaaggcataggatt
gtacatttcttataaaaaataaaacaatatataaaattcagttgtatttttattgctcaaataagttactacacccaattctccccctctctagtgag
agatcccaatcactttctactgtcactgttgctgctgtttgttgttcctcttcttcctcctcttcttcctcttcatcttcatcttctttagttcgttcttcattat
atgctttaattatcctcaatacatttaaaatggtgcgctcttctcggttaaacttctgattctttccctcaccaccaacatcgaccctacgtccatcct
ggtatactgtttcaccatcaccagaaaaaattctaatagacggatttttatagaacgtacaacaatgtccactaatgattctggttctggtgctaa
ctgatcatcgccaataaaagataaaacaactccagaaacacatggataatattgcacattgaattgatcatataaagtattgaagatgataggg
taagatgttggcccgtaggcagacatgatcaaacggggcctgaaaccaccctcatctctcattgtacaataatttgaacacatgaatctgattc
ctctttctgtaccagaatataggggtgtgtcttcattgaatctgtttgtgaggaaaaagtctcgtgcagataatgcaaacacgagacacgtacttt
tgtctcctattcttcctatttcgtttccatattcttcccttgcttcttcatcgatagcatccaatttgttggtttcctctagtttttcttctaggtatttcctcct
tgcggcgtcaaggtatacctcgtttggcatatggatgaccttattacgttcatgcttaccgtccattttttcggccgaatgtgtaacaactgcattc
aaaaataatgccatgcattgatacgtctttagaatgtctcggaatgtcatatttctttgtgcgtagtacacaaaatgtggcacttttcaagccacca
cctctacatcccagtgaggcggtggaggtggacgtcatatttaactggaatatgtgtaataatatagtccaattcttaaaaaatcaatagaccaa
taaaacaacaaaaaacattttgtaatttatttattcaactaaaatctcatcaaataatcttctcagcaataaacttacatttcatatcacagtgtacaat
actattcgtgcatttaccaacacggtacacgtaattcttactcgtcgagaaacatgaactacaaatattccctttgtacacatttctggggataact
tcaatctttacaaacaacgctctctcctccttgagaacgtcaataaaccagttatgtgcatcattcaattcttgaataattggttccattttcttgaca
tcttcttccctgtaaaccatcacatggaaaactggcttgtatttgacccactcggcgacacatgctttgaattcttctccatacatctccaaatact
ccctcaatccaatttttgattctggaaactgttctgtgtcgtgcctccacatgcctgtgaaaaagttcaatgatgatccaaacattccatccacga
gcctgggacgttcctggtaatcggatgcacgcaaatgctcttgaatggtccatgggaagatgggtaaatagtccctataattggtgtacatga
agttgaggaaagactgtttgccctttccggcagttctcatgttatccacccatgaagttttcccagttaggggagatccttcgacgaatacaatc
atgttgaagggtaagacttgtctaactctctcgagcgagctctctctcttttatagaggcaaaaatgtagttgccagccaccaccttaaaattcgt
gcattaataatatataatttttatattttgtatcattcaaccgtagtccatctaagagtcaacatgtcctcctcctcgtcgtcgtcgttctcgttccgca
tctccacctaccagacctttctcaaggctctggctcacccagatctggtggacaagattacccagaaatgtgacgagacaggaagaaacca
gaagtgccccatccagtttctggccgacatctcgcacctgatccaaggagaaagaaatggaggaaatctgttccctttgcacccgttcaaga
accaaccacatctggaaccaagaatagtgggaagtcttcacgggagaacattggacaatgacattgaagaatcatactgttattttgtcaagg
atctgtataatggagtattttcctatgtgaacggcgtcaaggagttacagggcgttctggacaagaaaatatctggatctggatctggagaatc
ctcctcttctagagctcccctgatcccaataactgatgtggatttgctgtacattttcggtactttggtagttcttccccccagatctaaagcgtacc
gagtcatcactgaagctgttctagcactccccttcaatgaattcagtaacaactggcctcctacaaatatcaaaggagcatacgtgtctagaga
tttcaggatgtttaatctgttggccggtttagatcatatagaaggagaagttgggggagaaagtgaatgggaatccatacacgcatctgtcgtc

aagcgaatggtcaccattatgcgcaacaaagctgagaagaaacctccatcaacatctagaatttttagagtgtatgtggctgaaccagttaat
gatgcagtgacaaagatccctatacgtgtactcagtaaattattcggttcaagactcgccggtattctccagaaagtgtactcttattcaatgcta
aatcttccatatctcctgtcttcaaattctatagacatcaagcaaggagtaaaaggaattacattatctataccatctgcaagaaaattgggattct
atttactccaaaaggatacaacattacaatcttccctatcacaagatgttgccgactgcatagtttcaatcaacgctggtattattggtgatgattt
ctctgaaaaaatacgacagtgcattgaggagaaaaacaagccagaaaactgttgtatgtgcttttgtgaaattgacaagacgcccgatttttctt
atagtgaacatgtggcaaggcacaatttcttccccgtccacgcattctcctcatcacatgatgacaagtgttgtggagcaaagatttgttccgaa
tgtatattcccgtacatcatttccctgtatgagaaaatgactggtgtggcaggtgtaaaagttgtagatttgttccagtgccctggttgtaaaagtg
gcatgctcaatctaaagggaagatgttacgagttttcaaatttgtgtaagagaatgatactaccatacacatcgactcattgttcttctctctttgat
gctaccataaatcgcgcagaggcttgtttttactccctagagtttctccagtatgattttgaaactgcgaggagaattgcgcatggagctaaaga
cattccccatgtttacaataaggtagtaaagaatgtaaaagatttggatagactgtgtgccttgtactgttacaaatgcgtatctcctgtcgtatgt
gatgagccgaatgaaagtacggactatgaaatggtggatgtaactccccctctaattaatcttaccgagattgttgattcggaagagtatgatg
atggtcccggaaatcatatgtggccagcaaaatttacctgtaactttattgcgggttccagtggagaaacacccaccattagtacgtgcagag
atgctgtaacttttctaggaagagcaccgaggaagaaaatggcaggatgggatgatcaatcggcagtggggcaagccattatagcgctag
ccaactggagaaagagtggggaattgcccaaaaatatgtttgatttactagaaggggtaaatgccgtactttatagaggcgacagtttcttgtt
acgtgcgataaactacccctgtgttattggtagatccatgagccctagtctggaactcgttaaaagaaaggtgaataaaattgctttaataaag
gccttcttccacgagaaaagggtgcgtccagacgcatctaaaaagttacttgaatgggcagaactattagtcaaaagttatctcatggaagttt
tacttcagacgccagaatgtgtcatacaccgcgcccattcatttgtaggcaaaactctcctcattactgacgaattggttcacatgcgtccagat
gatgccacaagaaacgcctatatccagaacctaaatgcggccagacagaatgcggctgctgcagcctcattttctgggtcgctccccaaac
ctgaatttgtcccctgcaaagaaaggacgattgaatggatgtatgaaaaggacaatgatgatgttagagttgtaaattgtccttcatgtaaaaag
gctatccagaaatatggaggttgtgtgaatgtgttttgtgaatgtggaacaaacatgtgctggatatgtgaagagaaggtttctcctgctgattct
aatcattgtgtggagaaacacaggattgtttatagtaactgtgttagggttaaatatgccttagaaagtatgtacgggtttgagatttgtaccatga
aaaatgtagaagaaggagttaaaaattattatgtaatggagaatggatttttctttgatgtacaagaaatggttgctaagaaataataaaacactt
gtgaaatgttataacttgttttattgtccattatattcaacaaactttaatttctttgcatgccttgttcaaaagggcaacataatcgccttctttctcctc
cctctttctctttgttcctccactgcttgtggctacttcttcttcttccccacgttcctcgttgtacgtctcaattgccttaactgtgttgaaaatcatgc
gacgctctcggcgcagacgcttcctttcctcctctggtaatgtttcttcctttttttattatgtccttgccttctttagaagaggggtcaatcacttttatt
tctacaccatcttgatatacttttccattatccatgaagattttgatgtctgatccagtaacctctctgaccatctccatcaacaacctttcattcgga
ggatcgatgaatgatatttcattaccctttccggatgcaatgaaatggttattaaaatggtttcgtatactatctctgatgcatataggtgcatctga
gccagttatggacatgaccaatctgggcctacgttcatccacgtccctctttctgtagtcgtccgaagagacgtatctgattccttcttctgtggc
agagtaaaaggaaccatttccattggttctgcacaacaaggagaaatctcggtcagaaaaggcaaagaccagacaggcaccgccgtcata
tctggatgatgcaattgaggactttttatgcacatctcgaagagaaatcctttgttcttctgaggcagcatatagctttttggattctttaagtttttctt
ccagaacacttctttcgctaactagaaggttgacttcatttttttgcaaatctgatttccttttcgaaattatctaatttttgattactcttttctagcgtgta
tacaactgcagcattgaaaaggcatgacatgtaatttccaatgcttttagctgacgggtgtgtgctgttgtagaaaagtaatccaagtttatccat
tggtacaattaggcacatatttgaaaaactgccaactatggggtttgcgatacttaagccattgtacgttaacctatcacagatatcatacactga
tacacgagaagaggtaccaccgtgttgcactaacacttgttccatattcttaagagtacgggtacaatttctaggagaaaacccacaccctact
aagaagtctgttgtgattatcatataatcacatttcttaacggccgcatttgaaagacgttcatgtatgattctgtataggggtgaaaatttaaagtt
cttgtcattaactagagccttttcttccacaattttgttcatttcctggaatgggcttggtagtgattcgttccatttttttccataattagggtcttcatc
ttctaagaaactaccttgttttgggagactggatgtgtattttagtccaatctgaagcgtaggatctcctgttaattgtctcaatctttcgttaccgtt
cttgatgtggagtacttccttgtctctaagacgaactataaaatgtccttcctttagctcacttctcctgacactcttgactgatttgaatgttaatagt
acccgttcataattttcggtattttccccagacaaacaaataccatttttttacttttaattgtttaaagtcttccatttttataccgtgtaaccctaacaat
ggtgtgccagtataataaatctacggaccatgctcttgatctgggaagtgtactcgtgatttctgtagaaattgctatttcctattcattatttccatt
acaggaaaagatgataaaatgaagacgtgcgatgagatatgtgtttttcaatatcgtatatcaaatttgggcaaattacatattttttagcgtcaa
cactcatcttcatgcaagactttttggagtgaaattactaaaaaaggctatatttcatgtacaacataaaacacatgtgcagacattaaattggttt
gctcaccccccccaacatcgttttcatcaactctttcctcacctctatataaaaccagcctgcaccccctcggaacccacagtcgacttgtaccct

ccgtcgagatgacatctccagctccatcaccctcttccacccccaaatccagttgtaccactattgtaaaccgatgtggtttcctccttgacaac
aacaaggaagtggtcatctacgacaccaattccaaattcaagtgtgaacccaaaaatctggaactaattggtgtactttctggagtctctgata
atgttgttacccagatatcccccgaccagatatttgtgggaacatatatggtcaaatataactggtctaaatctggtcatgaacgcttcagtgac
atgagtaacaactgtctggacaatattacacgcccttcagaagtgattgaaagtgtgataaagaaaacgtccagcgactttaaaatgaagtac
acacgttccttgatggaccacaccgagaaatactatttttctggtgaccaaaaattgagcaaaattagtagttggtgtacaacccctataacgac
agtgggtatgcaactccgtctagaaaactttgtcaaggaagaacatgaaactgtcgtggtgcacaacccttctggaatgactggattcaacat
atttaatagttcccccgtgtattttgaagtgcacaatgagatggacgccctaatttttatggcggctttcttgaagcacaatagtttatggggaga
aattaacgccaatatggacttgtacacgtttgattatgcgggtgcttttctggacgaaagatggtgccaccacgagaagagtttttctgtcgtcc
gagcacaacttatcaactcgtattacaagtgcaggagaaaaatcatgcaagccctggacaataactacaacaacaagaataagaagagga
agaatgttggtggagcacctgcgttcacatttatgagcggggacggagagggaggaaaggaagccctagaagctagtttcgatgtgattgg
gggaacaagaggaggaagatttggtgttgattcaacaccatgcccccattcttcagccatgcaactaaaactggacaatgaaggaaactatg
gatgtattgcctgcttcgcatcaatgttctttgtattggagaacccaggtgatgaatcttccttcatatcaacggatgcctctaaaattggacaag
cgcaagcatggatagatgaacgactacgaaacaatgaaaatggaggagaagaaaataatgtctttaaaaagaccttccatatgctggctgat
attacccaaaaggctcatgaaactgcctattccaataccatcccacttggacccaatggcaggcagtggaattggcctactcacactgtggaa
cctattgcccatgaatttgttacccattctctagtaaacacattgaaaaatctaggggatagaaaacttccccgattcaattttgatatcttgtacaa
cttgcttaatccatttggaaaaatgttgctagtgtttattcaaaattgtcacattttaactggacataaaaacaatgaaaatgtggtgcctcgaggtt
ctgcttctgggaagtggtggactattaattttgtgggtgtgaacatgtggacttttcaagtaacaaaatgtaaagttgaaaaggatagaaaaata
tccgatttggcctgtatggaaactctccctcgtctacctaatccaggaagcactaccgtcgatgacagaatagtttttaagggattctgtagagg
ggaaaatctagggagtgtaggtgaagtcgtatccgacattacacagagtgtcaagaatttttgtctcatggttgaaaataggaaatttagtgtg
gataaagaaactggtttcatctcttcagaatcgatagtgtctgatcccttcttttcactagaagtgactggctgtagatctaatcgtgcccaagata
ctattaataatggccgagttagtgctcgtgtaatgaggatcctaaagtcacgtgaaggtgctcgtgtatggttggccaaggatgaaaatgccat
catctttgaaaacgttaaccacgatacggccatctctacggacgctatggagcgagctatagggcagcacaagatactgtactatgatattga
aacaacagataaagatttcaccgacaaaaaaatcagtcatcacatctattgggttctgtttgtgtacgggaggcgatatgacacatggaggaga
gagaggagtatttggactggttgcacctggatccgacgtggaaaaggtgaaagagactataataaattcgtacgatcctgaagaaaaggaa
gacattatgaaacagtgccctcaagtgattgaaattttcaccaacgagtttgaaatgttgcttggttttggaaagtatatagataaagtgaagcct
cacgtgattagtgggtggaacaatgtagcttttgacgacccctttgtctttactcgtatcgtcaaacatttgagtgatcacaccaaagacatgtct
tattgtgtagcagatgcatctacagcagaatctgtccttcctagagcaacagaaggaggaggaggagaaactccatatagattgagcacccc
tcaagaaagaatacaactagcaagcactggtattttcaataaattgggaaaatttgtagacaagaaaactggcatgttgaaacctgaaatgact
gcagatttattggccggggcagaaagtcaggccaataccaagtttaaggaacgcaacaagttatcctccagtaataaaggatcagcaggat
ggttccagaaaattattggcggtatgtgcagtgctattcggttggatctcatgaaagtgtgcgaaaaggcctataaagaatccctctctgaattt
aatttgaacgccgtgctcgccaaagtgagtagtgtcggcgacaaggttaaaaatgtaaaagatgaagtagacctacactttcatctattggga
ttcttgaagctgaagaaggcccaggatcaggcaaaagtacacgtctattgttgcaaggatgcctacttgactggtatagtttctacctccatca
acaaggaaggggagatttttaggctgtgtatggactctgctttaaccgaggcggtcgtgacagccaacctggccactcctctatgtatagga
gaaggagcaatctgtagaaatatgggagaagaaagggcagatagaagaggtgtgggagtaagaagacactctattgccacagacacaaa
gggaggtatggtgagtcaacctatcgtcaatcatgttccctatcaaacgattgacatgacaagtttgtacccgatgaccatgtgtcagaataat
ctgtgcaccactacctttgtgacccatcgacaaattatgcaactgagggatagattggtacttgaaaaaatgaaaaacaaaaccaccgactctt
tattgttgttggacgttattgacgagtgcaatcagattgtgttgtccgagtacagacccattgatattgcagtcgcatcatggaagaatagcaact
ctaatagacaaactccaattactcgcatagaggaaagtttgggtctaagattcatagaaaatttggatgccgagaagacaaataataaaacgt
ggtgcaccaatacatctcccaatatgaatgtcactgccgcaggtatggattacttccccgagattgtgtgtgacattaatatgcagtttgcggcc
aaggtgaatgatgatatgcatatagccccagcaagtttagagtatatgcttcaagtattgcccataatgttaatcgacagaccgtacattggcg
cacacataacagctggaaaatgtcgtacattggaggatattctctcagaactcgaaaaggacttttctgttgaaaaagatgaggaaattataag
aacccattggacatttaagggtcaaaaacaatacgatttctgtcatagtcctgtgacccaaatggctcgtcacattattgaatctacgggaaga
aatatccgtgattatgaaggcaatgaaaaattcgagaggttggttagcttatcagacagaatttatcgccgcgttggcgcatttgattcggcca

atgatccagctgttagactgtggtcttctcgtctaatcaatgttggaatgttggttaggacatggaacgtaaaaactgacattcttaagggaatca
tccctcaaatgcaagccacttacagagccgatcgagttgtgatgcagaacaaggccaaggagtttgccaagatgggagacatgaaaagag
ctggtctaaacaaagttggacaaaatattatgaagctcggtatgaattccatgtatggtcatttggccctaagagcacgttcgagccgtaaaga
gtttgcgtctggatctgccaatactgcctcaagtatttccaacatgtcagccaccggaggaattggaggaggcacaaggcactcggtgacg
gccaatcagattacagaaaatgctcgatgtgtatttggcaatattggttgtggattacagatggctcttcctggtactaagcagacgtacgggg
atacagattctgtattctgcgtgcataatattgtaggtgatggaggaatgataccagaatatgatgaacaaactggcaaatattattatgtgatgg
atattgctctaaaaaataaaatggctgcaattattcccatcctagtcaactcgttaacaaagggcatccagtttgtagagcgccgagacgctgg
tgtgggcatgatgaatatcgcccatgaacgtctagctgtcgctggtcttttgtttgccaagaaaacataccatatgcttcactttaatgaaaatag
tgcagcgttcaatgacatgataaaattgaaatcaaccgataacaataataagtttgcatcctttatcaagagacctagccacgcagatgggtat
gttgttccccataatccttcattgattcttagagcggccgaaggacccgctggtaagaaattgaaaagctttttggaagaggaaggaattcatg
acgagaagagtatggaggaatggtttacctcttcacctacgtggatggccatggatgcttctgttatcaacaacttgtatgcctcacaaattgta
ggggtggagaagggtaactggattgacgccatgacttcccgccccatagaagcgggtacagaaatgatggaggcggtgacgcaagcga
atgcagctttcaccccttacaaaaagggagcctttgtgaagaagggaattacacccaccaccaaactaaagggtctccaatcattgattgcaa
gatttttaccaaaaatagaggaaaagaaaagctgttatttggatgtgatgaagaatcatgtggagaattttgcatcccatataacaaatcctgct
atgatgattactagttcccgagtcaacaagtttgatacgtcaaaagaacagagtagacctaatcctctagctctagcgataaataaccacctga
acccttcttcagaaatttcattggggcagaaatttaagacggtgacatcagtttcttcttggagtctttcggcagaggaaggggaagtccctgct
ggttattttaacgctggtagcgtgcgttgggatgccaccaacatgaagggaagtgttcctgcattttcagtcaagaatttatctgttgtgcccaa
cgccatcacatctgtatacaagatggttgagagcgataagacggcaataaaatccatgattgctaaaaatgtagaagtgttgtgttctacatct
gccaatactggattttctctgagaagaggagcattgtcatttaatacaggcgtcattgttacaaaggacgtggctatggcttgtatacgatctcta
aataataaacaaatgttattgtttgttggagggggaaaggattacggtgaagacgacgacgacgacgacgaagaagcagaagaagagga
cgaagaaaatggtgaaaacgaagagaacaaaggtgactgtgtcacggaaaaaaagatccctggacgaagcactaacaaggatgttggtg
aagaaactaaaacaagcgagaaaacggagggagaaagaaagggctctaagacggcaaagggaaagacggaggaaattgctagttcgtt
gagtaaatgtgggaagaaagatgcgagagatgtcattcttgaccgtttactaaaagcaacacattcttcttgcaccaacaatgaagagagaac
cagagtcttacaacaatatagcaattgtacattatcttcctatataacttcagtcatgaaattggaccaaagagtagcagaccaaatggaaaattt
aatatctcaattggatcaaatacgtaatctctccaacaagaagaggcaagaaaagggagggccttttaagtctgaattggacgccatggttgc
tgcagttaaggttaagtttttcccagttttagatgcgtctagaaaattgactcaagaccattggaaaaagtgccccgtgtccatcccagaaacgc
gtgaagaaaaaccattaatgggtgtgccttttgaagttgcactcaattctctaataggaaaacacaagtgcacagatacatgcgacatggcttg
ttgtcaatcattgtattttgtcctcttgtacacgctagctttaaaatttgagaacgaaagattggcccggcaaattggcctagatgactctgtagatt
tgatggctgagatgttgttcggaggggataaactattggcccaggaagtgttaaaaagggtaaaagatgctcaagatagaaagttggtgaaa
tctttattgcctttaaattataaccatgacacaaatacaattatatttttgtttgagtctttaaggtttgctcagaaacctgtagctggtatgagtgttag
tgaaataaaagacgctgttagaggtctggccttttctaccactacaggtactgtgtggaattatactgatgaaagatttttggaccattgtataac
atggatgaactttgtaacgaacgtgtcaatggaaattgtaaattgtcctttataactggtatttatcatacggcagcagtagaattggctgctgcat
gtctatcttgtgttttgtaagaaatgataataaaaaacatgtataatttcattctggtttttattttatatcctcaagagttgtagtagtagcagcagca
gcagcaatagcagcattattttcagcgacagcttcttcatttctattgattgatatctggtcattttctacagcccatggtcgtttaaggaatattcttt
gaagcagaaacacatacccagcaatgattgcaatgatcaatattaatagtagcgaattaattaaaataagcattttgtccacattctggctaaaa
taaaaaagtttatctctcaatgaaaaatttttattgaccaatcacttcgggaagttttgcagcatgacacactattactccaaaactatgctcactc
gtatctagtcccacaacatcacgcgtcatatttaaaacatgcacctttaaagcctgtttttcttatacaattctgtattgattatagggtttacaacct
ttaggccttcctttagagatactattttcatgtattttacaatcgaatatgagttgtcttgtgcaccagacgggagcctgtcatgaaactctacttctc
tcccacgtgtattgagcgagagatcattaagcacaacttctgaagaaacgagaggttgtaaaatagagttatatttaggtttaattaggaaacat
ttgtggaaacttgaactgttaatatcaatagaaactgagctatatgctgacgacatggcttgcattttcaaatcatgttcaaatcttactacataac
ccaagaaattataatctgcttcttcaccatcatcttcttccatggcatcctctcctcttggctcttttagtacaatttcttttggttggttagtcatgttgt
atatgttcattgagctatgatttcccacaacatcatcaaagaaaaccccttctgggataatgcatgcaccgacagaggtgtagaataatcctcc
atatctaatgacaccgactgcggtagcgttctcttcatcatagtaaccgagcttgattttcacatgactcatgggtccgatagagatattggaag

ccggttcagtagcggaaaaaatagacacttgttggaacattttgtcgtcaggaagagtctcgaataaaataccgacaggggaattatcttcaa
cactattaaagtttgttgcataagatagccgccctgaactacttgtattataaattgttatatcggatgctatttcagaacaaagatctcctataaaa
ttgtgattagggcagaccaacatcccgtctccagtgtaagacgatacatgaagtcctggtccagttggcacgaaaatggcagccgttcgtga
aggatttctcacgtcaactcgtctcaccttcactggtacttttccaaatcgtcgtggatctacattgaaactaaaatctctttcgtggggatcacttc
catcctttaaagagaagagaggcactctgtaaacaggagaccaatctgaatagtaccttaaagctgttctattttttcttctagagtgggtaccg
ctctgcggtgcattcggtacacctccaaatccgtttcctttacttgaatcgattttgctccttcattttcccagagcgcatcagaatcgtacctgtcg
tcgaacgaatcttcacctcccattttatttattttcctctatatgataaaaaccacacctaaaatgtctaatattttactacactgaaaaa

**Table S2**. Composition of LAMP primer evaluation reactions, Cas sgRNA evaluation reactions, one-pot SHERLOCK reactions, 10X LAMP primer mix, and sample dilutions used in one-pot SHERLOCK reactions.

**a)** Composition of LAMP primer evaluation reactions.

| LAMP Reaction component | Initial Concentration | Final Concentration | WSSV reaction volume (µL) | TSV reaction volume (µL) |
|---|---|---|---|---|
| Isothermal Amp Buffer (NEB) | 10x | 1x | 2 | 2 |
| MgSO4 (NEB) | 100 mM | 8 mM | 1.6 | 1.6 |
| glycine | 2 M | 200 mM | 2 | |
| taurine | 500 mM | 50 mM | | 2 |
| water | | | 4.3 | 5.8 |
| dNTPs (NEB) | 10 mM | 1.4 mM | 2.8 | 2.8 |
| WarmStart RTx Reverse Transcriptase (NEB) | 15000 units/mL | 300 units/mL | | 0.4 |
| WarmStart Bst 2.0 (NEB) | 8000 units/mL | 320 units/mL | 0.8 | 0.8 |
| Primer mix | 10x | 1x | 2 | 2 |
| Template | | | 4 | 2 |
| SYTO-82 LAMP dye (Thermo) | 100 µM | 0.5 µM (1 µM for TSV) | 0.1 | 0.2 |
| 5’ FAM-polyT(5) reporter (IDT) | 100 µM | 2 µM | 0.4 | 0.4 |
| **Total** | | | 20 | 20 |

**b)** Composition of Cas sgRNA evaluation reactions.

| Cas/gRNA evaluation reaction component | Initial Concentration | Final Concentration | WSSV reaction volume (µL) | TSV reaction volume (µL) |
|---|---|---|---|---|
| NE Buffer 2.1 (NEB) | 10x | 1x | 1 | 0 |
| Isothermal Amp Buffer (NEB) | 10x | 1x | 0 | 2 |
| MgSO4 (NEB) | 100 mM | 8 mM | 0 | 1.6 |
| Cas12b (GenScript) | 8.73 µM | 200 nM (250 nM for TSV) | 0.229 | 0.570 |
| sgRNA | 5 µM (2.5 µM for TSV) | 400 nM (250 nM for TSV) | 0.8 | 2 |
| water | | | 4.571 | 7.83 |
| Template | | | 3 | 5 |
| 5' HEX-polyT(20) reporter (IDT) | 5 µM | 200 nM | 0.4 | 0 |
| 5' FAM-polyT(5) reporter (IDT) | 50 µM | 2.5 µM | 0 | 1 |
| **Total** | | | 10 | 20 |

**c)** Composition of one-pot SHERLOCK reactions.

| One-Pot SHERLOCK Reaction component | Initial Concentration | Final Concentration | WSSV reaction volume (µL) | TSV reaction volume (µL) |
|---|---|---|---|---|
| Isothermal Amp Buffer | 10x | 1x | 2 | 2 |
| MgSO4 | 100 mM | 8 mM | 1.6 | 1.6 |
| glycine | 2 M | 200 mM | 2 | |
| taurine | 500 mM | 50 mM | | 2 |
| Cas12b | 8.73 µM | 200 nM (150 nM for TSV) | 0.458 | 0.344 |
| sgRNA | 10 µM | 200 nM (600 nM for TSV) | 0.4 | 1.2 |
| water | | | 3.44 | 2.46 |
| dNTPs | 10 mM | 1.4 mM | 2.8 | 2.8 |
| **Incubate for 15 minutes at room temperature then add…** | | | | |
| WarmStart RTx Reverse Transcriptase | 15000 units/mL | 150 units/mL | | 0.2 |
| WarmStart Bst 2.0 | 8000 units/mL | 320 units/mL | 0.8 | 0.8 |
| Primer mix | 10x (20x for TSV) | 1x | 2 | 1 |

| | | | | |
|---|---|---|---|---|
| DNA/RNA | | | 4 | 5 |
| SYTO-82 LAMP dye | 100 µM | 0.5 µM (1 µM for TSV) | 0.1 | 0.2 |
| 5' FAM-polyT(5) reporter | 100 µM | 2 µM | 0.4 | 0.4 |
| Total | | | 20 | 20 |

**d)** Composition of 10X primer mix.

| 10x Primer Mix component (100 µM) | 10x concentration | Reaction Concentration | Volume (µL) |
|---|---|---|---|
| F3 | 2 µM | 0.2 µM | 2 |
| B3 | 2 µM | 0.2 µM | 2 |
| BIP | 16 µM | 1.6 µM | 16 |
| FIP | 16 µM | 1.6 µM | 16 |
| LF | 4 µM | 0.4 µM | 4 |
| LB | 4 µM | 0.4 µM | 4 |
| Water | | | 56 |
| Total | | | 100 |

**e)** Composition of sample dilutions used in SHERLOCK reactions.

| Sample dilution component | WSSV | TSV |
|---|---|---|
| Sample DNA/RNA | 1ng | 10 ng |
| Pathogen free shrimp nucleic acid (DNA for WSSV, RNA for TSV) | 20 ng | 5 ng |
| water | variable | variable |
| Total volume/one-pot reaction | 4 µL | 5 µL |

**Table S3.** Viral concentrations estimated by qPCR, SHERLOCKv1, and SHERLOCKv2 assays.

| Target | Sample Name | qPCR predicted copies/uL (Mean) | qPCR predicted copies/uL (Std Dev) | SHERLOCKv1 predicted copies/uL (mean) | SHERLOCKv1 predicted copies/uL (Std Dev) | SHERLOCKv2 predicted copies/uL (Mean) | SHERLOCKv2 predicted copies/uL (Std Dev) |
|---|---|---|---|---|---|---|---|
| WSSV | Pvan_001 | 1261101.25 | 48246.34 | 1776906.337 | 625688.6194 | 5467145.199 | 1359933.112 |
| WSSV | Pvan_002 | 1718498 | 50077.57 | 1669218.927 | 545295.3689 | 4609541.442 | 1110242.961 |
| WSSV | Pvan_003 | 14366123 | 4244339 | 9001832.705 | 7178704.302 | 96679789.06 | 6755260.046 |
| WSSV | Pvan_004 | 813641.19 | 83353.48 | 349363.9842 | 193391.5695 | 1780740.143 | 525445.2165 |
| WSSV | Pvan_005 | 14909322 | 917087.06 | 11799814.91 | 6901524.777 | 77038795.42 | 9319605.555 |
| WSSV | Pvan_006 | 245472.52 | 10655.48 | 329394.7243 | 50657.36744 | 1162038.4 | 87541.39027 |
| WSSV | Pvan_007 | 1989807 | 60248.59 | 2411408.409 | 601446.7782 | 8350530.544 | 3014319.528 |
| WSSV | Pvan_008 | 8500098 | 1241473.75 | 8461070.295 | 9651892.371 | 46890709.6 | 4588834.125 |
| WSSV | Pvan_009 | 20427184 | 2885616.5 | 14534990.17 | 5324836.213 | 58414315.31 | 14373889.11 |

| | | | | | | | |
|---|---|---|---|---|---|---|---|
| WSSV | Pvan_009 | 20427184 | 2885616.5 | 14534990.17 | 5324836.213 | 58414315.31 | 14373889.11 |
| WSSV | Pvan_010 | 34825116 | 5418532.5 | 33151839.4 | 56799437.01 | 1292036201 | 208554040.5 |
| TSV | Val.1 | 9448820.368 | 29459.50244 | 0 | 0 | 3666974.695 | 1240577.439 |
| TSV | Val.10 | 4251.754357 | 252.5155991 | 0 | 0 | 246.50393 | 162.6205235 |
| TSV | Val.12 | 20.01454529 | 6.875195902 | 0 | 0 | NA | NA |
| TSV | Val.13 | 793198.6619 | 26800.87853 | 0 | 0 | 83022.78106 | 84986.27011 |
| TSV | Val.14 | 4779.881361 | 1094.038964 | 0 | 0 | 0.836411494 | 0.853683394 |
| TSV | Val.15 | 19474.14741 | 624.4697417 | 0 | 0 | 520.3408561 | 639.3912582 |
| TSV | Val.16 | 250289.7597 | 9312.733832 | 0 | 0 | 225158.857 | 135312.568 |
| TSV | Val.18 | 8488.771518 | 1668.779531 | 0 | 0 | 1435.485604 | 893.2158789 |
| TSV | Val.19 | 12984.57389 | 244.4843919 | 0 | 0 | 1403.516582 | 716.6397924 |
| TSV | Val.2 | 647.4646646 | 84.41761308 | 0 | 0 | NA | NA |
| TSV | Val.20 | 4244.294329 | 1342.586357 | 0 | 0 | 469.9003654 | 248.8232393 |
| TSV | Val.21 | 96172719.47 | 5849525.621 | 0 | 0 | 16669019.45 | 17751739.13 |
| TSV | Val.22 | 33075.42455 | 1023.116759 | 0 | 0 | 283.739625 | 308.0032653 |
| TSV | Val.23 | 5701.93041 | 297.856471 | 0 | 0 | 522.6309481 | 171.5829891 |
| TSV | Val.25 | 864.5498815 | 98.85942432 | 0 | 0 | NA | NA |
| TSV | Val.26 | 5214504.385 | 117015.3335 | 0 | 0 | 1168056.433 | 1211323.704 |
| TSV | Val.28 | 6021.620089 | 144.5154752 | 0 | 0 | 6.070351404 | 4.024323014 |
| TSV | Val.29 | 82264608.33 | 3938815.797 | 0 | 0 | 816313.871 | 232529.5993 |
| TSV | Val.30 | 18316.14497 | 82.72721706 | 0 | 0 | 139.8481296 | 84.16646448 |
| TSV | Val.31 | 5549.223316 | 295.3453823 | 0 | 0 | 227.937411 | 117.4144615 |
| TSV | Val.32 | 104.4052704 | 5.120732148 | 0 | 0 | 0.001045791 | 0.00098074 |
| TSV | Val.33 | 5149.949616 | 526.0034592 | 0 | 0 | NA | NA |
| TSV | Val.34 | 3735.120502 | 312.3589272 | 0 | 0 | 445.5397799 | 241.5433639 |
| TSV | Val.35 | 30803660.68 | 1027489.064 | 0 | 0 | 6239838.009 | 7670933.831 |
| TSV | Val.4 | 2039523802 | 458395462.7 | 0 | 0 | 171523702.7 | 52581346.83 |
| TSV | Val.5 | 9906995.501 | 376073.0159 | 0 | 0 | 1131242.4 | 154845.6221 |
| TSV | Val.6 | 29425.11971 | 1009.758788 | 0 | 0 | 4000.363056 | 3286.554796 |
| TSV | Val.7 | 2339.799066 | 251.7950218 | 0 | 0 | 115.0096636 | 79.80866549 |
| TSV | Val.8 | 945.3381209 | 184.3990381 | 0 | 0 | 42.00158029 | 38.60886497 |
| TSV | Val.9 | 9985.187813 | 458.6134716 | 0 | 0 | 11.98072859 | 12.88052937 |
| WSSV | Pvan_011 | 755.29 | 133.87 | 2281.542796 | 492.227141 | 1354.997249 | 1868.550003 |
| WSSV | Pvan_020 | 285116.16 | 13440.42 | 44.09142332 | 15.16275833 | 7085.491244 | 2078.220615 |
| WSSV | Pvan_021 | 26514780 | 481846.59 | 12782848.25 | 13237136.16 | 62602087.93 | 13390504.59 |
| WSSV | Pvan_022 | 2378394.5 | 221400.8 | 13226061.55 | 26445331.41 | 140109.3994 | 89870.25916 |
| WSSV | Pvan_023 | 5684949 | 877728.44 | 10942413.08 | 21520314.2 | 89051.86532 | 45108.3077 |
| WSSV | Pvan_023 | 5684949 | 877728.44 | 10942413.08 | 21520314.2 | 89051.86532 | 45108.3077 |
| WSSV | Pvan_024 | 662213.5 | 104129.52 | 629611.5187 | 1090265.603 | 365.9336519 | 442.7075628 |
| WSSV | Pvan_025 | 714.86 | 130.02 | 9502.144932 | 18919.57272 | 531.6808182 | 461.9524433 |
| WSSV | Pvan_027 | 471.29 | 21.12 | 8036.450868 | 15629.78019 | 4571.293956 | 6643.848727 |

| | | | | | | | |
|---|---|---|---|---|---|---|---|
| WSSV | Pvan_028 | 25092454 | 711439.94 | 40332523.81 | 69845123.78 | 30000013.14 | 12499625.04 |
| WSSV | Pvan_029 | 38.11 | 0.71 | 34.39503838 | 1.733003669 | NA | NA |
| WSSV | Pvan_012 | 825.26 | 87.39 | 1709.071993 | 788.8357409 | NA | NA |
| WSSV | Pvan_030 | 401.69 | 127.6 | 740.9000416 | 1216.460549 | 1842.406105 | 387.4690796 |
| WSSV | Pvan_031 | 51570296 | 10850528 | 54938270.64 | 56418505.53 | 63016260.34 | 20629485.5 |
| WSSV | Pvan_032 | 327981.19 | 38229.46 | 135315.522 | 161675.6649 | 132525.1577 | 46161.97828 |
| WSSV | Pvan_033 | 45211092 | 13916455 | 22713145.35 | 15334499.78 | 152120367.3 | 40322972.92 |
| WSSV | Pvan_034 | 2560871.5 | 455237.47 | 2179494.771 | 3745570.388 | 17598.10636 | 10033.61246 |
| WSSV | Pvan_035 | 6992533 | 879068.81 | 5778090.783 | 9373828.135 | 13863.52793 | 6544.163593 |
| WSSV | Pvan_035 | 6992533 | 879068.81 | 5778090.783 | 9373828.135 | 13863.52793 | 6544.163593 |
| WSSV | Pvan_013 | 984.16 | 521.7 | 2258.146377 | 840.0177409 | 738.1652517 | 544.7067557 |
| WSSV | Pvan_014 | 352.03 | 407.36 | 570.3737806 | 185.8082441 | 169.5386181 | 168.2183909 |
| WSSV | Pvan_015 | 2769921 | 613732.06 | 2963397.817 | 185310.9457 | 56621.76724 | 30254.24571 |
| WSSV | Pvan_015 | 2769921 | 613732.06 | 2963397.817 | 185310.9457 | 56621.76724 | 30254.24571 |
| WSSV | Pvan_016 | 3145114.25 | 296301.78 | 3008043.672 | 1440982.249 | 25501.11722 | 23491.10288 |
| WSSV | Pvan_016 | 3145114.25 | 296301.78 | 3008043.672 | 1440982.249 | 25501.11722 | 23491.10288 |
| WSSV | Pvan_017 | 7378686.5 | 467776.59 | 3945336.838 | 311435.2825 | 325062.9864 | 374691.1292 |
| WSSV | Pvan_017 | 7378686.5 | 467776.59 | 3945336.838 | 311435.2825 | 325062.9864 | 374691.1292 |
| WSSV | Pvan_018 | 5524195 | 261646.83 | 12106948.81 | 24213010.04 | 51171.2666 | 57834.1861 |
| WSSV | Pvan_018 | 5524195 | 261646.83 | 12106948.81 | 24213010.04 | 51171.2666 | 57834.1861 |
| WSSV | Pvan_019 | 58975944 | 12307867 | 61274590.18 | 120998437.4 | 32490015.21 | 9313367.484 |
